# Supporting materials: Endothelial cells differentiated from patient dermal fibroblast-derived induced pluripotent stem cells resemble vascular malformations of Port Wine Birthmark

**DOI:** 10.1101/2023.07.02.547408

**Authors:** Vi Nguyen, Chao Gao, Marcelo L Hochman, Jacob Kravitz, Elliott H Chen, Harold I Friedman, Camilla F Wenceslau, Dongbao Chen, Yunguan Wang, J Stuart Nelson, Anil G. Jegga, Wenbin Tan

## Abstract

**Background:** Port wine birthmark (PWB) is a congenital vascular malformation resulting from developmentally defective endothelial cells (ECs). Developing clinically relevant disease models for PWB studies is currently an unmet need.

**Objective:** Our study aims to generate PWB-derived induced pluripotent stem cells (iPSCs) and iPSC-derived ECs that preserve disease-related phenotypes.

**Methods:** PWB iPSCs were generated by reprogramming lesional dermal fibroblasts and differentiated into ECs. RNA-seq was performed to identify differentially expressed genes (DEGs) and enriched pathways. The functional phenotypes of iPSC-derived ECs were characterized by capillary-like structure (CLS) formation *in vitro* and Geltrex plug-in assay *in vivo*.

**Results:** Human PWB and control iPSC lines were generated through reprogramming of dermal fibroblasts by introducing the “Yamanaka factors” (Oct3/4, Sox2, Klf4, c-Myc) into them; the iPSCs were successfully differentiated into ECs. These iPSCs and their derived ECs were validated by expression of a series of stem cell and EC biomarkers, respectively. PWB iPSC-derived ECs showed impaired CLS *in vitro* with larger perimeters and thicker branches as compared to control iPSC-derived ECs. In the plug-in assay, perfused human vasculature formed by PWB iPSC- derived ECs showed bigger perimeters and greater densities than those formed by control iPSC-derived ECs in severe combined immune deficient (SCID) mice. The transcriptome analysis showed that dysregulated pathways of stem cell differentiation, Hippo, Wnt, and focal adhesion persisted through differentiation of PWB iPSCs to ECs. Functional enrichment analysis showed that Hippo and Wnt pathway-related PWB DEGs are enriched for vasculature development, tube morphology, endothelium development, and EC differentiation. Further, members of the zinc finger (ZNF) gene family were overrepresented among the DEGs in PWB iPSCs. ZNF DEGs confer significant functions in transcriptional regulation, chromatin remodeling, protein ubiquitination, and retinoic acid receptor signaling. Furthermore, NF-kappa B, TNF, MAPK, and cholesterol metabolism pathways were dysregulated in PWB ECs as readouts of impaired differentiation.

**Conclusions:** PWB iPSC-derived ECs render a novel and clinically-relevant disease model by retaining pathological phenotypes. Our data demonstrate multiple pathways, such as Hippo and Wnt, NF-kappa B, TNF, MAPK, and cholesterol metabolism, are dysregulated, which may contribute to the development of differentiation-defective ECs in PWB.

**Bulleted statements:** *What is already known about this topic?:*

- Port Wine Birthmark (PWB) is a congenital vascular malformation with an incidence rate of 0.1 – 0.3 % per live births.
- PWB results from developmental defects in the dermal vasculature; PWB endothelial cells (ECs) have differentiational impairments.
- Pulse dye laser (PDL) is currently the preferred treatment for PWB; unfortunately, the efficacy of PDL treatment of PWB has not improved over the past three decades.

*What does this study add?:*

- Induced pluripotent stem cells (iPSCs) were generated from PWB skin fibroblasts and differentiated into ECs.
- PWB ECs recapitulated their pathological phenotypes such as forming enlarged blood vessels in vitro and in vivo.
- Hippo and Wnt pathways were dysregulated in PWB iPSCs and ECs.
- Zinc-finger family genes were overrepresented among the differentially expressed genes in PWB iPSCs.
- Dysregulated NF-kappa B, TNF, MAPK, and cholesterol metabolism pathways were enriched in PWB ECs.

*What is the translational message?:*

- Targeting Hippo and Wnt pathways and Zinc-finger family genes could restore the physiological differentiation of ECs.
- Targeting NF-kappa B, TNF, MAPK, and cholesterol metabolism pathways could mitigate the pathological progression of PWB.
- These mechanisms may lead to the development of paradigm-shifting therapeutic interventions for PWB.

## Background

Congenital vascular malformations (CVMs) are a major threat to public health with a 1.5% incidence rate in the general population.^1^ Congenital capillary vascular malformation, also known as port-wine birthmarks or stains (PWB or PWS), is the most common type of CVMs with an estimated prevalence of 0.1–0.3% per live births.^2^ PWB can exist alone or be associated with many other CVMs occurring in children, such as Sturge-Weber syndrome (SWS), Parkes-Weber syndrome, and arteriovenous malformations.^3^ About 25% of PWB in infants with forehead lesions present SWS, which results in an ipsilateral leptomeningeal angiomatosis syndrome.^4^ PWBs appear as flat red macules in childhood, which tend to progressively darken to purple at the patient ages. By middle age, PWB often become raised as a result of vascular nodule development and are susceptible to spontaneous bleeding or hemorrhage.^2^ Moreover, PWB is a disease with devastating lifelong psychological and social implications that greatly impair the quality of life of afflicted children during their development and growth.^2^

The vascular phenotypes of PWB lesions typically show the proliferation of ECs and smooth muscle cells (SMCs), replication of basement membranes, disruption of vascular barriers, and the progressive dilatation of vasculature.^5, 6^ Pathologically, PWB ECs exhibit stem-cell-like phenotypes, which are considered aberrant endothelial progenitor cells (EPCs),^5, 7^ leading to differentiation-defective ECs.^5^ We posit that those differentiation- defective PWB ECs are responsible for these clinical manifestations. Pulsed dye laser (PDL) is the treatment of choice for PWB. Unfortunately, complete removal of PWB occurs in less than 10% of patients.^8^ Approximately about 20% of lesions do not respond to laser treatment.^9^ Between 16 - 50% of patients experience re-darkening of their PWB as early as 5 years after having received multiple PDL treatments.^9^ There has been no improvement in clinical outcomes using laser-based modalities for PWB treatment over the past three decades.^10^ Differentiation-defective PWB ECs likely persist after laser treatments, resulting in the revascularization of PWB blood vessels after laser exposure.^5, 11^

One long-term obstacle to our understanding of the pathogenesis of PWB and a significant impediment to the development of improved approaches to treat PWB has been the lack of clinically-relevant cell and animal models. One strategy to overcome this barrier is to generate PWB patient-derived-induced pluripotent stem cells (iPSCs). IPSCs can be reprogrammed from somatic cells by introducing the “Yamanaka factors” (Oct3/4, Sox2, Klf4, c-Myc), bypassing the need for embryonic cells but retains a patient-matching phenotype.^12^ IPSCs are capable of propagating indefinitely and producing many lineages, such as ECs. In this study, we successfully generated PWB patient-derived iPSCs which were differentiated into clinically-relevant ECs. These iPSC-derived ECs recapitulated many pathological phenotypes of PWB, including formation of blood vessels with large diameters *in vitro* and *in vivo*. The dysregulated Hippo and Wnt pathways could contribute to the differential defects of ECs, thus leading to the development of PWB vascular lesions.

## Methods

### Tissue preparation

The study (#1853132) was approved by the Institutional Review Board at the Prisma Health Midlands. One surgically excised nodular PWB lesion from one patient (Caucasian male, 46 years old) and one age and gender-matched de-identified surgically discarded normal skin tissue were collected for this study. The donor had very large lesions with nodular and hypertrophic PWB on back, chest, arm, and hand and received multiple rounds of PDL treatments prior to surgical procedures. The skin biopsies were cut into a size of 5 x 5 mm^2^ pieces and cultured in a Dulbecco’s Modified Eagle’s Medium (DMEM) with 10% fetal bovine serum (FBS) for outgrowths of human dermal fibroblasts (hDFs). The hDFs were propagated and harvested for generating iPSCs.

### Generation of iPSCs and differentiation of iPSCs into ECs

The generation of iPSCs was performed using a CytoTune-iPS Sendai Reprogramming Kit (ThermoFisher, Waltham, MA). Briefly, hDFs were cultured in DMEM with 10% FBS to reach 80- 90% confluence. The Sendai viral (SenV) particles were used to deliver Yamanaka factors through three vectors, e.g., polycistronic Klf4–Oct3/4–Sox2, c-Myc, and Klf4, into hDFs using the manufacturer’ manual (ThermoFisher, Waltham, MA). The iPSC colonies were selected, isolated, and propagated under feeder-free conditions using Essential 8 Medium on a Geltrex-coated plate (ThermoFisher, Waltham, MA).

The protocol of the differentiation of iPSCs into ECs followed several reports,^13–16^ with tailored modifications. First, the iPSCs were induced to mesenchymal stem cells (MSCs) with a tendency towards ECs. Briefly, iPSCs were incubated in DMEM/F12 with human Activin A (Peprotech, 125 ng/mL), B27 without insulin (1X, ThermoFisher) and geltrex (1X, ThermoFisher) on day 1. On days 2 - 4, the cells were incubated in DMEM/F12 with human BMP4 (Peprotech, 30 ng/mL), human bFGF (Peprotech, 10 ng/mL), human VEGF_165_ (Peprotech, 50 ng/mL) and geltrex (1X).

The medium was changed every other day. Second, EC induction was carried out on days 5 - 7. The MSCs were incubated in StemPro complete medium (ThermoFisher, Waltham, MA) with VEGF_165_ (200 ng/mL), 8-Br-cAMP (0.2 μM), and forskolin (2 μM). The medium was changed every other day. On day 8, the ECs were passed using StemPro Accutase:Trypsin (3:1 ratio) and cultured in StemPro complete medium with VEGF (200 ng/mL) and 8-Br-cAMP (0.2 μM). On day 9, the medium was replaced with StemPro complete medium with VEGF (200 ng/mL), 8-Br-cAMP (0.2 μM), and bFGF (10 ng/mL). On day 11, the ECs were incubated in a regular EC growth medium (PromoCell, Heidelberg, Germany).

### RNA-sequencing (RNA-seq) and data analysis

Total RNAs from iPSCs, MSCs, and ECs were extracted using a Qiagen RNA isolation kit. The ribosomal RNAs were depleted using a Qiagen rRNA HMR kit. RNA-seq libraries were constructed using a Qiagen Standard RNA Library kit as instructed by the manufacture’s manual. The insertion size of the libraries was determined using an Agilent 2100 Bioanalyzer system (Santa Clara, CA, USA). Library quantification was carried out using a next generation sequencing (NGS) library quantification kit from Takara Bio. Paired-end sequencing was performed on a NovaSeq 6000 system (Illumina, San Diego, CA, USA) for 25-50 M reads per library. A standard pipeline of RNA-seq analysis was followed. The STAR (spliced transcripts alignment to a reference; https://github.com/alexdobin/STAR) RNA-seq aligner was used for raw reads’ alignment.^17^ Human genome fasta and gtf annotation files from ensemble (Homo_sapiens.GRCh38.dna.primary_assembly.fa.gz; Homo_sapiens.GRCh38.105.chr.gtf.gz) were retrieved for building the STAR genome index files; the raw reads (fastq files) were then inputted, aligned and mapped to human reference genome using STAR. The bam files generated by STAR were used as inputs to FeatureCounts (http://bioconductor.org/packages/Rsubread) for further validation and gene counts of RNA transcripts.^18^ Biostatistical analysis for differentially expressed genes (DEGs) among groups were carried out using edgeR (https://bioconductor.org/packages/edgeR/) that implements *exact* and *generalized linear models* (*glms*)-based statistical methods.^19^ A set of scaling factors (*effective library sizes*) were used for downstream analyses after normalization of the original library sizes. Differential expressions were determined using either quasi-likelihood (QL) F-test or likelihood ratio test by *glm* functions in edgeR. A false discovery rate (FDR) < 0.05 was considered significant. Gene ontology (GO) for enrichment analysis on gene sets and Kyoto Encyclopedia of Genes and Genomes (KEGG) for enrichment pathway analysis were performed using the *goana* and *kegga* functions in edgeR.

For further pathway and functional enrichment analysis by using GO and various pathway database gene annotations, we compiled a list of known Wnt and Hippo signaling pathway genes. In addition, for Hippo signaling, we used the Hippo pathway interaction proteome, which included 343 new protein interactions for human hippo components^20^. For Zinc finger gene set compilation, we used gene families from the HUGO (June 2027; HGNC Database, HUGO Gene Nomenclature Committee (HGNC), European Molecular Biology Laboratory, European Bioinformatics Institute (EMBL-EBI), Wellcome Genome Campus, Hinxton, Cambridge CB10 1SD, United Kingdom www.genenames.org). These compiled lists of genes (Supplementary File 1) were used to obtain the EC, iPSC, MSC DEGs associated with Hippo or Wnt signaling pathways or were known to encode Zinc-finger proteins (ZNFs). Details of the intersections are included in the Supplementary File 2. Venn diagrams to illustrate the overlap between DEGs from iPSC, MSC, EC and compiled sets of known Hippo or Wnt signaling pathway genes or ZNFs were generated using Venny (https://bioinfogp.cnb.csic.es/tools/venny/index.html). Heatmap representations of the intersecting genes were generated using the Morpheus application (https://software.broadinstitute.org/morpheus). For functional characterization of the intersecting gene sets, we used the ToppCluster (https://pubmed.ncbi.nlm.nih.gov/20484371/), an extended version of ToppFun application of the ToppGene Suite.^21^ Briefly, ToppFun computes enrichment using the hypergeometric model, comparing the differences between the input genes and all genes, to identify the statistical significances of enriched annotations. We used the Benjamini-Hochberg (BH) multiple correction method to adjust the raw p-values, with 0.05 as the threshold. ToppCluster is built on the ToppGene Knowledgebase, which has a compilation of all gene annotations, and can analyze multiple gene lists at a time. Select enriched terms that pass the BH 0.05 p-value (converted to negative log p-values), along with the enriched genes, were exported as XGMML (eXtensible Graph Markup and Modeling Language) file from the ToppCluster application. The XGMML file was then imported into the Cytoscape application^22^ to generate the functional enrichment networks.

### Geltrex *in vitro* capillary-like structure (CLS) formation

The CLS formation of angiogenesis from ECs can be modeled *in vitro* by a Geltrex or Matrigel-based tube formation assay.^23, 24^ ECs plated on Geltrex/Matrigel at low densities form a network of branching structures which can be photographed and quantified by measuring the length, perimeter or area of CLS.^23, 24^ CLS formation was performed in 24-well plates. Briefly, Geltrex (ThermoFisher, Waltham, MA) (200 µl per well) was added to a 24-well plate and incubated at 37°C for 30 minutes. ECs (n=4 for each cell model) were trypsinized and resuspended in EC basal medium without supplements. Each cell type (4.5×10^4^ in 200 µl) was then added into each well and incubated at 37°C with 5% CO_2_ for 12-16 hrs to form CLS. The cells were fixed by 4% buffered formalin at the end of experiments. Images were acquired using an EVOS system (ThermoFisher, Waltham, MA). The CLS wall thickness and perimeter were measured using the ImageJ software.

### In vivo xenograft assay in severe combined immune deficient (SCID) mice

In vivo xenograft assay was performed to determine the angiogenic potentials of ECs derived from control and PWB iPSCs. The experimental procedures have been approved in an animal protocol (# 2581-101681-020822, approval date: 07/29/2022) according to the University of South Carolina institutional animal care and use committee (IACUC). We implanted iPSC-derived control and PWB ECs and mesenchymal stem cells (MSCs) with Geltrex into the subcutaneous layer of the skin of SCID mice using a protocol courtesy of Dr. Joyce Bischoff at Boston Children’s Hospital, Harvard University.^25,26^ Male SCID mice (Nude NIH-III) purchased from Charles River were used. For the transdermal xenografting, each of the normal or PWB EC/MSC (2 x 10^6^ cells with a ratio of EC:MSC as 2:3) was mixed with Geltrex in a total volume of 200 µl per injection. A negative control with Geltrex only without cells was included. Each animal was assigned to three distinct injection sites in the abdominal flanks. A pair of control and PWB EC/MSC/Geltrex mixtures and the Geltrex only negative control were injected into one of three sites in a SCID mouse, respectively. This allowed for side-by-side comparisons to minimize inter-animal variation and reduce animal number usage. The animals were sanctified on day 10 post-injection. The plug-ins were removed, fixed, embedded, and sectioned to examine the formation and remodeling of PWB-like dermal vasculatures *in vivo*.

#### Statistics

The paired samples t-test was performed to evaluate the statistical differences of morphological parameters among vasculature, and differences between PWB and normal control data sets. Data were presented as “mean ± SD”, and *p* < 0.05 was considered significant.

## Results

About 12 - 14 days after the delivery of Yamanaka factors into hDFs, the formation of iPSC colonization was observed (Fig 1A). The iPSC colonies were confirmed using alkaline phosphate live staining (Fig 1B). At the weeks of 3 – 4, the initial iPSC colonies were picked, isolated, and propagated. Some colonies were prone to differentiation, and it was difficult to maintain pluripotency status during screening and propagation. For scientific rigor, these colonies were thus discarded. Two iPSC lines from a human PWB lesion (#4221_3 and #4221_6) and three lines from normal skin (#52521_4, #52521_8 and #52521_9) were successfully expanded and maintained for more than 50 passages, respectively. These cells were further verified to express the iPSC biomarkers, Tra1-60, Nanog, Oct4, and Sox2 using IF staining (Figs 1C-F). In the following EC differentiation and functional characterizations, we mainly focused on PWB iPSC 4221_3 and control iPSC 52521_9 lines.

**Figure 1:**
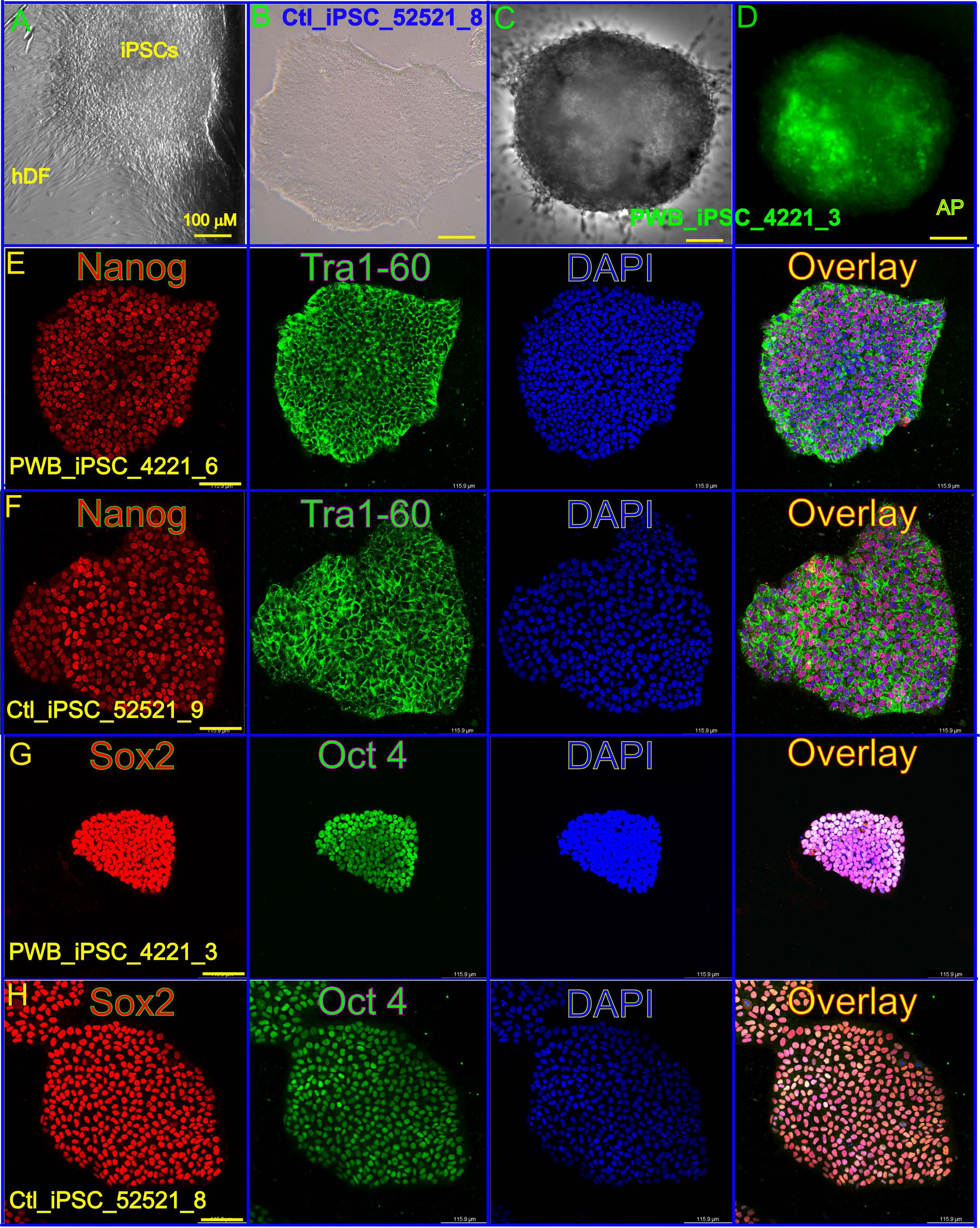
Generation of normal skin and PWB-derived iPSCs. (A) an iPSC colony raised from hDF after reprogramming. (B) typical morphology of a control iPSC colony, Ctl_iPSC_52521_8. (C) PWB iPSC 4221_3 under bright field. (D) AP staining on the same iPSC colony. (E) and (F) control and PWB iPSC colonies expressing stem cell biomarkers Nanog and Tra1-60. (G) and (H) stem cell biomarkers Sox2 and Oct4 were used to verify the control and PWB iPSC cells. DAPI staining was used for nuclei.

Our tailored EC differentiation protocol is outlined in Fig 2A. The differentiation of PWB iPSCs into MSCs was induced on the second day (Fig 2B). Fully differentiated monolayer ECs from PWB iPSCs was present on the eighth day after induction. Approximately 95% purity (based on CD31+ cell profile in flow cytometry, data not shown) of the EC population was achieved from MSCs. The molecular phenotypes of iPSC-derived ECs were verified using IF staining with antibodies recognizing specific biomarkers CD31 and CD144 (Figs 2C, D).

**Figure 2:**
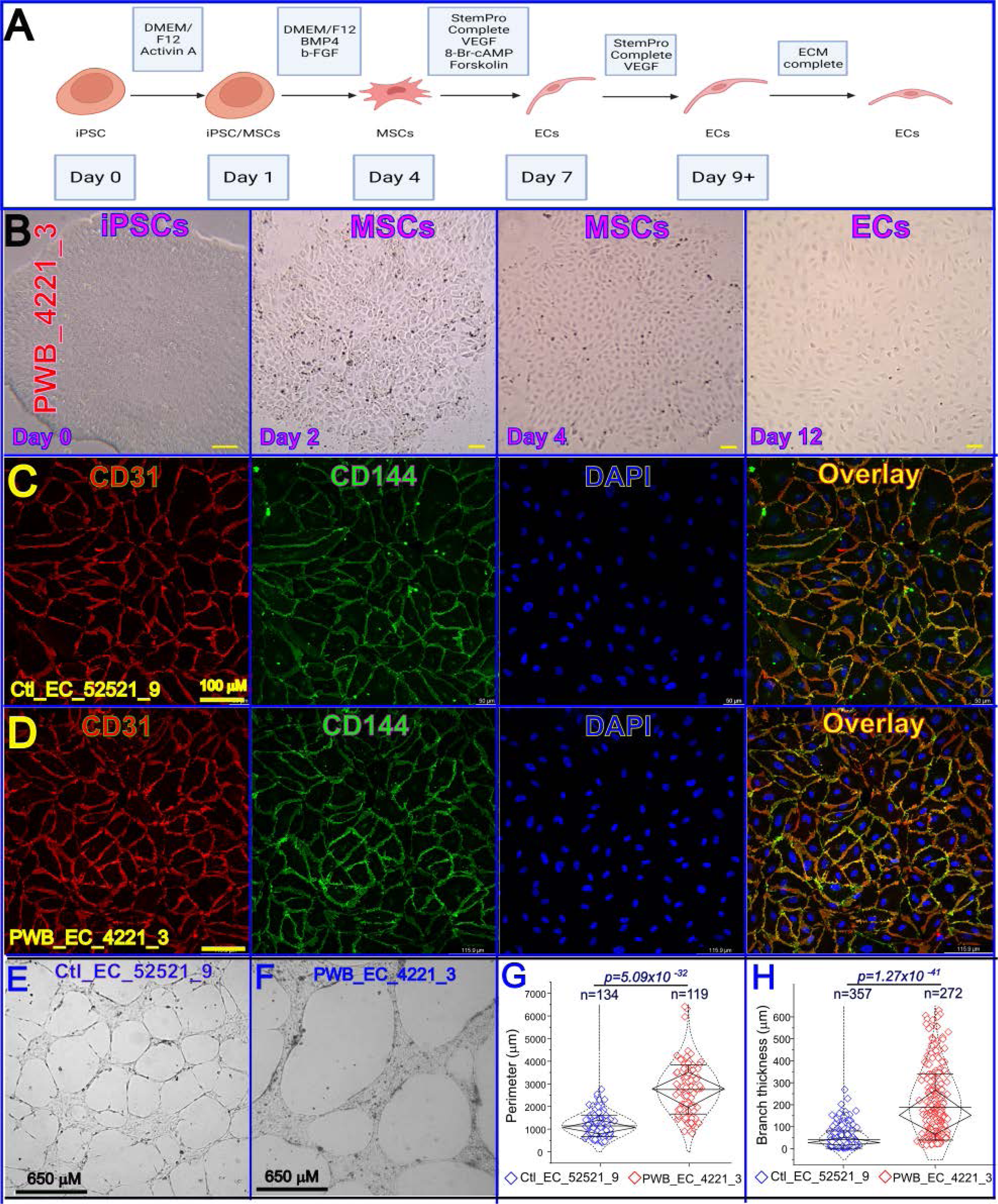
Differentiation of iPSCs into ECs and functional characterizations of PWB iPSC-derived ECs. (A) Flowchart of iPSC differentiation protocol. (B) Morphological changes during PWB iPSC differentiation through MSCs into ECs. MSCs were induced from iPSCs on day 2-4 with a course to EC lineage. The EC induction was performed through day 5 - 12. Fully differentiated monolayer ECs from PWB IPSCs were observed on day 8. (C) and (D) control and PWB iPSC-derived ECs were expressed EC membrane biomarkers CD31 (red) and CD144 (green). Nuclei were stained by DAPI. (E) and (F) control EC_52521_9 and PWB EC_4221_3 formed CLS on Geltrex. (G) and (H) PWB EC_4221_3 showed impaired CLS *in vitro* with larger perimeters (G) (*p*=5.09×10^-32^) and thicker branches (H) *(p*=1.27×10^−41^) as compared to control EC_52521_9. Whiskers: mean±S.D.; Diamond box: interquartile range (IQR); Dotted curve: data distribution. A Mann-Whitney U test was used.

The CLS formation has been used to assess morphological differentiations among a variety of ECs.^27^ In an effort to examine whether PWB ECs could recapitulate pathological phenotypes of PWB vasculature, we first performed CLS formation on Geltrex *in vitro* to characterize morphologies of vasculatures formed by PWB iPSC-derived ECs (Figs 2E-H). The CLS formed by PWB EC_4221 in Geltrex *in vitro* had larger perimeters and greater branches thickness than those formed by normal EC_52521 (Figs 2E-H). Second, we implanted iPSC-derived normal and PWB ECs and MSCs with Geltrex into the subcutaneous layer of the skin of SCID mice. Perfused PWB or normal vasculatures were formed in dermal implants ten days after xenografting (Figs 3A-B), which comprised human ECs recognized by anti-human UEA1 antibody (Figs 3C-D). The vasculature formed by PWB ECs had larger perimeters and higher density than those formed by normal ECs (Figs 3E-G); size distribution showed that the total percentage of perfused large vessels (perimeters > 100 µm) formed by PWB iPSC-derived ECs was significantly higher as compared to those formed by the control iPSC-derived ECs (41.2% vs 16.5%). This was consistent with data from patients’ lesions in our previous report^5^ and a xenografted animal model developed by Dr. Joyce Bischoff group.^25^

**Figure 3:**
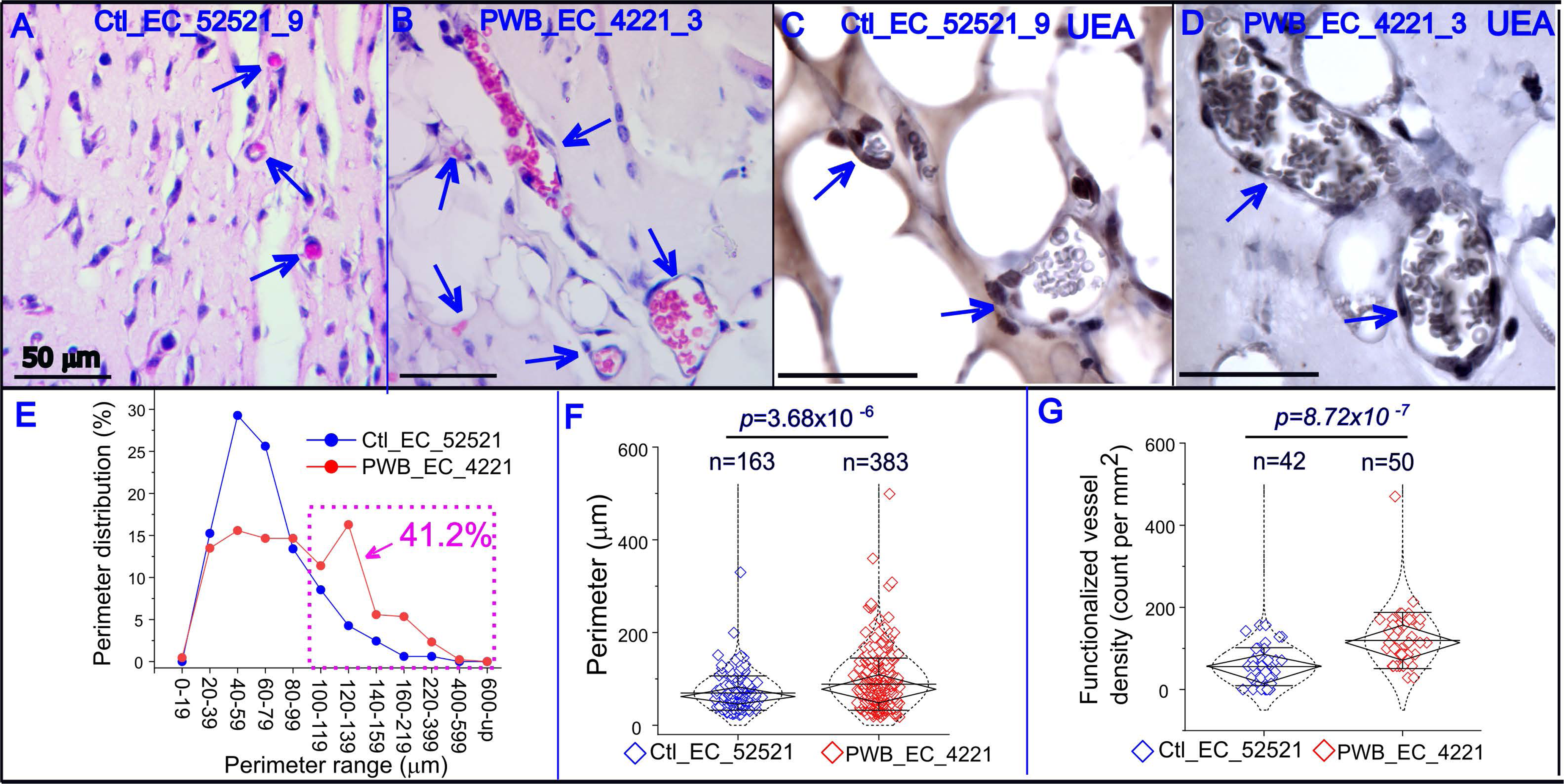
PWB iPSC-derived ECs formed physiologically phenotypic blood vessels *in vivo*. (A) and (B) Formation of perfused human vasculature 10 days after intradermal xenograft of control (A) and PWB (B) ECs with the corresponding MSCs into SCID mice. Arrows: perfused blood vessels in xenografts (H&E staining). (C) and (D) Perfused vasculature comprising human ECs was confirmed using immunohistochemistry (IHC) by an anti-human UEA1 antibody in both control and PWB ECs. (E) Perimeter distribution of xenografted vasculature formed by PWB ECs compared to control vasculature. Pink dashed rectangle: the total percentage (41.2%) of perfused vessels formed by PWB iPSC-derived ECs as compared with 16.5% of perfused vessels formed by control iPSC-derived ECs with perimeters over 100 µm. (F and G) Perfused vasculature formed by PWB iPSC-derived ECs showed (F) bigger perimeters (*p*=3.68×10^-6^) and (G) greater densities (per mm^2^) (*p*=8.72×10^-7^) than those formed by normal ECs. Whiskers: mean±S.D.; Diamond boxes: IQR; Dotted curves: data distribution. A Mann-Whitney U test was used.

Next, we compared the transcriptomes of PWB iPSCs, MSCs, and ECs with their control counterparts using RNA-seq. The full list of DEGs was included in the Supplementary File 1. PWB iPSC 4221 lines showed 1375 upregulated DEGs and 1717 downregulated DEGs compared with control iPSC 52521 lines (FDR<0.05) (Fig 4). PWB MSC 4221 lines showed 644 upregulated DEGs and 744 downregulated DEGs (FDR<0.05) (Fig 5). PWB EC 4221 lines showed 655 upregulated DEGs and 371 downregulated DEGs (FDR<0.05) (Fig 6). DEGs related to Hippo and Wnt pathways were consistently present in iPSC and ECs (p<0.05), as the main significantly enriched pathways during the entire course of differentiation (Figs 4-6). Members of the ZNF gene family were overrepresented among the DEGs in PWB iPSCs (p<0.05; Fig 7) with 108 downregulated and 93 upregulated DEGs related to the ZNF family (Fig 7). The lists of Wnt, Hippo and ZNF signatures and their overlaps crossing iPSCs, MSCs, and ECs were shown in the Supplementary File 2.

**Figure 4:**
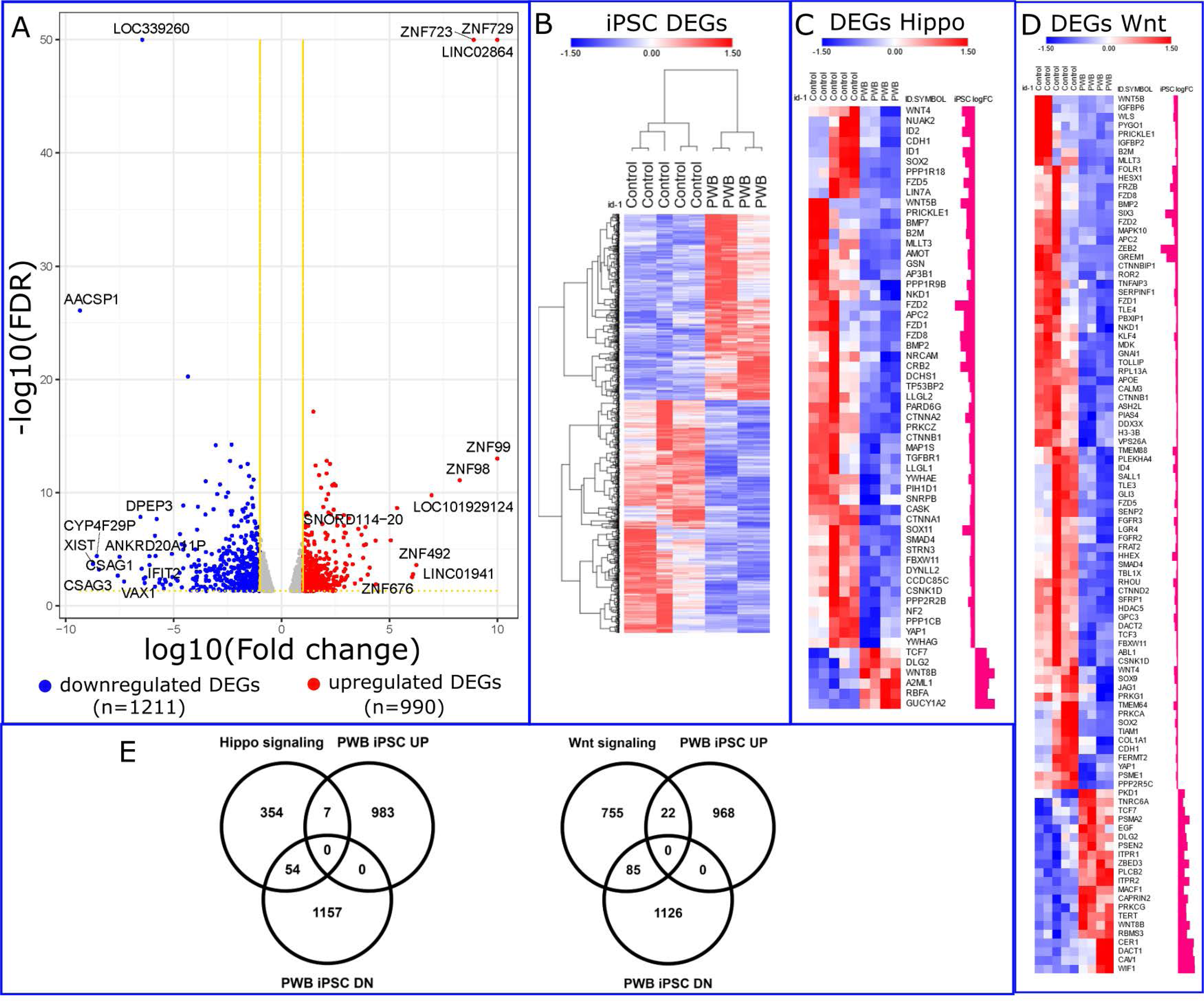
RNA-seq analysis and DEG enrichment in Hippo and Wnt pathways in PWB iPSCs. (A) Volcano plot showing the DEGs in PWB iPSCs as compared to control iPSCs. (B) Heatmap showing the total DEGs in PWB iPSCs vs control iPSCs. (C) Heatmap presenting DEGs in the Hippo pathway. (D) Heatmap showing DEGs in the Wnt pathway. (E) Venn diagram to illustrate up- and down- regulated DEGs in Hippo and Wnt pathways in PWB iPSCs as compared to control iPSCs.

**Figure 5:**
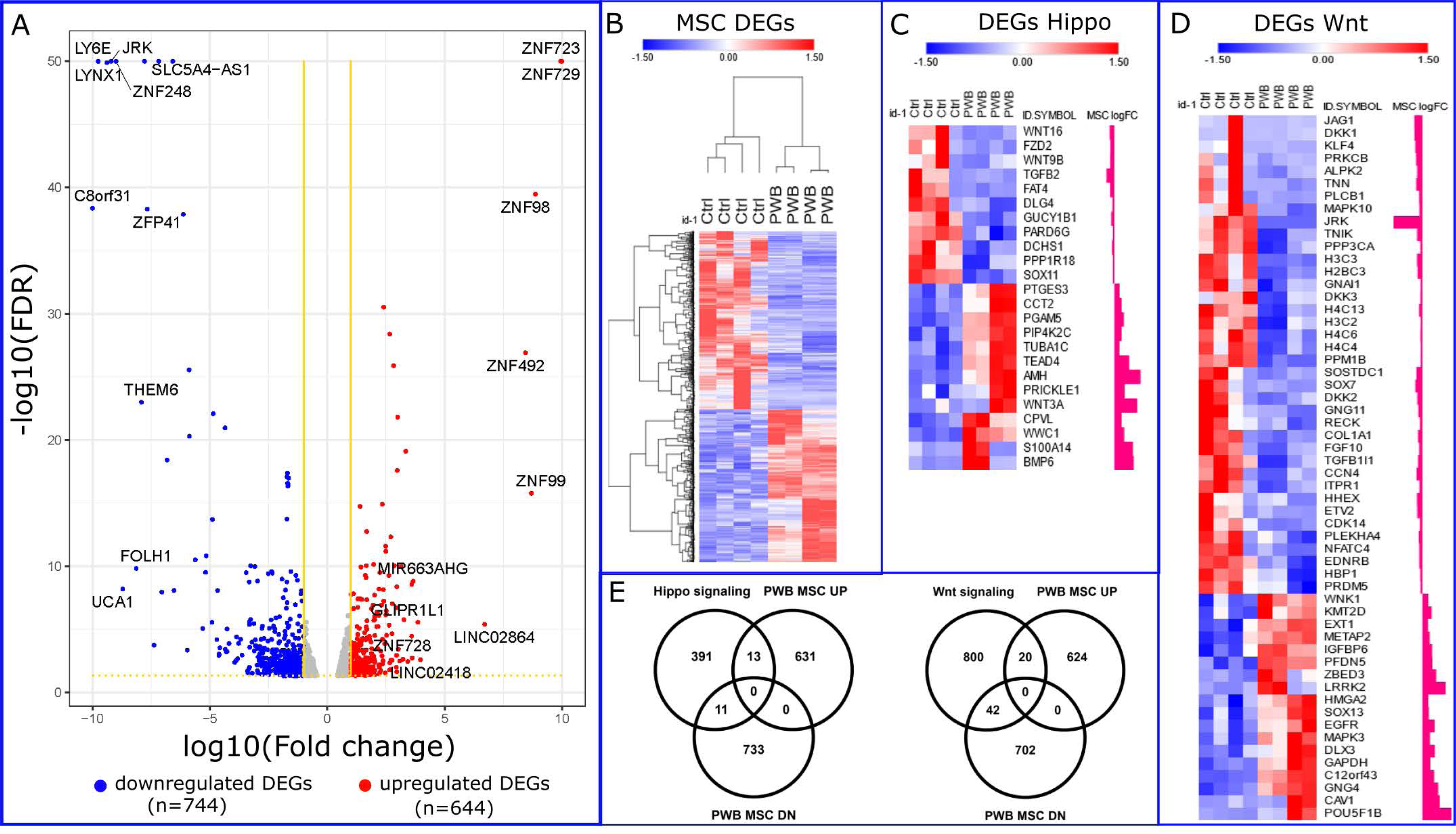
RNA-seq analysis and DEG enrichment in Hippo and Wnt pathways in PWB iPSC- derived MSCs. (A) Volcano plot showing the DEGs in PWB iPSC-derived MSCs as compared to control iPSC-derived MSCs. (B) Heatmap showing the total DEGs in PWB iPSC-derived MSCs vs control iPSC-derived MSCs. (C) Heatmap presenting DEGs in Hippo pathway. (D) Heatmap showing DEGs in Wnt pathway. (E) Venn diagram to illustrate up- and down- regulated DEGs in Hippo and Wnt pathways in PWB iPSC-derived MSCs as compared to control iPSC-derived MSCs.

**Figure 6:**
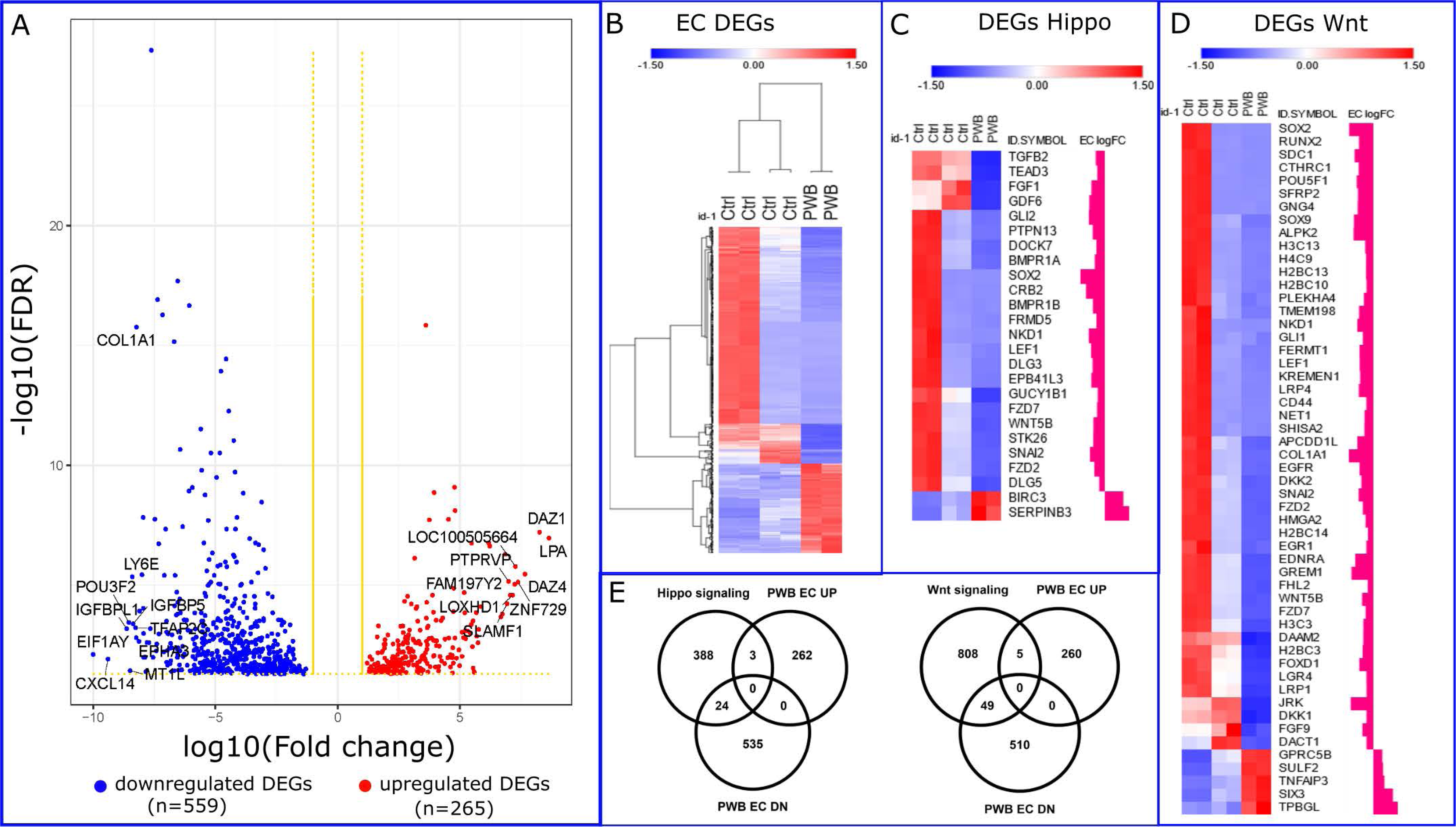
RNA-seq analysis and DEG enrichment in Hippo and Wnt pathways in PWB iPSC- derived ECs. (A) Volcano plot showing the DEGs in PWB iPSC-derived ECs as compared to control iPSC-derived ECs. (B) Heatmap showing the total DEGs in PWB iPSC-derived ECs vs control iPSC-derived ECs. (C) Heatmap representing DEGs in Hippo pathway. (D) Heatmap showing DEGs in Wnt pathway. (E) Venn diagram to illustrate up- and down- regulated DEGs in Hippo and Wnt pathways in PWB iPSC-derived ECs as compared to control iPSC-derived ECs.

**Figure 7:**
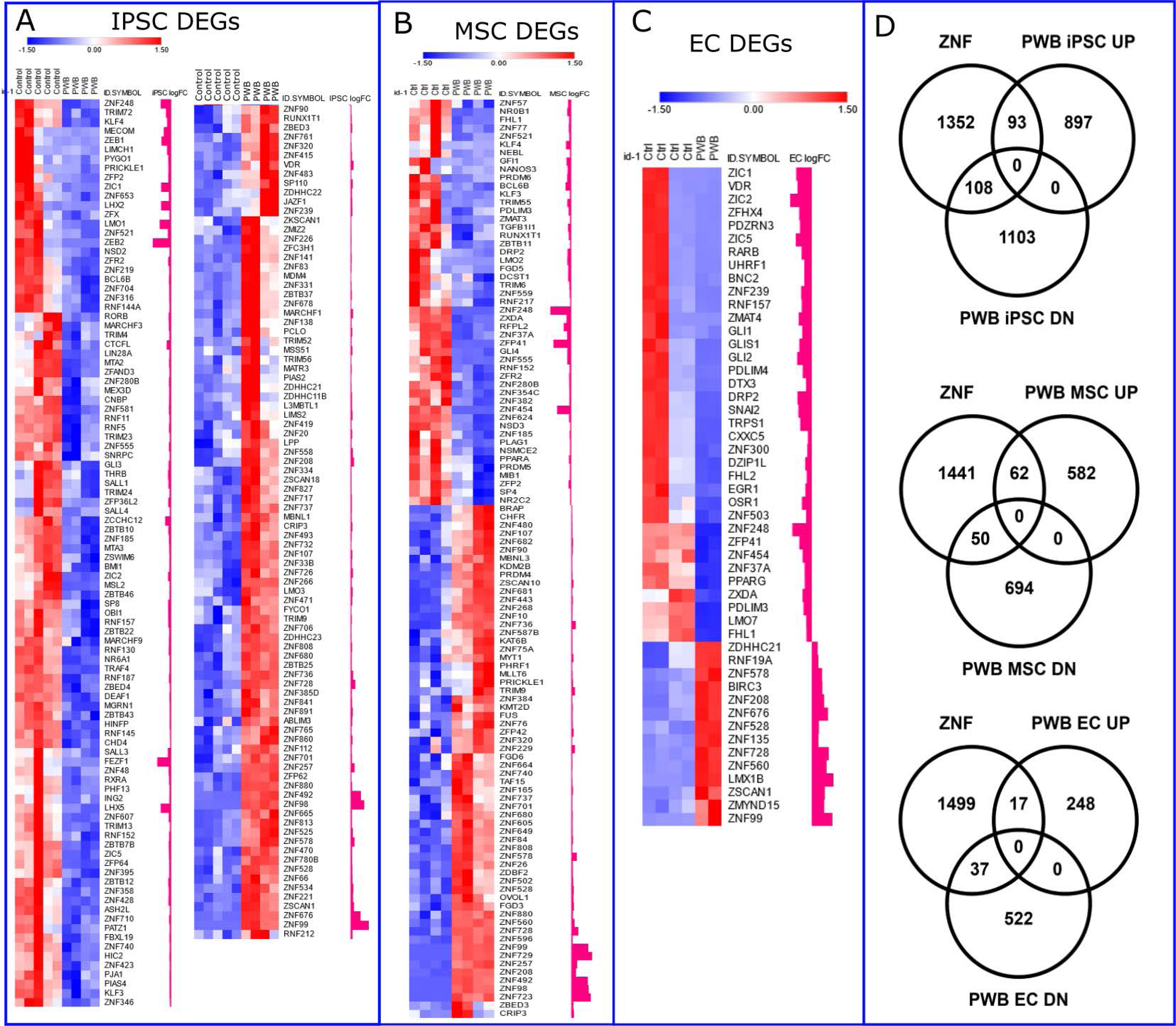
ZNF family members are overrepresented among the DEGs. (A), (B) and (C) Heatmaps representing ZNFs differentially expressed in PWB iPSCs (A), MSCs (B), and ECs (C) as compared with their control counterparts. (D) Venn diagrams to illustrate up- and down- regulated ZNFs in PWB iPSCs, MSCs, and ECs compared with their control counterparts.

Functional enrichment networks showed that DEGs in Hippo and Wnt pathways confer significant functions in vasculature development, tube morphology, endothelium development, and EC differentiation (Figs 8A and 8B). The ZNF DEGs showed significant enrichment for transcriptional regulation, chromatin remodeling, protein ubiquitination, and the retinol acid pathway (Fig 8C). The functional characterizations of the intersecting gene sets related to Wnt, Hippo and ZNFs were shown in the Supplementary File 3. The significantly dysregulated pathways in PWB iPSCs included calcium signaling, gap junction, fatty acid biosynthesis, and vascular smooth muscle cell contraction (Fig 8D). Dysregulation of TNF, NF-kappa B, MAPK, and cholesterol metabolism was found in PWB ECs only, suggesting that these pathways were likely secondary pathologies, resulting from differentiation defects during vascular development (Fig 8D). The overlapping dysregulated pathways in PWB iPSCs and ECs included Hippo, Wnt, regulating pluripotency of stem cells, TGF-beta, axon guidance, and focal adhesion, suggesting their potential roles in both disease development and progression (Fig 8E). Tight junction and cell adhesion were among the significantly dysregulated pathways in PWB iPSCs (Fig 8E). GO enrichment analysis showed top dysregulated biological processes including regulation of developmental processes, tissue/blood vessel differentiation and development, cell differentiation, tube morphogenesis, and cell adhesion in PWB iPSCs, MSCs, and ECs (Supplementary File 4).

**Figure 8:**
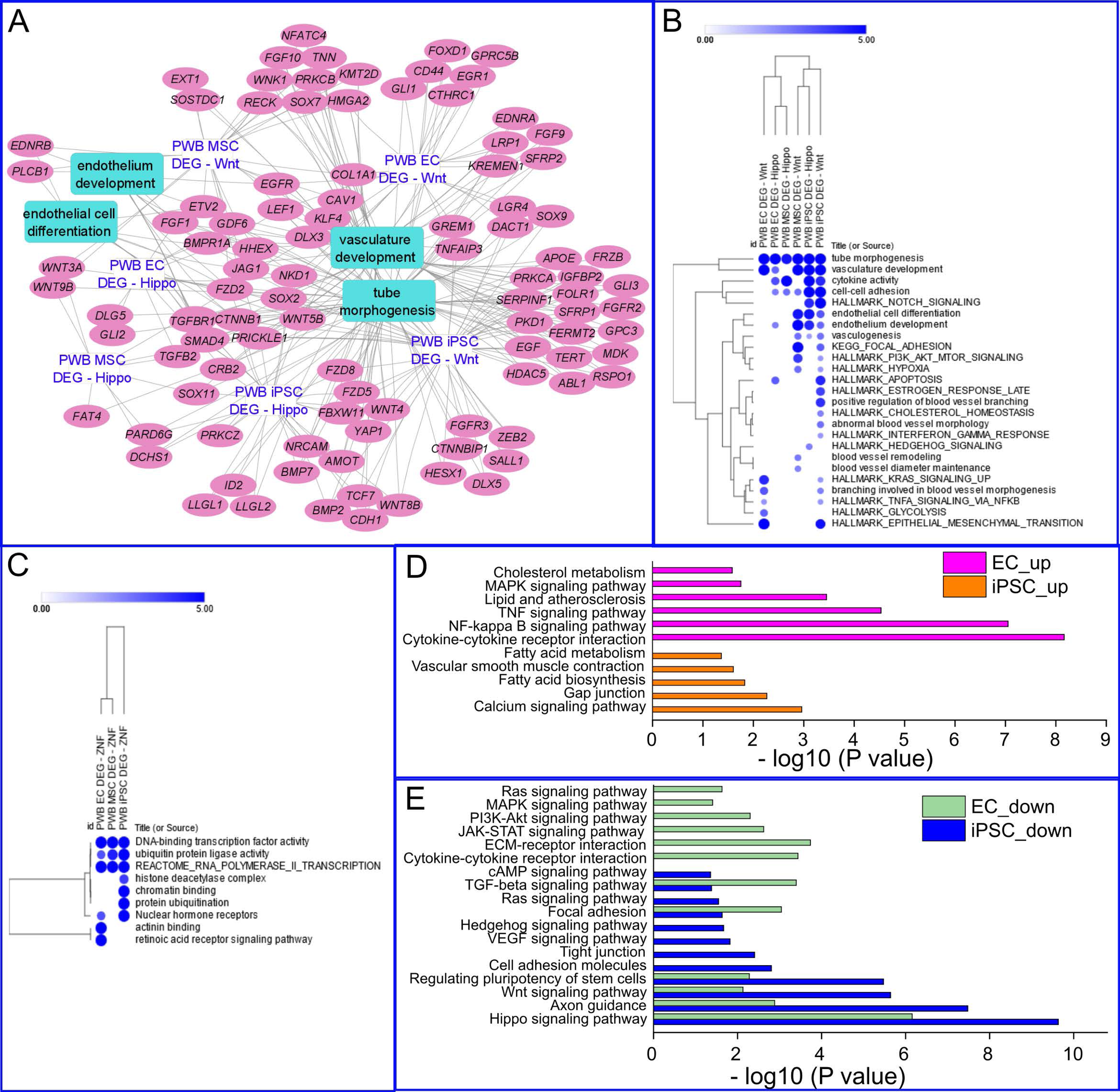
Functional interaction networks of DEGs in Hippo and Wnt pathways and ZNF family. (A) Functional interactive network of Hippo- and Wnt-related DEGs in PWB vasculature showing significant enrichment (FDR<0.05) for tube morphogenesis, endothelium and vasculature development, and EC differentiation. (B) Functional characterization of intersecting DEGs in Hippo and Wnt pathways in affected biological processes. (C) Functional modulations of biological pathways by DEGs in the ZNFs. (D) and (E) KEGG enrichment analysis showing significantly upregulated (D) and downregulated (E) pathways related to vascular differentiation and development in PWB iPSCs and ECs.

## Discussion

In this study, we successfully generated PWB patient-derived iPSCs and differentiated them into ECs. These cells recapitulate many of the vascular phenotypes from donor lesions, which can be used as a novel disease model for PWB. Dysregulated pathways such as Hippo, Wnt, regulating pluripotency of stem cells persist through PWB iPSCs to ECs during the differentiation process. These data suggest their contributive roles in both disease development and progression. Dysregulations of TNF, NF-kappa B, MAPK, and cholesterol metabolism are in PWB ECs but not evident in iPSCs, which suggests that impairments of these pathways are downstream results from differentiation defects during vascular development. Our data also suggest that dysregulated Hippo and Wnt pathways could be potential targets for therapeutic intervention development for PWB.

PWB results from developmental defects of the vasculature during embryonic stages. This is supported by the evidence that PWB ECs exhibit stem-cell-like phenotypes.^5, 7^ There is an urgent need for knowledge concerning the developmental defects of PWB ECs and underlying mechanisms. The Hippo pathway plays a fundamental role in controlling organ size, stem cell proliferation, and tissue repair and regeneration.^28, 29^ The Wnt pathway conveys crucial signaling that regulates cell fate, migration, polarity, and organogenesis.^30, 31^ The crosstalk between these two signalosomes coordinates their functions during development.^32, 33^ Due to the developmental vascular defects and soft tissue hypertrophy overgrowth in PWB, it is reasonable to posit the critical roles of impaired Hippo and Wnt pathways in the developmental defects and progression of vascular phenotypes in PWB lesions. Indeed, dysregulated Hippo and Wnt pathways have been found through a whole transcriptome atlas analysis of individual lesional vasculature on PWB formalin-fixed paraffin-embedded (FFPE) sections using a GeoMx digital spatial profiler (Nguyen, Wang, Jegga and Tan, personal communications). This data demonstrates that the dysregulated Hippo and Wnt pathways in iPSCs could be inherited from PWB lesional tissues and passed into MSCs and ECs during differentiation. There are two possible underlying mechanisms: (1) PWB iPSCs replicate the genome of lesional hDFs, such as processing related genetic abnormalities from lesional hDFs, thus preserving dysregulated Hippo and Wnt pathways which are incorporated into their differentiated ECs; and (2) PWB iPSCs retain an ’epigenetic memory’ of lesional hDFs, through which disease-related DNA methylation signatures are preserved from the original somatic cells.^34^ This could make PWB iPSCs prone to lineage differentiation propensity of donor ECs, thus matching disease-associated phenotypes.

Members of ZNFs were overrepresented among the DEGs in PWB iPSCs, MSCs, and ECs. The zinc-finger domains of ZNFs confer interactions with DNA, RNA, and proteins, involving many cellular processes such as transcriptional regulation, chromatin remodeling, ubiquitin-mediated proteasome, retinol acid signaling.^35^ As a major group of transcriptional modulators, ZNFs may play an important role in regulating Hippo and Wnt pathways in PWB. For example, both ZNF E-box-binding homebox (ZEB) 1 and 2 were significantly downregulated in PWB iPSCs. ZEBs function as a transcriptional repressor for epithelial genes to maintain an undifferentiated phenotype during epithelial-mesenchymal transition (EMT).^36^ TGF-β responsive genes and cell adhesion and junction genes are the transcriptional targets of ZEBs.^36^ Evidence also has shown the direct transcriptional regulation of ZNFs on Hippo pathways. For example, ZEB1 can directly interact with YAP, switching ZEB1 from a repressor to a transcriptional activator to cause cancer-promoting effects.^37^ This suggests the potential role of ZNFs in disease development of PWB, but the detailed mechanisms are yet to be determined in future studies.

There are a few limitations to this study. First, this study is the first to reveal the dysregulation of Hippo and Wnt pathways in PWB iPSCs and ECs; however, the detailed mechanisms of these dysregulated pathways underlying the development of PWB ECs are yet to be determined. The following studies will be performed in the future: (a) a series of functionally perturbative experiments will be carried out to identify the crucial DEGs related to Hippo and Wnt pathways in pathological development of vascular malformations; and (b) inhibitors and modulators to Hippo and Wnt pathways will be screened and investigated for their mitigative roles in vascular malformations. Second, the two PWB iPSC lines generated in this study were from one patient; they recapitulate major vascular phenotypes of PWB but cannot harbor a full spectrum of pathology crossing patient populations. We are aware that one general and ubiquitous challenge in iPSC-based studies is the “statistic power inefficiency”. Therefore, generalization of conclusions to the broader patient population needs to be cautious. This barrier can be addressed through (a) the establishment of a biobank of iPSC lines derived from PWB patients with various demographics at different lesional stages, (b) generation of isogenic iPSC lines, and (c) validation of those perturbative pathways in PWB vasculature from patients’ biopsies. These will be major aims of our future studies. Nevertheless, this study has first demonstrated the scientific value and feasibility of patient-derived iPSCs as PWB disease models.

In conclusion, PWB iPSCs and their-derived ECs retain pathological vascular phenotypes which can serve clinically relevant models. Our data demonstrate multiple pathways, such as Hippo and Wnt, NF-kappa B, TNF, MAPK, and cholesterol metabolism, are dysregulated, which may contribute to the development of differentiation-defective ECs in PWB.

## Acknowledgement

We are very thankful to the support and assistance from Instrumentation Resource Facility at University of South Carolina School of Medicine. We are also grateful to Arieleus Taine from the Spring Valley High School, Columbia, SC, who participated and performed the RNA-seq data analysis from MSCs.

## Acknowledgement for the DoD award HT9425-23-10008

i “The U.S. Army Medical Research Acquisition Activity, 820 Chandler Street, Fort Detrick MD 21702-5014 is the awarding and administering acquisition office.” and;
ii “This work was supported by The Assistant Secretary of Defense for Health Affairs endorsed by the Department of Defense, in the amount of $297,956.00, through the Peer Reviewed Medical Research Program under Award Number (HT9425-23-10008). Opinions, interpretations, conclusions, and recommendations contained herein are those of the author(s) and are not necessarily endorsed by the Department of Defense.”
i “In conducting research using animals, the investigator(s) adhere(s) to the laws of the United States and regulations of the Department of Agriculture.”
i “In the conduct of research utilizing recombinant DNA, the investigator(s) adhered to NIH Guidelines for research involving recombinant DNA molecules.”
ii “In the conduct of research involving hazardous organisms or toxins, the investigator(s) adhered to the CDC-NIH Guide for Biosafety in Microbiological and Biomedical Laboratories.”

## Database deposition

The processed bam files of RNA-seq are deposited into NIH SRA (Sequence Read Archive) (BioProject ID: PRJNA997591). The transcriptome data (raw counts and pseudo counts for each sample) are to be deposited into the NCBI GEO (Gene Expression Omnibus) (Accession number: GSE240770). Data are available to the public.

