## Supplementary material for "Supporting materials: Endothelial cells differentiated from patient dermal fibroblast-derived induced pluripotent stem cells resemble vascular malformations of Port Wine Birthmark": Suppl File

#### Supplementary File 1: DEGs for iPSCs, MSCs and ECs (PWB vs control, FDR < 0.05)

| ID. | ENTREZID | Group | ID.SYMBOL | logFC | ID. | ENTREZID | Group | ID.SYMBOL | logFC | ID. | ENTREZID | Group | ID.SYMBOL | logFC |
|---|---|---|---|---|---|---|---|---|---|---|---|---|---|---|
| ENSG00000196350 | 100287226 | PWB-iPSC UP | ZNF729 | 11.13839 | ENSG00000107882 | 51684 | PWB-iPSC DN | SUFU | -0.758855213 | ENSG00000105472 | 6320 | PWB-EC DN | CLEC11A | -5.94554 |
| ENSG00000213973 | 7652 | PWB-iPSC UP | ZNF99 | 11.12093 | ENSG00000167771 | 283248 | PWB-iPSC DN | RCOR2 | -0.760534011 | ENSG00000149633 | 85449 | PWB-EC DN | KIAA1755 | -6.025065 |
| ENSG00000263711 | 400655 | PWB-iPSC UP | LINC02864 | 10.47124 | ENSG00000119280 | 84886 | PWB-iPSC DN | C1orf198 | -0.760916436 | ENSG00000104783 | 3783 | PWB-EC DN | KCNN4 | -6.035076 |
| ENSG00000268696 | 646864 | PWB-iPSC UP | ZNF723 | 8.902628 | ENSG00000105355 | 10226 | PWB-iPSC DN | PLIN3 | -0.761309548 | ENSG00000168542 | 1281 | PWB-EC DN | COL3A1 | -6.066131 |
| ENSG00000197360 | 148198 | PWB-iPSC UP | ZNF98 | 8.254292 | ENSG00000124194 | 78997 | PWB-iPSC DN | GDAP1L1 | -0.76642267 | ENSG00000151617 | 1909 | PWB-EC DN | EDNRA | -6.079957 |
| ENSG00000260599 | 101929124 | PWB-iPSC UP | LOC101929124 | 6.950827 | ENSG00000171345 | 3880 | PWB-iPSC DN | KRT19 | -0.771561437 | ENSG00000198910 | 3897 | PWB-EC DN | L1CAM | -6.131435 |
| ENSG00000229676 | 57615 | PWB-iPSC UP | ZNF492 | 6.24507 | ENSG00000177469 | 284119 | PWB-iPSC DN | CAVIN1 | -0.771664818 | ENSG00000165810 | 153579 | PWB-EC DN | BTNL9 | -6.151035 |
| ENSG00000226813 | 105373600 | PWB-iPSC UP | LINC01941 | 6.095638 | ENSG00000170604 | 26145 | PWB-iPSC DN | IRF2BP1 | -0.772065956 | ENSG00000162745 | 25903 | PWB-EC DN | OLFML2B | -6.184051 |
| ENSG00000196109 | 163223 | PWB-iPSC UP | ZNF676 | 6.051482 | ENSG00000103647 | 10391 | PWB-iPSC DN | CORO2B | -0.772490234 | ENSG00000148204 | 286204 | PWB-EC DN | CRB2 | -6.273775 |
| ENSG00000202048 | 767598 | PWB-iPSC UP | SNORD114-20 | 5.355448 | ENSG00000189067 | 9516 | PWB-iPSC DN | LITAF | -0.772621495 | ENSG00000108679 | 3959 | PWB-EC DN | LGALS3BP | -6.281347 |
| ENSG00000147724 | 51059 | PWB-iPSC UP | FAM135B | 5.0603 | ENSG00000146066 | 192286 | PWB-iPSC DN | HIGD2A | -0.772768633 | ENSG00000115884 | 6382 | PWB-EC DN | SDC1 | -6.316411 |
| ENSG00000022556 | 55655 | PWB-iPSC UP | NLRP2 | 4.38072 | ENSG00000139800 | 85416 | PWB-iPSC DN | ZIC5 | -0.773023299 | ENSG00000069482 | 51083 | PWB-EC DN | GAL | -6.338386 |
| ENSG00000279000 | 390093 | PWB-iPSC UP | OR10A6 | 4.35506 | ENSG00000101945 | 6839 | PWB-iPSC DN | SUV39H1 | -0.773315551 | ENSG00000131459 | 9945 | PWB-EC DN | GFPT2 | -6.340887 |
| ENSG00000226258 | 101927347 | PWB-iPSC UP | GRM7-AS3 | 4.086098 | ENSG00000130669 | 10298 | PWB-iPSC DN | PAK4 | -0.773887614 | ENSG00000146910 | 285888 | PWB-EC DN | CNPY1 | -6.378452 |
| ENSG00000248228 | 100505893 | PWB-iPSC UP | SLIT2-IT1 | 3.997047 | ENSG00000165029 | 19 | PWB-iPSC DN | ABCA1 | -0.773946252 | ENSG00000148344 | 9536 | PWB-EC DN | PTGES | -6.384785 |
| ENSG00000242781 | 105377175 | PWB-iPSC UP | LINC02050 | 3.909852 | ENSG00000054277 | 23596 | PWB-iPSC DN | OPN3 | -0.774620941 | ENSG00000084636 | 1307 | PWB-EC DN | COL16A1 | -6.431039 |
| ENSG00000246228 | 727677 | PWB-iPSC UP | CASC8 | 3.892527 | ENSG00000161940 | 255877 | PWB-iPSC DN | BCL6B | -0.774849808 | ENSG00000087116 | 9509 | PWB-EC DN | ADAMTS2 | -6.440986 |
| ENSG00000241336 | 101928190 | PWB-iPSC UP | LINC01487 | 3.817409 | ENSG00000113657 | 1809 | PWB-iPSC DN | DPYSL3 | -0.775277086 | ENSG00000249992 | 25907 | PWB-EC DN | TMEM158 | -6.44303 |
| ENSG00000182261 | 338322 | PWB-iPSC UP | NLRP10 | 3.783649 | ENSG00000168159 | 149603 | PWB-iPSC DN | RNF187 | -0.775342901 | ENSG00000106236 | 4885 | PWB-EC DN | NPTX2 | -6.450207 |
| ENSG00000251450 | 102524628 | PWB-iPSC UP | RASGRF2-AS1 | 3.703892 | ENSG00000167680 | 10501 | PWB-iPSC DN | SEMA6B | -0.775652007 | ENSG00000100842 | 10278 | PWB-EC DN | EFS | -6.461771 |
| ENSG00000142583 | 6518 | PWB-iPSC UP | SLC2A5 | 3.678567 | ENSG00000158164 | 11013 | PWB-iPSC DN | TMSB15A | -0.776028167 | ENSG00000254377 | 100130155 | PWB-EC DN | MIR124-2HG | -6.469124 |
| ENSG00000285691 | 105377982 | PWB-iPSC UP | LOC105377982 | 3.557985 | ENSG00000147394 | 7739 | PWB-iPSC DN | ZNF185 | -0.77654113 | ENSG00000206432 | 645369 | PWB-EC DN | TMEM200C | -6.474355 |
| ENSG00000248995 | 109729131 | PWB-iPSC UP | LINC02231 | 3.397926 | ENSG00000108840 | 10014 | PWB-iPSC DN | HDAC5 | -0.77897507 | ENSG00000221698 | 100302287 | PWB-EC DN | MIR548H3 | -6.512905 |
| ENSG00000268964 | 100271846 | PWB-iPSC UP | ERVV-2 | 3.297647 | ENSG00000077150 | 4791 | PWB-iPSC DN | NFKB2 | -0.781887779 | ENSG00000177508 | 79191 | PWB-EC DN | IRX3 | -6.532727 |
| ENSG00000266976 | 102724908 | PWB-iPSC UP | LOC102724908 | 3.162525 | ENSG00000275004 | 140883 | PWB-iPSC DN | ZNF280B | -0.781940304 | ENSG00000125398 | 6662 | PWB-EC DN | SOX9 | -6.537044 |
| ENSG00000115353 | 6869 | PWB-iPSC UP | TACR1 | 3.123025 | ENSG00000168502 | 23255 | PWB-iPSC DN | MTCL1 | -0.782159454 | ENSG00000165023 | 54769 | PWB-EC DN | DIRAS2 | -6.542702 |
| ENSG00000183378 | 341277 | PWB-iPSC UP | OVCH2 | 3.073622 | ENSG00000063177 | 6141 | PWB-iPSC DN | RPL18 | -0.782825648 | ENSG00000133110 | 10631 | PWB-EC DN | POSTN | -6.544618 |
| ENSG00000277526 | 101927815 | PWB-iPSC UP | LOC101927815 | 3.052435 | ENSG00000148180 | 2934 | PWB-iPSC DN | GSN | -0.783529967 | ENSG00000233639 | 100506421 | PWB-EC DN | PANTR1 | -6.55402 |
| ENSG00000249267 | 400084 | PWB-iPSC UP | LINC00939 | 3.015017 | ENSG00000213465 | 402 | PWB-iPSC DN | ARL2 | -0.784121352 | ENSG00000157445 | 55799 | PWB-EC DN | CACNA2D3 | -6.554523 |
| ENSG00000279486 | 144125 | PWB-iPSC UP | OR2AG1 | 2.986184 | ENSG00000179242 | 1002 | PWB-iPSC DN | CDH4 | -0.787247996 | ENSG00000173546 | 1464 | PWB-EC DN | CSPG4 | -6.557707 |
| ENSG00000271860 | 101927314 | PWB-iPSC UP | LOC101927314 | 2.973097 | ENSG00000104856 | 5971 | PWB-iPSC DN | RELB | -0.787453823 | ENSG00000152377 | 6695 | PWB-EC DN | SPOCK1 | -6.576275 |
| ENSG00000179813 | 144809 | PWB-iPSC UP | FAM216B | 2.951989 | ENSG00000138356 | 316 | PWB-iPSC DN | AOX1 | -0.788062662 | ENSG00000144218 | 3899 | PWB-EC DN | AFF3 | -6.596727 |
| ENSG00000188596 | 144535 | PWB-iPSC UP | CFAP54 | 2.951143 | ENSG00000164307 | 51752 | PWB-iPSC DN | ERAP1 | -0.788348465 | ENSG00000198796 | 115701 | PWB-EC DN | ALPK2 | -6.645784 |
| ENSG00000137473 | 83894 | PWB-iPSC UP | TTC29 | 2.931544 | ENSG00000139266 | 92979 | PWB-iPSC DN | MARCHF9 | -0.788556205 | ENSG00000128342 | 3976 | PWB-EC DN | LIF | -6.670042 |
| ENSG00000254951 | 283299 | PWB-iPSC UP | LOC283299 | 2.92762 | ENSG00000170540 | 23204 | PWB-iPSC DN | ARL6IP1 | -0.789178564 | ENSG00000113721 | 5159 | PWB-EC DN | PDGFRB | -6.687117 |
| ENSG00000255693 | 400046 | PWB-iPSC UP | LINC02389 | 2.903714 | ENSG00000186300 | 148254 | PWB-iPSC DN | ZNF555 | -0.79104507 | ENSG00000264163 | 100500906 | PWB-EC DN | MIR3689B | -6.68912 |
| ENSG00000269067 | 388523 | PWB-iPSC UP | ZNF728 | 2.892835 | ENSG00000090097 | 57060 | PWB-iPSC DN | PCBP4 | -0.791849633 | ENSG00000138944 | 85352 | PWB-EC DN | SHISAL1 | -6.77378 |
| ENSG00000163017 | 72 | PWB-iPSC UP | ACTG2 | 2.888181 | ENSG00000176532 | 222171 | PWB-iPSC DN | PRR15 | -0.792386016 | ENSG00000128564 | 7425 | PWB-EC DN | VGF | -6.777475 |
| ENSG00000259163 | 101928909 | PWB-iPSC UP | LOC101928909 | 2.887461 | ENSG00000160783 | 11243 | PWB-iPSC DN | PMF1 | -0.793687123 | ENSG00000163814 | 64866 | PWB-EC DN | CDCP1 | -6.793521 |
| ENSG00000214548 | 55384 | PWB-iPSC UP | MEG3 | 2.878751 | ENSG00000129194 | 6665 | PWB-iPSC DN | SOX15 | -0.793714631 | ENSG00000215808 | 339535 | PWB-EC DN | LINC01139 | -6.819761 |
| ENSG00000261615 | 101929144 | PWB-iPSC UP | LINC01858 | 2.774397 | ENSG00000159842 | 29 | PWB-iPSC DN | ABR | -0.79429245 | ENSG00000090104 | 5996 | PWB-EC DN | RGS1 | -6.831377 |
| ENSG00000123201 | 2974 | PWB-iPSC UP | GUCY1B2 | 2.772505 | ENSG00000171823 | 144699 | PWB-iPSC DN | FBXL14 | -0.797092119 | ENSG00000240563 | 54596 | PWB-EC DN | L1TD1 | -6.845537 |
| ENSG00000134533 | 85004 | PWB-iPSC UP | RERG | 2.763656 | ENSG00000237945 | 100506334 | PWB-iPSC DN | LINC00649 | -0.798144477 | ENSG00000240694 | 10687 | PWB-EC DN | PNMA2 | -6.849107 |
| ENSG00000286329 | 105377177 | PWB-iPSC UP | LOC105377177 | 2.755526 | ENSG00000270885 | 91608 | PWB-iPSC DN | RASL10B | -0.798158637 | ENSG00000101463 | 79953 | PWB-EC DN | SYNDIG1 | -6.937867 |
| ENSG00000268257 | 100271873 | PWB-iPSC UP | AIRN | 2.680856 | ENSG00000187735 | 6917 | PWB-iPSC DN | TCEA1 | -0.798409073 | ENSG00000231107 | 101927873 | PWB-EC DN | LINC01508 | -6.96125 |
| ENSG00000250295 | 101926926 | PWB-iPSC UP | RDH10-AS1 | 2.615551 | ENSG00000151948 | 144423 | PWB-iPSC DN | GLT1D1 | -0.7994763 | ENSG00000101938 | 91851 | PWB-EC DN | CHRDL1 | -6.966936 |
| ENSG00000233993 | 101929526 | PWB-iPSC UP | DTD1-AS1 | 2.604932 | ENSG00000160014 | 808 | PWB-iPSC DN | CALM3 | -0.799560326 | ENSG00000137959 | 10964 | PWB-EC DN | IFI44L | -6.99139 |
| ENSG00000139915 | 161357 | PWB-iPSC UP | MDGA2 | 2.584314 | ENSG00000068976 | 5837 | PWB-iPSC DN | PYGM | -0.801778881 | ENSG00000258602 | 105370578 | PWB-EC DN | LINC01629 | -6.993306 |
| ENSG00000166573 | 2587 | PWB-iPSC UP | GALR1 | 2.565215 | ENSG00000089289 | 3476 | PWB-iPSC DN | IGBP1 | -0.801964022 | ENSG00000135903 | 5077 | PWB-EC DN | PAX3 | -6.999931 |
| ENSG00000254656 | 388015 | PWB-iPSC UP | RTL1 | 2.548115 | ENSG00000189143 | 1364 | PWB-iPSC DN | CLDN4 | -0.802491518 | ENSG00000095752 | 3589 | PWB-EC DN | IL11 | -7.035367 |
| ENSG00000226031 | 100129662 | PWB-iPSC UP | FGF13-AS1 | 2.546518 | ENSG00000125650 | 5623 | PWB-iPSC DN | PSPN | -0.8049657 | ENSG00000120708 | 7045 | PWB-EC DN | TGFBI | -7.088866 |
| ENSG00000240875 | 730091 | PWB-iPSC UP | LINC00886 | 2.533619 | ENSG00000161914 | 115950 | PWB-iPSC DN | ZNF653 | -0.806589607 | ENSG00000172935 | 116535 | PWB-EC DN | MRGPRF | -7.103307 |
| ENSG00000152595 | 56955 | PWB-iPSC UP | MEPE | 2.525259 | ENSG00000164104 | 3148 | PWB-iPSC DN | HMGB2 | -0.807135205 | ENSG00000198105 | 57209 | PWB-EC DN | ZNF248 | -7.161316 |
| ENSG00000197182 | 400931 | PWB-iPSC UP | MIRLET7BHG | 2.515902 | ENSG00000116661 | 26232 | PWB-iPSC DN | FBXO2 | -0.808841449 | ENSG00000113361 | 1004 | PWB-EC DN | CDH6 | -7.312584 |
| ENSG00000212807 | 402317 | PWB-iPSC UP | OR2A42 | 2.509695 | ENSG00000129038 | 4016 | PWB-iPSC DN | LOXL1 | -0.809669465 | ENSG00000061337 | 11178 | PWB-EC DN | LZTS1 | -7.316474 |
| ENSG00000241832 | 27442 | PWB-iPSC UP | CECR3 | 2.491559 | ENSG00000204580 | 780 | PWB-iPSC DN | DDR1 | -0.80997626 | ENSG00000166923 | 26585 | PWB-EC DN | GREM1 | -7.36636 |
| ENSG00000197134 | 113835 | PWB-iPSC UP | ZNF257 | 2.474883 | ENSG00000126016 | 154796 | PWB-iPSC DN | AMOT | -0.812916679 | ENSG00000164778 | 2020 | PWB-EC DN | EN2 | -7.39788 |
| ENSG00000182376 | 339059 | PWB-iPSC UP | LOC339059 | 2.464517 | ENSG00000169857 | 57099 | PWB-iPSC DN | AVEN | -0.813201602 | ENSG00000144834 | 29114 | PWB-EC DN | TAGLN3 | -7.420508 |
| ENSG00000272727 | 105377466 | PWB-iPSC UP | LINC02266 | 2.45893 | ENSG00000204175 | 9721 | PWB-iPSC DN | GPRIN2 | -0.813744604 | ENSG00000163359 | 1293 | PWB-EC DN | COL6A3 | -7.476669 |
| ENSG00000206113 | 402160 | PWB-iPSC UP | CFAP99 | 2.434527 | ENSG00000179603 | 2918 | PWB-iPSC DN | GRM8 | -0.8148132 | ENSG00000155093 | 5799 | PWB-EC DN | PTPRN2 | -7.54804 |
| ENSG00000258405 | 147660 | PWB-iPSC UP | ZNF578 | 2.431609 | ENSG00000171425 | 51545 | PWB-iPSC DN | ZNF581 | -0.814904178 | ENSG00000234616 | 8629 | PWB-EC DN | JRK | -7.620144 |
| ENSG00000250155 | 107986412 | PWB-iPSC UP | LOC107986412 | 2.429601 | ENSG00000140577 | 64784 | PWB-iPSC DN | CRTC3 | -0.815557793 | ENSG00000198914 | 5455 | PWB-EC DN | POU3F3 | -7.66098 |
| ENSG00000186439 | 10345 | PWB-iPSC UP | TRDN | 2.425607 | ENSG00000158292 | 387509 | PWB-iPSC DN | GPR153 | -0.816291361 | ENSG00000157851 | 56896 | PWB-EC DN | DPYSL5 | -7.717933 |
| ENSG00000243896 | 401427 | PWB-iPSC UP | OR2A7 | 2.416343 | ENSG00000206503 | 3105 | PWB-iPSC DN | HLA-A | -0.817901705 | ENSG00000198729 | 81706 | PWB-EC DN | PPP1R14C | -7.79527 |
| ENSG00000180658 | 79541 | PWB-iPSC UP | OR2A4 | 2.409157 | ENSG00000108821 | 1277 | PWB-iPSC DN | COL1A1 | -0.821255423 | ENSG00000123496 | 3598 | PWB-EC DN | IL13RA2 | -7.844511 |
| ENSG00000162365 | 284541 | PWB-iPSC UP | CYP4A22 | 2.400109 | ENSG00000165886 | 80019 | PWB-iPSC DN | UBTD1 | -0.821796139 | ENSG00000176971 | 387758 | PWB-EC DN | FIBIN | -7.879474 |
| ENSG00000105205 | 1178 | PWB-iPSC UP | CLC | 2.399292 | ENSG00000213563 | 414919 | PWB-iPSC DN | C8orf82 | -0.822464856 | ENSG00000162493 | 10630 | PWB-EC DN | PDPN | -7.93398 |
| ENSG00000103546 | 6530 | PWB-iPSC UP | SLC6A2 | 2.373419 | ENSG00000173926 | 115123 | PWB-iPSC DN | MARCHF3 | -0.823113425 | ENSG00000266524 | 2662 | PWB-EC DN | GDF10 | -7.939609 |
| ENSG00000197437 | 441933 | PWB-iPSC UP | OR13G1 | 2.373394 | ENSG00000178078 | 55620 | PWB-iPSC DN | STAP2 | -0.823436262 | ENSG00000043355 | 7546 | PWB-EC DN | ZIC2 | -7.943627 |
| ENSG00000226686 | 101927667 | PWB-iPSC UP | LINC01535 | 2.354086 | ENSG00000105701 | 23770 | PWB-iPSC DN | FKBP8 | -0.824810302 | ENSG00000130193 | 51337 | PWB-EC DN | THEM6 | -7.954109 |
| ENSG00000103522 | 50615 | PWB-iPSC UP | IL21R | 2.341609 | ENSG00000103160 | 83693 | PWB-iPSC DN | HSDL1 | -0.825086801 | ENSG00000115648 | 79083 | PWB-EC DN | MLPH | -8.003183 |
| ENSG00000163810 | 7047 | PWB-iPSC UP | TGM4 | 2.30618 | ENSG00000102802 | 84935 | PWB-iPSC DN | MEDAG | -0.826894527 | ENSG00000186340 | 7058 | PWB-EC DN | THBS2 | -8.101366 |
| ENSG00000221970 | 346528 | PWB-iPSC UP | OR2A1 | 2.279414 | ENSG00000167654 | 85300 | PWB-iPSC DN | ATCAY | -0.830720344 | ENSG00000181449 | 6657 | PWB-EC DN | SOX2 | -8.124795 |
| ENSG00000206609 | 100033423 | PWB-iPSC UP | SNORD116-11 | 2.278033 | ENSG00000187210 | 2650 | PWB-iPSC DN | GCNT1 | -0.830858394 | ENSG00000108821 | 1277 | PWB-EC DN | COL1A1 | -8.231896 |
| ENSG00000235436 | 442523 | PWB-iPSC UP | DPY19L2P4 | 2.261143 | ENSG00000143603 | 3782 | PWB-iPSC DN | KCNN3 | -0.83129045 | ENSG00000044524 | 2042 | PWB-EC DN | EPHA3 | -8.239504 |
| ENSG00000271079 | 441294 | PWB-iPSC UP | CTAGE15 | 2.247785 | ENSG00000151725 | 79682 | PWB-iPSC DN | CENPU | -0.834167113 | ENSG00000087510 | 7022 | PWB-EC DN | TFAP2C | -8.262976 |
| ENSG00000173401 | 256710 | PWB-iPSC UP | GLIPR1L1 | 2.243351 | ENSG00000106992 | 203 | PWB-iPSC DN | AK1 | -0.834616616 | ENSG00000115461 | 3488 | PWB-EC DN | IGFBP5 | -8.363224 |
| ENSG00000231652 | 101928489 | PWB-iPSC UP | LOC101928489 | 2.240117 | ENSG00000162972 | 79568 | PWB-iPSC DN | MAIP1 | -0.83577532 | ENSG00000160932 | 4061 | PWB-EC DN | LY6E | -8.405012 |
| ENSG00000187922 | 414332 | PWB-iPSC UP | LCN10 | 2.216362 | ENSG00000146278 | 10957 | PWB-iPSC DN | PNRC1 | -0.836942599 | ENSG00000260549 | 4500 | PWB-EC DN | MT1L | -8.493733 |
| ENSG00000225937 | 50652 | PWB-iPSC UP | PCA3 | 2.20454 | ENSG00000176244 | 414149 | PWB-iPSC DN | ACBD7 | -0.838970146 | ENSG00000184486 | 5454 | PWB-EC DN | POU3F2 | -8.540634 |
| ENSG00000163285 | 2565 | PWB-iPSC UP | GABRG1 | 2.199985 | ENSG00000173898 | 6712 | PWB-iPSC DN | SPTBN2 | -0.839838474 | ENSG00000137142 | 347252 | PWB-EC DN | IGFBPL1 | -8.628631 |
| ENSG00000243709 | 10637 | PWB-iPSC UP | LEFTY1 | 2.189227 | ENSG00000120594 | 84898 | PWB-iPSC DN | PLXDC2 | -0.840020299 | ENSG00000145824 | 9547 | PWB-EC DN | CXCL14 | -9.392574 |
| ENSG00000229404 | 170425 | PWB-iPSC UP | LINC00858 | 2.182576 | ENSG00000139722 | 79720 | PWB-iPSC DN | VPS37B | -0.840358905 | ENSG00000198692 | 9086 | PWB-EC DN | EIF1AY | -10.2853 |
| ENSG00000113319 | 5924 | PWB-iPSC UP | RASGRF2 | 2.169281 | ENSG00000142541 | 23521 | PWB-iPSC DN | RPL13A | -0.841651525 | ENSG00000196350 | 100287226 | PWB-MSC UP | ZNF729 | 10.640449 |
| ENSG00000172965 | 541471 | PWB-iPSC UP | MIR4435-2HG | 2.163685 | ENSG00000158623 | 26958 | PWB-iPSC DN | COPG2 | -0.841692002 | ENSG00000268696 | 646864 | PWB-MSC UP | ZNF723 | 9.9493975 |
| ENSG00000116785 | 10878 | PWB-iPSC UP | CFHR3 | 2.160043 | ENSG00000167526 | 6137 | PWB-iPSC DN | RPL13 | -0.841720666 | ENSG00000197360 | 148198 | PWB-MSC UP | ZNF98 | 8.8699038 |
| ENSG00000262999 | 101927131 | PWB-iPSC UP | LOC101927131 | 2.151801 | ENSG00000063660 | 2817 | PWB-iPSC DN | GPC1 | -0.842488814 | ENSG00000213973 | 7652 | PWB-MSC UP | ZNF99 | 8.6996621 |
| ENSG00000255929 | 105369438 | PWB-iPSC UP | LOC105369438 | 2.148867 | ENSG00000277443 | 4082 | PWB-iPSC DN | MARCKS | -0.842631745 | ENSG00000229676 | 57615 | PWB-MSC UP | ZNF492 | 8.4409213 |
| ENSG00000200661 | 100033422 | PWB-iPSC UP | SNORD116-10 | 2.142946 | ENSG00000183044 | 18 | PWB-iPSC DN | ABAT | -0.844212563 | ENSG00000263711 | 400655 | PWB-MSC UP | LINC02864 | 6.7078758 |
| ENSG00000274428 | 115409981 | PWB-iPSC UP | RNVU1-25 | 2.13524 | ENSG00000168476 | 80346 | PWB-iPSC DN | REEP4 | -0.844566759 | ENSG00000214039 | 100190940 | PWB-MSC UP | LINC02418 | 3.9744538 |
| ENSG00000201183 | 101954272 | PWB-iPSC UP | RNVU1-3 | 2.132698 | ENSG00000130055 | 54857 | PWB-iPSC DN | GDPD2 | -0.845995197 | ENSG00000173401 | 256710 | PWB-MSC UP | GLIPR1L1 | 3.86317 |
| ENSG00000163359 | 1293 | PWB-iPSC UP | COL6A3 | 2.130205 | ENSG00000198182 | 84775 | PWB-iPSC DN | ZNF607 | -0.84657892 | ENSG00000227195 | 284801 | PWB-MSC UP | MIR663AHG | 3.6609328 |
| ENSG00000229619 | 401093 | PWB-iPSC UP | MBNL1-AS1 | 2.119199 | ENSG00000083857 | 2195 | PWB-iPSC DN | FAT1 | -0.846750382 | ENSG00000269067 | 388523 | PWB-MSC UP | ZNF728 | 3.6184754 |
| ENSG00000152467 | 284312 | PWB-iPSC UP | ZSCAN1 | 2.118985 | ENSG00000113269 | 55819 | PWB-iPSC DN | RNF130 | -0.846785709 | ENSG00000206159 | 352887 | PWB-MSC UP | GYG2P1 | 3.5995384 |
| ENSG00000181211 | 100127950 | PWB-iPSC UP | HECW1-IT1 | 2.113219 | ENSG00000106211 | 3315 | PWB-iPSC DN | HSPB1 | -0.847829569 | ENSG00000260599 | 101929124 | PWB-MSC UP | LOC101929124 | 3.598479 |
| ENSG00000160321 | 7757 | PWB-iPSC UP | ZNF208 | 2.103232 | ENSG00000078902 | 54472 | PWB-iPSC DN | TOLLIP | -0.847851595 | ENSG00000138823 | 4547 | PWB-MSC UP | MTTP | 3.4339387 |
| ENSG00000196104 | 50859 | PWB-iPSC UP | SPOCK3 | 2.100272 | ENSG00000127220 | 79575 | PWB-iPSC DN | ABHD8 | -0.849024629 | ENSG00000206656 | 100033429 | PWB-MSC UP | SNORD116-17 | 3.3485292 |
| ENSG00000222041 | 112597 | PWB-iPSC UP | CYTOR | 2.091647 | ENSG00000111678 | 113246 | PWB-iPSC DN | C12orf57 | -0.849844997 | ENSG00000248473 | 102577426 | PWB-MSC UP | LINC01962 | 3.2670678 |
| ENSG00000236761 | 643854 | PWB-iPSC UP | CTAGE9 | 2.087488 | ENSG00000160685 | 51043 | PWB-iPSC DN | ZBTB7B | -0.849922906 | ENSG00000243775 | 202459 | PWB-MSC UP | OSTCP1 | 3.2246253 |
| ENSG00000136167 | 3936 | PWB-iPSC UP | LCP1 | 2.076574 | ENSG00000161714 | 113026 | PWB-iPSC DN | PLCD3 | -0.850226844 | ENSG00000283160 | 100616406 | PWB-MSC UP | MIR4521 | 3.1429837 |
| ENSG00000108932 | 9120 | PWB-iPSC UP | SLC16A6 | 2.072639 | ENSG00000169184 | 4330 | PWB-iPSC DN | MN1 | -0.851557792 | ENSG00000113722 | 1044 | PWB-MSC UP | CDX1 | 3.0925078 |
| ENSG00000243323 | 148713 | PWB-iPSC UP | PTPRVP | 2.061147 | ENSG00000213722 | 23564 | PWB-iPSC DN | DDAH2 | -0.852713836 | ENSG00000147724 | 51059 | PWB-MSC UP | FAM135B | 3.0257051 |
| ENSG00000053747 | 3909 | PWB-iPSC UP | LAMA3 | 2.056635 | ENSG00000095587 | 7093 | PWB-iPSC DN | TLL2 | -0.853249332 | ENSG00000197134 | 113835 | PWB-MSC UP | ZNF257 | 2.9982669 |
| ENSG00000277610 | 101954264 | PWB-iPSC UP | RNVU1-4 | 2.047772 | ENSG00000130147 | 23677 | PWB-iPSC DN | SH3BP4 | -0.854436629 | ENSG00000202354 | 6085 | PWB-MSC UP | RNY3 | 2.981773 |
| ENSG00000236051 | 100874212 | PWB-iPSC UP | MYCBP2-AS1 | 2.041092 | ENSG00000269994 | 440173 | PWB-iPSC DN | LOC440173 | -0.854528661 | ENSG00000201321 | 100169760 | PWB-MSC UP | RNA5S9 | 2.9651621 |

| | | | | | | | | | | | | | | |
|---|---|---|---|---|---|---|---|---|---|---|---|---|---|---|
| ENSG00000250302 | 152578 | PWB-iPSC UP | LINC01618 | 2.03706 | ENSG00000175793 | 2810 | PWB-iPSC DN | SFN | -0.855392857 | ENSG00000164379 | 94234 | PWB-MSC UP | FOXQ1 | 2.9626555 |
| ENSG00000261105 | 101927155 | PWB-iPSC UP | LMO7-AS1 | 2.036591 | ENSG00000198816 | 140467 | PWB-iPSC DN | ZNF358 | -0.856280764 | ENSG00000104722 | 4741 | PWB-MSC UP | NEFM | 2.9573533 |
| ENSG00000189398 | 10821 | PWB-iPSC UP | OR7E12P | 2.031985 | ENSG00000175274 | 9537 | PWB-iPSC DN | TP53I11 | -0.859679837 | ENSG00000276380 | 389898 | PWB-MSC UP | UBE2NL | 2.9510822 |
| ENSG00000236581 | 100874241 | PWB-iPSC UP | STARD13-AS | 2.031757 | ENSG00000135480 | 3855 | PWB-iPSC DN | KRT7 | -0.861874605 | ENSG00000167434 | 762 | PWB-MSC UP | CA4 | 2.9400915 |
| ENSG00000251175 | 101929529 | PWB-iPSC UP | GSTCD-AS1 | 2.025559 | ENSG00000170906 | 4696 | PWB-iPSC DN | NDUFA3 | -0.863763736 | ENSG00000233780 | 101930748 | PWB-MSC UP | LINC00367 | 2.8629538 |
| ENSG00000254838 | 387751 | PWB-iPSC UP | GVINP1 | 2.021239 | ENSG00000160325 | 11094 | PWB-iPSC DN | CACFD1 | -0.864105465 | ENSG00000258405 | 147660 | PWB-MSC UP | ZNF578 | 2.823931 |
| ENSG00000146047 | 255626 | PWB-iPSC UP | H2BC1 | 2.018817 | ENSG00000178233 | 441151 | PWB-iPSC DN | TMEM151B | -0.864750385 | ENSG00000213538 | 283102 | PWB-MSC UP | KRT8P41 | 2.77863 |
| ENSG00000269289 | 100505851 | PWB-iPSC UP | LOC100505851 | 2.003046 | ENSG00000154548 | 135295 | PWB-iPSC DN | SRSF12 | -0.864814453 | ENSG00000069482 | 51083 | PWB-MSC UP | GAL | 2.7642119 |
| ENSG00000188124 | 338755 | PWB-iPSC UP | OR2AG2 | 1.995265 | ENSG00000183186 | 126567 | PWB-iPSC DN | C2CD4C | -0.870041348 | ENSG00000120498 | 56159 | PWB-MSC UP | TEX11 | 2.7641089 |
| ENSG00000261934 | 56107 | PWB-iPSC UP | PCDHGA9 | 1.989075 | ENSG00000100842 | 10278 | PWB-iPSC DN | EFS | -0.871693143 | ENSG00000156574 | 4838 | PWB-MSC UP | NODAL | 2.7115447 |
| ENSG00000171217 | 49861 | PWB-iPSC UP | CLDN20 | 1.978372 | ENSG00000166886 | 4665 | PWB-iPSC DN | NAB2 | -0.872282506 | ENSG00000226686 | 101927667 | PWB-MSC UP | LINC01535 | 2.7073025 |
| ENSG00000258548 | 100505967 | PWB-iPSC UP | LINC00645 | 1.944061 | ENSG00000177045 | 147912 | PWB-iPSC DN | SIX5 | -0.873326158 | ENSG00000167780 | 8435 | PWB-MSC UP | SOAT2 | 2.6671195 |
| ENSG00000204956 | 56114 | PWB-iPSC UP | PCDHGA1 | 1.940556 | ENSG00000283093 | 441495 | PWB-iPSC DN | CENPVL2 | -0.874345969 | ENSG00000264940 | 780853 | PWB-MSC UP | SNORD3C | 2.6661118 |
| ENSG00000262884 | 101927911 | PWB-iPSC UP | LOC101927911 | 1.937405 | ENSG00000105290 | 333 | PWB-iPSC DN | APLP1 | -0.874796994 | ENSG00000269028 | 100462981 | PWB-MSC UP | MTRNR2L2 | 2.6565776 |
| ENSG00000230542 | 100359394 | PWB-iPSC UP | LINC00102 | 1.933651 | ENSG00000071246 | 22846 | PWB-iPSC DN | VASH1 | -0.875972425 | ENSG00000187537 | 404785 | PWB-MSC UP | POTEG | 2.6370777 |
| ENSG00000271321 | 340307 | PWB-iPSC UP | CTAGE6 | 1.933221 | ENSG00000148175 | 2040 | PWB-iPSC DN | STOM | -0.876193294 | ENSG00000207187 | 109616960 | PWB-MSC UP | SNORA10B | 2.624742 |
| ENSG00000261175 | 102724344 | PWB-iPSC UP | LINC02188 | 1.925615 | ENSG00000180035 | 197407 | PWB-iPSC DN | ZNF48 | -0.877512073 | ENSG00000277526 | 101927815 | PWB-MSC UP | LOC101927815 | 2.5235114 |
| ENSG00000286575 | 107984265 | PWB-iPSC UP | LOC107984265 | 1.924248 | ENSG00000223591 | 389857 | PWB-iPSC DN | CENPVL1 | -0.879447258 | ENSG00000228224 | 83955 | PWB-MSC UP | NACA4P | 2.519291 |
| ENSG00000246422 | 100505658 | PWB-iPSC UP | DIAPH1-AS1 | 1.919892 | ENSG00000109265 | 57482 | PWB-iPSC DN | CRACD | -0.879768144 | ENSG00000179023 | 127707 | PWB-MSC UP | KLHDC7A | 2.5090717 |
| ENSG00000101448 | 57119 | PWB-iPSC UP | EPPIN | 1.909439 | ENSG00000132640 | 22903 | PWB-iPSC DN | BTBD3 | -0.883147271 | ENSG00000219451 | 222901 | PWB-MSC UP | RPL23P8 | 2.4972311 |
| ENSG00000154316 | 157739 | PWB-iPSC UP | TDH | 1.904173 | ENSG00000116667 | 81563 | PWB-iPSC DN | C1orf21 | -0.883198602 | ENSG00000135502 | 65012 | PWB-MSC UP | SLC26A10 | 2.4963146 |
| ENSG00000197584 | 10242 | PWB-iPSC UP | KCNMB2 | 1.89952 | ENSG00000236552 | 728658 | PWB-iPSC DN | RPL13AP5 | -0.883235838 | ENSG00000230667 | 646817 | PWB-MSC UP | SETSIP | 2.4886497 |
| ENSG00000250582 | 101927659 | PWB-iPSC UP | SMAD1-AS2 | 1.897316 | ENSG00000047249 | 51606 | PWB-iPSC DN | ATP6V1H | -0.88542264 | ENSG00000148702 | 3026 | PWB-MSC UP | HABP2 | 2.473147 |
| ENSG00000253457 | 100507341 | PWB-iPSC UP | SMIM18 | 1.895924 | ENSG00000081087 | 28962 | PWB-iPSC DN | OSTM1 | -0.886573023 | ENSG00000169436 | 169044 | PWB-MSC UP | COL22A1 | 2.4492242 |
| ENSG00000180353 | 3059 | PWB-iPSC UP | HCLS1 | 1.888801 | ENSG00000137691 | 85016 | PWB-iPSC DN | CFAP300 | -0.888073958 | ENSG00000253506 | 342538 | PWB-MSC UP | NACA2 | 2.4069269 |
| ENSG00000226416 | 100133545 | PWB-iPSC UP | MRPL23-AS1 | 1.888368 | ENSG00000115255 | 92840 | PWB-iPSC DN | REEP6 | -0.888671223 | ENSG00000160321 | 7757 | PWB-MSC UP | ZNF208 | 2.4064664 |
| ENSG00000133636 | 4922 | PWB-iPSC UP | NTS | 1.883253 | ENSG00000152229 | 9050 | PWB-iPSC DN | PSTPIP2 | -0.889417333 | ENSG00000188375 | 440093 | PWB-MSC UP | H3-5 | 2.3934658 |
| ENSG00000057468 | 4438 | PWB-iPSC UP | MSH4 | 1.878908 | ENSG00000100906 | 4792 | PWB-iPSC DN | NFKBIA | -0.890069497 | ENSG00000171560 | 2243 | PWB-MSC UP | FGA | 2.3805663 |
| ENSG00000175189 | 3626 | PWB-iPSC UP | INHBC | 1.874035 | ENSG00000215271 | 57594 | PWB-iPSC DN | HOMEZ | -0.890794933 | ENSG00000277918 | 115409983 | PWB-MSC UP | RNVU1-28 | 2.3559571 |
| ENSG00000227115 | 100287225 | PWB-iPSC UP | LINC01630 | 1.87043 | ENSG00000198963 | 6096 | PWB-iPSC DN | RORB | -0.891743618 | ENSG00000212993 | 5462 | PWB-MSC UP | POU5F1B | 2.3517398 |
| ENSG00000275585 | 101929798 | PWB-iPSC UP | LOC101929798 | 1.855427 | ENSG00000021762 | 114879 | PWB-iPSC DN | OSBPL5 | -0.894814343 | ENSG00000201098 | 6084 | PWB-MSC UP | RNY1 | 2.3496033 |
| ENSG00000239704 | 284040 | PWB-iPSC UP | CDRT4 | 1.838237 | ENSG00000198937 | 154467 | PWB-iPSC DN | CCDC167 | -0.895303759 | ENSG00000124233 | 6406 | PWB-MSC UP | SEMG1 | 2.3341888 |
| ENSG00000235885 | 101927661 | PWB-iPSC UP | LOC101927661 | 1.838168 | ENSG00000023902 | 51177 | PWB-iPSC DN | PLEKHO1 | -0.895535474 | ENSG00000104899 | 268 | PWB-MSC UP | AMH | 2.3181623 |
| ENSG00000196187 | 9725 | PWB-iPSC UP | TMEM63A | 1.832433 | ENSG00000082014 | 6604 | PWB-iPSC DN | SMARCD3 | -0.896782045 | ENSG00000185155 | 83881 | PWB-MSC UP | MIXL1 | 2.2793879 |
| ENSG00000111424 | 7421 | PWB-iPSC UP | VDR | 1.823528 | ENSG00000146352 | 134829 | PWB-iPSC DN | CLVS2 | -0.898394373 | ENSG00000234444 | 728927 | PWB-MSC UP | ZNF736 | 2.2765486 |
| ENSG00000207721 | 406962 | PWB-iPSC UP | MIR186 | 1.822746 | ENSG00000153721 | 154043 | PWB-iPSC DN | CNKSR3 | -0.899583943 | ENSG00000188674 | 389073 | PWB-MSC UP | C2orf80 | 2.248989 |
| ENSG00000248810 | 100507639 | PWB-iPSC UP | LINC02432 | 1.820168 | ENSG00000174428 | 389524 | PWB-iPSC DN | GTF2IRD2B | -0.900450566 | ENSG00000204963 | 56141 | PWB-MSC UP | PCDHA7 | 2.2457581 |
| ENSG00000118473 | 84251 | PWB-iPSC UP | SGIP1 | 1.818858 | ENSG00000249395 | 101805492 | PWB-iPSC DN | CASC9 | -0.900528002 | ENSG00000266976 | 102724908 | PWB-MSC UP | LOC102724908 | 2.2187567 |
| ENSG00000240338 | 100506281 | PWB-iPSC UP | LOC100506281 | 1.813149 | ENSG00000181274 | 23401 | PWB-iPSC DN | FRAT2 | -0.901012444 | ENSG00000252316 | 6086 | PWB-MSC UP | RNY4 | 2.2146694 |
| ENSG00000254166 | 103164619 | PWB-iPSC UP | PCAT2 | 1.812657 | ENSG00000105419 | 56917 | PWB-iPSC DN | MEIS3 | -0.901319676 | ENSG00000139540 | 283375 | PWB-MSC UP | SLC39A5 | 2.2093092 |
| ENSG00000235865 | 57000 | PWB-iPSC UP | GSN-AS1 | 1.79621 | ENSG00000106571 | 2737 | PWB-iPSC DN | GLI3 | -0.901373824 | ENSG00000248546 | 23520 | PWB-MSC UP | ANP32C | 2.2080496 |
| ENSG00000125551 | 5342 | PWB-iPSC UP | PLGLB2 | 1.78814 | ENSG00000135709 | 9764 | PWB-iPSC DN | KIAA0513 | -0.90250056 | ENSG00000221676 | 100151684 | PWB-MSC UP | RNU6ATAC | 2.200918 |
| ENSG00000206262 | 401089 | PWB-iPSC UP | FOXL2NB | 1.784202 | ENSG00000171132 | 5581 | PWB-iPSC DN | PRKCE | -0.904602728 | ENSG00000205436 | 91828 | PWB-MSC UP | EXOC3L4 | 2.197781 |
| ENSG00000204965 | 56143 | PWB-iPSC UP | PCDHA5 | 1.783389 | ENSG00000272886 | 55802 | PWB-iPSC DN | DCP1A | -0.904613624 | ENSG00000107742 | 9806 | PWB-MSC UP | SPOCK2 | 2.167972 |
| ENSG00000130612 | 22952 | PWB-iPSC UP | CYP2G1P | 1.782919 | ENSG00000108819 | 84687 | PWB-iPSC DN | PPP1R9B | -0.905726595 | ENSG00000146001 | 54660 | PWB-MSC UP | PCDHB18P | 2.1535457 |
| ENSG00000173124 | 142827 | PWB-iPSC UP | ACSM6 | 1.777046 | ENSG00000125652 | 84266 | PWB-iPSC DN | ALKBH7 | -0.905936506 | ENSG00000272674 | 57717 | PWB-MSC UP | PCDHB16 | 2.1255292 |
| ENSG00000236296 | 441046 | PWB-iPSC UP | GUSBP5 | 1.776203 | ENSG00000205213 | 55366 | PWB-iPSC DN | LGR4 | -0.909218961 | ENSG00000198573 | 64663 | PWB-MSC UP | SPANXC | 2.1027732 |
| ENSG00000203907 | 441161 | PWB-iPSC UP | OOEP | 1.773941 | ENSG00000171150 | 9655 | PWB-iPSC DN | SOCS5 | -0.912859424 | ENSG00000207008 | 677833 | PWB-MSC UP | SNORA54 | 2.0988231 |
| ENSG00000169752 | 145957 | PWB-iPSC UP | NRG4 | 1.768411 | ENSG00000144655 | 64651 | PWB-iPSC DN | CSRNP1 | -0.913648587 | ENSG00000167588 | 2819 | PWB-MSC UP | GPD1 | 2.0329995 |
| ENSG00000207340 | 101954273 | PWB-iPSC UP | RNVU1-1 | 1.762879 | ENSG00000161682 | 284069 | PWB-iPSC DN | FAM171A2 | -0.914152895 | ENSG00000207344 | 109616965 | PWB-MSC UP | SNORA22C | 2.0243354 |
| ENSG00000167578 | 53916 | PWB-iPSC UP | RAB4B | 1.761254 | ENSG00000165895 | 143872 | PWB-iPSC DN | ARHGAP42 | -0.914776699 | ENSG00000154342 | 89780 | PWB-MSC UP | WNT3A | 2.0195031 |
| ENSG00000278709 | 105416157 | PWB-iPSC UP | NKILA | 1.759686 | ENSG00000058799 | 54432 | PWB-iPSC DN | YIPF1 | -0.917193946 | ENSG00000283475 | 103504734 | PWB-MSC UP | MIR1244-4 | 2.018694 |
| ENSG00000255084 | 101928896 | PWB-iPSC UP | LOC101928896 | 1.759412 | ENSG00000183778 | 10317 | PWB-iPSC DN | B3GALT5 | -0.918000428 | ENSG00000131746 | 84951 | PWB-MSC UP | TNS4 | 2.0178546 |
| ENSG00000167895 | 147138 | PWB-iPSC UP | TMC8 | 1.752839 | ENSG00000147324 | 9258 | PWB-iPSC DN | MFHAS1 | -0.920379221 | ENSG00000270722 | 115482718 | PWB-MSC UP | RNVU1-31 | 2.0174547 |
| ENSG00000259120 | 100130933 | PWB-iPSC UP | SMIM6 | 1.752103 | ENSG00000101144 | 655 | PWB-iPSC DN | BMP7 | -0.922741673 | ENSG00000200156 | 26832 | PWB-MSC UP | RNU5B-1 | 2.0156847 |
| ENSG00000261121 | 107986266 | PWB-iPSC UP | LINC02473 | 1.750146 | ENSG00000099203 | 11018 | PWB-iPSC DN | TMED1 | -0.922905109 | ENSG00000227234 | 728695 | PWB-MSC UP | SPANXB1 | 2.0153099 |
| ENSG00000223482 | 728190 | PWB-iPSC UP | NUTM2A-AS1 | 1.734789 | ENSG00000131116 | 126299 | PWB-iPSC DN | ZNF428 | -0.924934758 | ENSG00000120328 | 56124 | PWB-MSC UP | PCDHB12 | 2.012739 |
| ENSG00000232803 | 100127888 | PWB-iPSC UP | SLCO4A1-AS1 | 1.728585 | ENSG00000072958 | 8907 | PWB-iPSC DN | AP1M1 | -0.925429161 | ENSG00000255378 | 100133251 | PWB-MSC UP | PRR23D2 | 1.9933607 |
| ENSG00000229867 | 100874111 | PWB-iPSC UP | STEAP3-AS1 | 1.728122 | ENSG00000051128 | 9454 | PWB-iPSC DN | HOMER3 | -0.925662467 | ENSG00000255251 | 100131608 | PWB-MSC UP | PRR23D1 | 1.99336 |
| ENSG00000256690 | 105369332 | PWB-iPSC UP | LOC105369332 | 1.727441 | ENSG00000117461 | 8503 | PWB-iPSC DN | PIK3R3 | -0.926036717 | ENSG00000180658 | 79541 | PWB-MSC UP | OR2A4 | 1.9885397 |
| ENSG00000253485 | 56110 | PWB-iPSC UP | PCDHGA5 | 1.717722 | ENSG00000123146 | 976 | PWB-iPSC DN | ADGRE5 | -0.927331543 | ENSG00000207191 | 100033417 | PWB-MSC UP | SNORD116-5 | 1.9852933 |
| ENSG00000171084 | 100125556 | PWB-iPSC UP | FAM86JP | 1.71718 | ENSG00000171223 | 3726 | PWB-iPSC DN | JUNB | -0.928306551 | ENSG00000199568 | 26831 | PWB-MSC UP | RNU5A-1 | 1.9755475 |
| ENSG00000145020 | 275 | PWB-iPSC UP | AMT | 1.704939 | ENSG00000084710 | 22979 | PWB-iPSC DN | EFR3B | -0.929215155 | ENSG00000257084 | 105369635 | PWB-MSC UP | MIR200CHG | 1.9605114 |
| ENSG00000198633 | 147658 | PWB-iPSC UP | ZNF534 | 1.703435 | ENSG00000115239 | 51130 | PWB-iPSC DN | ASB3 | -0.935427645 | ENSG00000178401 | 79962 | PWB-MSC UP | DNAJC22 | 1.9572274 |
| ENSG00000226891 | 101927084 | PWB-iPSC UP | LINC01359 | 1.697249 | ENSG00000033170 | 2530 | PWB-iPSC DN | FUT8 | -0.936434912 | ENSG00000184860 | 93517 | PWB-MSC UP | SDR42E1 | 1.9564197 |
| ENSG00000163492 | 285025 | PWB-iPSC UP | CCDC141 | 1.694875 | ENSG00000155966 | 2334 | PWB-iPSC DN | AFF2 | -0.938082764 | ENSG00000251898 | 677780 | PWB-MSC UP | SCARNA11 | 1.9385893 |
| ENSG00000251287 | 644974 | PWB-iPSC UP | ALG1L2 | 1.687204 | ENSG00000249992 | 25907 | PWB-iPSC DN | TMEM158 | -0.938111033 | ENSG00000162391 | 338094 | PWB-MSC UP | FAM151A | 1.9117828 |
| ENSG00000281731 | 101929741 | PWB-iPSC UP | METTL14-DT | 1.680866 | ENSG00000171126 | 170850 | PWB-iPSC DN | KCNG3 | -0.939591792 | ENSG00000100505 | 114088 | PWB-MSC UP | TRIM9 | 1.9030728 |
| ENSG00000125965 | 8200 | PWB-iPSC UP | GDF5 | 1.680492 | ENSG00000139174 | 144165 | PWB-iPSC DN | PRICKLE1 | -0.9401662 | ENSG00000243896 | 401427 | PWB-MSC UP | OR2A7 | 1.8951571 |
| ENSG00000282556 | 101927825 | PWB-iPSC UP | LOC101927825 | 1.677146 | ENSG00000103449 | 6299 | PWB-iPSC DN | SALL1 | -0.94161085 | ENSG00000104435 | 11075 | PWB-MSC UP | STMN2 | 1.8947811 |
| ENSG00000125872 | 164312 | PWB-iPSC UP | LRRN4 | 1.675309 | ENSG00000207389 | 6060 | PWB-iPSC DN | RNU1-4 | -0.94250722 | ENSG00000084674 | 338 | PWB-MSC UP | APOB | 1.8936274 |
| ENSG00000286177 | 105372401 | PWB-iPSC UP | LOC105372401 | 1.674407 | ENSG00000135333 | 2045 | PWB-iPSC DN | EPHA7 | -0.943266579 | ENSG00000188906 | 120892 | PWB-MSC UP | LRRK2 | 1.8888841 |
| ENSG00000164398 | 23305 | PWB-iPSC UP | ACSL6 | 1.674352 | ENSG00000207005 | 26870 | PWB-iPSC DN | RNU1-2 | -0.943434861 | ENSG00000207313 | 677794 | PWB-MSC UP | SNORA2B | 1.8805483 |
| ENSG00000162399 | 7809 | PWB-iPSC UP | BSND | 1.671513 | ENSG00000273768 | 115409989 | PWB-iPSC DN | RNVU1-29 | -0.945070346 | ENSG00000136011 | 55576 | PWB-MSC UP | STAB2 | 1.8790443 |
| ENSG00000180332 | 386618 | PWB-iPSC UP | KCTD4 | 1.669898 | ENSG00000163344 | 10654 | PWB-iPSC DN | PMVK | -0.945355313 | ENSG00000206633 | 100302743 | PWB-MSC UP | SNORA80B | 1.8669953 |
| ENSG00000134538 | 10599 | PWB-iPSC UP | SLCO1B1 | 1.656313 | ENSG00000122335 | 84947 | PWB-iPSC DN | SERAC1 | -0.945703461 | ENSG00000237872 | 645682 | PWB-MSC UP | POU5F1P4 | 1.8549125 |
| ENSG00000124701 | 10930 | PWB-iPSC UP | APOBEC2 | 1.654695 | ENSG00000136960 | 5168 | PWB-iPSC DN | ENPP2 | -0.946479904 | ENSG00000278318 | 7772 | PWB-MSC UP | ZNF229 | 1.8454588 |
| ENSG00000242193 | 730102 | PWB-iPSC UP | CRYZL2P | 1.650477 | ENSG00000127399 | 65999 | PWB-iPSC DN | LRRC61 | -0.946624066 | ENSG00000127399 | 65999 | PWB-MSC UP | LRRC61 | 1.8177599 |
| ENSG00000259583 | 101927751 | PWB-iPSC UP | LOC101927751 | 1.640518 | ENSG00000174469 | 26047 | PWB-iPSC DN | CNTNAP2 | -0.947423331 | ENSG00000158825 | 978 | PWB-MSC UP | CDA | 1.8104438 |
| ENSG00000234883 | 114614 | PWB-iPSC UP | MIR155HG | 1.638836 | ENSG00000206652 | 26871 | PWB-iPSC DN | RNU1-1 | -0.948373889 | ENSG00000255622 | 54661 | PWB-MSC UP | PCDHB17P | 1.8081175 |
| ENSG00000286071 | 105373170 | PWB-iPSC UP | LOC105373170 | 1.63213 | ENSG00000207513 | 26869 | PWB-iPSC DN | RNU1-3 | -0.949207639 | ENSG00000206612 | 677793 | PWB-MSC UP | SNORA2A | 1.7922681 |
| ENSG00000258667 | 105370526 | PWB-iPSC UP | HIF1A-AS3 | 1.629379 | ENSG00000206737 | 26863 | PWB-iPSC DN | RNVU1-18 | -0.951253947 | ENSG00000213412 | 728643 | PWB-MSC UP | HNRNPA1P33 | 1.7914612 |
| ENSG00000118785 | 6696 | PWB-iPSC UP | SPP1 | 1.611435 | ENSG00000139971 | 145407 | PWB-iPSC DN | ARMH4 | -0.951844287 | ENSG00000081277 | 5317 | PWB-MSC UP | PKP1 | 1.7763587 |
| ENSG00000229311 | 101927827 | PWB-iPSC UP | LOC101927827 | 1.609948 | ENSG00000007255 | 79090 | PWB-iPSC DN | TRAPPC6A | -0.95210828 | ENSG00000197635 | 1803 | PWB-MSC UP | DPP4 | 1.7565049 |
| ENSG00000132639 | 6616 | PWB-iPSC UP | SNAP25 | 1.609221 | ENSG00000206585 | 26864 | PWB-iPSC DN | RNVU1-7 | -0.952362367 | ENSG00000260389 | 441818 | PWB-MSC UP | WBP11P1 | 1.7497753 |
| ENSG00000141052 | 93649 | PWB-iPSC UP | MYOCD | 1.60893 | ENSG00000134775 | 80206 | PWB-iPSC DN | FHOD3 | -0.954774481 | ENSG00000125787 | 2797 | PWB-MSC UP | GNRH2 | 1.7149675 |
| ENSG00000197085 | 404744 | PWB-iPSC UP | NPSR1-AS1 | 1.605468 | ENSG00000101347 | 25939 | PWB-iPSC DN | SAMHD1 | -0.955174726 | ENSG00000153162 | 654 | PWB-MSC UP | BMP6 | 1.7069493 |
| ENSG00000205890 | 100128770 | PWB-iPSC UP | LOC100128770 | 1.604806 | ENSG00000169071 | 4920 | PWB-iPSC DN | ROR2 | -0.955535149 | ENSG00000151790 | 6999 | PWB-MSC UP | TDO2 | 1.7014081 |
| ENSG00000156574 | 4838 | PWB-iPSC UP | NODAL | 1.602917 | ENSG00000050438 | 9498 | PWB-iPSC DN | SLC4A8 | -0.956524997 | ENSG00000132677 | 57127 | PWB-MSC UP | RHBG | 1.700175 |
| ENSG00000247473 | 100852407 | PWB-iPSC UP | CARS-AS1 | 1.602282 | ENSG00000189171 | 6284 | PWB-iPSC DN | S100A13 | -0.957136002 | ENSG00000271079 | 441294 | PWB-MSC UP | CTAGE15 | 1.6944308 |
| ENSG00000176928 | 51301 | PWB-iPSC UP | GCNT4 | 1.60095 | ENSG00000125266 | 1948 | PWB-iPSC DN | EFNB2 | -0.958028757 | ENSG00000139219 | 1280 | PWB-MSC UP | COL2A1 | 1.69349 |
| ENSG00000231851 | 101927492 | PWB-iPSC UP | MRPS9-AS1 | 1.600276 | ENSG00000115226 | 64838 | PWB-iPSC DN | FNDC4 | -0.958695454 | ENSG00000164647 | 26872 | PWB-MSC UP | STEAP1 | 1.6881114 |
| ENSG00000156076 | 11197 | PWB-iPSC UP | WIF1 | 1.598686 | ENSG00000117280 | 8934 | PWB-iPSC DN | RAB29 | -0.958875541 | ENSG00000206634 | 677807 | PWB-MSC UP | SNORA22 | 1.6855095 |
| ENSG00000204869 | 444882 | PWB-iPSC UP | IGFL4 | 1.595494 | ENSG00000110492 | 4192 | PWB-iPSC DN | MDK | -0.959519571 | ENSG00000198028 | 147741 | PWB-MSC UP | ZNF560 | 1.6830773 |
| ENSG00000258932 | 728755 | PWB-iPSC UP | LOC728755 | 1.59485 | ENSG00000118503 | 7128 | PWB-iPSC DN | TNFAIP3 | -0.961167322 | ENSG00000184344 | 9573 | PWB-MSC UP | GDF3 | 1.6703606 |
| ENSG00000249835 | 105379054 | PWB-iPSC UP | VCAN-AS1 | 1.594077 | ENSG00000108679 | 3959 | PWB-iPSC DN | LGALS3BP | -0.962157204 | ENSG00000208892 | 677829 | PWB-MSC UP | SNORA49 | 1.6684061 |
| ENSG00000173295 | 286042 | PWB-iPSC UP | FAM86B3P | 1.59094 | ENSG00000087116 | 9509 | PWB-iPSC DN | ADAMTS2 | -0.963689487 | ENSG00000265480 | 284085 | PWB-MSC UP | KRT18P55 | 1.6661592 |
| ENSG00000162241 | 283130 | PWB-iPSC UP | SLC25A45 | 1.580716 | ENSG00000105642 | 3780 | PWB-iPSC DN | KCNN1 | -0.963879073 | ENSG00000231389 | 3113 | PWB-MSC UP | HLA-DPA1 | 1.6659251 |
| ENSG00000260735 | 100505915 | PWB-iPSC UP | LOC100505915 | 1.578223 | ENSG00000213420 | 221914 | PWB-iPSC DN | GPC2 | -0.966932884 | ENSG00000259706 | 7190 | PWB-MSC UP | HSP90B2P | 1.6659094 |
| ENSG00000236064 | 101928335 | PWB-iPSC UP | LOC101928335 | 1.577225 | ENSG00000153093 | 55289 | PWB-iPSC DN | ACOXL | -0.967930879 | ENSG00000223572 | 548596 | PWB-MSC UP | CKMT1A | 1.6652673 |
| ENSG00000234444 | 728927 | PWB-iPSC UP | ZNF736 | 1.576101 | ENSG00000179222 | 9500 | PWB-iPSC DN | MAGED1 | -0.968809202 | ENSG00000139344 | 144193 | PWB-MSC UP | AMDHD1 | 1.6622401 |
| ENSG00000287299 | 101927741 | PWB-iPSC UP | LOC101927741 | 1.575325 | ENSG00000007376 | 113000 | PWB-iPSC DN | RPUSD1 | -0.970820103 | ENSG00000175697 | 165829 | PWB-MSC UP | GPR156 | 1.6572208 |
| ENSG00000250365 | 101927124 | PWB-iPSC UP | LOC101927124 | 1.573536 | ENSG00000205189 | 65986 | PWB-iPSC DN | ZBTB10 | -0.97309462 | ENSG00000239039 | 692084 | PWB-MSC UP | SNORD13 | 1.6552752 |
| ENSG00000117600 | 9890 | PWB-iPSC UP | PLPPR4 | 1.571351 | ENSG00000099864 | 5064 | PWB-iPSC DN | PALM | -0.973168293 | ENSG00000179412 | 101060301 | PWB-MSC UP | HNRNPCL4 | 1.6494889 |
| ENSG00000270011 | 100529215 | PWB-iPSC UP | ZNF559-ZNF177 | 1.565568 | ENSG00000119711 | 4329 | PWB-iPSC DN | ALDH6A1 | -0.974234101 | ENSG00000135454 | 2583 | PWB-MSC UP | B4GALNT1 | 1.6421462 |
| ENSG00000125414 | 4620 | PWB-iPSC UP | MYH2 | 1.564091 | ENSG00000204366 | 221527 | PWB-iPSC DN | ZBTB12 | -0.979559254 | ENSG00000233384 | 102723817 | PWB-MSC UP | CNIH3-AS2 | 1.6417382 |
| ENSG00000225868 | 100631378 | PWB-iPSC UP | LOC100631378 | 1.547337 | ENSG00000180616 | 6752 | PWB-iPSC DN | SSTR2 | -0.979860983 | ENSG00000120322 | 56128 | PWB-MSC UP | PCDHB8 | 1.6413164 |

| | | | | | | | | | | | | | | | | |
|---|---|---|---|---|---|---|---|---|---|---|---|---|---|---|---|---|
| ENSG00000159905 | 7638 | PWB-iPSC UP | ZNF221 | 1.546926 | | ENSG00000182013 | 55228 | PWB-iPSC DN | PNMA8A | -0.983032802 | | ENSG00000189334 | 57402 | PWB-MSC UP | S100A14 | 1.6382292 |
| ENSG00000228727 | 401251 | PWB-iPSC UP | SAPCD1 | 1.544423 | | ENSG00000198939 | 80108 | PWB-iPSC DN | ZFP2 | -0.986594647 | | ENSG00000008735 | 23542 | PWB-MSC UP | MAPK8IP2 | 1.6370771 |
| ENSG00000235927 | 374987 | PWB-iPSC UP | NEXN-AS1 | 1.542493 | | ENSG00000135736 | 92922 | PWB-iPSC DN | CCDC102A | -0.988759018 | | ENSG00000110944 | 51561 | PWB-MSC UP | IL23A | 1.6242269 |
| ENSG00000105974 | 857 | PWB-iPSC UP | CAV1 | 1.541909 | | ENSG00000141576 | 114804 | PWB-iPSC DN | RNF157 | -0.993809063 | | ENSG00000197870 | 5544 | PWB-MSC UP | PRB3 | 1.6217967 |
| ENSG00000205181 | 149837 | PWB-iPSC UP | LINC00654 | 1.539807 | | ENSG00000140455 | 9960 | PWB-iPSC DN | USP3 | -0.993902858 | | ENSG00000155265 | 401647 | PWB-MSC UP | GOLGA7B | 1.6201561 |
| ENSG00000154736 | 11096 | PWB-iPSC UP | ADAMTS5 | 1.530345 | | ENSG00000171016 | 26108 | PWB-iPSC DN | PYGO1 | -0.995643278 | | ENSG00000050327 | 7984 | PWB-MSC UP | ARHGEF5 | 1.6200285 |
| ENSG00000248445 | 101927233 | PWB-iPSC UP | SEMA6A-AS1 | 1.530039 | | ENSG00000180336 | 284071 | PWB-iPSC DN | MEIOC | -0.997119766 | | ENSG00000185974 | 6011 | PWB-MSC UP | GRK1 | 1.609412 |
| ENSG00000184343 | 26576 | PWB-iPSC UP | SRPK3 | 1.529021 | | ENSG00000091136 | 3912 | PWB-iPSC DN | LAMB1 | -0.997879307 | | ENSG00000101276 | 113278 | PWB-MSC UP | SLC52A3 | 1.6061284 |
| ENSG00000237399 | 100507034 | PWB-iPSC UP | PITRM1-AS1 | 1.524787 | | ENSG00000152518 | 678 | PWB-iPSC DN | ZFP36L2 | -1.003193857 | | ENSG00000259516 | 723972 | PWB-MSC UP | ANP32AP1 | 1.6001715 |
| ENSG00000126790 | 112849 | PWB-iPSC UP | L3HYPDH | 1.523504 | | ENSG00000166710 | 567 | PWB-iPSC DN | B2M | -1.003599264 | | ENSG00000182866 | 3932 | PWB-MSC UP | LCK | 1.5996372 |
| ENSG00000284874 | 100526833 | PWB-iPSC UP | SEPT5-GP1BB | 1.51944 | | ENSG00000091129 | 4897 | PWB-iPSC DN | NRCAM | -1.005926174 | | ENSG00000168505 | 2637 | PWB-MSC UP | GBX2 | 1.590353 |
| ENSG00000041515 | 23026 | PWB-iPSC UP | MYO16 | 1.516211 | | ENSG00000152193 | 79596 | PWB-iPSC DN | OBI1 | -1.006935738 | | ENSG00000271321 | 340307 | PWB-MSC UP | CTAGE6 | 1.5895743 |
| ENSG00000145416 | 55016 | PWB-iPSC UP | MARCHF1 | 1.515364 | | ENSG00000105278 | 23217 | PWB-iPSC DN | ZFR2 | -1.007752146 | | ENSG00000136928 | 9568 | PWB-MSC UP | GABBR2 | 1.5885538 |
| ENSG00000171722 | 284680 | PWB-iPSC UP | SPATA46 | 1.509804 | | ENSG00000131203 | 3620 | PWB-iPSC DN | IDO1 | -1.008269911 | | ENSG00000261175 | 102724344 | PWB-MSC UP | LINC02188 | 1.5877896 |
| ENSG00000145700 | 256006 | PWB-iPSC UP | ANKRD31 | 1.508099 | | ENSG00000067113 | 8611 | PWB-iPSC DN | PLPP1 | -1.008829556 | | ENSG00000205846 | 93978 | PWB-MSC UP | CLEC6A | 1.575613 |
| ENSG00000232490 | 100874206 | PWB-iPSC UP | OSBPL10-AS1 | 1.505515 | | ENSG00000214530 | 10809 | PWB-iPSC DN | STARD10 | -1.009692195 | | ENSG00000237289 | 1159 | PWB-MSC UP | CKMT1B | 1.5690548 |
| ENSG00000186645 | 102723849 | PWB-iPSC UP | SPDYE17 | 1.502731 | | ENSG00000178585 | 56998 | PWB-iPSC DN | CTNNBIP1 | -1.010633757 | | ENSG00000053108 | 23105 | PWB-MSC UP | FSTL4 | 1.5680339 |
| ENSG00000236008 | 101929567 | PWB-iPSC UP | LINC01814 | 1.50273 | | ENSG00000251322 | 85358 | PWB-iPSC DN | SHANK3 | -1.011699786 | | ENSG00000165556 | 1045 | PWB-MSC UP | CDX2 | 1.5628919 |
| ENSG00000177483 | 375316 | PWB-iPSC UP | RBM44 | 1.50241 | | ENSG00000114854 | 7134 | PWB-iPSC DN | TNNC1 | -1.011770286 | | ENSG00000184363 | 11187 | PWB-MSC UP | PKP3 | 1.5525303 |
| ENSG00000121211 | 84057 | PWB-iPSC UP | MND1 | 1.496126 | | ENSG00000167815 | 7001 | PWB-iPSC DN | PRDX2 | -1.011842461 | | ENSG00000199436 | 692053 | PWB-MSC UP | SNORD9 | 1.5524373 |
| ENSG00000205038 | 93035 | PWB-iPSC UP | PKHD1L1 | 1.494436 | | ENSG00000108830 | 8153 | PWB-iPSC DN | RND2 | -1.014361361 | | ENSG00000165025 | 6850 | PWB-MSC UP | SYK | 1.53216 |
| ENSG00000172460 | 124221 | PWB-iPSC UP | PRSS30P | 1.489949 | | ENSG00000141854 | 113230 | PWB-iPSC DN | MISP3 | -1.015126631 | | ENSG00000171631 | 5031 | PWB-MSC UP | P2RY6 | 1.5319321 |
| ENSG00000110455 | 84680 | PWB-iPSC UP | ACCS | 1.48744 | | ENSG00000176595 | 9920 | PWB-iPSC DN | KBTBD11 | -1.016483145 | | ENSG00000225602 | 100873935 | PWB-MSC UP | MTOR-AS1 | 1.5181548 |
| ENSG00000213967 | 730087 | PWB-iPSC UP | ZNF726 | 1.486681 | | ENSG00000170561 | 153572 | PWB-iPSC DN | IRX2 | -1.017529992 | | ENSG00000159166 | 3898 | PWB-MSC UP | LAD1 | 1.5114277 |
| ENSG00000162620 | 127255 | PWB-iPSC UP | LRRIQ3 | 1.48519 | | ENSG00000152953 | 55351 | PWB-iPSC DN | STK32B | -1.018498737 | | ENSG00000081853 | 56113 | PWB-MSC UP | PCDHGA2 | 1.5097301 |
| ENSG00000136928 | 9568 | PWB-iPSC UP | GABBR2 | 1.484041 | | ENSG00000176171 | 664 | PWB-iPSC DN | BNIP3 | -1.019115404 | | ENSG00000236761 | 643854 | PWB-MSC UP | CTAGE9 | 1.5069517 |
| ENSG00000147869 | 9350 | PWB-iPSC UP | CER1 | 1.472728 | | ENSG00000167716 | 124997 | PWB-iPSC DN | WDR81 | -1.019632232 | | ENSG00000134873 | 9071 | PWB-MSC UP | CLDN10 | 1.5043833 |
| ENSG00000050327 | 7984 | PWB-iPSC UP | ARHGEF5 | 1.472383 | | ENSG00000158966 | 57685 | PWB-iPSC DN | CACHD1 | -1.020848064 | | ENSG00000200879 | 85391 | PWB-MSC UP | SNORD14E | 1.4997608 |
| ENSG00000232265 | 100505824 | PWB-iPSC UP | LINC02805 | 1.472328 | | ENSG00000146700 | 136853 | PWB-iPSC DN | SSC4D | -1.021217805 | | ENSG00000259717 | 105370683 | PWB-MSC UP | LINC00677 | 1.4855503 |
| ENSG00000138641 | 8916 | PWB-iPSC UP | HERC3 | 1.472193 | | ENSG00000144857 | 91653 | PWB-iPSC DN | BOC | -1.02194835 | | ENSG00000135127 | 92558 | PWB-MSC UP | BICDL1 | 1.4757164 |
| ENSG00000250266 | 101928223 | PWB-iPSC UP | LINC01612 | 1.466403 | | ENSG00000185112 | 131583 | PWB-iPSC DN | FAM43A | -1.026741737 | | ENSG00000251669 | 348926 | PWB-MSC UP | FAM86EP | 1.4737231 |
| ENSG00000245534 | 101928784 | PWB-iPSC UP | RORA-AS1 | 1.464427 | | ENSG00000105289 | 27134 | PWB-iPSC DN | TJP3 | -1.028193186 | | ENSG00000185304 | 729857 | PWB-MSC UP | RGPD2 | 1.4689987 |
| ENSG00000236991 | 101927983 | PWB-iPSC UP | EDRF1-AS1 | 1.460666 | | ENSG00000154822 | 23228 | PWB-iPSC DN | PLCL2 | -1.032761104 | | ENSG00000166482 | 4239 | PWB-MSC UP | MFAP4 | 1.4587804 |
| ENSG00000155657 | 7273 | PWB-iPSC UP | TTN | 1.458075 | | ENSG00000095932 | 284422 | PWB-iPSC DN | SMIM24 | -1.033746249 | | ENSG00000142494 | 55244 | PWB-MSC UP | SLC47A1 | 1.4558008 |
| ENSG00000262209 | 56102 | PWB-iPSC UP | PCDHGB3 | 1.456565 | | ENSG00000108557 | 10743 | PWB-iPSC DN | RAI1 | -1.033946537 | | ENSG00000207088 | 677797 | PWB-MSC UP | SNORA7B | 1.4538926 |
| ENSG00000283982 | 101929563 | PWB-iPSC UP | LOC101929563 | 1.455098 | | ENSG00000286190 | 728392 | PWB-iPSC DN | LOC728392 | -1.034110983 | | ENSG00000225868 | 100631378 | PWB-MSC UP | LOC100631378 | 1.4492816 |
| ENSG00000187699 | 84281 | PWB-iPSC UP | C2orf88 | 1.453282 | | ENSG00000128311 | 7263 | PWB-iPSC DN | TST | -1.036188122 | | ENSG00000234498 | 387841 | PWB-MSC UP | RPL13AP20 | 1.4417221 |
| ENSG00000259881 | 101927793 | PWB-iPSC UP | LOC101927793 | 1.44881 | | ENSG00000132589 | 2319 | PWB-iPSC DN | FLOT2 | -1.036654794 | | ENSG00000276476 | 100506622 | PWB-MSC UP | LINC00540 | 1.4391666 |
| ENSG00000197934 | 100996571 | PWB-iPSC UP | CYYR1-AS1 | 1.44764 | | ENSG00000129946 | 25759 | PWB-iPSC DN | SHC2 | -1.040205752 | | ENSG00000183801 | 283298 | PWB-MSC UP | OLFML1 | 1.4357733 |
| ENSG00000188869 | 342125 | PWB-iPSC UP | TMC3 | 1.447077 | | ENSG00000241697 | 8577 | PWB-iPSC DN | TMEFF1 | -1.042496095 | | ENSG00000122863 | 9469 | PWB-MSC UP | CHST3 | 1.4351702 |
| ENSG00000203999 | 284751 | PWB-iPSC UP | LINC01270 | 1.445946 | | ENSG00000164651 | 221833 | PWB-iPSC DN | SP8 | -1.049335566 | | ENSG00000113319 | 5924 | PWB-MSC UP | RASGRF2 | 1.4249447 |
| ENSG00000262576 | 56111 | PWB-iPSC UP | PCDHGA4 | 1.442555 | | ENSG00000103490 | 29108 | PWB-iPSC DN | PYCARD | -1.049338189 | | ENSG00000139155 | 53919 | PWB-MSC UP | SLCO1C1 | 1.4177929 |
| ENSG00000232229 | 643529 | PWB-iPSC UP | LINC00865 | 1.438957 | | ENSG00000166165 | 1152 | PWB-iPSC DN | CKB | -1.049908525 | | ENSG00000179172 | 343069 | PWB-MSC UP | HNRNPCL1 | 1.416334 |
| ENSG00000189136 | 388165 | PWB-iPSC UP | UBE2Q2P1 | 1.437035 | | ENSG00000185909 | 200942 | PWB-iPSC DN | KLHDC8B | -1.050241654 | | ENSG00000182489 | 402415 | PWB-MSC UP | XKRX | 1.4157786 |
| ENSG00000253230 | 157627 | PWB-iPSC UP | LINC00599 | 1.436219 | | ENSG00000137078 | 27240 | PWB-iPSC DN | SIT1 | -1.051442279 | | ENSG00000187627 | 400966 | PWB-MSC UP | RGPD1 | 1.4107236 |
| ENSG00000112212 | 222642 | PWB-iPSC UP | TSPO2 | 1.434751 | | ENSG00000183971 | 283869 | PWB-iPSC DN | NPW | -1.054245298 | | ENSG00000177181 | 284716 | PWB-MSC UP | RIMKLA | 1.4105863 |
| ENSG00000197753 | 222662 | PWB-iPSC UP | LHFPL5 | 1.432431 | | ENSG00000105559 | 57664 | PWB-iPSC DN | PLEKHA4 | -1.055871243 | | ENSG00000110876 | 6404 | PWB-MSC UP | SELPLG | 1.4098188 |
| ENSG00000106686 | 55064 | PWB-iPSC UP | SPATA6L | 1.42978 | | ENSG00000148123 | 54886 | PWB-iPSC DN | PLPPR1 | -1.055943717 | | ENSG00000204956 | 56114 | PWB-MSC UP | PCDHGA1 | 1.4089807 |
| ENSG00000167785 | 148156 | PWB-iPSC UP | ZNF558 | 1.427665 | | ENSG00000130707 | 445 | PWB-iPSC DN | ASS1 | -1.057149312 | | ENSG00000105974 | 857 | PWB-MSC UP | CAV1 | 1.4066933 |
| ENSG00000064787 | 8537 | PWB-iPSC UP | BCAS1 | 1.426986 | | ENSG00000152779 | 387700 | PWB-iPSC DN | SLC16A12 | -1.05717793 | | ENSG00000165794 | 29986 | PWB-MSC UP | SLC39A2 | 1.4066788 |
| ENSG00000189099 | 345062 | PWB-iPSC UP | PRSS48 | 1.426943 | | ENSG00000043355 | 7546 | PWB-iPSC DN | ZIC2 | -1.05762944 | | ENSG00000148600 | 92211 | PWB-MSC UP | CDHR1 | 1.4021519 |
| ENSG00000244161 | 105377105 | PWB-iPSC UP | FLNB-AS1 | 1.426806 | | ENSG00000124126 | 57580 | PWB-iPSC DN | PREX1 | -1.059730915 | | ENSG00000166426 | 1381 | PWB-MSC UP | CRABP1 | 1.4015813 |
| ENSG00000228592 | 266917 | PWB-iPSC UP | D21S2088E | 1.425086 | | ENSG00000126218 | 2159 | PWB-iPSC DN | F10 | -1.060737855 | | ENSG00000173262 | 144195 | PWB-MSC UP | SLC2A14 | 1.3966266 |
| ENSG00000174600 | 1240 | PWB-iPSC UP | CMKLR1 | 1.415939 | | ENSG00000169862 | 1501 | PWB-iPSC DN | CTNND2 | -1.061700016 | | ENSG00000118137 | 335 | PWB-MSC UP | APOA1 | 1.392961 |
| ENSG00000240280 | 146771 | PWB-iPSC UP | TCAM1P | 1.41511 | | ENSG00000147872 | 123 | PWB-iPSC DN | PLIN2 | -1.063831095 | | ENSG00000146215 | 401262 | PWB-MSC UP | CRIP3 | 1.3882307 |
| ENSG00000255152 | 100532732 | PWB-iPSC UP | MSH5-SAPCD1 | 1.412199 | | ENSG00000198822 | 2913 | PWB-iPSC DN | GRM3 | -1.065885158 | | ENSG00000105143 | 6511 | PWB-MSC UP | SLC1A6 | 1.3864119 |
| ENSG00000204740 | 340895 | PWB-iPSC UP | MALRD1 | 1.411231 | | ENSG00000066032 | 1496 | PWB-iPSC DN | CTNNA2 | -1.069254057 | | ENSG00000255408 | 56145 | PWB-MSC UP | PCDHA3 | 1.3835444 |
| ENSG00000232788 | 101929947 | PWB-iPSC UP | ITGA6-AS1 | 1.407857 | | ENSG00000113763 | 90249 | PWB-iPSC DN | UNC5A | -1.071136341 | | ENSG00000127084 | 89846 | PWB-MSC UP | FGD3 | 1.3745833 |
| ENSG00000238062 | 348761 | PWB-iPSC UP | SPATA3-AS1 | 1.407574 | | ENSG00000184500 | 5627 | PWB-iPSC DN | PROS1 | -1.072167287 | | ENSG00000115758 | 4953 | PWB-MSC UP | ODC1 | 1.3705741 |
| ENSG00000256988 | 105369614 | PWB-iPSC UP | LOC105369614 | 1.405902 | | ENSG00000115318 | 84695 | PWB-iPSC DN | LOXL3 | -1.073630986 | | ENSG00000235602 | 642559 | PWB-MSC UP | POU5F1P3 | 1.358205 |
| ENSG00000058866 | 1608 | PWB-iPSC UP | DGKG | 1.40533 | | ENSG00000129116 | 23022 | PWB-iPSC DN | PALLD | -1.075316611 | | ENSG00000182107 | 161291 | PWB-MSC UP | TMEM30B | 1.3572234 |
| ENSG00000214940 | 101059953 | PWB-iPSC UP | NPIPA8 | 1.404635 | | ENSG00000197019 | 29950 | PWB-iPSC DN | SERTAD1 | -1.076788586 | | ENSG00000255337 | 101928424 | PWB-MSC UP | LOC101928424 | 1.3562972 |
| ENSG00000205592 | 283463 | PWB-iPSC UP | MUC19 | 1.404266 | | ENSG00000182175 | 56963 | PWB-iPSC DN | RGMA | -1.078057807 | | ENSG00000113212 | 56129 | PWB-MSC UP | PCDHB7 | 1.3559463 |
| ENSG00000184465 | 253769 | PWB-iPSC UP | WDR27 | 1.403458 | | ENSG00000158764 | 142683 | PWB-iPSC DN | ITLN2 | -1.078500724 | | ENSG00000215769 | 109286553 | PWB-MSC UP | ARHGAP27P1-BF | 1.3493399 |
| ENSG00000203326 | 170958 | PWB-iPSC UP | ZNF525 | 1.401841 | | ENSG00000181449 | 6657 | PWB-iPSC DN | SOX2 | -1.083419919 | | ENSG00000120899 | 2185 | PWB-MSC UP | PTK2B | 1.3472186 |
| ENSG00000165617 | 51339 | PWB-iPSC UP | DACT1 | 1.399589 | | ENSG00000151090 | 7068 | PWB-iPSC DN | THRB | -1.085599067 | | ENSG00000162975 | 3754 | PWB-MSC UP | KCNF1 | 1.3393348 |
| ENSG00000178440 | 113218477 | PWB-iPSC UP | LINC00843 | 1.396674 | | ENSG00000137142 | 347252 | PWB-iPSC DN | IGFBPL1 | -1.08636551 | | ENSG00000131019 | 79465 | PWB-MSC UP | ULBP3 | 1.3373623 |
| ENSG00000187833 | 388960 | PWB-iPSC UP | C2orf78 | 1.395728 | | ENSG00000163545 | 81788 | PWB-iPSC DN | NUAK2 | -1.088985175 | | ENSG00000139767 | 84530 | PWB-MSC UP | SRRM4 | 1.3355553 |
| ENSG00000250116 | 100506142 | PWB-iPSC UP | LOC100506142 | 1.3952 | | ENSG00000244694 | 442213 | PWB-iPSC DN | PTCHD4 | -1.092171066 | | ENSG00000170439 | 196410 | PWB-MSC UP | METTL7B | 1.3216791 |
| ENSG00000184672 | 138046 | PWB-iPSC UP | RALYL | 1.393949 | | ENSG00000166863 | 6866 | PWB-iPSC DN | TAC3 | -1.09329291 | | ENSG00000143786 | 149111 | PWB-MSC UP | CNIH3 | 1.3093637 |
| ENSG00000184434 | 64922 | PWB-iPSC UP | LRRC19 | 1.393432 | | ENSG00000243449 | 401115 | PWB-iPSC DN | C4orf48 | -1.097820747 | | ENSG00000197905 | 7004 | PWB-MSC UP | TEAD4 | 1.3073091 |
| ENSG00000170356 | 401428 | PWB-iPSC UP | OR2A20P | 1.390933 | | ENSG00000143502 | 55061 | PWB-iPSC DN | SUSD4 | -1.098988897 | | ENSG00000221740 | 692210 | PWB-MSC UP | SNORD93 | 1.30629 |
| ENSG00000172215 | 10663 | PWB-iPSC UP | CXCR6 | 1.3897 | | ENSG00000162496 | 9249 | PWB-iPSC DN | DHRS3 | -1.099271367 | | ENSG00000223482 | 728190 | PWB-MSC UP | NUTM2A-AS1 | 1.3019692 |
| ENSG00000157765 | 10568 | PWB-iPSC UP | SLC34A2 | 1.389129 | | ENSG00000115325 | 1796 | PWB-iPSC DN | DOK1 | -1.102909969 | | ENSG00000110921 | 4598 | PWB-MSC UP | MVK | 1.3018583 |
| ENSG00000121380 | 79370 | PWB-iPSC UP | BCL2L14 | 1.388821 | | ENSG00000119185 | 9270 | PWB-iPSC DN | ITGB1BP1 | -1.103490924 | | ENSG00000228570 | 283008 | PWB-MSC UP | NUTM2E | 1.2980042 |
| ENSG00000054179 | 954 | PWB-iPSC UP | ENTPD2 | 1.386462 | | ENSG00000205542 | 7114 | PWB-iPSC DN | TMSB4X | -1.104011773 | | ENSG00000200169 | 26830 | PWB-MSC UP | RNU5D-1 | 1.2967304 |
| ENSG00000181798 | 151477 | PWB-iPSC UP | LINC00471 | 1.383988 | | ENSG00000150625 | 2823 | PWB-iPSC DN | GPM6A | -1.105658429 | | ENSG00000185818 | 339983 | PWB-MSC UP | NAT8L | 1.2956715 |
| ENSG00000283050 | 101926918 | PWB-iPSC UP | GTF2IP12 | 1.383563 | | ENSG00000197705 | 57565 | PWB-iPSC DN | KLHL14 | -1.106256768 | | ENSG00000165105 | 158158 | PWB-MSC UP | RASEF | 1.2921197 |
| ENSG00000258525 | 100506071 | PWB-iPSC UP | LOC100506071 | 1.377913 | | ENSG00000185668 | 5453 | PWB-iPSC DN | POU3F1 | -1.106315004 | | ENSG00000151136 | 121551 | PWB-MSC UP | BTBD11 | 1.2910209 |
| ENSG00000226314 | 651302 | PWB-iPSC UP | ZNF192P1 | 1.375248 | | ENSG00000237923 | 105375014 | PWB-iPSC DN | LINC02570 | -1.107234134 | | ENSG00000202400 | 25826 | PWB-MSC UP | SNORD82 | 1.2878902 |
| ENSG00000223972 | 100287102 | PWB-iPSC UP | DDX11L1 | 1.372144 | | ENSG00000185664 | 6490 | PWB-iPSC DN | PMEL | -1.108787901 | | ENSG00000228695 | 51716 | PWB-MSC UP | CES1P1 | 1.2866872 |
| ENSG00000224597 | 102724316 | PWB-iPSC UP | SVIL-AS1 | 1.369501 | | ENSG00000174007 | 84984 | PWB-iPSC DN | CEP19 | -1.10881771 | | ENSG00000168243 | 2786 | PWB-MSC UP | GNG4 | 1.2805622 |
| ENSG00000273173 | 8926 | PWB-iPSC UP | SNURF | 1.368434 | | ENSG00000185567 | 113146 | PWB-iPSC DN | AHNAK2 | -1.10911473 | | ENSG00000254221 | 56104 | PWB-MSC UP | PCDHGB1 | 1.2719792 |
| ENSG00000101977 | 4168 | PWB-iPSC UP | MCF2 | 1.367676 | | ENSG00000123240 | 10133 | PWB-iPSC DN | OPTN | -1.109847352 | | ENSG00000167555 | 84436 | PWB-MSC UP | ZNF528 | 1.2703323 |
| ENSG00000286058 | 440905 | PWB-iPSC UP | FAR2P1 | 1.365555 | | ENSG00000173281 | 79660 | PWB-iPSC DN | PPP1R3B | -1.110653399 | | ENSG00000188211 | 374383 | PWB-MSC UP | NCR3LG1 | 1.2648965 |
| ENSG00000270392 | 767846 | PWB-iPSC UP | PFN1P2 | 1.363282 | | ENSG00000250510 | 27239 | PWB-iPSC DN | GPR162 | -1.11520344 | | ENSG00000104112 | 29106 | PWB-MSC UP | SCG3 | 1.2640851 |
| ENSG00000214029 | 101060200 | PWB-iPSC UP | ZNF891 | 1.358039 | | ENSG00000133216 | 2048 | PWB-iPSC DN | EPHB2 | -1.115245483 | | ENSG00000123405 | 4778 | PWB-MSC UP | NFE2 | 1.2610452 |
| ENSG00000233393 | 105369306 | PWB-iPSC UP | LOC105369306 | 1.356469 | | ENSG00000234745 | 3106 | PWB-iPSC DN | HLA-B | -1.118239531 | | ENSG00000207233 | 677819 | PWB-MSC UP | SNORA37 | 1.2602235 |
| ENSG00000165046 | 137994 | PWB-iPSC UP | LETM2 | 1.354011 | | ENSG00000129951 | 79948 | PWB-iPSC DN | PLPPR3 | -1.118716969 | | ENSG00000130829 | 1852 | PWB-MSC UP | DUSP9 | 1.2561069 |
| ENSG00000162630 | 8707 | PWB-iPSC UP | B3GALT2 | 1.353645 | | ENSG00000174099 | 253827 | PWB-iPSC DN | MSRB3 | -1.119607205 | | ENSG00000212283 | 692205 | PWB-MSC UP | SNORD89 | 1.2530439 |
| ENSG00000236383 | 100874261 | PWB-iPSC UP | CCDC200 | 1.351103 | | ENSG00000183067 | 150084 | PWB-iPSC DN | IGSF5 | -1.123042664 | | ENSG00000200394 | 100124536 | PWB-MSC UP | SNORA38B | 1.2471552 |
| ENSG00000188916 | 642938 | PWB-iPSC UP | INSYN2A | 1.3409 | | ENSG00000141570 | 57332 | PWB-iPSC DN | CBX8 | -1.126379496 | | ENSG00000111077 | 23371 | PWB-MSC UP | TNS2 | 1.2465347 |
| ENSG00000273520 | 102723849 | PWB-iPSC UP | SPDYE17 | 1.339967 | | ENSG00000256463 | 27164 | PWB-iPSC DN | SALL3 | -1.128225442 | | ENSG00000112812 | 10279 | PWB-MSC UP | PRSS16 | 1.2406042 |
| ENSG00000277942 | 102465874 | PWB-iPSC UP | MIR8075 | 1.338389 | | ENSG00000179604 | 23580 | PWB-iPSC DN | CDC42EP4 | -1.129124428 | | ENSG00000276547 | 56101 | PWB-MSC UP | PCDHGB5 | 1.2401818 |
| ENSG00000235488 | 100506681 | PWB-iPSC UP | JARID2-AS1 | 1.338037 | | ENSG00000184897 | 8971 | PWB-iPSC DN | H1-10 | -1.130051742 | | ENSG00000201302 | 26783 | PWB-MSC UP | SNORA65 | 1.2316766 |
| ENSG00000269834 | 102724105 | PWB-iPSC UP | ZNF528-AS1 | 1.336454 | | ENSG00000196358 | 84628 | PWB-iPSC DN | NTNG2 | -1.130401581 | | ENSG00000184613 | 4753 | PWB-MSC UP | NELL2 | 1.2233012 |
| ENSG00000171951 | 7857 | PWB-iPSC UP | SCG2 | 1.335763 | | ENSG00000121900 | 113452 | PWB-iPSC DN | TMEM54 | -1.13043717 | | ENSG00000076555 | 32 | PWB-MSC UP | ACACB | 1.2218845 |
| ENSG00000153823 | 55022 | PWB-iPSC UP | PID1 | 1.334905 | | ENSG00000120708 | 7045 | PWB-iPSC DN | TGFBI | -1.131762839 | | ENSG00000111319 | 6337 | PWB-MSC UP | SCNN1A | 1.2215694 |
| ENSG00000042781 | 7399 | PWB-iPSC UP | USH2A | 1.333995 | | ENSG00000121966 | 7852 | PWB-iPSC DN | CXCR4 | -1.133637495 | | ENSG00000197410 | 54798 | PWB-MSC UP | DCHS2 | 1.2205358 |
| ENSG00000204623 | 80862 | PWB-iPSC UP | ZNRD1ASP | 1.333247 | | ENSG00000058335 | 5923 | PWB-iPSC DN | RASGRF1 | -1.135341667 | | ENSG00000179059 | 132625 | PWB-MSC UP | ZFP42 | 1.2191428 |
| ENSG00000197959 | 26052 | PWB-iPSC UP | DNM3 | 1.332626 | | ENSG00000198795 | 25925 | PWB-iPSC DN | ZNF521 | -1.135474816 | | ENSG00000163686 | 57406 | PWB-MSC UP | ABHD6 | 1.2185375 |
| ENSG00000230782 | 340512 | PWB-iPSC UP | LOC340512 | 1.332591 | | ENSG00000214336 | 344167 | PWB-iPSC DN | FOXI3 | -1.138040628 | | ENSG00000147383 | 50814 | PWB-MSC UP | NSDHL | 1.2139774 |
| ENSG00000197816 | 100499483 | PWB-iPSC UP | CCDC180 | 1.330352 | | ENSG00000135750 | 3775 | PWB-iPSC DN | KCNK1 | -1.139046823 | | ENSG00000110455 | 84680 | PWB-MSC UP | ACCS | 1.2083065 |
| ENSG00000168234 | 125488 | PWB-iPSC UP | TTC39C | 1.33015 | | ENSG00000105855 | 3696 | PWB-iPSC DN | ITGB8 | -1.139223203 | | ENSG00000268095 | 101928571 | PWB-MSC UP | ZNF649-AS1 | 1.2042467 |
| ENSG00000184144 | 6900 | PWB-iPSC UP | CNTN2 | 1.330068 | | ENSG00000105894 | 5764 | PWB-iPSC DN | PTN | -1.140110338 | | ENSG00000124302 | 64377 | PWB-MSC UP | CHST8 | 1.2026245 |
| ENSG00000145975 | 222826 | PWB-iPSC UP | FAM217A | 1.32969 | | ENSG00000106853 | 22949 | PWB-iPSC DN | PTGR1 | -1.144150312 | | ENSG00000167767 | 144501 | PWB-MSC UP | KRT80 | 1.1949152 |
| ENSG00000204323 | 643008 | PWB-iPSC UP | SMIM5 | 1.327264 | | ENSG00000102547 | 81617 | PWB-iPSC DN | CAB39L | -1.144447357 | | ENSG00000145623 | 9180 | PWB-MSC UP | OSMR | 1.1941799 |

| | | | | | | | | | | | | | | |
|---|---|---|---|---|---|---|---|---|---|---|---|---|---|---|
| ENSG00000237940 | 102723927 | PWB-iPSC UP | LINC01238 | 1.322747 | ENSG00000176485 | 11145 | PWB-iPSC DN | PLAAT3 | -1.152985871 | ENSG00000200795 | 26835 | PWB-MSC UP | RNU4-1 | 1.1868772 |
| ENSG00000251432 | 100507487 | PWB-iPSC UP | LINC02615 | 1.322312 | ENSG00000183844 | 54097 | PWB-iPSC DN | FAM3B | -1.154336568 | ENSG00000108950 | 54757 | PWB-MSC UP | FAM20A | 1.1854327 |
| ENSG00000146477 | 6581 | PWB-iPSC UP | SLC22A3 | 1.322276 | ENSG00000197956 | 6277 | PWB-iPSC DN | S100A6 | -1.155793141 | ENSG00000200792 | 677846 | PWB-MSC UP | SNORA80A | 1.1845011 |
| ENSG00000159640 | 1636 | PWB-iPSC UP | ACE | 1.321845 | ENSG00000164488 | 168002 | PWB-iPSC DN | DACT2 | -1.163198009 | ENSG00000172817 | 9420 | PWB-MSC UP | CYP7B1 | 1.1843772 |
| ENSG00000103888 | 57214 | PWB-iPSC UP | CEMIP | 1.319809 | ENSG00000140937 | 1009 | PWB-iPSC DN | CDH11 | -1.172067695 | ENSG00000146070 | 7941 | PWB-MSC UP | PLA2G7 | 1.182987 |
| ENSG00000183535 | 378832 | PWB-iPSC UP | COL18A1-AS1 | 1.31885 | ENSG00000180592 | 387640 | PWB-iPSC DN | SKIDA1 | -1.174519219 | ENSG00000184845 | 1812 | PWB-MSC UP | DRD1 | 1.1771299 |
| ENSG00000135899 | 3431 | PWB-iPSC UP | SP110 | 1.316284 | ENSG00000196177 | 36 | PWB-iPSC DN | ACADSB | -1.175936941 | ENSG00000214562 | 728130 | PWB-MSC UP | NUTM2D | 1.1767945 |
| ENSG00000118432 | 1268 | PWB-iPSC UP | CNR1 | 1.315657 | ENSG00000158825 | 978 | PWB-iPSC DN | CDA | -1.178191373 | ENSG00000179178 | 128218 | PWB-MSC UP | TMEM125 | 1.1695218 |
| ENSG00000262461 | 102723849 | PWB-iPSC UP | SPDYE17 | 1.313744 | ENSG00000114315 | 3280 | PWB-iPSC DN | HES1 | -1.182346333 | ENSG00000200816 | 677820 | PWB-MSC UP | SNORA38 | 1.1685335 |
| ENSG00000260279 | 105371414 | PWB-iPSC UP | LOC105371414 | 1.311931 | ENSG00000184012 | 7113 | PWB-iPSC DN | TMPRSS2 | -1.18450196 | ENSG00000207340 | 101954273 | PWB-MSC UP | RNVU1-1 | 1.1677701 |
| ENSG00000280136 | 101929823 | PWB-iPSC UP | LOC101929823 | 1.310998 | ENSG00000196116 | 23424 | PWB-iPSC DN | TDRD7 | -1.186701748 | ENSG00000150990 | 57647 | PWB-MSC UP | DHX37 | 1.1676262 |
| ENSG00000256594 | 374443 | PWB-iPSC UP | LOC374443 | 1.310612 | ENSG00000274333 | 102724219 | PWB-iPSC DN | LOC10272421 | -1.186960361 | ENSG00000266402 | 105376843 | PWB-MSC UP | SNHG25 | 1.1658303 |
| ENSG00000152822 | 2911 | PWB-iPSC UP | GRM1 | 1.310336 | ENSG00000182575 | 11248 | PWB-iPSC DN | NXPH3 | -1.189346894 | ENSG00000197444 | 55753 | PWB-MSC UP | OGDHL | 1.1630018 |
| ENSG00000136449 | 84073 | PWB-iPSC UP | MYCBPAP | 1.308347 | ENSG00000131771 | 84152 | PWB-iPSC DN | PPP1R1B | -1.189587332 | ENSG00000111641 | 4839 | PWB-MSC UP | NOP2 | 1.1533242 |
| ENSG00000205089 | 645121 | PWB-iPSC UP | CCNI2 | 1.307918 | ENSG00000173673 | 390992 | PWB-iPSC DN | HES3 | -1.190848453 | ENSG00000162746 | 127943 | PWB-MSC UP | FCRLB | 1.1463236 |
| ENSG00000246477 | 101929290 | PWB-iPSC UP | LOC101929290 | 1.307672 | ENSG00000062282 | 84649 | PWB-iPSC DN | DGAT2 | -1.193214971 | ENSG00000140563 | 55784 | PWB-MSC UP | MCTP2 | 1.1442714 |
| ENSG00000232995 | 8490 | PWB-iPSC UP | RGS5 | 1.307646 | ENSG00000170801 | 94031 | PWB-iPSC DN | HTRA3 | -1.206503602 | ENSG00000212443 | 677832 | PWB-MSC UP | SNORA53 | 1.1398071 |
| ENSG00000008226 | 9940 | PWB-iPSC UP | DLEC1 | 1.303826 | ENSG00000142794 | 84224 | PWB-iPSC DN | NBPF3 | -1.207290586 | ENSG00000172818 | 5017 | PWB-MSC UP | OVOL1 | 1.1367453 |
| ENSG00000254221 | 56104 | PWB-iPSC UP | PCDHGB1 | 1.302782 | ENSG00000256713 | 5222 | PWB-iPSC DN | PGA5 | -1.213158781 | ENSG00000208772 | 692225 | PWB-MSC UP | SNORD94 | 1.1352791 |
| ENSG00000104889 | 10535 | PWB-iPSC UP | RNASEH2A | 1.2982 | ENSG00000277991 | 102723360 | PWB-iPSC DN | LOC10272336 | -1.214384042 | ENSG00000233718 | 10408 | PWB-MSC UP | MYCNOS | 1.132606 |
| ENSG00000267871 | 105372476 | PWB-iPSC UP | ZNF460-AS1 | 1.297139 | ENSG00000173093 | 160762 | PWB-iPSC DN | CCDC63 | -1.216910969 | ENSG00000147408 | 55790 | PWB-MSC UP | CSGALNACT1 | 1.1310548 |
| ENSG00000204963 | 56141 | PWB-iPSC UP | PCDHA7 | 1.296465 | ENSG00000203952 | 347475 | PWB-iPSC DN | CCDC160 | -1.217131644 | ENSG00000184564 | 84189 | PWB-MSC UP | SLITRK6 | 1.1290332 |
| ENSG00000124019 | 79843 | PWB-iPSC UP | FAM124B | 1.296405 | ENSG00000115266 | 10297 | PWB-iPSC DN | APC2 | -1.219422439 | ENSG00000157557 | 2114 | PWB-MSC UP | ETS2 | 1.1246928 |
| ENSG00000172209 | 2845 | PWB-iPSC UP | GPR22 | 1.294269 | ENSG00000197467 | 1305 | PWB-iPSC DN | COL13A1 | -1.221249032 | ENSG00000158856 | 2039 | PWB-MSC UP | DMTN | 1.1234975 |
| ENSG00000267470 | 100507433 | PWB-iPSC UP | ZNF571-AS1 | 1.291469 | ENSG00000157240 | 8321 | PWB-iPSC DN | FZD1 | -1.221472508 | ENSG00000132846 | 84327 | PWB-MSC UP | ZBED3 | 1.1223808 |
| ENSG00000048540 | 55885 | PWB-iPSC UP | LMO3 | 1.288774 | ENSG00000109099 | 5376 | PWB-iPSC DN | PMP22 | -1.223309483 | ENSG00000130182 | 84891 | PWB-MSC UP | ZSCAN10 | 1.118 |
| ENSG00000196167 | 399948 | PWB-iPSC UP | COLCA1 | 1.288103 | ENSG00000105251 | 56961 | PWB-iPSC DN | SHD | -1.225090952 | ENSG00000232677 | 100506930 | PWB-MSC UP | LINC00665 | 1.1166226 |
| ENSG00000172403 | 171024 | PWB-iPSC UP | SYNPO2 | 1.288003 | ENSG00000171004 | 90161 | PWB-iPSC DN | HS6ST2 | -1.227897905 | ENSG00000233087 | 150786 | PWB-MSC UP | RAB6D | 1.1157654 |
| ENSG00000253910 | 56103 | PWB-iPSC UP | PCDHGB2 | 1.287907 | ENSG00000168672 | 157638 | PWB-iPSC DN | LRATD2 | -1.230931129 | ENSG00000178882 | 144347 | PWB-MSC UP | RFLNA | 1.1069854 |
| ENSG00000189350 | 165186 | PWB-iPSC UP | TOGARAM2 | 1.287821 | ENSG00000248098 | 593 | PWB-iPSC DN | BCKDHA | -1.231884462 | ENSG00000162496 | 9249 | PWB-MSC UP | DHRS3 | 1.104812 |
| ENSG00000198786 | 4540 | PWB-iPSC UP | ND5 | 1.285334 | ENSG00000095321 | 1384 | PWB-iPSC DN | CRAT | -1.23596416 | ENSG00000196132 | 4661 | PWB-MSC UP | MYT1 | 1.1022955 |
| ENSG00000255920 | 103752584 | PWB-iPSC UP | CCND2-AS1 | 1.285155 | ENSG00000085185 | 63035 | PWB-iPSC DN | BCORL1 | -1.23827714 | ENSG00000111261 | 54682 | PWB-MSC UP | MANSC1 | 1.1009734 |
| ENSG00000198626 | 6262 | PWB-iPSC UP | RYR2 | 1.280434 | ENSG00000188483 | 389792 | PWB-iPSC DN | IER5L | -1.239107375 | ENSG00000171084 | 100125556 | PWB-MSC UP | FAM86JP | 1.1003395 |
| ENSG00000248449 | 56120 | PWB-iPSC UP | PCDHGB8P | 1.274553 | ENSG00000125398 | 6662 | PWB-iPSC DN | SOX9 | -1.240022812 | ENSG00000202515 | 56662 | PWB-MSC UP | VTRNA1-3 | 1.097145 |
| ENSG00000185304 | 729857 | PWB-iPSC UP | RGPD2 | 1.272277 | ENSG00000103154 | 54550 | PWB-iPSC DN | NECAB2 | -1.240899554 | ENSG00000144645 | 114884 | PWB-MSC UP | OSBPL10 | 1.0923255 |
| ENSG00000132872 | 6860 | PWB-iPSC UP | SYT4 | 1.268764 | ENSG00000271133 | 101927811 | PWB-iPSC DN | LOC10192781 | -1.241679606 | ENSG00000081760 | 65985 | PWB-MSC UP | AACS | 1.091436 |
| ENSG00000253767 | 9708 | PWB-iPSC UP | PCDHGA8 | 1.268667 | ENSG00000162174 | 80150 | PWB-iPSC DN | ASRGL1 | -1.245923332 | ENSG00000224597 | 102724316 | PWB-MSC UP | SVIL-AS1 | 1.0880645 |
| ENSG00000132297 | 10086 | PWB-iPSC UP | HHLA1 | 1.268052 | ENSG00000141639 | 5596 | PWB-iPSC DN | MAPK4 | -1.250642841 | ENSG00000228649 | 109729180 | PWB-MSC UP | SNHG26 | 1.0871379 |
| ENSG00000169116 | 25849 | PWB-iPSC UP | PARM1 | 1.267575 | ENSG00000104332 | 6422 | PWB-iPSC DN | SFRP1 | -1.252665554 | ENSG00000179101 | 115482706 | PWB-MSC UP | H3P47 | 1.0848329 |
| ENSG00000197410 | 54798 | PWB-iPSC UP | DCHS2 | 1.267136 | ENSG00000177098 | 6330 | PWB-iPSC DN | SCN4B | -1.25441772 | ENSG00000223865 | 3115 | PWB-MSC UP | HLA-DPB1 | 1.0816194 |
| ENSG00000189367 | 9729 | PWB-iPSC UP | KIAA0408 | 1.265827 | ENSG00000189184 | 54510 | PWB-iPSC DN | PCDH18 | -1.254652589 | ENSG00000166801 | 63901 | PWB-MSC UP | FAM111A | 1.0787443 |
| ENSG00000174501 | 400986 | PWB-iPSC UP | ANKRD36C | 1.264972 | ENSG00000163251 | 7855 | PWB-iPSC DN | FZD5 | -1.260880932 | ENSG00000143469 | 255928 | PWB-MSC UP | SYT14 | 1.0731682 |
| ENSG00000138316 | 140766 | PWB-iPSC UP | ADAMTS14 | 1.261221 | ENSG00000165949 | 3429 | PWB-iPSC DN | IFI27 | -1.261318833 | ENSG00000113209 | 26167 | PWB-MSC UP | PCDHB5 | 1.071608 |
| ENSG00000177788 | 105373159 | PWB-iPSC UP | LOC105373159 | 1.253593 | ENSG00000182580 | 2049 | PWB-iPSC DN | EPHB3 | -1.26169046 | ENSG00000233654 | 105376748 | PWB-MSC UP | LOC105376748 | 1.0669737 |
| ENSG00000173157 | 80070 | PWB-iPSC UP | ADAMTS20 | 1.25247 | ENSG00000128606 | 10234 | PWB-iPSC DN | LRRC17 | -1.262763794 | ENSG00000233673 | 100286979 | PWB-MSC UP | ANAPC1P1 | 1.0640409 |
| ENSG00000268235 | 100996631 | PWB-iPSC UP | TCP11X1 | 1.250921 | ENSG00000161103 | 102725072 | PWB-iPSC DN | LOC10272507 | -1.26685139 | ENSG00000155465 | 9056 | PWB-MSC UP | SLC7A7 | 1.0639856 |
| ENSG00000275291 | 115409982 | PWB-iPSC UP | RNVU1-26 | 1.247436 | ENSG00000149256 | 26011 | PWB-iPSC DN | TENM4 | -1.269485408 | ENSG00000037897 | 4234 | PWB-MSC UP | METTL1 | 1.0626453 |
| ENSG00000271270 | 100507032 | PWB-iPSC UP | TMCC1-AS1 | 1.247322 | ENSG00000219451 | 222901 | PWB-iPSC DN | RPL23P8 | -1.269551768 | ENSG00000055732 | 55283 | PWB-MSC UP | MCOLN3 | 1.0572987 |
| ENSG00000136404 | 53346 | PWB-iPSC UP | TM6SF1 | 1.24488 | ENSG00000165194 | 57526 | PWB-iPSC DN | PCDH19 | -1.272972947 | ENSG00000005471 | 5244 | PWB-MSC UP | ABCB4 | 1.0539761 |
| ENSG00000274210 | 115409988 | PWB-iPSC UP | RNVU1-27 | 1.244689 | ENSG00000100604 | 1113 | PWB-iPSC DN | CHGA | -1.277536577 | ENSG00000103196 | 83716 | PWB-MSC UP | CRISPLD2 | 1.0509235 |
| ENSG00000162631 | 22854 | PWB-iPSC UP | NTNG1 | 1.234501 | ENSG00000179796 | 116135 | PWB-iPSC DN | LRRC3B | -1.278811878 | ENSG00000121380 | 79370 | PWB-MSC UP | BCL2L14 | 1.0439796 |
| ENSG00000071967 | 79901 | PWB-iPSC UP | CYBRD1 | 1.234057 | ENSG00000188505 | 342897 | PWB-iPSC DN | NCCRP1 | -1.286359454 | ENSG00000262074 | 780852 | PWB-MSC UP | SNORD3B-2 | 1.0436161 |
| ENSG00000170827 | 1057 | PWB-iPSC UP | CELP | 1.231922 | ENSG00000169851 | 5099 | PWB-iPSC DN | PCDH7 | -1.289556465 | ENSG00000177192 | 80324 | PWB-MSC UP | PUS1 | 1.0423558 |
| ENSG00000138193 | 51196 | PWB-iPSC UP | PLCE1 | 1.231554 | ENSG00000152092 | 460 | PWB-iPSC DN | ASTN1 | -1.290732649 | ENSG00000276170 | 101929494 | PWB-MSC UP | LOC101929494 | 1.041048 |
| ENSG00000159173 | 7135 | PWB-iPSC UP | TNNI1 | 1.231045 | ENSG00000158859 | 9507 | PWB-iPSC DN | ADAMTS4 | -1.296335747 | ENSG00000013503 | 55703 | PWB-MSC UP | POLR3B | 1.035469 |
| ENSG00000278099 | 6063 | PWB-iPSC UP | RNVU1-2A | 1.230628 | ENSG00000095752 | 3589 | PWB-iPSC DN | IL11 | -1.29723972 | ENSG00000112294 | 7915 | PWB-MSC UP | ALDH5A1 | 1.0329681 |
| ENSG00000276547 | 56101 | PWB-iPSC UP | PCDHGB5 | 1.229886 | ENSG00000087076 | 51171 | PWB-iPSC DN | HSD17B14 | -1.298468807 | ENSG00000256232 | 101929027 | PWB-MSC UP | LINC02387 | 1.0292615 |
| ENSG00000225037 | 100874078 | PWB-iPSC UP | EIF1AX-AS1 | 1.229465 | ENSG00000165323 | 120114 | PWB-iPSC DN | FAT3 | -1.298866024 | ENSG00000265185 | 26851 | PWB-MSC UP | SNORD3B-1 | 1.0285656 |
| ENSG00000216031 | 100126296 | PWB-iPSC UP | MIR298 | 1.228804 | ENSG00000101210 | 1917 | PWB-iPSC DN | EEF1A2 | -1.300572732 | ENSG00000135451 | 10024 | PWB-MSC UP | TROAP | 1.0284774 |
| ENSG00000162601 | 114803 | PWB-iPSC UP | MYSM1 | 1.227768 | ENSG00000088992 | 54997 | PWB-iPSC DN | TESC | -1.300750069 | ENSG00000177432 | 266812 | PWB-MSC UP | NAP1L5 | 1.0212356 |
| ENSG00000241484 | 23779 | PWB-iPSC UP | ARHGAP8 | 1.222587 | ENSG00000149212 | 143686 | PWB-iPSC DN | SESN3 | -1.304228668 | ENSG00000197728 | 6231 | PWB-MSC UP | RPS26 | 1.0206817 |
| ENSG00000230937 | 642587 | PWB-iPSC UP | MIR205HG | 1.221856 | ENSG00000154319 | 83648 | PWB-iPSC DN | FAM167A | -1.304835012 | ENSG00000089091 | 55184 | PWB-MSC UP | DZANK1 | 1.0188025 |
| ENSG00000248360 | 201853 | PWB-iPSC UP | LINC00504 | 1.219903 | ENSG00000063180 | 770 | PWB-iPSC DN | CA11 | -1.31051902 | ENSG00000167703 | 124935 | PWB-MSC UP | SLC43A2 | 1.0182788 |
| ENSG00000274993 | 105375431 | PWB-iPSC UP | LOC105375431 | 1.218272 | ENSG00000205336 | 9289 | PWB-iPSC DN | ADGRG1 | -1.318270395 | ENSG00000243678 | 4831 | PWB-MSC UP | NME2 | 1.0157897 |
| ENSG00000198727 | 4519 | PWB-iPSC UP | CYTB | 1.216995 | ENSG00000137672 | 7225 | PWB-iPSC DN | TRPC6 | -1.320250321 | ENSG00000269834 | 102724105 | PWB-MSC UP | ZNF528-AS1 | 1.0131566 |
| ENSG00000178222 | 285498 | PWB-iPSC UP | RNF212 | 1.216873 | ENSG00000276077 | 102724951 | PWB-iPSC DN | LOC10272495 | -1.32037741 | ENSG00000166166 | 115708 | PWB-MSC UP | TRMT61A | 1.012964 |
| ENSG00000070669 | 440 | PWB-iPSC UP | ASNS | 1.216172 | ENSG00000159674 | 10417 | PWB-iPSC DN | SPON2 | -1.322269568 | ENSG00000029153 | 56938 | PWB-MSC UP | ARNTL2 | 1.0087216 |
| ENSG00000188933 | 162632 | PWB-iPSC UP | USP32P1 | 1.216006 | ENSG00000115468 | 80303 | PWB-iPSC DN | EFHD1 | -1.323260437 | ENSG00000206625 | 26827 | PWB-MSC UP | RNU6-1 | 1.0030765 |
| ENSG00000072163 | 55679 | PWB-iPSC UP | LIMS2 | 1.215678 | ENSG00000277067 | 102724843 | PWB-iPSC DN | LOC10272484 | -1.324562494 | ENSG00000064195 | 1747 | PWB-MSC UP | DLX3 | 1.0014802 |
| ENSG00000250508 | 105369364 | PWB-iPSC UP | LINC02701 | 1.214868 | ENSG00000100097 | 3956 | PWB-iPSC DN | LGALS1 | -1.325208654 | ENSG00000134323 | 4613 | PWB-MSC UP | MYCN | 1.001007 |
| ENSG00000198723 | 374877 | PWB-iPSC UP | TEX45 | 1.214575 | ENSG00000100031 | 2678 | PWB-iPSC DN | GGT1 | -1.325269763 | ENSG00000163694 | 54502 | PWB-MSC UP | RBM47 | 0.9945202 |
| ENSG00000256862 | 105369609 | PWB-iPSC UP | LOC105369609 | 1.213575 | ENSG00000134532 | 6660 | PWB-iPSC DN | SOX5 | -1.326687359 | ENSG00000139428 | 326625 | PWB-MSC UP | MMAB | 0.9893385 |
| ENSG00000155897 | 114 | PWB-iPSC UP | ADCY8 | 1.2131 | ENSG00000127578 | 117166 | PWB-iPSC DN | WFIKKN1 | -1.327482561 | ENSG00000146648 | 1956 | PWB-MSC UP | EGFR | 0.9888014 |
| ENSG00000205838 | 153657 | PWB-iPSC UP | TTC23L | 1.212903 | ENSG00000176428 | 155382 | PWB-iPSC DN | VPS37D | -1.329550269 | ENSG00000189410 | 400745 | PWB-MSC UP | SH2D5 | 0.9880645 |
| ENSG00000197385 | 344787 | PWB-iPSC UP | ZNF860 | 1.212862 | ENSG00000120875 | 1846 | PWB-iPSC DN | DUSP4 | -1.330796846 | ENSG00000196975 | 307 | PWB-MSC UP | ANXA4 | 0.9871981 |
| ENSG00000124313 | 23096 | PWB-iPSC UP | IQSEC2 | 1.212677 | ENSG00000172602 | 27289 | PWB-iPSC DN | RND1 | -1.335168523 | ENSG00000174206 | 144577 | PWB-MSC UP | C12orf66 | 0.9818761 |
| ENSG00000234494 | 100506325 | PWB-iPSC UP | SP2-AS1 | 1.211218 | ENSG00000021300 | 58473 | PWB-iPSC DN | PLEKHB1 | -1.336199039 | ENSG00000213214 | 445328 | PWB-MSC UP | ARHGEF35 | 0.9796014 |
| ENSG00000156006 | 10 | PWB-iPSC UP | NAT2 | 1.210973 | ENSG00000143507 | 11221 | PWB-iPSC DN | DUSP10 | -1.340555922 | ENSG00000196653 | 91392 | PWB-MSC UP | ZNF502 | 0.9782161 |
| ENSG00000203930 | 286411 | PWB-iPSC UP | LINC00632 | 1.210021 | ENSG00000240771 | 115557 | PWB-iPSC DN | ARHGEF25 | -1.341529398 | ENSG00000110975 | 341359 | PWB-MSC UP | SYT10 | 0.9758554 |
| ENSG00000176358 | 255061 | PWB-iPSC UP | TAC4 | 1.209945 | ENSG00000172247 | 114900 | PWB-iPSC DN | C1QTNF4 | -1.342676835 | ENSG00000224914 | 439994 | PWB-MSC UP | LINC00863 | 0.9682227 |
| ENSG00000180525 | 414235 | PWB-iPSC UP | PRR26 | 1.208743 | ENSG00000151376 | 10873 | PWB-iPSC DN | ME3 | -1.34323332 | ENSG00000133943 | 80017 | PWB-MSC UP | DGLUCY | 0.9653446 |
| ENSG00000168038 | 54986 | PWB-iPSC UP | ULK4 | 1.207944 | ENSG00000225968 | 392617 | PWB-iPSC DN | ELFN1 | -1.344304249 | ENSG00000166562 | 90701 | PWB-MSC UP | SEC11C | 0.9576447 |
| ENSG00000106133 | 260294 | PWB-iPSC UP | NSUN5P2 | 1.207627 | ENSG00000180113 | 221400 | PWB-iPSC DN | TDRD6 | -1.346779948 | ENSG00000274559 | 102724334 | PWB-MSC UP | LOC102724334 | 0.9569393 |
| ENSG00000263069 | 100294362 | PWB-iPSC UP | RNF213-AS1 | 1.207215 | ENSG00000175084 | 1674 | PWB-iPSC DN | DES | -1.346954999 | ENSG00000184923 | 728118 | PWB-MSC UP | NUTM2A | 0.9503495 |
| ENSG00000160229 | 7617 | PWB-iPSC UP | ZNF66 | 1.205981 | ENSG00000164434 | 2173 | PWB-iPSC DN | FABP7 | -1.349273834 | ENSG00000202538 | 26834 | PWB-MSC UP | RNU4-2 | 0.9495647 |
| ENSG00000265142 | 102723167 | PWB-iPSC UP | MIR133A1HG | 1.204602 | ENSG00000179431 | 24147 | PWB-iPSC DN | FJX1 | -1.355758073 | ENSG00000140479 | 5046 | PWB-MSC UP | PCSK6 | 0.9483549 |
| ENSG00000251364 | 100506258 | PWB-iPSC UP | LOC100506258 | 1.201618 | ENSG00000125968 | 3397 | PWB-iPSC DN | ID1 | -1.359042548 | ENSG00000157895 | 64897 | PWB-MSC UP | C12orf43 | 0.9474779 |
| ENSG00000135063 | 9413 | PWB-iPSC UP | FAM189A2 | 1.200625 | ENSG00000261308 | 401720 | PWB-iPSC DN | FIGNL2 | -1.359552246 | ENSG00000157570 | 90139 | PWB-MSC UP | TSPAN18 | 0.9449033 |
| ENSG00000280055 | 641384 | PWB-iPSC UP | TMEM75 | 1.198214 | ENSG00000111331 | 4940 | PWB-iPSC DN | OAS3 | -1.359612304 | ENSG00000111203 | 55846 | PWB-MSC UP | ITFG2 | 0.9425283 |
| ENSG00000102287 | 2564 | PWB-iPSC UP | GABRE | 1.194609 | ENSG00000049769 | 89801 | PWB-iPSC DN | PPP1R3F | -1.359656439 | ENSG00000115107 | 55240 | PWB-MSC UP | STEAP3 | 0.9424352 |
| ENSG00000276170 | 101929494 | PWB-iPSC UP | LOC101929494 | 1.193864 | ENSG00000131620 | 55107 | PWB-iPSC DN | ANO1 | -1.361509757 | ENSG00000168056 | 4054 | PWB-MSC UP | LTBP3 | 0.9417984 |
| ENSG00000281128 | 101243555 | PWB-iPSC UP | PTENP1-AS | 1.192836 | ENSG00000275496 | 102724701 | PWB-iPSC DN | LOC10272470 | -1.366745913 | ENSG00000143819 | 2052 | PWB-MSC UP | EPHX1 | 0.9388261 |
| ENSG00000185689 | 404220 | PWB-iPSC UP | C6orf201 | 1.19179 | ENSG00000069188 | 54549 | PWB-iPSC DN | SDK2 | -1.368188305 | ENSG00000066629 | 2009 | PWB-MSC UP | EML1 | 0.9373756 |
| ENSG00000101188 | 4923 | PWB-iPSC UP | NTSR1 | 1.191192 | ENSG00000136826 | 9314 | PWB-iPSC DN | KLF4 | -1.377634939 | ENSG00000123415 | 23583 | PWB-MSC UP | SMUG1 | 0.9351605 |
| ENSG00000232712 | 101929591 | PWB-iPSC UP | KIZ-AS1 | 1.188708 | ENSG00000002586 | 4267 | PWB-iPSC DN | CD99 | -1.383785599 | ENSG00000100599 | 79890 | PWB-MSC UP | RIN3 | 0.9334114 |
| ENSG00000272034 | 26822 | PWB-iPSC UP | SNORD14A | 1.187769 | ENSG00000057252 | 6646 | PWB-iPSC DN | SOAT1 | -1.385907255 | ENSG00000093072 | 51816 | PWB-MSC UP | ADA2 | 0.9297258 |
| ENSG00000187627 | 400966 | PWB-iPSC UP | RGPD1 | 1.187491 | ENSG00000036448 | 9172 | PWB-iPSC DN | MYOM2 | -1.386915383 | ENSG00000135476 | 9700 | PWB-MSC UP | ESPL1 | 0.9264609 |
| ENSG00000118194 | 7139 | PWB-iPSC UP | TNNT2 | 1.185972 | ENSG00000196503 | 132946 | PWB-iPSC DN | ARL9 | -1.387552204 | ENSG00000081803 | 93664 | PWB-MSC UP | CADPS2 | 0.9249437 |
| ENSG00000270580 | 105369154 | PWB-iPSC UP | PKD1P6-NPIPP1 | 1.18526 | ENSG00000176887 | 6664 | PWB-iPSC DN | SOX11 | -1.393631448 | ENSG00000226210 | 100288778 | PWB-MSC UP | WASH8P | 0.9186621 |
| ENSG00000250608 | 339874 | PWB-iPSC UP | LOC339874 | 1.185176 | ENSG00000065320 | 9423 | PWB-iPSC DN | NTN1 | -1.395036078 | ENSG00000130787 | 9026 | PWB-MSC UP | HIP1R | 0.907595 |
| ENSG00000122254 | 9956 | PWB-iPSC UP | HS3ST2 | 1.179848 | ENSG00000162009 | 6755 | PWB-iPSC DN | SSTR5 | -1.399896428 | ENSG00000139044 | 283358 | PWB-MSC UP | B4GALNT3 | 0.9055961 |
| ENSG00000163378 | 285203 | PWB-iPSC UP | EOGT | 1.177612 | ENSG00000132386 | 5176 | PWB-iPSC DN | SERPINF1 | -1.4007058 | ENSG00000139174 | 144165 | PWB-MSC UP | PRICKLE1 | 0.9051431 |
| ENSG00000169885 | 163688 | PWB-iPSC UP | CALML6 | 1.17713 | ENSG00000160801 | 5745 | PWB-iPSC DN | PTH1R | -1.402384012 | ENSG00000124160 | 57727 | PWB-MSC UP | NCOA5 | 0.9038608 |
| ENSG00000163596 | 130026 | PWB-iPSC UP | ICA1L | 1.176821 | ENSG00000130720 | 84929 | PWB-iPSC DN | FIBCD1 | -1.407540736 | ENSG00000162650 | 127002 | PWB-MSC UP | ATXN7L2 | 0.9030372 |
| ENSG00000130038 | 84766 | PWB-iPSC UP | CRACR2A | 1.17632 | ENSG00000064042 | 22998 | PWB-iPSC DN | LIMCH1 | -1.408213171 | ENSG00000013573 | 1663 | PWB-MSC UP | DDX11 | 0.9005788 |
| ENSG00000156239 | 29104 | PWB-iPSC UP | N6AMT1 | 1.173661 | ENSG00000129991 | 7137 | PWB-iPSC DN | TNNI3 | -1.408668578 | ENSG00000144063 | 7851 | PWB-MSC UP | MALL | 0.8989256 |
| ENSG00000146592 | 9586 | PWB-iPSC UP | CREB5 | 1.171515 | ENSG00000165757 | 57608 | PWB-iPSC DN | JCAD | -1.410348598 | ENSG00000204525 | 3107 | PWB-MSC UP | HLA-C | 0.8985059 |

| | | | | | | | | | | | | | | |
|---|---|---|---|---|---|---|---|---|---|---|---|---|---|---|
| ENSG00000183977 | 151649 | PWB-iPSC UP | PP2D1 | 1.170013 | ENSG00000159335 | 5763 | PWB-iPSC DN | PTMS | -1.411503411 | ENSG00000197124 | 91120 | PWB-MSC UP | ZNF682 | 0.8971912 |
| ENSG00000140543 | 55070 | PWB-iPSC UP | DET1 | 1.169864 | ENSG00000204347 | 388419 | PWB-iPSC DN | BTBD17 | -1.411928776 | ENSG00000106003 | 3955 | PWB-MSC UP | LFNG | 0.8958158 |
| ENSG00000226833 | 100505774 | PWB-iPSC UP | LOC100505774 | 1.169537 | ENSG00000130600 | 283120 | PWB-iPSC DN | H19 | -1.415453828 | ENSG00000139546 | 6895 | PWB-MSC UP | TARBP2 | 0.8952054 |
| ENSG00000226648 | 101927117 | PWB-iPSC UP | PLCG1-AS1 | 1.165989 | ENSG00000185942 | 286183 | PWB-iPSC DN | NKAIN3 | -1.416355962 | ENSG00000215866 | 100996702 | PWB-MSC UP | LINC01356 | 0.893141 |
| ENSG00000146215 | 401262 | PWB-iPSC UP | CRIP3 | 1.164985 | ENSG00000115738 | 3398 | PWB-iPSC DN | ID2 | -1.420243318 | ENSG00000073060 | 949 | PWB-MSC UP | SCARB1 | 0.8867959 |
| ENSG00000166321 | 25961 | PWB-iPSC UP | NUDT13 | 1.1646 | ENSG00000099617 | 1943 | PWB-iPSC DN | EFNA2 | -1.424545745 | ENSG00000139629 | 11226 | PWB-MSC UP | GALNT6 | 0.8847543 |
| ENSG00000269821 | 10984 | PWB-iPSC UP | KCNQ1OT1 | 1.162713 | ENSG00000253831 | 440695 | PWB-iPSC DN | ETV3L | -1.424608428 | ENSG00000120800 | 27340 | PWB-MSC UP | UTP20 | 0.8847127 |
| ENSG00000198763 | 4536 | PWB-iPSC UP | ND2 | 1.160883 | ENSG00000171617 | 8507 | PWB-iPSC DN | ENC1 | -1.427730532 | ENSG00000198040 | 7637 | PWB-MSC UP | ZNF84 | 0.8837326 |
| ENSG00000039139 | 1767 | PWB-iPSC UP | DNAH5 | 1.160827 | ENSG00000186007 | 93273 | PWB-iPSC DN | LEMD1 | -1.430461601 | ENSG00000221923 | 400713 | PWB-MSC UP | ZNF880 | 0.8721759 |
| ENSG00000115648 | 79083 | PWB-iPSC UP | MLPH | 1.158615 | ENSG00000161896 | 117283 | PWB-iPSC DN | IP6K3 | -1.432686176 | ENSG00000285756 | 389906 | PWB-MSC UP | LOC389906 | 0.8708378 |
| ENSG00000143127 | 8515 | PWB-iPSC UP | ITGA10 | 1.156783 | ENSG00000183421 | 54101 | PWB-iPSC DN | RIPK4 | -1.433197665 | ENSG00000197279 | 7718 | PWB-MSC UP | ZNF165 | 0.8704396 |
| ENSG00000123572 | 203447 | PWB-iPSC UP | NRK | 1.155355 | ENSG00000162552 | 54361 | PWB-iPSC DN | WNT4 | -1.440582016 | ENSG00000135111 | 6926 | PWB-MSC UP | TBX3 | 0.8680336 |
| ENSG00000166343 | 118490 | PWB-iPSC UP | MSS51 | 1.155245 | ENSG00000069696 | 1815 | PWB-iPSC DN | DRD4 | -1.443945956 | ENSG00000167272 | 51367 | PWB-MSC UP | POP5 | 0.8675054 |
| ENSG00000115361 | 33 | PWB-iPSC UP | ACADL | 1.149377 | ENSG00000068078 | 2261 | PWB-iPSC DN | FGFR3 | -1.448502201 | ENSG00000105699 | 51599 | PWB-MSC UP | LSR | 0.8669849 |
| ENSG00000170236 | 373509 | PWB-iPSC UP | USP50 | 1.143901 | ENSG00000159212 | 54102 | PWB-iPSC DN | CLIC6 | -1.451367432 | ENSG00000234289 | 54145 | PWB-MSC UP | H2BS1 | 0.8661254 |
| ENSG00000206656 | 100033429 | PWB-iPSC UP | SNORD116-17 | 1.143816 | ENSG00000135097 | 4440 | PWB-iPSC DN | MSI1 | -1.453525145 | ENSG00000113645 | 23286 | PWB-MSC UP | WWC1 | 0.8656269 |
| ENSG00000175463 | 374403 | PWB-iPSC UP | TBC1D10C | 1.142427 | ENSG00000105048 | 7138 | PWB-iPSC DN | TNNT1 | -1.455508001 | ENSG00000168038 | 54986 | PWB-MSC UP | ULK4 | 0.8552035 |
| ENSG00000151882 | 56477 | PWB-iPSC UP | CCL28 | 1.14196 | ENSG00000115267 | 64135 | PWB-iPSC DN | IFIH1 | -1.458007931 | ENSG00000253958 | 137075 | PWB-MSC UP | CLDN23 | 0.8531682 |
| ENSG00000074660 | 8578 | PWB-iPSC UP | SCARF1 | 1.140423 | ENSG00000185201 | 10581 | PWB-iPSC DN | IFITM2 | -1.464270412 | ENSG00000253731 | 56109 | PWB-MSC UP | PCDHGA6 | 0.8527639 |
| ENSG00000112309 | 135152 | PWB-iPSC UP | B3GAT2 | 1.140047 | ENSG00000181649 | 7262 | PWB-iPSC DN | PHLDA2 | -1.464290211 | ENSG00000247077 | 192111 | PWB-MSC UP | PGAM5 | 0.8413098 |
| ENSG00000238133 | 339751 | PWB-iPSC UP | MAP3K20-AS1 | 1.139602 | ENSG00000164929 | 79870 | PWB-iPSC DN | BAALC | -1.467974885 | ENSG00000235173 | 51236 | PWB-MSC UP | HGH1 | 0.8400589 |
| ENSG00000245148 | 100506020 | PWB-iPSC UP | ARAP1-AS2 | 1.138608 | ENSG00000082556 | 4986 | PWB-iPSC DN | OPRK1 | -1.471422012 | ENSG00000162433 | 205 | PWB-MSC UP | AK4 | 0.8388119 |
| ENSG00000196793 | 8187 | PWB-iPSC UP | ZNF239 | 1.137234 | ENSG00000170390 | 166614 | PWB-iPSC DN | DCLK2 | -1.474099478 | ENSG00000122965 | 9904 | PWB-MSC UP | RBM19 | 0.8382132 |
| ENSG00000142698 | 84970 | PWB-iPSC UP | C1orf94 | 1.136145 | ENSG00000167779 | 3489 | PWB-iPSC DN | IGFBP6 | -1.484526777 | ENSG00000134709 | 51361 | PWB-MSC UP | HOOK1 | 0.8380582 |
| ENSG00000154734 | 9510 | PWB-iPSC UP | ADAMTS1 | 1.134981 | ENSG00000136842 | 7111 | PWB-iPSC DN | TMOD1 | -1.489133543 | ENSG00000139343 | 6636 | PWB-MSC UP | SNRPF | 0.8375264 |
| ENSG00000216937 | 79741 | PWB-iPSC UP | CCDC7 | 1.132375 | ENSG00000253626 | 143244 | PWB-iPSC DN | EIF5AL1 | -1.494113324 | ENSG00000184047 | 56616 | PWB-MSC UP | DIABLO | 0.8374892 |
| ENSG00000196549 | 4311 | PWB-iPSC UP | MME | 1.131697 | ENSG00000177359 | 144203 | PWB-iPSC DN | OVOS2 | -1.494906267 | ENSG00000188199 | 729262 | PWB-MSC UP | NUTM2B | 0.8364368 |
| ENSG00000241155 | 100874246 | PWB-iPSC UP | ARHGAP31-AS1 | 1.129988 | ENSG00000197769 | 440738 | PWB-iPSC DN | MAP1LC3C | -1.505528463 | ENSG00000205352 | 54458 | PWB-MSC UP | PRR13 | 0.835468 |
| ENSG00000180264 | 347088 | PWB-iPSC UP | ADGRD2 | 1.125092 | ENSG00000147257 | 2719 | PWB-iPSC DN | GPC3 | -1.509935419 | ENSG00000177084 | 5426 | PWB-MSC UP | POLE | 0.8349188 |
| ENSG00000236438 | 728262 | PWB-iPSC UP | FAM157A | 1.124612 | ENSG00000165215 | 1365 | PWB-iPSC DN | CLDN3 | -1.510583111 | ENSG00000020129 | 23154 | PWB-MSC UP | NCDN | 0.8344962 |
| ENSG00000152402 | 2977 | PWB-iPSC UP | GUCY1A2 | 1.123581 | ENSG00000109339 | 5602 | PWB-iPSC DN | MAPK10 | -1.510607462 | ENSG00000149292 | 54970 | PWB-MSC UP | TTC12 | 0.832784 |
| ENSG00000251595 | 79963 | PWB-iPSC UP | ABCA11P | 1.120229 | ENSG00000090776 | 1947 | PWB-iPSC DN | EFNB1 | -1.523260935 | ENSG00000179041 | 23212 | PWB-MSC UP | RRS1 | 0.8324658 |
| ENSG00000167555 | 84436 | PWB-iPSC UP | ZNF528 | 1.11957 | ENSG00000271601 | 128077 | PWB-iPSC DN | LIX1L | -1.525170118 | ENSG00000123064 | 79039 | PWB-MSC UP | DDX54 | 0.8318223 |
| ENSG00000244560 | 155060 | PWB-iPSC UP | LOC155060 | 1.118865 | ENSG00000167535 | 784 | PWB-iPSC DN | CACNB3 | -1.532567451 | ENSG00000125848 | 23767 | PWB-MSC UP | FLRT3 | 0.831256 |
| ENSG00000152580 | 285313 | PWB-iPSC UP | IGSF10 | 1.11606 | ENSG00000100298 | 164668 | PWB-iPSC DN | APOBEC3H | -1.532956382 | ENSG00000004478 | 2288 | PWB-MSC UP | FKBP4 | 0.8310726 |
| ENSG00000227733 | 101927468 | PWB-iPSC UP | LOC101927468 | 1.1151 | ENSG00000124092 | 140690 | PWB-iPSC DN | CTCFL | -1.533814666 | ENSG00000200087 | 26768 | PWB-MSC UP | SNORA73B | 0.8302499 |
| ENSG00000184384 | 84441 | PWB-iPSC UP | MAML2 | 1.113999 | ENSG00000280780 | 153469 | PWB-iPSC DN | JAKMIP2-AS1 | -1.539343779 | ENSG00000135409 | 269 | PWB-MSC UP | AMHR2 | 0.824918 |
| ENSG00000154065 | 147463 | PWB-iPSC UP | ANKRD29 | 1.112771 | ENSG00000165507 | 11067 | PWB-iPSC DN | DEPP1 | -1.561011675 | ENSG00000132773 | 114034 | PWB-MSC UP | TOE1 | 0.8236723 |
| ENSG00000205885 | 283314 | PWB-iPSC UP | C1RL-AS1 | 1.110961 | ENSG00000236106 | 112268411 | PWB-iPSC DN | LOC11226841 | -1.561825205 | ENSG00000256546 | 100506691 | PWB-MSC UP | LOC100506691 | 0.8167921 |
| ENSG00000077522 | 88 | PWB-iPSC UP | ACTN2 | 1.110725 | ENSG00000167074 | 7008 | PWB-iPSC DN | TEF | -1.564314731 | ENSG00000121316 | 79887 | PWB-MSC UP | PLBD1 | 0.8137805 |
| ENSG00000156049 | 9630 | PWB-iPSC UP | GNA14 | 1.109719 | ENSG00000164442 | 10370 | PWB-iPSC DN | CITED2 | -1.567688019 | ENSG00000123395 | 60673 | PWB-MSC UP | ATG101 | 0.8105782 |
| ENSG00000132196 | 51478 | PWB-iPSC UP | HSD17B7 | 1.109297 | ENSG00000157601 | 4599 | PWB-iPSC DN | MX1 | -1.568798818 | ENSG00000204186 | 57683 | PWB-MSC UP | ZDBF2 | 0.8079522 |
| ENSG00000253972 | 105375726 | PWB-iPSC UP | MAL2-AS1 | 1.109209 | ENSG00000167614 | 57348 | PWB-iPSC DN | TTYH1 | -1.570189995 | ENSG00000123349 | 5204 | PWB-MSC UP | PFDN5 | 0.8068896 |
| ENSG00000196247 | 51427 | PWB-iPSC UP | ZNF107 | 1.107905 | ENSG00000128266 | 2781 | PWB-iPSC DN | GNAZ | -1.572167836 | ENSG00000113205 | 56132 | PWB-MSC UP | PCDHB3 | 0.8061573 |
| ENSG00000165084 | 116328 | PWB-iPSC UP | C8orf34 | 1.107203 | ENSG00000188042 | 10123 | PWB-iPSC DN | ARL4C | -1.572958654 | ENSG00000155016 | 113612 | PWB-MSC UP | CYP2U1 | 0.8054372 |
| ENSG00000006534 | 221 | PWB-iPSC UP | ALDH3B1 | 1.106702 | ENSG00000125845 | 650 | PWB-iPSC DN | BMP2 | -1.582827815 | ENSG00000100029 | 23481 | PWB-MSC UP | PES1 | 0.8030419 |
| ENSG00000223705 | 155400 | PWB-iPSC UP | NSUN5P1 | 1.106651 | ENSG00000111186 | 81029 | PWB-iPSC DN | WNT5B | -1.584060303 | ENSG00000180855 | 10224 | PWB-MSC UP | ZNF443 | 0.8024736 |
| ENSG00000105146 | 6795 | PWB-iPSC UP | AURKC | 1.106603 | ENSG00000184985 | 57537 | PWB-iPSC DN | SORCS2 | -1.592211196 | ENSG00000139116 | 55605 | PWB-MSC UP | KIF21A | 0.8013764 |
| ENSG00000137266 | 63027 | PWB-iPSC UP | SLC22A23 | 1.103315 | ENSG00000180694 | 169200 | PWB-iPSC DN | TMEM64 | -1.594157784 | ENSG00000139180 | 4704 | PWB-MSC UP | NDUFA9 | 0.7972851 |
| ENSG00000170006 | 201799 | PWB-iPSC UP | TMEM154 | 1.101798 | ENSG00000138646 | 51191 | PWB-iPSC DN | HERC5 | -1.596124215 | ENSG00000167173 | 56905 | PWB-MSC UP | C15orf39 | 0.7962833 |
| ENSG00000234945 | 100505624 | PWB-iPSC UP | GTF3C2-AS1 | 1.100678 | ENSG00000148204 | 286204 | PWB-iPSC DN | CRB2 | -1.606766144 | ENSG00000128973 | 54982 | PWB-MSC UP | CLN6 | 0.7960653 |
| ENSG00000198785 | 116443 | PWB-iPSC UP | GRIN3A | 1.100452 | ENSG00000189060 | 3005 | PWB-iPSC DN | H1-0 | -1.611092719 | ENSG00000198393 | 7574 | PWB-MSC UP | ZNF26 | 0.7880674 |
| ENSG00000215424 | 114044 | PWB-iPSC UP | MCM3AP-AS1 | 1.100403 | ENSG00000066230 | 6550 | PWB-iPSC DN | SLC9A3 | -1.61730095 | ENSG00000167779 | 3489 | PWB-MSC UP | IGFBP6 | 0.7875913 |
| ENSG00000075290 | 7479 | PWB-iPSC UP | WNT8B | 1.099688 | ENSG00000286940 | 102725068 | PWB-iPSC DN | MICB-DT | -1.623563719 | ENSG00000231752 | 647121 | PWB-MSC UP | EMBP1 | 0.7854018 |
| ENSG00000198899 | 4508 | PWB-iPSC UP | ATP6 | 1.09749 | ENSG00000107796 | 59 | PWB-iPSC DN | ACTA2 | -1.634458972 | ENSG00000176834 | 54621 | PWB-MSC UP | VSIG10 | 0.7839191 |
| ENSG00000227533 | 440584 | PWB-iPSC UP | SLC2A1-AS1 | 1.097366 | ENSG00000177283 | 8325 | PWB-iPSC DN | FZD8 | -1.635587435 | ENSG00000011105 | 10867 | PWB-MSC UP | TSPAN9 | 0.7830196 |
| ENSG00000223756 | 650368 | PWB-iPSC UP | TSSC2 | 1.096121 | ENSG00000253716 | 100507316 | PWB-iPSC DN | MINCR | -1.637210035 | ENSG00000199753 | 692227 | PWB-MSC UP | SNORD104 | 0.7824748 |
| ENSG00000163806 | 245711 | PWB-iPSC UP | SPDYA | 1.095307 | ENSG00000134363 | 10468 | PWB-iPSC DN | FST | -1.647746183 | ENSG00000132382 | 10514 | PWB-MSC UP | MYBBP1A | 0.7796964 |
| ENSG00000243701 | 344595 | PWB-iPSC UP | DUBR | 1.091378 | ENSG00000000005 | 64102 | PWB-iPSC DN | TNMD | -1.648688409 | ENSG00000134809 | 26519 | PWB-MSC UP | TIMM10 | 0.7794953 |
| ENSG00000225855 | 284618 | PWB-iPSC UP | RUSC1-AS1 | 1.090287 | ENSG00000158473 | 912 | PWB-iPSC DN | CD1D | -1.654056206 | ENSG00000139437 | 84260 | PWB-MSC UP | TCHP | 0.7790422 |
| ENSG00000250337 | 643401 | PWB-iPSC UP | PURPL | 1.089867 | ENSG00000259129 | 100506433 | PWB-iPSC DN | LINC00648 | -1.654124113 | ENSG00000089248 | 10961 | PWB-MSC UP | ERP29 | 0.7745684 |
| ENSG00000203739 | 101928673 | PWB-iPSC UP | LOC101928673 | 1.084251 | ENSG00000116729 | 79971 | PWB-iPSC DN | WLS | -1.659838486 | ENSG00000109084 | 27346 | PWB-MSC UP | TMEM97 | 0.7743048 |
| ENSG00000249592 | 100129917 | PWB-iPSC UP | LOC100129917 | 1.083903 | ENSG00000171621 | 80176 | PWB-iPSC DN | SPSB1 | -1.659954831 | ENSG00000053372 | 51154 | PWB-MSC UP | MRTO4 | 0.7739556 |
| ENSG00000248538 | 157273 | PWB-iPSC UP | LOC157273 | 1.080095 | ENSG00000230006 | 645784 | PWB-iPSC DN | ANKRD36BP2 | -1.666914286 | ENSG00000111665 | 83461 | PWB-MSC UP | CDCA3 | 0.7728765 |
| ENSG00000223960 | 101927027 | PWB-iPSC UP | CHROMR | 1.080062 | ENSG00000169499 | 59339 | PWB-iPSC DN | PLEKHA2 | -1.667511911 | ENSG00000127328 | 117177 | PWB-MSC UP | RAB3IP | 0.7712133 |
| ENSG00000106588 | 5683 | PWB-iPSC UP | PSMA2 | 1.07915 | ENSG00000142530 | 112703 | PWB-iPSC DN | FAM71E1 | -1.673603048 | ENSG00000135446 | 1019 | PWB-MSC UP | CDK4 | 0.7659238 |
| ENSG00000254413 | 386593 | PWB-iPSC UP | CHKB-CPT1B | 1.078751 | ENSG00000172572 | 5139 | PWB-iPSC DN | PDE3A | -1.674186597 | ENSG00000174718 | 55196 | PWB-MSC UP | RESF1 | 0.7657541 |
| ENSG00000173210 | 22885 | PWB-iPSC UP | ABLIM3 | 1.078715 | ENSG00000124588 | 4835 | PWB-iPSC DN | NQO2 | -1.674745998 | ENSG00000111737 | 11021 | PWB-MSC UP | RAB35 | 0.7649475 |
| ENSG00000111799 | 1303 | PWB-iPSC UP | COL12A1 | 1.077527 | ENSG00000155052 | 129684 | PWB-iPSC DN | CNTNAP5 | -1.697634224 | ENSG00000065029 | 7629 | PWB-MSC UP | ZNF76 | 0.7583448 |
| ENSG00000169436 | 169044 | PWB-iPSC UP | COL22A1 | 1.073901 | ENSG00000130287 | 1463 | PWB-iPSC DN | NCAN | -1.701933299 | ENSG00000135424 | 3679 | PWB-MSC UP | ITGA7 | 0.757418 |
| ENSG00000163531 | 23114 | PWB-iPSC UP | NFASC | 1.071837 | ENSG00000116574 | 58480 | PWB-iPSC DN | RHOU | -1.711679732 | ENSG00000213988 | 7643 | PWB-MSC UP | ZNF90 | 0.7563479 |
| ENSG00000166762 | 117155 | PWB-iPSC UP | CATSPER2 | 1.071288 | ENSG00000163565 | 3428 | PWB-iPSC DN | IFI16 | -1.712288007 | ENSG00000168906 | 4144 | PWB-MSC UP | MAT2A | 0.7556303 |
| ENSG00000185306 | 115749 | PWB-iPSC UP | C12orf56 | 1.071261 | ENSG00000185652 | 4908 | PWB-iPSC DN | NTF3 | -1.714066392 | ENSG00000170473 | 84305 | PWB-MSC UP | PYM1 | 0.7553845 |
| ENSG00000196263 | 57573 | PWB-iPSC UP | ZNF471 | 1.071039 | ENSG00000184232 | 220323 | PWB-iPSC DN | OAF | -1.717023084 | ENSG00000225366 | 6998 | PWB-MSC UP | TDGF1P3 | 0.7553708 |
| ENSG00000079337 | 10411 | PWB-iPSC UP | RAPGEF3 | 1.07058 | ENSG00000213023 | 84258 | PWB-iPSC DN | SYT3 | -1.718004738 | ENSG00000176871 | 55884 | PWB-MSC UP | WSB2 | 0.7541422 |
| ENSG00000255408 | 56145 | PWB-iPSC UP | PCDHA3 | 1.070343 | ENSG00000187608 | 9636 | PWB-iPSC DN | ISG15 | -1.731323084 | ENSG00000129347 | 65095 | PWB-MSC UP | KRI1 | 0.7532502 |
| ENSG00000278811 | 100289211 | PWB-iPSC UP | LINC00624 | 1.069866 | ENSG00000188620 | 340784 | PWB-iPSC DN | HMX3 | -1.736598108 | ENSG00000167548 | 8085 | PWB-MSC UP | KMT2D | 0.7529804 |
| ENSG00000238197 | 100506215 | PWB-iPSC UP | PAXBP1-AS1 | 1.067714 | ENSG00000232938 | 388574 | PWB-iPSC DN | RPL23AP87 | -1.738075243 | ENSG00000111206 | 2305 | PWB-MSC UP | FOXM1 | 0.7526974 |
| ENSG00000151025 | 57512 | PWB-iPSC UP | GPR158 | 1.067303 | ENSG00000272636 | 8447 | PWB-iPSC DN | DOC2B | -1.740767847 | ENSG00000120875 | 1846 | PWB-MSC UP | DUSP4 | 0.7504057 |
| ENSG00000153814 | 221895 | PWB-iPSC UP | JAZF1 | 1.066795 | ENSG00000157322 | 348174 | PWB-iPSC DN | CLEC18A | -1.741131544 | ENSG00000076248 | 7374 | PWB-MSC UP | UNG | 0.7498056 |
| ENSG00000186472 | 27445 | PWB-iPSC UP | PCLO | 1.065256 | ENSG00000172201 | 3400 | PWB-iPSC DN | ID4 | -1.746412963 | ENSG00000207205 | 101954267 | PWB-MSC UP | RNVU1-15 | 0.7473746 |
| ENSG00000123104 | 3709 | PWB-iPSC UP | ITPR2 | 1.064733 | ENSG00000099284 | 55506 | PWB-iPSC DN | MACROH2A2 | -1.746769492 | ENSG00000174233 | 112 | PWB-MSC UP | ADCY6 | 0.7431534 |
| ENSG00000127311 | 92797 | PWB-iPSC UP | HELB | 1.062795 | ENSG00000186094 | 84871 | PWB-iPSC DN | AGBL4 | -1.747949493 | ENSG00000141759 | 10907 | PWB-MSC UP | TXNL4A | 0.7427462 |
| ENSG00000101292 | 128674 | PWB-iPSC UP | PROKR2 | 1.060803 | ENSG00000235034 | 342918 | PWB-iPSC DN | C19orf81 | -1.757171876 | ENSG00000257337 | 283335 | PWB-MSC UP | LOC283335 | 0.7381145 |
| ENSG00000177432 | 266812 | PWB-iPSC UP | NAP1L5 | 1.05969 | ENSG00000157851 | 56896 | PWB-iPSC DN | DPYSL5 | -1.763875926 | ENSG00000083307 | 79977 | PWB-MSC UP | GRHL2 | 0.7376973 |
| ENSG00000178852 | 124989 | PWB-iPSC UP | EFCAB13 | 1.059581 | ENSG00000163749 | 339965 | PWB-iPSC DN | CCDC158 | -1.770416132 | ENSG00000226784 | 441531 | PWB-MSC UP | PGAM4 | 0.7365163 |
| ENSG00000279491 | 399900 | PWB-iPSC UP | LOC399900 | 1.056886 | ENSG00000162909 | 824 | PWB-iPSC DN | CAPN2 | -1.772203456 | ENSG00000159445 | 117145 | PWB-MSC UP | THEM4 | 0.7364513 |
| ENSG00000132010 | 7568 | PWB-iPSC UP | ZNF20 | 1.056149 | ENSG00000177238 | 493829 | PWB-iPSC DN | TRIM72 | -1.775199377 | ENSG00000144036 | 23233 | PWB-MSC UP | EXOC6B | 0.7362851 |
| ENSG00000069869 | 4734 | PWB-iPSC UP | NEDD4 | 1.054292 | ENSG00000132821 | 128434 | PWB-iPSC DN | VSTM2L | -1.775731059 | ENSG00000112759 | 2030 | PWB-MSC UP | SLC29A1 | 0.7360838 |
| ENSG00000020577 | 23034 | PWB-iPSC UP | SAMD4A | 1.047279 | ENSG00000196154 | 6275 | PWB-iPSC DN | S100A4 | -1.810158524 | ENSG00000134287 | 377 | PWB-MSC UP | ARF3 | 0.7360391 |
| ENSG00000138061 | 1545 | PWB-iPSC UP | CYP1B1 | 1.046887 | ENSG00000133107 | 7223 | PWB-iPSC DN | TRPC4 | -1.825244083 | ENSG00000135457 | 7024 | PWB-MSC UP | TFCP2 | 0.7358949 |
| ENSG00000009694 | 10178 | PWB-iPSC UP | TENM1 | 1.044905 | ENSG00000118946 | 27253 | PWB-iPSC DN | PCDH17 | -1.828986025 | ENSG00000111786 | 8683 | PWB-MSC UP | SRSF9 | 0.7358708 |
| ENSG00000103257 | 8140 | PWB-iPSC UP | SLC7A5 | 1.044383 | ENSG00000078018 | 4133 | PWB-iPSC DN | MAP2 | -1.829460873 | ENSG00000111412 | 79794 | PWB-MSC UP | C12orf49 | 0.7338627 |
| ENSG00000231365 | 101929147 | PWB-iPSC UP | WARS2-AS1 | 1.04356 | ENSG00000105499 | 8605 | PWB-iPSC DN | PLA2G4C | -1.829650997 | ENSG00000187778 | 10445 | PWB-MSC UP | MCRS1 | 0.732021 |
| ENSG00000235138 | 100130548 | PWB-iPSC UP | LOC100130548 | 1.042227 | ENSG00000137331 | 8870 | PWB-iPSC DN | IER3 | -1.831397147 | ENSG00000173041 | 340252 | PWB-MSC UP | ZNF680 | 0.7311363 |
| ENSG00000275954 | 84218 | PWB-iPSC UP | TBC1D3F | 1.041397 | ENSG00000204711 | 138255 | PWB-iPSC DN | C9orf135 | -1.834029854 | ENSG00000077044 | 8527 | PWB-MSC UP | DGKD | 0.730582 |
| ENSG00000132846 | 84327 | PWB-iPSC UP | ZBED3 | 1.041394 | ENSG00000163666 | 8820 | PWB-iPSC DN | HESX1 | -1.835947127 | ENSG00000145882 | 78991 | PWB-MSC UP | PCYOX1L | 0.7297154 |
| ENSG00000280109 | 191585 | PWB-iPSC UP | PLAC4 | 1.037685 | ENSG00000176842 | 10265 | PWB-iPSC DN | IRX5 | -1.85388363 | ENSG00000172748 | 169270 | PWB-MSC UP | ZNF596 | 0.7287009 |
| ENSG00000162599 | 4774 | PWB-iPSC UP | NFIA | 1.033696 | ENSG00000240204 | 100287482 | PWB-iPSC DN | SMKR1 | -1.862013821 | ENSG00000181396 | 79701 | PWB-MSC UP | OGFOD3 | 0.7285678 |
| ENSG00000204632 | 3135 | PWB-iPSC UP | HLA-G | 1.027929 | ENSG00000143320 | 1382 | PWB-iPSC DN | CRABP2 | -1.867028653 | ENSG00000155974 | 23426 | PWB-MSC UP | GRIP1 | 0.7208644 |
| ENSG00000158483 | 55199 | PWB-iPSC UP | FAM86C1 | 1.024304 | ENSG00000129990 | 6861 | PWB-iPSC DN | SYT5 | -1.867422211 | ENSG00000123353 | 29095 | PWB-MSC UP | ORMDL2 | 0.7196191 |
| ENSG00000228526 | 106614088 | PWB-iPSC UP | MIR34AHG | 1.018812 | ENSG00000171873 | 146 | PWB-iPSC DN | ADRA1D | -1.873454445 | ENSG00000150977 | 196383 | PWB-MSC UP | RILPL2 | 0.7187356 |
| ENSG00000227403 | 100996579 | PWB-iPSC UP | LINC01806 | 1.018693 | ENSG00000134686 | 1912 | PWB-iPSC DN | PHC2 | -1.876491094 | ENSG00000139620 | 54934 | PWB-MSC UP | KANSL2 | 0.7178336 |
| ENSG00000135298 | 577 | PWB-iPSC UP | ADGRB3 | 1.017929 | ENSG00000088881 | 57593 | PWB-iPSC DN | EBF4 | -1.880229542 | ENSG00000182871 | 80781 | PWB-MSC UP | COL18A1 | 0.7157234 |
| ENSG00000232656 | 55853 | PWB-iPSC UP | IDI2-AS1 | 1.017471 | ENSG00000129221 | 23746 | PWB-iPSC DN | AIPL1 | -1.8804559 | ENSG00000278771 | 378707 | PWB-MSC UP | RN7SL3 | 0.7100855 |
| ENSG00000284770 | 6905 | PWB-iPSC UP | TBCE | 1.016803 | ENSG00000090530 | 55214 | PWB-iPSC DN | P3H2 | -1.891304398 | ENSG00000198093 | 65251 | PWB-MSC UP | ZNF649 | 0.7093423 |
| ENSG00000119943 | 84795 | PWB-iPSC UP | PYROXD2 | 1.01447 | ENSG00000162878 | 91461 | PWB-iPSC DN | PKDCC | -1.904064788 | ENSG00000084112 | 54434 | PWB-MSC UP | SSH1 | 0.7086988 |

| | | | | | | | | | | | | | | |
|---|---|---|---|---|---|---|---|---|---|---|---|---|---|---|
| ENSG00000161381 | 57125 | PWB-iPSC UP | PLXDC1 | 1.01408 | ENSG00000266074 | 57597 | PWB-iPSC DN | BAHCC1 | -1.905153582 | ENSG00000160256 | 85395 | PWB-MSC UP | FAM207A | 0.7076872 |
| ENSG00000241472 | 100506994 | PWB-iPSC UP | PTPRG-AS1 | 1.011285 | ENSG00000167874 | 92162 | PWB-iPSC DN | TMEM88 | -1.907587101 | ENSG00000065361 | 2065 | PWB-MSC UP | ERBB3 | 0.7073213 |
| ENSG00000118507 | 9465 | PWB-iPSC UP | AKAP7 | 1.010883 | ENSG00000140015 | 27133 | PWB-iPSC DN | KCNH5 | -1.918483118 | ENSG00000242802 | 9907 | PWB-MSC UP | AP5Z1 | 0.7054303 |
| ENSG00000188921 | 401494 | PWB-iPSC UP | HACD4 | 1.00888 | ENSG00000287260 | 101928868 | PWB-iPSC DN | LOC10192886 | -1.942741735 | ENSG00000185684 | 347918 | PWB-MSC UP | EP400P1 | 0.7032147 |
| ENSG00000223773 | 401577 | PWB-iPSC UP | CD99P1 | 1.00718 | ENSG00000128683 | 2571 | PWB-iPSC DN | GAD1 | -1.951433594 | ENSG00000090612 | 10795 | PWB-MSC UP | ZNF268 | 0.701602 |
| ENSG00000189292 | 285016 | PWB-iPSC UP | ALKAL2 | 1.004247 | ENSG00000125730 | 718 | PWB-iPSC DN | C3 | -1.956736431 | ENSG00000182199 | 6472 | PWB-MSC UP | SHMT2 | 0.6995919 |
| ENSG00000161010 | 51149 | PWB-iPSC UP | MRNIP | 1.003374 | ENSG00000008118 | 57172 | PWB-iPSC DN | CAMK1G | -1.960495548 | ENSG00000111361 | 1967 | PWB-MSC UP | EIF2B1 | 0.6995633 |
| ENSG00000176593 | 100128398 | PWB-iPSC UP | LOC100128398 | 1.003273 | ENSG00000167711 | 5345 | PWB-iPSC DN | SERPINF2 | -1.97430437 | ENSG00000182544 | 84975 | PWB-MSC UP | MFSD5 | 0.6981114 |
| ENSG00000067840 | 57595 | PWB-iPSC UP | PDZD4 | 1.001435 | ENSG00000125726 | 970 | PWB-iPSC DN | CD70 | -1.993422362 | ENSG00000275023 | 4302 | PWB-MSC UP | MLLT6 | 0.6950066 |
| ENSG00000140092 | 10516 | PWB-iPSC UP | FBLN5 | 1.001248 | ENSG00000215912 | 100287898 | PWB-iPSC DN | TTC34 | -1.996996724 | ENSG00000198324 | 144717 | PWB-MSC UP | PHETA1 | 0.6932888 |
| ENSG00000253438 | 100750225 | PWB-iPSC UP | PCAT1 | 0.997803 | ENSG00000110148 | 887 | PWB-iPSC DN | CCKBR | -1.997449958 | ENSG00000089280 | 2521 | PWB-MSC UP | FUS | 0.6910138 |
| ENSG00000243989 | 95 | PWB-iPSC UP | ACY1 | 0.996062 | ENSG00000165495 | 63876 | PWB-iPSC DN | PKNOX2 | -1.999949964 | ENSG00000130779 | 6249 | PWB-MSC UP | CLIP1 | 0.6889974 |
| ENSG00000200534 | 594839 | PWB-iPSC UP | SNORA33 | 0.994508 | ENSG00000163661 | 5806 | PWB-iPSC DN | PTX3 | -2.000689864 | ENSG00000139718 | 23067 | PWB-MSC UP | SETD1B | 0.6885484 |
| ENSG00000128000 | 163131 | PWB-iPSC UP | ZNF780B | 0.992303 | ENSG00000267909 | 56936 | PWB-iPSC DN | CCDC177 | -2.000837441 | ENSG00000139405 | 84934 | PWB-MSC UP | RITA1 | 0.6883676 |
| ENSG00000135976 | 375248 | PWB-iPSC UP | ANKRD36 | 0.990904 | ENSG00000005421 | 5444 | PWB-iPSC DN | PON1 | -2.004953844 | ENSG00000178921 | 5198 | PWB-MSC UP | PFAS | 0.6835979 |
| ENSG00000234690 | 101927043 | PWB-iPSC UP | EPCAM-DT | 0.990041 | ENSG00000198597 | 9745 | PWB-iPSC DN | ZNF536 | -2.008727378 | ENSG00000111666 | 56994 | PWB-MSC UP | CHPT1 | 0.683025 |
| ENSG00000145536 | 170690 | PWB-iPSC UP | ADAMTS16 | 0.989066 | ENSG00000180447 | 2619 | PWB-iPSC DN | GAS1 | -2.016163739 | ENSG00000165271 | 65083 | PWB-MSC UP | NOL6 | 0.6828399 |
| ENSG00000129682 | 2258 | PWB-iPSC UP | FGF13 | 0.988538 | ENSG00000174460 | 170261 | PWB-iPSC DN | ZCCHC12 | -2.028907774 | ENSG00000132361 | 23277 | PWB-MSC UP | CLUH | 0.6817719 |
| ENSG00000134042 | 83876 | PWB-iPSC UP | MRO | 0.988328 | ENSG00000184486 | 5454 | PWB-iPSC DN | POU3F2 | -2.035803996 | ENSG00000196458 | 100289635 | PWB-MSC UP | ZNF605 | 0.678517 |
| ENSG00000231689 | 104355152 | PWB-iPSC UP | LINC01090 | 0.988073 | ENSG00000111181 | 6539 | PWB-iPSC DN | SLC6A12 | -2.040018142 | ENSG00000089693 | 8079 | PWB-MSC UP | MLF2 | 0.6783405 |
| ENSG00000204352 | 445577 | PWB-iPSC UP | C9orf129 | 0.986558 | ENSG00000142319 | 6531 | PWB-iPSC DN | SLC6A3 | -2.061383746 | ENSG00000104731 | 54758 | PWB-MSC UP | KLHDC4 | 0.6767027 |
| ENSG00000170891 | 54360 | PWB-iPSC UP | CYTL1 | 0.986505 | ENSG00000110195 | 2348 | PWB-iPSC DN | FOLR1 | -2.070284193 | ENSG00000146834 | 56257 | PWB-MSC UP | MEPCE | 0.6763964 |
| ENSG00000158220 | 83850 | PWB-iPSC UP | ESYT3 | 0.983545 | ENSG00000227640 | 100507533 | PWB-iPSC DN | SOX21-AS1 | -2.080450259 | ENSG00000184967 | 79050 | PWB-MSC UP | NOC4L | 0.6754784 |
| ENSG00000243753 | 3139 | PWB-iPSC UP | HLA-L | 0.983531 | ENSG00000152804 | 3087 | PWB-iPSC DN | HHEX | -2.096912072 | ENSG00000010292 | 9918 | PWB-MSC UP | NCAPD2 | 0.6750514 |
| ENSG00000249859 | 5820 | PWB-iPSC UP | PVT1 | 0.979196 | ENSG00000161681 | 50944 | PWB-iPSC DN | SHANK1 | -2.109178091 | ENSG00000151131 | 121053 | PWB-MSC UP | C12orf45 | 0.6697093 |
| ENSG00000236778 | 100507398 | PWB-iPSC UP | INTS6-AS1 | 0.976466 | ENSG00000105472 | 6320 | PWB-iPSC DN | CLEC11A | -2.114967242 | ENSG00000167118 | 81605 | PWB-MSC UP | URM1 | 0.6696491 |
| ENSG00000212907 | 4539 | PWB-iPSC UP | ND4L | 0.974529 | ENSG00000178623 | 2859 | PWB-iPSC DN | GPR35 | -2.118554166 | ENSG00000139726 | 8562 | PWB-MSC UP | DENR | 0.6694697 |
| ENSG00000119900 | 79627 | PWB-iPSC UP | OGFRL1 | 0.973411 | ENSG00000172478 | 79919 | PWB-iPSC DN | MAB21L4 | -2.139255157 | ENSG00000111669 | 7167 | PWB-MSC UP | TPI1 | 0.6691631 |
| ENSG00000136895 | 84253 | PWB-iPSC UP | GARNL3 | 0.972389 | ENSG00000101327 | 5173 | PWB-iPSC DN | PDYN | -2.151235238 | ENSG00000139182 | 9746 | PWB-MSC UP | CLSTN3 | 0.6680067 |
| ENSG00000198482 | 388558 | PWB-iPSC UP | ZNF808 | 0.971104 | ENSG00000092607 | 6913 | PWB-iPSC DN | TBX15 | -2.17280142 | ENSG00000222489 | 109616976 | PWB-MSC UP | SNORA79B | 0.6669639 |
| ENSG00000205771 | 440278 | PWB-iPSC UP | CATSPER2P1 | 0.970942 | ENSG00000170542 | 5272 | PWB-iPSC DN | SERPINB9 | -2.173458632 | ENSG00000111639 | 51258 | PWB-MSC UP | MRPL51 | 0.6648577 |
| ENSG00000255423 | 55096 | PWB-iPSC UP | EBLN2 | 0.970244 | ENSG00000125657 | 8744 | PWB-iPSC DN | TNFSF9 | -2.196837752 | ENSG00000136720 | 9394 | PWB-MSC UP | HS6ST1 | 0.6647818 |
| ENSG00000231999 | 400761 | PWB-iPSC UP | LRRC8C-DT | 0.969858 | ENSG00000171533 | 4135 | PWB-iPSC DN | MAP6 | -2.203255324 | ENSG00000066117 | 6602 | PWB-MSC UP | SMARCD1 | 0.6608567 |
| ENSG00000170160 | 9720 | PWB-iPSC UP | CCDC144A | 0.969688 | ENSG00000258484 | 246777 | PWB-iPSC DN | SPESP1 | -2.204695223 | ENSG00000257599 | 101055625 | PWB-MSC UP | OVCH1-AS1 | 0.6601265 |
| ENSG00000198198 | 23334 | PWB-iPSC UP | SZT2 | 0.96912 | ENSG00000180340 | 2535 | PWB-iPSC DN | FZD2 | -2.204746062 | ENSG00000047621 | 57102 | PWB-MSC UP | C12orf4 | 0.6590771 |
| ENSG00000163050 | 56997 | PWB-iPSC UP | COQ8A | 0.969019 | ENSG00000177508 | 79191 | PWB-iPSC DN | IRX3 | -2.212918726 | ENSG00000060138 | 8531 | PWB-MSC UP | YBX3 | 0.6587259 |
| ENSG00000197016 | 388566 | PWB-iPSC UP | ZNF470 | 0.967839 | ENSG00000168874 | 84913 | PWB-iPSC DN | ATOH8 | -2.220290595 | ENSG00000111676 | 1822 | PWB-MSC UP | ATN1 | 0.6566089 |
| ENSG00000179406 | 285908 | PWB-iPSC UP | LINC00174 | 0.967 | ENSG00000165140 | 2203 | PWB-iPSC DN | FBP1 | -2.221845653 | ENSG00000162086 | 7627 | PWB-MSC UP | ZNF75A | 0.6537009 |
| ENSG00000203875 | 387066 | PWB-iPSC UP | SNHG5 | 0.964517 | ENSG00000180422 | 283860 | PWB-iPSC DN | LINC00304 | -2.244119602 | ENSG00000135148 | 10906 | PWB-MSC UP | TRAFD1 | 0.6485263 |
| ENSG00000122481 | 25950 | PWB-iPSC UP | RWDD3 | 0.964209 | ENSG00000182870 | 50614 | PWB-iPSC DN | GALNT9 | -2.24544766 | ENSG00000196814 | 89853 | PWB-MSC UP | MVB12B | 0.6483021 |
| ENSG00000197608 | 284371 | PWB-iPSC UP | ZNF841 | 0.961383 | ENSG00000196433 | 438 | PWB-iPSC DN | ASMT | -2.253441857 | ENSG00000126749 | 10436 | PWB-MSC UP | EMG1 | 0.6448774 |
| ENSG00000258725 | 100507118 | PWB-iPSC UP | PRC1-AS1 | 0.956894 | ENSG00000155093 | 5799 | PWB-iPSC DN | PTPRN2 | -2.258914473 | ENSG00000161981 | 79622 | PWB-MSC UP | SNRNP25 | 0.6446972 |
| ENSG00000198712 | 4513 | PWB-iPSC UP | COX2 | 0.956037 | ENSG00000165588 | 5015 | PWB-iPSC DN | OTX2 | -2.281648473 | ENSG00000166908 | 79837 | PWB-MSC UP | PIP4K2C | 0.644411 |
| ENSG00000162069 | 146439 | PWB-iPSC UP | BICDL2 | 0.955194 | ENSG00000162692 | 7412 | PWB-iPSC DN | VCAM1 | -2.286384773 | ENSG00000111057 | 3875 | PWB-MSC UP | KRT18 | 0.6435957 |
| ENSG00000101104 | 80336 | PWB-iPSC UP | PABPC1L | 0.954304 | ENSG00000101336 | 3055 | PWB-iPSC DN | HCK | -2.295625567 | ENSG00000174437 | 488 | PWB-MSC UP | ATP2A2 | 0.6435541 |
| ENSG00000137501 | 54843 | PWB-iPSC UP | SYTL2 | 0.949976 | ENSG00000104435 | 11075 | PWB-iPSC DN | STMN2 | -2.299868393 | ENSG00000129317 | 83448 | PWB-MSC UP | PUS7L | 0.6430212 |
| ENSG00000183091 | 4703 | PWB-iPSC UP | NEB | 0.949196 | ENSG00000178573 | 4094 | PWB-iPSC DN | MAF | -2.320211396 | ENSG00000111640 | 2597 | PWB-MSC UP | GAPDH | 0.642267 |
| ENSG00000185513 | 26013 | PWB-iPSC UP | L3MBTL1 | 0.948666 | ENSG00000149557 | 9638 | PWB-iPSC DN | FEZ1 | -2.321628434 | ENSG00000068654 | 25885 | PWB-MSC UP | POLR1A | 0.6422142 |
| ENSG00000260287 | 101060321 | PWB-iPSC UP | TBC1D3G | 0.946526 | ENSG00000112619 | 5961 | PWB-iPSC DN | PRPH2 | -2.327772563 | ENSG00000156650 | 23522 | PWB-MSC UP | KAT6B | 0.6418618 |
| ENSG00000081052 | 1286 | PWB-iPSC UP | COL4A4 | 0.941413 | ENSG00000167617 | 148170 | PWB-iPSC DN | CDC42EP5 | -2.330776453 | ENSG00000111696 | 51559 | PWB-MSC UP | NT5DC3 | 0.6417483 |
| ENSG00000221923 | 400713 | PWB-iPSC UP | ZNF880 | 0.94103 | ENSG00000198125 | 4151 | PWB-iPSC DN | MB | -2.336186991 | ENSG00000181222 | 5430 | PWB-MSC UP | POLR2A | 0.6409192 |
| ENSG00000184507 | 256646 | PWB-iPSC UP | NUTM1 | 0.940548 | ENSG00000064195 | 1747 | PWB-iPSC DN | DLX3 | -2.347724364 | ENSG00000125877 | 3704 | PWB-MSC UP | ITPA | 0.640306 |
| ENSG00000033122 | 57554 | PWB-iPSC UP | LRRC7 | 0.93946 | ENSG00000137872 | 80031 | PWB-iPSC DN | SEMA6D | -2.35756769 | ENSG00000106066 | 54504 | PWB-MSC UP | CPVL | 0.6399024 |
| ENSG00000187105 | 399671 | PWB-iPSC UP | HEATR4 | 0.939147 | ENSG00000144481 | 79054 | PWB-iPSC DN | TRPM8 | -2.370296426 | ENSG00000061273 | 51564 | PWB-MSC UP | HDAC7 | 0.6396043 |
| ENSG00000250579 | 101929176 | PWB-iPSC UP | CTD-2297D10.2 | 0.937877 | ENSG00000273604 | 100170841 | PWB-iPSC DN | EPOP | -2.389643226 | ENSG00000167535 | 784 | PWB-MSC UP | CACNB3 | 0.6395633 |
| ENSG00000245213 | 101930370 | PWB-iPSC UP | LOC101930370 | 0.937514 | ENSG00000071909 | 140469 | PWB-iPSC DN | MYO3B | -2.400899831 | ENSG00000166226 | 10576 | PWB-MSC UP | CCT2 | 0.636378 |
| ENSG00000266173 | 92335 | PWB-iPSC UP | STRADA | 0.936983 | ENSG00000145284 | 79966 | PWB-iPSC DN | SCD5 | -2.403853483 | ENSG00000162013 | 1476 | PWB-MSC UP | CSTB | 0.6356142 |
| ENSG00000076067 | 5939 | PWB-iPSC UP | RBMS2 | 0.935485 | ENSG00000070031 | 6343 | PWB-iPSC DN | SCT | -2.425917569 | ENSG00000089220 | 5037 | PWB-MSC UP | PEBP1 | 0.6341424 |
| ENSG00000104728 | 9639 | PWB-iPSC UP | ARHGEF10 | 0.9339 | ENSG00000127863 | 55504 | PWB-iPSC DN | TNFRSF19 | -2.438659625 | ENSG00000091972 | 4345 | PWB-MSC UP | CD200 | 0.6339261 |
| ENSG00000162341 | 219931 | PWB-iPSC UP | TPCN2 | 0.932984 | ENSG00000120149 | 4488 | PWB-iPSC DN | MSX2 | -2.440956912 | ENSG00000151650 | 27287 | PWB-MSC UP | VENTX | 0.6335106 |
| ENSG00000205238 | 441273 | PWB-iPSC UP | SPDYE2 | 0.932193 | ENSG00000135248 | 84691 | PWB-iPSC DN | FAM71F1 | -2.482772448 | ENSG00000196247 | 51427 | PWB-MSC UP | ZNF107 | 0.6326303 |
| ENSG00000237298 | 100506866 | PWB-iPSC UP | TTN-AS1 | 0.931693 | ENSG00000125848 | 23767 | PWB-iPSC DN | FLRT3 | -2.483712968 | ENSG00000110851 | 11108 | PWB-MSC UP | PRDM4 | 0.6304886 |
| ENSG00000110888 | 65981 | PWB-iPSC UP | CAPRIN2 | 0.930761 | ENSG00000179344 | 3119 | PWB-iPSC DN | HLA-DQB1 | -2.487813038 | ENSG00000110107 | 27339 | PWB-MSC UP | PRPF19 | 0.6283751 |
| ENSG00000149292 | 54970 | PWB-iPSC UP | TTC12 | 0.93006 | ENSG00000085276 | 2122 | PWB-iPSC DN | MECOM | -2.491827209 | ENSG00000139197 | 5830 | PWB-MSC UP | PEX5 | 0.6274222 |
| ENSG00000186648 | 90668 | PWB-iPSC UP | CARMIL3 | 0.929688 | ENSG00000122756 | 1271 | PWB-iPSC DN | CNTFR | -2.510053062 | ENSG00000151148 | 89910 | PWB-MSC UP | UBE3B | 0.6271608 |
| ENSG00000198203 | 6819 | PWB-iPSC UP | SULT1C2 | 0.929507 | ENSG00000112183 | 221662 | PWB-iPSC DN | RBM24 | -2.527438593 | ENSG00000157837 | 121665 | PWB-MSC UP | SPPL3 | 0.6266291 |
| ENSG00000147862 | 4781 | PWB-iPSC UP | NFIB | 0.929413 | ENSG00000179750 | 9582 | PWB-iPSC DN | APOBEC3B | -2.533590687 | ENSG00000206527 | 201562 | PWB-MSC UP | HACD2 | 0.6260717 |
| ENSG00000251022 | 100499177 | PWB-iPSC UP | THAP9-AS1 | 0.929073 | ENSG00000188219 | 445582 | PWB-iPSC DN | POTEE | -2.535624974 | ENSG00000111364 | 57696 | PWB-MSC UP | DDX55 | 0.6238799 |
| ENSG00000231312 | 728730 | PWB-iPSC UP | MAP4K3-DT | 0.925646 | ENSG00000169218 | 284654 | PWB-iPSC DN | RSPO1 | -2.544863187 | ENSG00000146556 | 375260 | PWB-MSC UP | WASH2P | 0.6237409 |
| ENSG00000187775 | 8632 | PWB-iPSC UP | DNAH17 | 0.924106 | ENSG00000142606 | 79258 | PWB-iPSC DN | MMEL1 | -2.557404521 | ENSG00000165349 | 84889 | PWB-MSC UP | SLC7A3 | 0.6232327 |
| ENSG00000206077 | 653082 | PWB-iPSC UP | ZDHHC11B | 0.923794 | ENSG00000144152 | 129804 | PWB-iPSC DN | FBLN7 | -2.56137321 | ENSG00000080824 | 3320 | PWB-MSC UP | HSP90AA1 | 0.6226431 |
| ENSG00000132294 | 23167 | PWB-iPSC UP | EFR3A | 0.92343 | ENSG00000110799 | 7450 | PWB-iPSC DN | VWF | -2.577407604 | ENSG00000160214 | 8568 | PWB-MSC UP | RRP1 | 0.6190354 |
| ENSG00000255468 | 102724064 | PWB-iPSC UP | LOC102724064 | 0.920423 | ENSG00000149090 | 25891 | PWB-iPSC DN | PAMR1 | -2.577430374 | ENSG00000256223 | 7556 | PWB-MSC UP | ZNF10 | 0.6188424 |
| ENSG00000136010 | 160428 | PWB-iPSC UP | ALDH1L2 | 0.917711 | ENSG00000196415 | 5657 | PWB-iPSC DN | PRTN3 | -2.578471907 | ENSG00000007062 | 8842 | PWB-MSC UP | PROM1 | 0.6169195 |
| ENSG00000231721 | 378805 | PWB-iPSC UP | LINC-PINT | 0.916895 | ENSG00000128383 | 200315 | PWB-iPSC DN | APOBEC3A | -2.614787526 | ENSG00000198464 | 147657 | PWB-MSC UP | ZNF480 | 0.6139126 |
| ENSG00000198804 | 4512 | PWB-iPSC UP | COX1 | 0.916205 | ENSG00000174963 | 84107 | PWB-iPSC DN | ZIC4 | -2.615063029 | ENSG00000173598 | 11163 | PWB-MSC UP | NUDT4 | 0.6127892 |
| ENSG00000169314 | 150248 | PWB-iPSC UP | C22orf15 | 0.915971 | ENSG00000259439 | 107985879 | PWB-iPSC DN | LINC01833 | -2.626718424 | ENSG00000089154 | 10985 | PWB-MSC UP | GCN1 | 0.6116351 |
| ENSG00000218739 | 100505876 | PWB-iPSC UP | CEBPZOS | 0.915684 | ENSG00000113100 | 1007 | PWB-iPSC DN | CDH9 | -2.651846782 | ENSG00000102882 | 5595 | PWB-MSC UP | MAPK3 | 0.6113283 |
| ENSG00000225484 | 101060691 | PWB-iPSC UP | NUTM2B-AS1 | 0.915419 | ENSG00000134138 | 4212 | PWB-iPSC DN | MEIS2 | -2.665018337 | ENSG00000148719 | 54788 | PWB-MSC UP | DNAJB12 | 0.6112102 |
| ENSG00000267254 | 284408 | PWB-iPSC UP | ZNF790-AS1 | 0.913939 | ENSG00000115461 | 3488 | PWB-iPSC DN | IGFBP5 | -2.674157673 | ENSG00000100379 | 79734 | PWB-MSC UP | KCTD17 | 0.6108201 |
| ENSG00000115339 | 2591 | PWB-iPSC UP | GALNT3 | 0.91338 | ENSG00000177335 | 286122 | PWB-iPSC DN | C8orf31 | -2.674568411 | ENSG00000179195 | 144348 | PWB-MSC UP | ZNF664 | 0.6099697 |
| ENSG00000152583 | 8404 | PWB-iPSC UP | SPARCL1 | 0.913058 | ENSG00000119242 | 80212 | PWB-iPSC DN | CCDC92 | -2.675773924 | ENSG00000063244 | 11338 | PWB-MSC UP | U2AF2 | 0.6089122 |
| ENSG00000090376 | 11213 | PWB-iPSC UP | IRAK3 | 0.907078 | ENSG00000273540 | 123624 | PWB-iPSC DN | AGBL1 | -2.678627943 | ENSG00000111229 | 10094 | PWB-MSC UP | ARPC3 | 0.607904 |
| ENSG00000234327 | 101928000 | PWB-iPSC UP | LOC101928000 | 0.905048 | ENSG00000007372 | 5080 | PWB-iPSC DN | PAX6 | -2.689675339 | ENSG00000239306 | 10432 | PWB-MSC UP | RBM14 | 0.607798 |
| ENSG00000214837 | 731275 | PWB-iPSC UP | LINC01347 | 0.904446 | ENSG00000169548 | 129025 | PWB-iPSC DN | ZNF280A | -2.705435502 | ENSG00000270647 | 8148 | PWB-MSC UP | TAF15 | 0.6060901 |
| ENSG00000186889 | 200728 | PWB-iPSC UP | TMEM17 | 0.904258 | ENSG00000133321 | 5920 | PWB-iPSC DN | PLAAT4 | -2.706067613 | ENSG00000135372 | 55226 | PWB-MSC UP | NAT10 | 0.6051855 |
| ENSG00000177108 | 283576 | PWB-iPSC UP | ZDHHC22 | 0.902038 | ENSG00000172782 | 283985 | PWB-iPSC DN | FADS6 | -2.753084567 | ENSG00000135503 | 91 | PWB-MSC UP | ACVR1B | 0.6019803 |
| ENSG00000144395 | 284992 | PWB-iPSC UP | CCDC150 | 0.901487 | ENSG00000148948 | 57689 | PWB-iPSC DN | LRRC4C | -2.760018525 | ENSG00000139644 | 7009 | PWB-MSC UP | TMBIM6 | 0.6016362 |
| ENSG00000215908 | 84809 | PWB-iPSC UP | CROCCP2 | 0.901245 | ENSG00000100867 | 10202 | PWB-iPSC DN | DHRS2 | -2.778695273 | ENSG00000105926 | 51678 | PWB-MSC UP | MPP6 | 0.6003744 |
| ENSG00000167562 | 55762 | PWB-iPSC UP | ZNF701 | 0.900196 | ENSG00000126709 | 2537 | PWB-iPSC DN | IFI6 | -2.792519622 | ENSG00000148334 | 80142 | PWB-MSC UP | PTGES2 | 0.5990659 |
| ENSG00000173209 | 130872 | PWB-iPSC UP | AHSA2P | 0.900083 | ENSG00000162998 | 2487 | PWB-iPSC DN | FRZB | -2.797135844 | ENSG00000269343 | 100293516 | PWB-MSC UP | ZNF587B | 0.5970652 |
| ENSG00000080947 | 114819 | PWB-iPSC UP | CROCCP3 | 0.897545 | ENSG00000144410 | 130749 | PWB-iPSC DN | CPO | -2.827466314 | ENSG00000197111 | 5094 | PWB-MSC UP | PCBP2 | 0.5968317 |
| ENSG00000136603 | 6498 | PWB-iPSC UP | SKIL | 0.895275 | ENSG00000233608 | 117581 | PWB-iPSC DN | TWIST2 | -2.83067947 | ENSG00000176915 | 23141 | PWB-MSC UP | ANKLE2 | 0.595745 |
| ENSG00000181019 | 1728 | PWB-iPSC UP | NQO1 | 0.894766 | ENSG00000040731 | 1008 | PWB-iPSC DN | CDH10 | -2.835853417 | ENSG00000167562 | 55762 | PWB-MSC UP | ZNF701 | 0.5951745 |
| ENSG00000178038 | 259173 | PWB-iPSC UP | ALS2CL | 0.893355 | ENSG00000125285 | 11166 | PWB-iPSC DN | SOX21 | -2.840083558 | ENSG00000100591 | 10598 | PWB-MSC UP | AHSA1 | 0.5937778 |
| ENSG00000257337 | 283335 | PWB-iPSC UP | LOC283335 | 0.892982 | ENSG00000182168 | 8633 | PWB-iPSC DN | UNC5C | -2.844820865 | ENSG00000111142 | 10988 | PWB-MSC UP | METAP2 | 0.5937508 |
| ENSG00000188130 | 6300 | PWB-iPSC UP | MAPK12 | 0.89264 | ENSG00000198914 | 5455 | PWB-iPSC DN | POU3F3 | -2.848321683 | ENSG00000111678 | 113246 | PWB-MSC UP | C12orf57 | 0.5924076 |
| ENSG00000147174 | 93953 | PWB-iPSC UP | GCNA | 0.889659 | ENSG00000131409 | 94030 | PWB-iPSC DN | LRRC4B | -2.855533018 | ENSG00000111602 | 8914 | PWB-MSC UP | TIMELESS | 0.5923872 |
| ENSG00000174353 | 378108 | PWB-iPSC UP | TRIM74 | 0.889143 | ENSG00000204335 | 389058 | PWB-iPSC DN | SP5 | -2.86689932 | ENSG00000237440 | 100129842 | PWB-MSC UP | ZNF737 | 0.5907176 |
| ENSG00000115705 | 7173 | PWB-iPSC UP | TPO | 0.888751 | ENSG00000130303 | 684 | PWB-iPSC DN | BST2 | -2.899685353 | ENSG00000183495 | 57634 | PWB-MSC UP | EP400 | 0.58938 |
| ENSG00000143801 | 5664 | PWB-iPSC UP | PSEN2 | 0.886978 | ENSG00000140678 | 3687 | PWB-iPSC DN | ITGAX | -2.910284425 | ENSG00000180263 | 55785 | PWB-MSC UP | FGD6 | 0.5882565 |
| ENSG00000230922 | 101927551 | PWB-iPSC UP | LOC101927551 | 0.884616 | ENSG00000185745 | 3434 | PWB-iPSC DN | IFIT1 | -2.936434373 | ENSG00000075856 | 9733 | PWB-MSC UP | SART3 | 0.5878438 |
| ENSG00000185864 | 440345 | PWB-iPSC UP | NPIPB4 | 0.884213 | ENSG00000256576 | 100996246 | PWB-iPSC DN | LINC02361 | -2.940281871 | ENSG00000175727 | 22877 | PWB-MSC UP | MLXIP | 0.5870609 |
| ENSG00000241343 | 6173 | PWB-iPSC UP | RPL36A | 0.884186 | ENSG00000283196 | 107985911 | PWB-iPSC DN | LOC10798591 | -2.956473106 | ENSG00000129757 | 1028 | PWB-MSC UP | CDKN1C | 0.5866109 |
| ENSG00000047662 | 27146 | PWB-iPSC UP | FAM184B | 0.883091 | ENSG00000095596 | 1592 | PWB-iPSC DN | CYP26A1 | -2.98323289 | ENSG00000071889 | 60343 | PWB-MSC UP | FAM3A | 0.5839519 |
| ENSG00000146856 | 340351 | PWB-iPSC UP | AGBL3 | 0.881104 | ENSG00000128918 | 8854 | PWB-iPSC DN | ALDH1A2 | -2.999904788 | ENSG00000101057 | 4605 | PWB-MSC UP | MYBL2 | 0.5820639 |
| ENSG00000171502 | 255631 | PWB-iPSC UP | COL24A1 | 0.880683 | ENSG00000116106 | 2043 | PWB-iPSC DN | EPHA4 | -3.054132981 | ENSG00000111445 | 5985 | PWB-MSC UP | RFC5 | 0.5816039 |

| | | | | | | | | | | | | | | |
|---|---|---|---|---|---|---|---|---|---|---|---|---|---|---|
| ENSG00000281183 | 101241892 | PWB-iPSC UP | NPTN-IT1 | 0.880579 | ENSG00000221882 | 4995 | PWB-iPSC DN | OR3A2 | -3.063672527 | ENSG00000059804 | 6515 | PWB-MSC UP | SLC2A3 | 0.5801648 |
| ENSG00000196912 | 57730 | PWB-iPSC UP | ANKRD36B | 0.880461 | ENSG00000196604 | 728378 | PWB-iPSC DN | POTEF | -3.068018975 | ENSG00000185480 | 55010 | PWB-MSC UP | PARPBP | 0.5791844 |
| ENSG00000196693 | 7582 | PWB-iPSC UP | ZNF33B | 0.879884 | ENSG00000180155 | 66004 | PWB-iPSC DN | LYNX1 | -3.081680748 | ENSG00000073111 | 4171 | PWB-MSC UP | MCM2 | 0.5784442 |
| ENSG00000170835 | 1056 | PWB-iPSC UP | CEL | 0.879384 | ENSG00000141449 | 80000 | PWB-iPSC DN | GREB1L | -3.084478862 | ENSG00000174243 | 9416 | PWB-MSC UP | DDX23 | 0.5775043 |
| ENSG00000134716 | 1573 | PWB-iPSC UP | CYP2J2 | 0.876065 | ENSG00000100095 | 23544 | PWB-iPSC DN | SEZ6L | -3.101113017 | ENSG00000111667 | 8078 | PWB-MSC UP | USP5 | 0.5756559 |
| ENSG00000170954 | 55786 | PWB-iPSC UP | ZNF415 | 0.874254 | ENSG00000261456 | 347688 | PWB-iPSC DN | TUBB8 | -3.13243299 | ENSG00000057294 | 5318 | PWB-MSC UP | PKP2 | 0.5741854 |
| ENSG00000225366 | 6998 | PWB-iPSC UP | TDGF1P3 | 0.874065 | ENSG00000171954 | 126410 | PWB-iPSC DN | CYP4F22 | -3.142279299 | ENSG00000165891 | 144455 | PWB-MSC UP | E2F7 | 0.5680358 |
| ENSG00000149591 | 6876 | PWB-iPSC UP | TAGLN | 0.872579 | ENSG00000148677 | 27063 | PWB-iPSC DN | ANKRD1 | -3.146291188 | ENSG00000139719 | 65082 | PWB-MSC UP | VPS33A | 0.5674914 |
| ENSG00000214967 | 101059938 | PWB-iPSC UP | NPIPA7 | 0.868816 | ENSG00000166450 | 283659 | PWB-iPSC DN | PRTG | -3.15350966 | ENSG00000161813 | 113251 | PWB-MSC UP | LARP4 | 0.5632319 |
| ENSG00000197497 | 79788 | PWB-iPSC UP | ZNF665 | 0.868549 | ENSG00000169877 | 51327 | PWB-iPSC DN | AHSP | -3.213369946 | ENSG00000126746 | 171017 | PWB-MSC UP | ZNF384 | 0.5618777 |
| ENSG00000132199 | 55556 | PWB-iPSC UP | ENOSF1 | 0.867799 | ENSG00000148516 | 6935 | PWB-iPSC DN | ZEB1 | -3.237677796 | ENSG00000169515 | 83987 | PWB-MSC UP | CCDC8 | 0.5614483 |
| ENSG00000247828 | 100505894 | PWB-iPSC UP | TMEM161B-AS1 | 0.867793 | ENSG00000229314 | 5004 | PWB-iPSC DN | ORM1 | -3.289014814 | ENSG00000171552 | 598 | PWB-MSC UP | BCL2L1 | 0.5611623 |
| ENSG00000166105 | 112937 | PWB-iPSC UP | GLB1L3 | 0.867719 | ENSG00000263563 | 23666 | PWB-iPSC DN | UBBP4 | -3.293640076 | ENSG00000074071 | 65993 | PWB-MSC UP | MRPS34 | 0.561118 |
| ENSG00000154040 | 26256 | PWB-iPSC UP | CABYR | 0.867362 | ENSG00000152977 | 7545 | PWB-iPSC DN | ZIC1 | -3.34934201 | ENSG00000198482 | 388558 | PWB-MSC UP | ZNF808 | 0.5609902 |
| ENSG00000166405 | 79608 | PWB-iPSC UP | RIC3 | 0.866694 | ENSG00000106689 | 9355 | PWB-iPSC DN | LHX2 | -3.366713708 | ENSG00000127586 | 63922 | PWB-MSC UP | CHTF18 | 0.5589614 |
| ENSG00000198346 | 126017 | PWB-iPSC UP | ZNF813 | 0.864763 | ENSG00000009709 | 5081 | PWB-iPSC DN | PAX7 | -3.461316223 | ENSG00000139433 | 51228 | PWB-MSC UP | GLTP | 0.5583463 |
| ENSG00000182108 | 28955 | PWB-iPSC UP | DEXI | 0.864694 | ENSG00000188257 | 5320 | PWB-iPSC DN | PLA2G2A | -3.481293421 | ENSG00000177889 | 7334 | PWB-MSC UP | UBE2N | 0.5566151 |
| ENSG00000110090 | 1374 | PWB-iPSC UP | CPT1A | 0.864576 | ENSG00000182393 | 282618 | PWB-iPSC DN | IFNL1 | -3.489138491 | ENSG00000123136 | 10212 | PWB-MSC UP | DDX39A | 0.5563958 |
| ENSG00000188897 | 400499 | PWB-iPSC UP | LOC400499 | 0.861611 | ENSG00000061492 | 7478 | PWB-iPSC DN | WNT8A | -3.491723116 | ENSG00000143569 | 9898 | PWB-MSC UP | UBAP2L | 0.5554468 |
| ENSG00000174669 | 3177 | PWB-iPSC UP | SLC29A2 | 0.860682 | ENSG00000089116 | 64211 | PWB-iPSC DN | LHX5 | -3.505777421 | ENSG00000170421 | 3856 | PWB-MSC UP | KRT8 | 0.553997 |
| ENSG00000140993 | 91151 | PWB-iPSC UP | TIGD7 | 0.860176 | ENSG00000108231 | 9211 | PWB-iPSC DN | LGI1 | -3.505829777 | ENSG00000004455 | 204 | PWB-MSC UP | AK2 | 0.5523727 |
| ENSG00000243716 | 100132247 | PWB-iPSC UP | NPIPB5 | 0.859588 | ENSG00000119917 | 3437 | PWB-iPSC DN | IFIT3 | -3.506758809 | ENSG00000196172 | 148213 | PWB-MSC UP | ZNF681 | 0.5523429 |
| ENSG00000140451 | 80119 | PWB-iPSC UP | PIF1 | 0.857295 | ENSG00000198105 | 57209 | PWB-iPSC DN | ZNF248 | -3.509235898 | ENSG00000184992 | 140707 | PWB-MSC UP | BRI3BP | 0.5518454 |
| ENSG00000157036 | 9941 | PWB-iPSC UP | EXOG | 0.857177 | ENSG00000039987 | 54831 | PWB-iPSC DN | BEST2 | -3.515563304 | ENSG00000087269 | 8602 | PWB-MSC UP | NOP14 | 0.5503452 |
| ENSG00000249915 | 10016 | PWB-iPSC UP | PDCD6 | 0.855301 | ENSG00000198732 | 64093 | PWB-iPSC DN | SMOC1 | -3.530478906 | ENSG00000186185 | 146909 | PWB-MSC UP | KIF18B | 0.5483955 |
| ENSG00000160172 | 645332 | PWB-iPSC UP | FAM86C2P | 0.853223 | ENSG00000230316 | 154860 | PWB-iPSC DN | FEZF1-AS1 | -3.530814907 | ENSG00000142449 | 84467 | PWB-MSC UP | FBN3 | 0.5467693 |
| ENSG00000249715 | 90342 | PWB-iPSC UP | FER1L5 | 0.849691 | ENSG00000164756 | 169026 | PWB-iPSC DN | SLC30A8 | -3.534010784 | ENSG00000110955 | 506 | PWB-MSC UP | ATP5F1B | 0.5406725 |
| ENSG00000103740 | 23205 | PWB-iPSC UP | ACSBG1 | 0.845286 | ENSG00000256288 | 113523642 | PWB-iPSC DN | LINC02617 | -3.576429226 | ENSG00000140365 | 54939 | PWB-MSC UP | COMMD4 | 0.5405695 |
| ENSG00000164620 | 285613 | PWB-iPSC UP | RELL2 | 0.841983 | ENSG00000169908 | 4071 | PWB-iPSC DN | TM4SF1 | -3.734995329 | ENSG00000072609 | 55743 | PWB-MSC UP | CHFR | 0.5403617 |
| ENSG00000171451 | 92126 | PWB-iPSC UP | DSEL | 0.841129 | ENSG00000075884 | 55843 | PWB-iPSC DN | ARHGAP15 | -3.782762618 | ENSG00000172336 | 10248 | PWB-MSC UP | POP7 | 0.5398274 |
| ENSG00000232859 | 201229 | PWB-iPSC UP | LYRM9 | 0.838034 | ENSG00000134438 | 30062 | PWB-iPSC DN | RAX | -3.823908484 | ENSG00000111328 | 8099 | PWB-MSC UP | CDK2AP1 | 0.5392317 |
| ENSG00000275832 | 57636 | PWB-iPSC UP | ARHGAP23 | 0.835872 | ENSG00000166407 | 4004 | PWB-iPSC DN | LMO1 | -3.838363517 | ENSG00000076770 | 55796 | PWB-MSC UP | MBNL3 | 0.5383898 |
| ENSG00000234719 | 729978 | PWB-iPSC UP | NPIPB2 | 0.834949 | ENSG00000153266 | 55079 | PWB-iPSC DN | FEZF2 | -3.844089835 | ENSG00000084463 | 51729 | PWB-MSC UP | WBP11 | 0.5382824 |
| ENSG00000273513 | 101060351 | PWB-iPSC UP | TBC1D3K | 0.833558 | ENSG00000105392 | 1406 | PWB-iPSC DN | CRX | -3.878659261 | ENSG00000135404 | 967 | PWB-MSC UP | CD63 | 0.5362801 |
| ENSG00000135317 | 57231 | PWB-iPSC UP | SNX14 | 0.830706 | ENSG00000137959 | 10964 | PWB-iPSC DN | IFI44L | -3.901519639 | ENSG00000172301 | 55352 | PWB-MSC UP | COPRS | 0.5349965 |
| ENSG00000242028 | 25764 | PWB-iPSC UP | HYPK | 0.830521 | ENSG00000205856 | 150297 | PWB-iPSC DN | C22orf42 | -3.917408371 | ENSG00000121064 | 59342 | PWB-MSC UP | SCPEP1 | 0.5327495 |
| ENSG00000253731 | 56109 | PWB-iPSC UP | PCDHGA6 | 0.828439 | ENSG00000187553 | 340665 | PWB-iPSC DN | CYP26C1 | -3.918202202 | ENSG00000166532 | 57494 | PWB-MSC UP | RIMKLB | 0.5313144 |
| ENSG00000103335 | 9780 | PWB-iPSC UP | PIEZO1 | 0.828336 | ENSG00000196092 | 5079 | PWB-iPSC DN | PAX5 | -3.919560741 | ENSG00000169627 | 654483 | PWB-MSC UP | BOLA2B | 0.5307954 |
| ENSG00000144642 | 27303 | PWB-iPSC UP | RBMS3 | 0.826735 | ENSG00000183206 | 388468 | PWB-iPSC DN | POTEC | -3.923833138 | ENSG00000123416 | 10376 | PWB-MSC UP | TUBA1B | 0.527258 |
| ENSG00000167525 | 147011 | PWB-iPSC UP | PROCA1 | 0.826409 | ENSG00000111335 | 4939 | PWB-iPSC DN | OAS2 | -3.944111063 | ENSG00000138778 | 1062 | PWB-MSC UP | CENPE | 0.526468 |
| ENSG00000274226 | 729877 | PWB-iPSC UP | TBC1D3H | 0.824463 | ENSG00000186675 | 139599 | PWB-iPSC DN | MAGEE2 | -3.945754674 | ENSG00000089094 | 84678 | PWB-MSC UP | KDM2B | 0.5249636 |
| ENSG00000254860 | 493900 | PWB-iPSC UP | TMEM9B-AS1 | 0.823489 | ENSG00000185904 | 84856 | PWB-iPSC DN | LINC00839 | -3.950252531 | ENSG00000143842 | 9580 | PWB-MSC UP | SOX13 | 0.5245169 |
| ENSG00000140563 | 55784 | PWB-iPSC UP | MCTP2 | 0.823476 | ENSG00000162761 | 4009 | PWB-iPSC DN | LMX1A | -3.959026266 | ENSG00000139372 | 6996 | PWB-MSC UP | TDG | 0.5233384 |
| ENSG00000101194 | 63910 | PWB-iPSC UP | SLC17A9 | 0.822891 | ENSG00000249310 | 100874530 | PWB-iPSC DN | APOBEC3B-AS | -3.991697469 | ENSG00000061794 | 60488 | PWB-MSC UP | MRPS35 | 0.5233371 |
| ENSG00000133121 | 90627 | PWB-iPSC UP | STARD13 | 0.822049 | ENSG00000167634 | 199713 | PWB-iPSC DN | NLRP7 | -3.997274769 | ENSG00000006831 | 79602 | PWB-MSC UP | ADIPOR2 | 0.5191833 |
| ENSG00000241015 | 147804 | PWB-iPSC UP | TPM3P9 | 0.822042 | ENSG00000140287 | 3067 | PWB-iPSC DN | HDC | -3.997858276 | ENSG00000111731 | 9847 | PWB-MSC UP | C2CD5 | 0.5147297 |
| ENSG00000002016 | 5893 | PWB-iPSC UP | RAD52 | 0.821763 | ENSG00000113361 | 1004 | PWB-iPSC DN | CDH6 | -4.041781653 | ENSG00000242485 | 55052 | PWB-MSC UP | MRPL20 | 0.5109267 |
| ENSG00000106991 | 2022 | PWB-iPSC UP | ENG | 0.819943 | ENSG00000136869 | 7099 | PWB-iPSC DN | TLR4 | -4.081011021 | ENSG00000196510 | 51434 | PWB-MSC UP | ANAPC7 | 0.5103448 |
| ENSG00000204961 | 9752 | PWB-iPSC UP | PCDHA9 | 0.81989 | ENSG00000166923 | 26585 | PWB-iPSC DN | GREM1 | -4.109505155 | ENSG00000070047 | 57661 | PWB-MSC UP | PHRF1 | 0.5093128 |
| ENSG00000185278 | 84614 | PWB-iPSC UP | ZBTB37 | 0.818886 | ENSG00000261667 | 101928108 | PWB-iPSC DN | LY6L | -4.179563641 | ENSG00000167965 | 64223 | PWB-MSC UP | MLST8 | 0.5073425 |
| ENSG00000050555 | 10319 | PWB-iPSC UP | LAMC3 | 0.818808 | ENSG00000179455 | 7681 | PWB-iPSC DN | MKRN3 | -4.205651388 | ENSG00000168476 | 80346 | PWB-MSC UP | REEP4 | 0.5071213 |
| ENSG00000227124 | 100131827 | PWB-iPSC UP | ZNF717 | 0.818749 | ENSG00000236373 | 101926942 | PWB-iPSC DN | LINC02653 | -4.331980036 | ENSG00000174106 | 23592 | PWB-MSC UP | LEMD3 | 0.5052974 |
| ENSG00000145623 | 9180 | PWB-iPSC UP | OSMR | 0.817951 | ENSG00000249378 | 401164 | PWB-iPSC DN | LINC01060 | -4.338443138 | ENSG00000132341 | 5901 | PWB-MSC UP | RAN | 0.5028283 |
| ENSG00000183426 | 9284 | PWB-iPSC UP | NPIPA1 | 0.816358 | ENSG00000145721 | 167410 | PWB-iPSC DN | LIX1 | -4.496248956 | ENSG00000123268 | 466 | PWB-MSC UP | ATF1 | 0.5016451 |
| ENSG00000151575 | 374618 | PWB-iPSC UP | TEX9 | 0.815669 | ENSG00000205212 | 339184 | PWB-iPSC DN | CCDC144NL | -4.554491305 | ENSG00000139613 | 6601 | PWB-MSC UP | SMARCC2 | 0.5013613 |
| ENSG00000134954 | 2113 | PWB-iPSC UP | ETS1 | 0.815583 | ENSG00000138083 | 6496 | PWB-iPSC DN | SIX3 | -4.55857251 | ENSG00000170515 | 5036 | PWB-MSC UP | PA2G4 | 0.5002988 |
| ENSG00000172197 | 154141 | PWB-iPSC UP | MBOAT1 | 0.815507 | ENSG00000128422 | 3872 | PWB-iPSC DN | KRT17 | -4.559574336 | ENSG00000003056 | 4074 | PWB-MSC UP | M6PR | 0.4963551 |
| ENSG00000187097 | 957 | PWB-iPSC UP | ENTPD5 | 0.814587 | ENSG00000197915 | 388697 | PWB-iPSC DN | HRNR | -4.618468401 | ENSG00000105486 | 3978 | PWB-MSC UP | LIG1 | 0.4959428 |
| ENSG00000169246 | 23117 | PWB-iPSC UP | NPIPB3 | 0.814429 | ENSG00000128610 | 389549 | PWB-iPSC DN | FEZF1 | -4.63701542 | ENSG00000182986 | 162967 | PWB-MSC UP | ZNF320 | 0.4954748 |
| ENSG00000180376 | 285331 | PWB-iPSC UP | CCDC66 | 0.814359 | ENSG00000130957 | 8789 | PWB-iPSC DN | FBP2 | -4.707748679 | ENSG00000104331 | 54928 | PWB-MSC UP | IMPAD1 | 0.493662 |
| ENSG00000237491 | 105378580 | PWB-iPSC UP | LINC01409 | 0.814039 | ENSG00000105880 | 1749 | PWB-iPSC DN | DLX5 | -4.733778953 | ENSG00000139651 | 283337 | PWB-MSC UP | ZNF740 | 0.4915677 |
| ENSG00000115963 | 390 | PWB-iPSC UP | RND3 | 0.812634 | ENSG00000236502 | 100506108 | PWB-iPSC DN | SIX3-AS1 | -4.823222028 | ENSG00000100401 | 5905 | PWB-MSC UP | RANGAP1 | 0.4902282 |
| ENSG00000151789 | 79750 | PWB-iPSC UP | ZNF385D | 0.810282 | ENSG00000135114 | 8638 | PWB-iPSC DN | OASL | -4.897032047 | ENSG00000184752 | 55967 | PWB-MSC UP | NDUFA12 | 0.4895867 |
| ENSG00000237440 | 100129842 | PWB-iPSC UP | ZNF737 | 0.810223 | ENSG00000271503 | 6352 | PWB-iPSC DN | CCL5 | -4.96525994 | ENSG00000041802 | 55341 | PWB-MSC UP | LSG1 | 0.4892902 |
| ENSG00000117616 | 57035 | PWB-iPSC UP | RSRP1 | 0.810207 | ENSG00000196735 | 3117 | PWB-iPSC DN | HLA-DQA1 | -5.087247878 | ENSG00000147684 | 4715 | PWB-MSC UP | NDUFB9 | 0.4876161 |
| ENSG00000274020 | 388685 | PWB-iPSC UP | LINC01138 | 0.808874 | ENSG00000237975 | 339400 | PWB-iPSC DN | FLG-AS1 | -5.148663428 | ENSG00000136938 | 10541 | PWB-MSC UP | ANP32B | 0.4859423 |
| ENSG00000006025 | 114881 | PWB-iPSC UP | OSBPL7 | 0.805664 | ENSG00000248874 | 439936 | PWB-iPSC DN | C5orf17 | -5.36752167 | ENSG00000149948 | 8091 | PWB-MSC UP | HMGA2 | 0.4847653 |
| ENSG00000164647 | 26872 | PWB-iPSC UP | STEAP1 | 0.805262 | ENSG00000168542 | 1281 | PWB-iPSC DN | COL3A1 | -5.40674362 | ENSG00000060237 | 65125 | PWB-MSC UP | WNK1 | 0.4845704 |
| ENSG00000248671 | 285407 | PWB-iPSC UP | ALG1L9P | 0.805201 | ENSG00000175745 | 7025 | PWB-iPSC DN | NR2F1 | -5.460449446 | ENSG00000196465 | 140465 | PWB-MSC UP | MYL6B | 0.4826053 |
| ENSG00000166482 | 4239 | PWB-iPSC UP | MFAP4 | 0.804811 | ENSG00000143631 | 2312 | PWB-iPSC DN | FLG | -5.511177403 | ENSG00000215021 | 11331 | PWB-MSC UP | PHB2 | 0.4791121 |
| ENSG00000074935 | 51175 | PWB-iPSC UP | TUBE1 | 0.803565 | ENSG00000089127 | 4938 | PWB-iPSC DN | OAS1 | -5.644851537 | ENSG00000070961 | 490 | PWB-MSC UP | ATP2B1 | 0.4748408 |
| ENSG00000247400 | 100289274 | PWB-iPSC UP | DNAJC3-DT | 0.802532 | ENSG00000138722 | 22915 | PWB-iPSC DN | MMRN1 | -5.669905574 | ENSG00000139645 | 283373 | PWB-MSC UP | ANKRD52 | 0.4733485 |
| ENSG00000163945 | 57654 | PWB-iPSC UP | UVSSA | 0.799098 | ENSG00000242082 | 110806273 | PWB-iPSC DN | SLC5A4-AS1 | -5.78637986 | ENSG00000023734 | 11171 | PWB-MSC UP | STRAP | 0.4732155 |
| ENSG00000226210 | 100288778 | PWB-iPSC UP | WASH8P | 0.798511 | ENSG00000147381 | 4103 | PWB-iPSC DN | MAGEA4 | -5.843426891 | ENSG00000175970 | 84747 | PWB-MSC UP | UNC119B | 0.4719627 |
| ENSG00000273899 | 79159 | PWB-iPSC UP | NOL12 | 0.79783 | ENSG00000185551 | 7026 | PWB-iPSC DN | NR2F2 | -5.870201013 | ENSG00000089234 | 8315 | PWB-MSC UP | BRAP | 0.4714591 |
| ENSG00000268471 | 54553 | PWB-iPSC UP | MIR4453HG | 0.796938 | ENSG00000185559 | 8788 | PWB-iPSC DN | DLK1 | -5.87129753 | ENSG00000110958 | 10728 | PWB-MSC UP | PTGES3 | 0.4682653 |
| ENSG00000132330 | 51540 | PWB-iPSC UP | SCLY | 0.795236 | ENSG00000188681 | 100132288 | PWB-iPSC DN | TEKT4P2 | -6.122908314 | ENSG00000111670 | 79158 | PWB-MSC UP | GNPTAB | 0.467812 |
| ENSG00000185338 | 8651 | PWB-iPSC UP | SOCS1 | 0.791758 | ENSG00000169554 | 9839 | PWB-iPSC DN | ZEB2 | -6.125403311 | ENSG00000139624 | 91012 | PWB-MSC UP | CERS5 | 0.4661019 |
| ENSG00000263956 | 200030 | PWB-iPSC UP | NBPF11 | 0.791692 | ENSG00000268902 | 102723547 | PWB-iPSC DN | CSAG2 | -6.317387547 | ENSG00000160877 | 112939 | PWB-MSC UP | NACC1 | 0.4659259 |
| ENSG00000150672 | 1740 | PWB-iPSC UP | DLG2 | 0.79061 | ENSG00000169245 | 3627 | PWB-iPSC DN | CXCL10 | -6.336455655 | ENSG00000075415 | 5250 | PWB-MSC UP | SLC25A3 | 0.4628142 |
| ENSG00000258289 | 91612 | PWB-iPSC UP | CHURC1 | 0.788386 | ENSG00000137965 | 10561 | PWB-iPSC DN | IFI44 | -6.389280982 | ENSG00000181924 | 51287 | PWB-MSC UP | COA4 | 0.4558581 |
| ENSG00000160712 | 3570 | PWB-iPSC UP | IL6R | 0.787698 | ENSG00000233098 | 339260 | PWB-iPSC DN | LOC339260 | -6.456010151 | ENSG00000079387 | 29843 | PWB-MSC UP | SENP1 | 0.4533851 |
| ENSG00000166342 | 81832 | PWB-iPSC UP | NETO1 | 0.787226 | ENSG00000119922 | 3433 | PWB-iPSC DN | IFIT2 | -6.484091449 | ENSG00000182197 | 2131 | PWB-MSC UP | EXT1 | 0.4513303 |
| ENSG00000151093 | 54995 | PWB-iPSC UP | OXSM | 0.787054 | ENSG00000141096 | 64180 | PWB-iPSC DN | DPEP3 | -6.542357726 | ENSG00000198301 | 55153 | PWB-MSC UP | SDAD1 | 0.44996 |
| ENSG00000100505 | 114088 | PWB-iPSC UP | TRIM9 | 0.786513 | ENSG00000148704 | 11023 | PWB-iPSC DN | VAX1 | -7.300401228 | ENSG00000123352 | 65244 | PWB-MSC UP | SPATS2 | 0.4497513 |
| ENSG00000143382 | 54507 | PWB-iPSC UP | ADAMTSL4 | 0.78456 | ENSG00000215559 | 391267 | PWB-iPSC DN | ANKRD20A11( | -7.508804891 | ENSG00000167553 | 84790 | PWB-MSC UP | TUBA1C | 0.4453941 |
| ENSG00000221843 | 84226 | PWB-iPSC UP | C2orf16 | 0.78356 | ENSG00000198930 | 158511 | PWB-iPSC DN | CSAG1 | -7.592560451 | ENSG00000089022 | 8550 | PWB-MSC UP | MAPKAPK5 | 0.4276815 |
| ENSG00000166535 | 144568 | PWB-iPSC UP | A2ML1 | 0.780583 | ENSG00000268916 | 389903 | PWB-iPSC DN | CSAG3 | -8.463140705 | ENSG00000168385 | 4735 | PWB-MSC DN | SEPTIN2 | -0.429361 |
| ENSG00000268658 | 400680 | PWB-iPSC UP | LINC00664 | 0.780516 | ENSG00000228314 | 54055 | PWB-iPSC DN | CYP4F29P | -8.564524129 | ENSG00000138829 | 2201 | PWB-MSC DN | FBN2 | -0.444198 |
| ENSG00000167103 | 138429 | PWB-iPSC UP | PIP5KL1 | 0.78041 | ENSG00000229807 | 7503 | PWB-iPSC DN | XIST | -8.742106023 | ENSG00000184014 | 23258 | PWB-MSC DN | DENND5A | -0.458513 |
| ENSG00000234171 | 100506054 | PWB-iPSC UP | RNASEH1-AS1 | 0.77917 | ENSG00000250420 | 729522 | PWB-iPSC DN | AACSP1 | -9.342967694 | ENSG00000128294 | 8459 | PWB-MSC DN | TPST2 | -0.458557 |
| ENSG00000214021 | 26140 | PWB-iPSC UP | TTLL3 | 0.778512 | ENSG00000198670 | 4018 | PWB-EC UP | LPA | 8.646644278 | ENSG00000197930 | 30001 | PWB-MSC DN | ERO1A | -0.473092 |
| ENSG00000164362 | 7015 | PWB-iPSC UP | TERT | 0.778307 | ENSG00000188120 | 1617 | PWB-EC UP | DAZ1 | 8.255095573 | ENSG00000144744 | 9039 | PWB-MSC DN | UBA3 | -0.475406 |
| ENSG00000170854 | 84864 | PWB-iPSC UP | RIOX2 | 0.777943 | ENSG00000205916 | 57135 | PWB-EC UP | DAZ4 | 7.66042705 | ENSG00000146373 | 154214 | PWB-MSC DN | RNF217 | -0.485233 |
| ENSG00000142347 | 4542 | PWB-iPSC UP | MYO1F | 0.776641 | ENSG00000196350 | 100287226 | PWB-EC UP | ZNF729 | 7.3773329 | ENSG00000095794 | 1390 | PWB-MSC DN | CREM | -0.486563 |
| ENSG00000251474 | 132241 | PWB-iPSC UP | RPL32P3 | 0.775948 | ENSG00000230387 | 100505664 | PWB-EC UP | LOC10050566 | 7.269999984 | ENSG00000044090 | 9820 | PWB-MSC DN | CUL7 | -0.489448 |
| ENSG00000249931 | 653125 | PWB-iPSC UP | GOLGA8K | 0.775757 | ENSG00000225560 | 105379268 | PWB-EC UP | FAM197Y8 | 7.239726767 | ENSG00000117020 | 10000 | PWB-MSC DN | AKT3 | -0.490124 |
| ENSG00000187957 | 92737 | PWB-iPSC UP | DNER | 0.77175 | ENSG00000167210 | 125336 | PWB-EC UP | LOXHD1 | 7.152950421 | ENSG00000116679 | 10625 | PWB-MSC DN | IVNS1ABP | -0.490698 |
| ENSG00000072840 | 2121 | PWB-iPSC UP | EVC | 0.770697 | ENSG00000234803 | 252946 | PWB-EC UP | FAM197Y2 | 7.059163062 | ENSG00000147548 | 54904 | PWB-MSC DN | NSD3 | -0.494542 |
| ENSG00000185917 | 54093 | PWB-iPSC UP | SETD4 | 0.770553 | ENSG00000243323 | 148713 | PWB-EC UP | PTPRVP | 6.987743152 | ENSG00000130821 | 6535 | PWB-MSC DN | SLC6A8 | -0.500994 |
| ENSG00000174652 | 10781 | PWB-iPSC UP | ZNF266 | 0.768493 | ENSG00000117090 | 6504 | PWB-EC UP | SLAMF1 | 6.930614589 | ENSG00000177463 | 7182 | PWB-MSC DN | NR2C2 | -0.504455 |
| ENSG00000246859 | 100505678 | PWB-iPSC UP | STARD4-AS1 | 0.767213 | ENSG00000215182 | 4586 | PWB-EC UP | MUC5AC | 6.851681648 | ENSG00000169379 | 200894 | PWB-MSC DN | ARL13B | -0.506308 |
| ENSG00000153904 | 23576 | PWB-iPSC UP | DDAH1 | 0.765766 | ENSG00000225516 | 100289150 | PWB-EC UP | FAM197Y5 | 6.736104348 | ENSG00000138650 | 57575 | PWB-MSC DN | PCDH10 | -0.506984 |
| ENSG00000163728 | 151613 | PWB-iPSC UP | TTC14 | 0.763784 | ENSG00000188681 | 100132288 | PWB-EC UP | TEKT4P2 | 6.681974013 | ENSG00000169519 | 196074 | PWB-MSC DN | METTL15 | -0.512708 |
| ENSG00000127957 | 5387 | PWB-iPSC UP | PMS2P3 | 0.763016 | ENSG00000100448 | 1511 | PWB-EC UP | CTSG | 6.217521281 | ENSG00000170017 | 214 | PWB-MSC DN | ALCAM | -0.512863 |
| ENSG00000100767 | 89932 | PWB-iPSC UP | PAPLN | 0.762593 | ENSG00000004468 | 952 | PWB-EC UP | CD38 | 6.176112447 | ENSG00000255529 | 81488 | PWB-MSC DN | POLR2M | -0.513347 |
| ENSG00000179046 | 205860 | PWB-iPSC UP | TRIML2 | 0.762201 | ENSG00000174450 | 283685 | PWB-EC UP | GOLGA6L2 | 5.849536424 | ENSG00000159388 | 7832 | PWB-MSC DN | BTG2 | -0.51674 |

| | | | | | | | | | | | | | | |
|---|---|---|---|---|---|---|---|---|---|---|---|---|---|---|
| ENSG00000103723 | 8120 | PWB-iPSC UP | AP3B2 | 0.760562 | ENSG00000277865 | 440243 | PWB-EC UP | GOLGA6L22 | 5.762680054 | ENSG00000120690 | 1997 | PWB-MSC DN | ELF1 | -0.519862 |
| ENSG00000167615 | 114823 | PWB-iPSC UP | LENG8 | 0.758439 | ENSG00000237850 | 645202 | PWB-EC UP | LOC645202 | 5.74307297 | ENSG00000167904 | 137695 | PWB-MSC DN | TMEM68 | -0.520981 |
| ENSG00000165923 | 79841 | PWB-iPSC UP | AGBL2 | 0.75792 | ENSG00000198788 | 4583 | PWB-EC UP | MUC2 | 5.68323975 | ENSG00000227051 | 56967 | PWB-MSC DN | C14orf132 | -0.523175 |
| ENSG00000170417 | 130827 | PWB-iPSC UP | TMEM182 | 0.757347 | ENSG00000236424 | 100289087 | PWB-EC UP | TSPY10 | 5.669026114 | ENSG00000242086 | 727956 | PWB-MSC DN | SDHAP2 | -0.524316 |
| ENSG00000126583 | 5582 | PWB-iPSC UP | PRKCG | 0.756776 | ENSG00000183625 | 1232 | PWB-EC UP | CCR3 | 5.585869917 | ENSG00000166265 | 116159 | PWB-MSC DN | CYYR1 | -0.525798 |
| ENSG00000180801 | 79642 | PWB-iPSC UP | ARSJ | 0.756616 | ENSG00000103546 | 6530 | PWB-EC UP | SLC6A2 | 5.585678611 | ENSG00000116209 | 9528 | PWB-MSC DN | TMEM59 | -0.526008 |
| ENSG00000078295 | 108 | PWB-iPSC UP | ADCY2 | 0.754655 | ENSG00000183668 | 5678 | PWB-EC UP | PSG9 | 5.538273445 | ENSG00000105866 | 6671 | PWB-MSC DN | SP4 | -0.53211 |
| ENSG00000073584 | 6605 | PWB-iPSC UP | SMARCE1 | 0.75443 | ENSG00000228927 | 728137 | PWB-EC UP | TSPY3 | 5.53236791 | ENSG00000105829 | 10282 | PWB-MSC DN | BET1 | -0.532226 |
| ENSG00000089775 | 7597 | PWB-iPSC UP | ZBTB25 | 0.753122 | ENSG00000261738 | 645355 | PWB-EC UP | MIR3976HG | 5.508526615 | ENSG00000119729 | 23433 | PWB-MSC DN | RHOQ | -0.533608 |
| ENSG00000146416 | 51390 | PWB-iPSC UP | AIG1 | 0.753006 | ENSG00000136944 | 4010 | PWB-EC UP | LMX1B | 5.477197137 | ENSG00000108861 | 1845 | PWB-MSC DN | DUSP3 | -0.535665 |
| ENSG00000062370 | 7771 | PWB-iPSC UP | ZNF112 | 0.75296 | ENSG00000229549 | 728403 | PWB-EC UP | TSPY8 | 5.471295326 | ENSG00000066422 | 27107 | PWB-MSC DN | ZBTB11 | -0.536052 |
| ENSG00000185829 | 51326 | PWB-iPSC UP | ARL17A | 0.752017 | ENSG00000006071 | 6833 | PWB-EC UP | ABCC8 | 5.426289845 | ENSG00000011028 | 9902 | PWB-MSC DN | MRC2 | -0.539491 |
| ENSG00000184619 | 124751 | PWB-iPSC UP | KRBA2 | 0.751458 | ENSG00000233803 | 728395 | PWB-EC UP | TSPY4 | 5.314063969 | ENSG00000125734 | 56927 | PWB-MSC DN | GPR108 | -0.539648 |
| ENSG00000103528 | 51760 | PWB-iPSC UP | SYT17 | 0.750747 | ENSG00000170927 | 5314 | PWB-EC UP | PKHD1 | 5.231172206 | ENSG00000169439 | 6383 | PWB-MSC DN | SDC2 | -0.543067 |
| ENSG00000143847 | 8497 | PWB-iPSC UP | PPFIA4 | 0.750429 | ENSG00000213973 | 7652 | PWB-EC UP | ZNF99 | 5.220981053 | ENSG00000088448 | 55608 | PWB-MSC DN | ANKRD10 | -0.544543 |
| ENSG00000186815 | 53373 | PWB-iPSC UP | TPCN1 | 0.74833 | ENSG00000218819 | 100129278 | PWB-EC UP | TDRD15 | 5.201924939 | ENSG00000277157 | 8360 | PWB-MSC DN | H4C4 | -0.544841 |
| ENSG00000153933 | 8526 | PWB-iPSC UP | DGKE | 0.745016 | ENSG00000225868 | 100631378 | PWB-EC UP | LOC10063137 | 5.174742866 | ENSG00000178184 | 84552 | PWB-MSC DN | PARD6G | -0.544944 |
| ENSG00000234912 | 654434 | PWB-iPSC UP | SNHG20 | 0.744058 | ENSG00000197921 | 388585 | PWB-EC UP | HES5 | 5.152655499 | ENSG00000101911 | 5634 | PWB-MSC DN | PRPS2 | -0.547699 |
| ENSG00000260034 | 100506655 | PWB-iPSC UP | LCMT1-AS2 | 0.742579 | ENSG00000089169 | 22895 | PWB-EC UP | RPH3A | 5.087922024 | ENSG00000166803 | 9768 | PWB-MSC DN | PCLAF | -0.548309 |
| ENSG00000006652 | 3475 | PWB-iPSC UP | IFRD1 | 0.742357 | ENSG00000169894 | 4584 | PWB-EC UP | MUC3A | 5.085648305 | ENSG00000176887 | 6664 | PWB-MSC DN | SOX11 | -0.549156 |
| ENSG00000268516 | 105372482 | PWB-iPSC UP | LOC105372482 | 0.742104 | ENSG00000207744 | 406903 | PWB-EC UP | MIR10B | 4.9893137 | ENSG00000168116 | 57691 | PWB-MSC DN | KIAA1586 | -0.550187 |
| ENSG00000184731 | 642273 | PWB-iPSC UP | FAM110C | 0.741231 | ENSG00000241186 | 6997 | PWB-EC UP | TDGF1 | 4.958946179 | ENSG00000187837 | 3006 | PWB-MSC DN | H1-2 | -0.551949 |
| ENSG00000079102 | 862 | PWB-iPSC UP | RUNX1T1 | 0.739963 | ENSG00000227717 | 100132202 | PWB-EC UP | LOC10013220 | 4.858537951 | ENSG00000286522 | 8358 | PWB-MSC DN | H3C2 | -0.552601 |
| ENSG00000283674 | 729732 | PWB-iPSC UP | LOC729732 | 0.738118 | ENSG00000197360 | 148198 | PWB-EC UP | ZNF98 | 4.805078901 | ENSG00000100934 | 10484 | PWB-MSC DN | SEC23A | -0.553488 |
| ENSG00000204681 | 2550 | PWB-iPSC UP | GABBR1 | 0.738074 | ENSG00000243137 | 5672 | PWB-EC UP | PSG4 | 4.796063452 | ENSG00000172667 | 64393 | PWB-MSC DN | ZMAT3 | -0.556121 |
| ENSG00000148600 | 92211 | PWB-iPSC UP | CDHR1 | 0.737965 | ENSG00000197134 | 113835 | PWB-EC UP | ZNF257 | 4.778797008 | ENSG00000176225 | 25914 | PWB-MSC DN | RTTN | -0.556334 |
| ENSG00000236088 | 100874058 | PWB-iPSC UP | COX10-AS1 | 0.736964 | ENSG00000181143 | 94025 | PWB-EC UP | MUC16 | 4.749073915 | ENSG00000148690 | 118924 | PWB-MSC DN | FRA10AC1 | -0.556485 |
| ENSG00000101901 | 79868 | PWB-iPSC UP | ALG13 | 0.736213 | ENSG00000214548 | 55384 | PWB-EC UP | MEG3 | 4.730485559 | ENSG00000133812 | 81846 | PWB-MSC DN | SBF2 | -0.557364 |
| ENSG00000228649 | 109729180 | PWB-iPSC UP | SNHG26 | 0.736134 | ENSG00000224271 | 284930 | PWB-EC UP | LOC284930 | 4.729652523 | ENSG00000146112 | 170954 | PWB-MSC DN | PPP1R18 | -0.558698 |
| ENSG00000269609 | 100505761 | PWB-iPSC UP | RPARP-AS1 | 0.735237 | ENSG00000225366 | 6998 | PWB-EC UP | TDGF1P3 | 4.706526938 | ENSG00000138032 | 5495 | PWB-MSC DN | PPM1B | -0.559083 |
| ENSG00000173041 | 340252 | PWB-iPSC UP | ZNF680 | 0.734278 | ENSG00000007908 | 6401 | PWB-EC UP | SELE | 4.697621388 | ENSG00000197223 | 10438 | PWB-MSC DN | C1D | -0.563728 |
| ENSG00000196268 | 284443 | PWB-iPSC UP | ZNF493 | 0.734099 | ENSG00000235142 | 100422737 | PWB-EC UP | LINC02532 | 4.687611863 | ENSG00000144935 | 7220 | PWB-MSC DN | TRPC1 | -0.56741 |
| ENSG00000135541 | 54806 | PWB-iPSC UP | AHI1 | 0.733193 | ENSG00000070886 | 2046 | PWB-EC UP | EPHA8 | 4.68667187 | ENSG00000113812 | 93973 | PWB-MSC DN | ACTR8 | -0.568148 |
| ENSG00000198064 | 613037 | PWB-iPSC UP | NPIPB13 | 0.732671 | ENSG00000273756 | 102723623 | PWB-EC UP | LOC10272362 | 4.625696916 | ENSG00000177917 | 151188 | PWB-MSC DN | ARL6IP6 | -0.572641 |
| ENSG00000114857 | 4820 | PWB-iPSC UP | NKTR | 0.731944 | ENSG00000263711 | 400655 | PWB-EC UP | LINC02864 | 4.616529055 | ENSG00000164291 | 153642 | PWB-MSC DN | ARSK | -0.573123 |
| ENSG00000168056 | 4054 | PWB-iPSC UP | LTBP3 | 0.731906 | ENSG00000226979 | 4049 | PWB-EC UP | LTA | 4.578847347 | ENSG00000156831 | 286053 | PWB-MSC DN | NSMCE2 | -0.573303 |
| ENSG00000188878 | 85302 | PWB-iPSC UP | FBF1 | 0.731219 | ENSG00000227403 | 100996579 | PWB-EC UP | LINC01806 | 4.547428281 | ENSG00000181904 | 134553 | PWB-MSC DN | C5orf24 | -0.573482 |
| ENSG00000267244 | 100288123 | PWB-iPSC UP | LOC100288123 | 0.731154 | ENSG00000053108 | 23105 | PWB-EC UP | FSTL4 | 4.53007392 | ENSG00000135926 | 64114 | PWB-MSC DN | TMBIM1 | -0.577427 |
| ENSG00000196756 | 388796 | PWB-iPSC UP | SNHG17 | 0.730952 | ENSG00000258867 | 283587 | PWB-EC UP | LINC01146 | 4.495016539 | ENSG00000033867 | 9497 | PWB-MSC DN | SLC4A7 | -0.578439 |
| ENSG00000180747 | 100271836 | PWB-iPSC UP | SMG1P3 | 0.730764 | ENSG00000276462 | 440896 | PWB-EC UP | LOC440896 | 4.388288454 | ENSG00000124570 | 5269 | PWB-MSC DN | SERPINB6 | -0.579 |
| ENSG00000173320 | 56977 | PWB-iPSC UP | STOX2 | 0.729394 | ENSG00000104938 | 10332 | PWB-EC UP | CLEC4M | 4.37641791 | ENSG00000124575 | 3007 | PWB-MSC DN | H1-3 | -0.581098 |
| ENSG00000166801 | 63901 | PWB-iPSC UP | FAM111A | 0.724178 | ENSG00000269067 | 388523 | PWB-EC UP | ZNF728 | 4.298092276 | ENSG00000127955 | 2770 | PWB-MSC DN | GNAI1 | -0.584302 |
| ENSG00000112972 | 3157 | PWB-iPSC UP | HMGCS1 | 0.722957 | ENSG00000273976 | 283767 | PWB-EC UP | GOLGA6L1 | 4.243334714 | ENSG00000138738 | 11107 | PWB-MSC DN | PRDM5 | -0.585291 |
| ENSG00000164114 | 79884 | PWB-iPSC UP | MAP9 | 0.722716 | ENSG00000224141 | 101927797 | PWB-EC UP | MIR548XHG | 4.2300046 | ENSG00000181690 | 5324 | PWB-MSC DN | PLAG1 | -0.587315 |
| ENSG00000185324 | 8558 | PWB-iPSC UP | CDK10 | 0.721849 | ENSG00000168757 | 64591 | PWB-EC UP | TSPY2 | 4.187336982 | ENSG00000117385 | 64175 | PWB-MSC DN | P3H1 | -0.587692 |
| ENSG00000106688 | 6505 | PWB-iPSC UP | SLC1A1 | 0.720815 | ENSG00000279516 | 26080 | PWB-EC UP | FAM230C | 4.15372288 | ENSG00000156162 | 286148 | PWB-MSC DN | DPY19L4 | -0.590089 |
| ENSG00000015479 | 9782 | PWB-iPSC UP | MATR3 | 0.71482 | ENSG00000261594 | 100507050 | PWB-EC UP | TPBGL | 4.136249058 | ENSG00000184867 | 9823 | PWB-MSC DN | ARMCX2 | -0.591456 |
| ENSG00000187678 | 81848 | PWB-iPSC UP | SPRY4 | 0.714607 | ENSG00000277322 | 727832 | PWB-EC UP | GOLGA6L6 | 4.126615465 | ENSG00000119900 | 79627 | PWB-MSC DN | OGFRL1 | -0.593472 |
| ENSG00000064012 | 841 | PWB-iPSC UP | CASP8 | 0.713064 | ENSG00000196109 | 163223 | PWB-EC UP | ZNF676 | 4.099648259 | ENSG00000078114 | 10529 | PWB-MSC DN | NEBL | -0.595307 |
| ENSG00000101546 | 79863 | PWB-iPSC UP | RBFA | 0.711506 | ENSG00000105499 | 8605 | PWB-EC UP | PLA2G4C | 4.052775913 | ENSG00000155275 | 152992 | PWB-MSC DN | TRMT44 | -0.595819 |
| ENSG00000183793 | 100288332 | PWB-iPSC UP | NPIPA5 | 0.710587 | ENSG00000183317 | 284656 | PWB-EC UP | EPHA10 | 4.040399378 | ENSG00000107679 | 59338 | PWB-MSC DN | PLEKHA1 | -0.59754 |
| ENSG00000152601 | 4154 | PWB-iPSC UP | MBNL1 | 0.709482 | ENSG00000149654 | 64405 | PWB-EC UP | CDH22 | 4.027754643 | ENSG00000137944 | 56267 | PWB-MSC DN | KYAT3 | -0.598281 |
| ENSG00000131127 | 7700 | PWB-iPSC UP | ZNF141 | 0.7092 | ENSG00000163739 | 2919 | PWB-EC UP | CXCL1 | 3.988278602 | ENSG00000135862 | 3915 | PWB-MSC DN | LAMC1 | -0.599778 |
| ENSG00000172461 | 10690 | PWB-iPSC UP | FUT9 | 0.707895 | ENSG00000180287 | 200150 | PWB-EC UP | PLD5 | 3.979050787 | ENSG00000113638 | 23548 | PWB-MSC DN | TTC33 | -0.60103 |
| ENSG00000282458 | 375690 | PWB-iPSC UP | WASH5P | 0.707762 | ENSG00000258279 | 283404 | PWB-EC UP | LINC00592 | 3.947871841 | ENSG00000058063 | 23200 | PWB-MSC DN | ATP11B | -0.60343 |
| ENSG00000143858 | 127833 | PWB-iPSC UP | SYT2 | 0.706675 | ENSG00000142910 | 64129 | PWB-EC UP | TINAGL1 | 3.945199034 | ENSG00000272886 | 55802 | PWB-MSC DN | DCP1A | -0.608307 |
| ENSG00000182389 | 785 | PWB-iPSC UP | CACNB4 | 0.706387 | ENSG00000229859 | 643834 | PWB-EC UP | PGA3 | 3.924117317 | ENSG00000053524 | 23101 | PWB-MSC DN | MCF2L2 | -0.608525 |
| ENSG00000235590 | 149775 | PWB-iPSC UP | GNAS-AS1 | 0.705674 | ENSG00000038945 | 4481 | PWB-EC UP | MSR1 | 3.918610022 | ENSG00000010270 | 83930 | PWB-MSC DN | STARD3NL | -0.613144 |
| ENSG00000151914 | 667 | PWB-iPSC UP | DST | 0.703685 | ENSG00000115507 | 5013 | PWB-EC UP | OTX1 | 3.914192672 | ENSG00000090006 | 8425 | PWB-MSC DN | LTBP4 | -0.614405 |
| ENSG00000163138 | 133015 | PWB-iPSC UP | PACRGL | 0.702816 | ENSG00000115008 | 3552 | PWB-EC UP | IL1A | 3.876823867 | ENSG00000128881 | 146057 | PWB-MSC DN | TTBK2 | -0.615436 |
| ENSG00000123552 | 85015 | PWB-iPSC UP | USP45 | 0.701523 | ENSG00000181984 | 729786 | PWB-EC UP | GOLGA8CP | 3.84536366 | ENSG00000153707 | 5789 | PWB-MSC DN | PTPRD | -0.619872 |
| ENSG00000226696 | 104355426 | PWB-iPSC UP | LENG8-AS1 | 0.701281 | ENSG00000286691 | 107986631 | PWB-EC UP | LOC10798663 | 3.784479008 | ENSG00000164619 | 168667 | PWB-MSC DN | BMPER | -0.623838 |
| ENSG00000235823 | 90271 | PWB-iPSC UP | OLMALINC | 0.700435 | ENSG00000198028 | 147741 | PWB-EC UP | ZNF560 | 3.74951482 | ENSG00000122126 | 4952 | PWB-MSC DN | OCRL | -0.624602 |
| ENSG00000224712 | 642778 | PWB-iPSC UP | NPIPA3 | 0.696699 | ENSG00000017427 | 3479 | PWB-EC UP | IGF1 | 3.733364359 | ENSG00000105655 | 51477 | PWB-MSC DN | ISYNA1 | -0.626433 |
| ENSG00000076043 | 25996 | PWB-iPSC UP | REXO2 | 0.696166 | ENSG00000277526 | 101927815 | PWB-EC UP | LOC10192781 | 3.703206789 | ENSG00000101752 | 57534 | PWB-MSC DN | MIB1 | -0.627118 |
| ENSG00000147364 | 26260 | PWB-iPSC UP | FBXO25 | 0.695718 | ENSG00000184530 | 352999 | PWB-EC UP | C6orf58 | 3.701914482 | ENSG00000102409 | 56271 | PWB-MSC DN | BEX4 | -0.628147 |
| ENSG00000259803 | 146429 | PWB-iPSC UP | SLC22A31 | 0.695558 | ENSG00000187037 | 353345 | PWB-EC UP | GPR141 | 3.693950736 | ENSG00000274618 | 8361 | PWB-MSC DN | H4C6 | -0.632819 |
| ENSG00000149150 | 8501 | PWB-iPSC UP | SLC43A1 | 0.693635 | ENSG00000229372 | 100873952 | PWB-EC UP | SZT2-AS1 | 3.66962589 | ENSG00000050165 | 27122 | PWB-MSC DN | DKK3 | -0.633173 |
| ENSG00000237296 | 641298 | PWB-iPSC UP | SMG1P1 | 0.691164 | ENSG00000196218 | 6261 | PWB-EC UP | RYR1 | 3.663908681 | ENSG00000182093 | 7485 | PWB-MSC DN | GET1 | -0.634989 |
| ENSG00000196141 | 26010 | PWB-iPSC UP | SPATS2L | 0.690759 | ENSG00000166183 | 374569 | PWB-EC UP | ASPG | 3.661328633 | ENSG00000182287 | 8905 | PWB-MSC DN | AP1S2 | -0.637294 |
| ENSG00000265479 | 441263 | PWB-iPSC UP | DTX2P1-UPK3BP1- | 0.688994 | ENSG00000146955 | 401409 | PWB-EC UP | RAB19 | 3.6496924 | ENSG00000156103 | 4325 | PWB-MSC DN | MMP16 | -0.641456 |
| ENSG00000254852 | 642799 | PWB-iPSC UP | NPIPA2 | 0.688772 | ENSG00000091879 | 285 | PWB-EC UP | ANGPT2 | 3.641972794 | ENSG00000103160 | 83693 | PWB-MSC DN | HSDL1 | -0.641524 |
| ENSG00000197444 | 55753 | PWB-iPSC UP | OGDHL | 0.686403 | ENSG00000183629 | 283768 | PWB-EC UP | GOLGA8G | 3.628631844 | ENSG00000134013 | 4017 | PWB-MSC DN | LOXL2 | -0.642052 |
| ENSG00000153898 | 255231 | PWB-iPSC UP | MCOLN2 | 0.68412 | ENSG00000141668 | 147381 | PWB-EC UP | CBLN2 | 3.627587795 | ENSG00000102804 | 8848 | PWB-MSC DN | TSC22D1 | -0.644692 |
| ENSG00000171757 | 151827 | PWB-iPSC UP | LRRC34 | 0.68213 | ENSG00000008517 | 9235 | PWB-EC UP | IL32 | 3.609092985 | ENSG00000109787 | 51274 | PWB-MSC DN | KLF3 | -0.644785 |
| ENSG00000240694 | 10687 | PWB-iPSC UP | PNMA2 | 0.681754 | ENSG00000163673 | 85443 | PWB-EC UP | DCLK3 | 3.598753994 | ENSG00000235823 | 90271 | PWB-MSC DN | OLMALINC | -0.645788 |
| ENSG00000047188 | 64848 | PWB-iPSC UP | YTHDC2 | 0.681229 | ENSG00000169429 | 3576 | PWB-EC UP | CXCL8 | 3.59313912 | ENSG00000112186 | 10486 | PWB-MSC DN | CAP2 | -0.64615 |
| ENSG00000122786 | 800 | PWB-iPSC UP | CALD1 | 0.680458 | ENSG00000233208 | 400238 | PWB-EC UP | LINC00642 | 3.572722687 | ENSG00000135338 | 167691 | PWB-MSC DN | LCA5 | -0.648247 |
| ENSG00000242125 | 8420 | PWB-iPSC UP | SNHG3 | 0.679685 | ENSG00000126251 | 2866 | PWB-EC UP | GPR42 | 3.524256075 | ENSG00000179454 | 54813 | PWB-MSC DN | KLHL28 | -0.651198 |
| ENSG00000134802 | 29015 | PWB-iPSC UP | SLC43A3 | 0.675976 | ENSG00000163218 | 57115 | PWB-EC UP | PGLYRP4 | 3.517249352 | ENSG00000155330 | 388272 | PWB-MSC DN | C16orf87 | -0.652268 |
| ENSG00000198286 | 84433 | PWB-iPSC UP | CARD11 | 0.674397 | ENSG00000057149 | 6317 | PWB-EC UP | SERPINB3 | 3.515835149 | ENSG00000162616 | 11080 | PWB-MSC DN | DNAJB4 | -0.653529 |
| ENSG00000183718 | 84851 | PWB-iPSC UP | TRIM52 | 0.670228 | ENSG00000163131 | 1520 | PWB-EC UP | CTSS | 3.498491984 | ENSG00000138814 | 5530 | PWB-MSC DN | PPP3CA | -0.653999 |
| ENSG00000153558 | 25827 | PWB-iPSC UP | FBXL2 | 0.66988 | ENSG00000107562 | 6387 | PWB-EC UP | CXCL12 | 3.453898977 | ENSG00000186104 | 120227 | PWB-MSC DN | CYP2R1 | -0.660071 |
| ENSG00000166669 | 80063 | PWB-iPSC UP | ATF7IP2 | 0.665146 | ENSG00000077327 | 9576 | PWB-EC UP | SPAG6 | 3.445244248 | ENSG00000106025 | 23554 | PWB-MSC DN | TSPAN12 | -0.660128 |
| ENSG00000196417 | 91661 | PWB-iPSC UP | ZNF765 | 0.66405 | ENSG00000232265 | 100505824 | PWB-EC UP | LINC02805 | 3.434967659 | ENSG00000269713 | 400818 | PWB-MSC DN | NBPF9 | -0.663782 |
| ENSG00000266338 | 284565 | PWB-iPSC UP | NBPF15 | 0.663082 | ENSG00000110848 | 969 | PWB-EC UP | CD69 | 3.414657668 | ENSG00000164292 | 22836 | PWB-MSC DN | RHOBTB3 | -0.673354 |
| ENSG00000139974 | 145389 | PWB-iPSC UP | SLC38A6 | 0.662061 | ENSG00000260220 | 399693 | PWB-EC UP | CCDC187 | 3.388426162 | ENSG00000132535 | 1742 | PWB-MSC DN | DLG4 | -0.675013 |
| ENSG00000103995 | 22995 | PWB-iPSC UP | CEP152 | 0.659935 | ENSG00000229425 | 101927745 | PWB-EC UP | LOC10192774 | 3.38052353 | ENSG00000100196 | 11015 | PWB-MSC DN | KDELR3 | -0.67547 |
| ENSG00000247708 | 100507266 | PWB-iPSC UP | STX18-AS1 | 0.65949 | ENSG00000229140 | 137196 | PWB-EC UP | CCDC26 | 3.373969508 | ENSG00000149256 | 26011 | PWB-MSC DN | TENM4 | -0.675496 |
| ENSG00000164989 | 203238 | PWB-iPSC UP | CCDC171 | 0.657495 | ENSG00000130234 | 59272 | PWB-EC UP | ACE2 | 3.353794417 | ENSG00000188321 | 84527 | PWB-MSC DN | ZNF559 | -0.677798 |
| ENSG00000169174 | 255738 | PWB-iPSC UP | PCSK9 | 0.657014 | ENSG00000138083 | 6496 | PWB-EC UP | SIX3 | 3.314537667 | ENSG00000168300 | 115294 | PWB-MSC DN | PCMTD1 | -0.687785 |
| ENSG00000138798 | 1950 | PWB-iPSC UP | EGF | 0.656362 | ENSG00000204161 | 170371 | PWB-EC UP | TMEM273 | 3.314122403 | ENSG00000197712 | 92689 | PWB-MSC DN | FAM114A1 | -0.688845 |
| ENSG00000279170 | 100130890 | PWB-iPSC UP | TSTD3 | 0.654972 | ENSG00000270547 | 401492 | PWB-EC UP | LINC01235 | 3.291354216 | ENSG00000164331 | 57763 | PWB-MSC DN | ANKRA2 | -0.701417 |
| ENSG00000198910 | 3897 | PWB-iPSC UP | L1CAM | 0.654904 | ENSG00000152467 | 284312 | PWB-EC UP | ZSCAN1 | 3.276690733 | ENSG00000164442 | 10370 | PWB-MSC DN | CITED2 | -0.701954 |
| ENSG00000133124 | 8471 | PWB-iPSC UP | IRS4 | 0.654389 | ENSG00000175445 | 4023 | PWB-EC UP | LPL | 3.267663549 | ENSG00000147202 | 1730 | PWB-MSC DN | DIAPH2 | -0.70645 |
| ENSG00000135127 | 92558 | PWB-iPSC UP | BICDL1 | 0.652937 | ENSG00000232810 | 7124 | PWB-EC UP | TNF | 3.265402041 | ENSG00000105856 | 26959 | PWB-MSC DN | HBP1 | -0.714578 |
| ENSG00000117533 | 8674 | PWB-iPSC UP | VAMP4 | 0.651295 | ENSG00000154478 | 2849 | PWB-EC UP | GPR26 | 3.262710988 | ENSG00000143924 | 27436 | PWB-MSC DN | EML4 | -0.715199 |
| ENSG00000204764 | 64901 | PWB-iPSC UP | RANBP17 | 0.651252 | ENSG00000215498 | 642633 | PWB-EC UP | FAM230B | 3.249005792 | ENSG00000196368 | 55190 | PWB-MSC DN | NUDT11 | -0.715218 |
| ENSG00000081059 | 6932 | PWB-iPSC UP | TCF7 | 0.651215 | ENSG00000090339 | 3383 | PWB-EC UP | ICAM1 | 3.243842667 | ENSG00000130522 | 3727 | PWB-MSC DN | JUND | -0.716064 |
| ENSG00000113721 | 5159 | PWB-iPSC UP | PDGFRB | 0.65065 | ENSG00000144644 | 339896 | PWB-EC UP | GADL1 | 3.23915575 | ENSG00000147394 | 7739 | PWB-MSC DN | ZNF185 | -0.720378 |
| ENSG00000148120 | 84909 | PWB-iPSC UP | AOPEP | 0.649982 | ENSG00000228340 | 284757 | PWB-EC UP | MIR646HG | 3.224720393 | ENSG00000148677 | 27063 | PWB-MSC DN | ANKRD1 | -0.720595 |
| ENSG00000196218 | 6261 | PWB-iPSC UP | RYR1 | 0.649904 | ENSG00000183273 | 160777 | PWB-EC UP | CCDC60 | 3.207805335 | ENSG00000185585 | 169611 | PWB-MSC DN | OLFML2A | -0.724375 |
| ENSG00000215252 | 440270 | PWB-iPSC UP | GOLGA8B | 0.648171 | ENSG00000120217 | 29126 | PWB-EC UP | CD274 | 3.184565294 | ENSG00000049323 | 4052 | PWB-MSC DN | LTBP1 | -0.725134 |
| ENSG00000145012 | 4026 | PWB-iPSC UP | LPP | 0.647688 | ENSG00000141497 | 84225 | PWB-EC UP | ZMYND15 | 3.1470131 | ENSG00000125354 | 23157 | PWB-MSC DN | SEPTIN6 | -0.725232 |
| ENSG00000226887 | 100288413 | PWB-iPSC UP | ERVMER34-1 | 0.645864 | ENSG00000108001 | 253738 | PWB-EC UP | EBF3 | 3.145789822 | ENSG00000164318 | 133584 | PWB-MSC DN | EGFLAM | -0.727742 |
| ENSG00000185946 | 55599 | PWB-iPSC UP | RNPC3 | 0.645762 | ENSG00000160321 | 7757 | PWB-EC UP | ZNF208 | 3.139716559 | ENSG00000071575 | 28951 | PWB-MSC DN | TRIB2 | -0.729022 |
| ENSG00000146350 | 221322 | PWB-iPSC UP | TBC1D32 | 0.644527 | ENSG00000234323 | 100996590 | PWB-EC UP | LINC01505 | 3.107162652 | ENSG00000145736 | 2966 | PWB-MSC DN | GTF2H2 | -0.730273 |

| | | | | | | | | | | | | | | | | |
|---|---|---|---|---|---|---|---|---|---|---|---|---|---|---|---|---|
| ENSG00000008710 | 5310 | PWB-iPSC UP | PKD1 | 0.643779 | | ENSG00000175676 | 100132979 | PWB-EC UP | GOLGA8DP | 3.097180629 | | ENSG00000075043 | 3785 | PWB-MSC DN | KCNQ2 | -0.737954 |
| ENSG00000184307 | 254887 | PWB-iPSC UP | ZDHHC23 | 0.643142 | | ENSG00000183378 | 341277 | PWB-EC UP | OVCH2 | 3.09347304 | | ENSG00000176641 | 220441 | PWB-MSC DN | RNF152 | -0.740844 |
| ENSG00000072274 | 7037 | PWB-iPSC UP | TFRC | 0.641004 | | ENSG00000166535 | 144568 | PWB-EC UP | A2ML1 | 3.077374448 | | ENSG00000150093 | 3688 | PWB-MSC DN | ITGB1 | -0.741682 |
| ENSG00000139629 | 11226 | PWB-iPSC UP | GALNT6 | 0.640719 | | ENSG00000206142 | 100996335 | PWB-EC UP | FAM230H | 3.032775927 | | ENSG00000138448 | 3685 | PWB-MSC DN | ITGAV | -0.742606 |
| ENSG00000197008 | 7697 | PWB-iPSC UP | ZNF138 | 0.640393 | | ENSG00000134201 | 2949 | PWB-EC UP | GSTM5 | 3.022262064 | | ENSG00000185201 | 10581 | PWB-MSC DN | IFITM2 | -0.746097 |
| ENSG00000161960 | 1973 | PWB-iPSC UP | EIF4A1 | 0.638945 | | ENSG00000102109 | 27344 | PWB-EC UP | PCSK1N | 3.012195183 | | ENSG00000095321 | 1384 | PWB-MSC DN | CRAT | -0.748059 |
| ENSG00000188234 | 119016 | PWB-iPSC UP | AGAP4 | 0.638511 | | ENSG00000153684 | 100132565 | PWB-EC UP | GOLGA8F | 3.002841346 | | ENSG00000112893 | 4124 | PWB-MSC DN | MAN2A1 | -0.749634 |
| ENSG00000169203 | 440353 | PWB-iPSC UP | NPIPB12 | 0.637629 | | ENSG00000156395 | 22986 | PWB-EC UP | SORCS3 | 2.940303305 | | ENSG00000070087 | 5217 | PWB-MSC DN | PFN2 | -0.752889 |
| ENSG00000023909 | 2730 | PWB-iPSC UP | GCLM | 0.63676 | | ENSG00000215845 | 100131187 | PWB-EC UP | TSTD1 | 2.928512734 | | ENSG00000021355 | 1992 | PWB-MSC DN | SERPINB1 | -0.753609 |
| ENSG00000140835 | 10164 | PWB-iPSC UP | CHST4 | 0.636635 | | ENSG00000255545 | 283177 | PWB-EC UP | LOC283177 | 2.906294016 | | ENSG00000181458 | 55076 | PWB-MSC DN | TMEM45A | -0.755259 |
| ENSG00000149483 | 51524 | PWB-iPSC UP | TMEM138 | 0.635449 | | ENSG00000172716 | 91607 | PWB-EC UP | SLFN11 | 2.899861203 | | ENSG00000163577 | 56648 | PWB-MSC DN | EIF5A2 | -0.755911 |
| ENSG00000163738 | 441024 | PWB-iPSC UP | MTHFD2L | 0.629995 | | ENSG00000205869 | 387264 | PWB-EC UP | KRTAP5-1 | 2.897031352 | | ENSG00000177380 | 8541 | PWB-MSC DN | PPFIA3 | -0.760414 |
| ENSG00000121904 | 114784 | PWB-iPSC UP | CSMD2 | 0.629728 | | ENSG00000234722 | 103724390 | PWB-EC UP | LINC01287 | 2.889150861 | | ENSG00000108515 | 2027 | PWB-MSC DN | ENO3 | -0.764576 |
| ENSG00000175455 | 64770 | PWB-iPSC UP | CCDC14 | 0.628961 | | ENSG00000225937 | 50652 | PWB-EC UP | PCA3 | 2.873660548 | | ENSG00000151725 | 79682 | PWB-MSC DN | CENPU | -0.768901 |
| ENSG00000188517 | 84570 | PWB-iPSC UP | COL25A1 | 0.628684 | | ENSG00000198515 | 1259 | PWB-EC UP | CNGA1 | 2.868645997 | | ENSG00000135740 | 6553 | PWB-MSC DN | SLC9A5 | -0.770663 |
| ENSG00000151612 | 152485 | PWB-iPSC UP | ZNF827 | 0.624168 | | ENSG00000226686 | 101927667 | PWB-EC UP | LINC01535 | 2.865886679 | | ENSG00000172318 | 8708 | PWB-MSC DN | B3GALT1 | -0.773362 |
| ENSG00000204271 | 169981 | PWB-iPSC UP | SPIN3 | 0.623786 | | ENSG00000111981 | 80329 | PWB-EC UP | ULBP1 | 2.86266574 | | ENSG00000136141 | 23143 | PWB-MSC DN | LRCH1 | -0.779455 |
| ENSG00000137841 | 5330 | PWB-iPSC UP | PLCB2 | 0.620166 | | ENSG00000134070 | 3656 | PWB-EC UP | IRAK2 | 2.857982949 | | ENSG00000151151 | 253430 | PWB-MSC DN | IPMK | -0.785826 |
| ENSG00000074370 | 489 | PWB-iPSC UP | ATP2A3 | 0.619448 | | ENSG00000226620 | 144920 | PWB-EC UP | LINC00343 | 2.805230542 | | ENSG00000106617 | 51422 | PWB-MSC DN | PRKAG2 | -0.786979 |
| ENSG00000105136 | 79744 | PWB-iPSC UP | ZNF419 | 0.618781 | | ENSG00000185155 | 83881 | PWB-EC UP | MIXL1 | 2.794969779 | | ENSG00000106853 | 22949 | PWB-MSC DN | PTGR1 | -0.787398 |
| ENSG00000119688 | 5826 | PWB-iPSC UP | ABCD4 | 0.618066 | | ENSG00000253910 | 56103 | PWB-EC UP | PCDHGB2 | 2.794012627 | | ENSG00000090530 | 55214 | PWB-MSC DN | P3H2 | -0.787507 |
| ENSG00000216775 | 730101 | PWB-iPSC UP | LOC730101 | 0.617517 | | ENSG00000116032 | 116444 | PWB-EC UP | GRIN3B | 2.790493398 | | ENSG00000069188 | 54549 | PWB-MSC DN | SDK2 | -0.788506 |
| ENSG00000182986 | 162967 | PWB-iPSC UP | ZNF320 | 0.616277 | | ENSG00000238121 | 100188949 | PWB-EC UP | LINC00426 | 2.790119236 | | ENSG00000153214 | 84910 | PWB-MSC DN | TMEM87B | -0.790046 |
| ENSG00000183340 | 8690 | PWB-iPSC UP | JRKL | 0.614726 | | ENSG00000140090 | 123041 | PWB-EC UP | SLC24A4 | 2.765106868 | | ENSG00000270231 | 728841 | PWB-MSC DN | NBPF8 | -0.794746 |
| ENSG00000175893 | 340481 | PWB-iPSC UP | ZDHHC21 | 0.614296 | | ENSG00000173083 | 10855 | PWB-EC UP | HPSE | 2.760364371 | | ENSG00000183091 | 4703 | PWB-MSC DN | NEB | -0.799288 |
| ENSG00000008311 | 10157 | PWB-iPSC UP | AASS | 0.613703 | | ENSG00000110324 | 3587 | PWB-EC UP | IL10RA | 2.750953572 | | ENSG00000206503 | 3105 | PWB-MSC DN | HLA-A | -0.799623 |
| ENSG00000186166 | 338657 | PWB-iPSC UP | CCDC84 | 0.611264 | | ENSG00000184515 | 340542 | PWB-EC UP | BEX5 | 2.733729169 | | ENSG00000119640 | 97 | PWB-MSC DN | ACYP1 | -0.803891 |
| ENSG00000185485 | 255812 | PWB-iPSC UP | SDHAP1 | 0.610917 | | ENSG00000278873 | 100133319 | PWB-EC UP | PRO1804 | 2.730581294 | | ENSG00000112406 | 51696 | PWB-MSC DN | HECA | -0.805144 |
| ENSG00000155850 | 1836 | PWB-iPSC UP | SLC26A2 | 0.608534 | | ENSG00000155926 | 6503 | PWB-EC UP | SLA | 2.696127724 | | ENSG00000107537 | 5264 | PWB-MSC DN | PHYH | -0.809588 |
| ENSG00000140534 | 90381 | PWB-iPSC UP | TICRR | 0.60847 | | ENSG00000103534 | 79838 | PWB-EC UP | TMC5 | 2.68754918 | | ENSG00000163683 | 201895 | PWB-MSC DN | SMIM14 | -0.810657 |
| ENSG00000130844 | 55422 | PWB-iPSC UP | ZNF331 | 0.607914 | | ENSG00000124343 | 7499 | PWB-EC UP | XG | 2.685351742 | | ENSG00000177932 | 30832 | PWB-MSC DN | ZNF354C | -0.811848 |
| ENSG00000167766 | 55769 | PWB-iPSC UP | ZNF83 | 0.607493 | | ENSG00000249574 | 442497 | PWB-EC UP | LOC442497 | 2.674382544 | | ENSG00000053438 | 4826 | PWB-MSC DN | NNAT | -0.812228 |
| ENSG00000236144 | 100506469 | PWB-iPSC UP | TMEM147-AS1 | 0.604444 | | ENSG00000183090 | 166752 | PWB-EC UP | FREM3 | 2.668536042 | | ENSG00000122707 | 8434 | PWB-MSC DN | RECK | -0.814296 |
| ENSG00000165684 | 6621 | PWB-iPSC UP | SNAPC4 | 0.602916 | | ENSG00000023445 | 330 | PWB-EC UP | BIRC3 | 2.667806672 | | ENSG00000155657 | 7273 | PWB-MSC DN | TTN | -0.816102 |
| ENSG00000154920 | 146956 | PWB-iPSC UP | EME1 | 0.601617 | | ENSG00000154262 | 23460 | PWB-EC UP | ABCA6 | 2.667431863 | | ENSG00000119185 | 9270 | PWB-MSC DN | ITGB1BP1 | -0.818174 |
| ENSG00000159314 | 201176 | PWB-iPSC UP | ARHGAP27 | 0.601181 | | ENSG00000205277 | 10071 | PWB-EC UP | MUC12 | 2.662381277 | | ENSG00000102780 | 160851 | PWB-MSC DN | DGKH | -0.820552 |
| ENSG00000072041 | 55117 | PWB-iPSC UP | SLC6A15 | 0.600755 | | ENSG00000142609 | 85452 | PWB-EC UP | CFAP74 | 2.653216724 | | ENSG00000116717 | 1647 | PWB-MSC DN | GADD45A | -0.82311 |
| ENSG00000107281 | 56654 | PWB-iPSC UP | NPDC1 | 0.600753 | | ENSG00000125538 | 3553 | PWB-EC UP | IL1B | 2.64729131 | | ENSG00000169851 | 5099 | PWB-MSC DN | PCDH7 | -0.823674 |
| ENSG00000164211 | 134429 | PWB-iPSC UP | STARD4 | 0.600692 | | ENSG00000177294 | 162517 | PWB-EC UP | FBXO39 | 2.597734405 | | ENSG00000058091 | 5218 | PWB-MSC DN | CDK14 | -0.827038 |
| ENSG00000187741 | 2175 | PWB-iPSC UP | FANCA | 0.600662 | | ENSG00000136560 | 10010 | PWB-EC UP | TANK | 2.584383364 | | ENSG00000161298 | 84911 | PWB-MSC DN | ZNF382 | -0.827984 |
| ENSG00000268119 | 105372321 | PWB-iPSC UP | LOC105372321 | 0.598463 | | ENSG00000152078 | 148534 | PWB-EC UP | TLCD4 | 2.576767872 | | ENSG00000113448 | 5144 | PWB-MSC DN | PDE4D | -0.828622 |
| ENSG00000196313 | 9883 | PWB-iPSC UP | POM121 | 0.597892 | | ENSG00000258405 | 147660 | PWB-EC UP | ZNF578 | 2.575258624 | | ENSG00000121236 | 117854 | PWB-MSC DN | TRIM6 | -0.839414 |
| ENSG00000247746 | 158880 | PWB-iPSC UP | USP51 | 0.597724 | | ENSG00000167195 | 653641 | PWB-EC UP | GOLGA6C | 2.571511023 | | ENSG00000167601 | 558 | PWB-MSC DN | AXL | -0.844803 |
| ENSG00000214331 | 283922 | PWB-iPSC UP | LOC283922 | 0.59701 | | ENSG00000119630 | 5228 | PWB-EC UP | PGF | 2.561439237 | | ENSG00000160233 | 81543 | PWB-MSC DN | LRRC3 | -0.851264 |
| ENSG00000067177 | 5255 | PWB-iPSC UP | PHKA1 | 0.596997 | | ENSG00000076706 | 4162 | PWB-EC UP | MCAM | 2.559220322 | | ENSG00000151388 | 81792 | PWB-MSC DN | ADAMTS12 | -0.852092 |
| ENSG00000121413 | 65982 | PWB-iPSC UP | ZSCAN18 | 0.596734 | | ENSG00000081853 | 56113 | PWB-EC UP | PCDHGA2 | 2.506414869 | | ENSG00000259330 | 100505573 | PWB-MSC DN | INAFM2 | -0.853686 |
| ENSG00000263753 | 339290 | PWB-iPSC UP | LINC00667 | 0.593814 | | ENSG00000181634 | 9966 | PWB-EC UP | TNFSF15 | 2.504889162 | | ENSG00000166341 | 8642 | PWB-MSC DN | DCHS1 | -0.85413 |
| ENSG00000157107 | 115548 | PWB-iPSC UP | FCHO2 | 0.593348 | | ENSG00000159289 | 342096 | PWB-EC UP | GOLGA6A | 2.503093979 | | ENSG00000099337 | 9424 | PWB-MSC DN | KCNK6 | -0.854585 |
| ENSG00000167380 | 7769 | PWB-iPSC UP | ZNF226 | 0.592077 | | ENSG00000146151 | 54511 | PWB-EC UP | HMGCLL1 | 2.491157231 | | ENSG00000213190 | 10962 | PWB-MSC DN | MLLT11 | -0.856076 |
| ENSG00000148690 | 118924 | PWB-iPSC UP | FRA10AC1 | 0.591878 | | ENSG00000135604 | 8676 | PWB-EC UP | STX11 | 2.491126512 | | ENSG00000198795 | 25925 | PWB-MSC DN | ZNF521 | -0.856302 |
| ENSG00000198185 | 55713 | PWB-iPSC UP | ZNF334 | 0.588831 | | ENSG00000227733 | 101927468 | PWB-EC UP | LOC10192746 | 2.486516114 | | ENSG00000197568 | 11147 | PWB-MSC DN | HHLA3 | -0.860252 |
| ENSG00000150995 | 3708 | PWB-iPSC UP | ITPR1 | 0.585531 | | ENSG00000137491 | 11309 | PWB-EC UP | SLCO2B1 | 2.483318639 | | ENSG00000156976 | 1974 | PWB-MSC DN | EIF4A2 | -0.860572 |
| ENSG00000106976 | 1759 | PWB-iPSC UP | DNM1 | 0.58524 | | ENSG00000170074 | 285596 | PWB-EC UP | FAM153A | 2.4787213 | | ENSG00000148814 | 80313 | PWB-MSC DN | LRRC27 | -0.862613 |
| ENSG00000116133 | 1718 | PWB-iPSC UP | DHCR24 | 0.58497 | | ENSG00000050030 | 340533 | PWB-EC UP | NEXMIF | 2.475402053 | | ENSG00000205542 | 7114 | PWB-MSC DN | TMSB4X | -0.863629 |
| ENSG00000163281 | 132789 | PWB-iPSC UP | GNPDA2 | 0.580489 | | ENSG00000244578 | 103344930 | PWB-EC UP | LINC01391 | 2.463291731 | | ENSG00000103723 | 8120 | PWB-MSC DN | AP3B2 | -0.869841 |
| ENSG00000198720 | 124930 | PWB-iPSC UP | ANKRD13B | 0.580487 | | ENSG00000109743 | 683 | PWB-EC UP | BST1 | 2.454205927 | | ENSG00000162623 | 127253 | PWB-MSC DN | TYW3 | -0.872493 |
| ENSG00000185189 | 340371 | PWB-iPSC UP | NRBP2 | 0.578751 | | ENSG00000250120 | 56139 | PWB-EC UP | PCDHA10 | 2.449634289 | | ENSG00000063241 | 79763 | PWB-MSC DN | ISOC2 | -0.87368 |
| ENSG00000196670 | 643836 | PWB-iPSC UP | ZFP62 | 0.578315 | | ENSG00000254951 | 283299 | PWB-EC UP | LOC283299 | 2.418903787 | | ENSG00000163644 | 152926 | PWB-MSC DN | PPM1K | -0.878191 |
| ENSG00000137460 | 85462 | PWB-iPSC UP | FHDC1 | 0.577155 | | ENSG00000178726 | 7056 | PWB-EC UP | THBD | 2.404911057 | | ENSG00000184005 | 256435 | PWB-MSC DN | ST6GALNAC3 | -0.878649 |
| ENSG00000178567 | 9852 | PWB-iPSC UP | EPM2AIP1 | 0.576484 | | ENSG00000206262 | 401089 | PWB-EC UP | FOXL2NB | 2.378693728 | | ENSG00000266472 | 54460 | PWB-MSC DN | MRPS21 | -0.882758 |
| ENSG00000268350 | 29057 | PWB-iPSC UP | FAM156A | 0.574094 | | ENSG00000229089 | 729171 | PWB-EC UP | ANKRD20A8P | 2.371524013 | | ENSG00000124440 | 64344 | PWB-MSC DN | HIF3A | -0.884999 |
| ENSG00000179304 | 727866 | PWB-iPSC UP | FAM156B | 0.571901 | | ENSG00000236790 | 339789 | PWB-EC UP | LINC00299 | 2.349779066 | | ENSG00000275004 | 140883 | PWB-MSC DN | ZNF280B | -0.891856 |
| ENSG00000162104 | 115 | PWB-iPSC UP | ADCY9 | 0.571104 | | ENSG00000253485 | 56110 | PWB-EC UP | PCDHGA5 | 2.342897952 | | ENSG00000115084 | 80255 | PWB-MSC DN | SLC35F5 | -0.893555 |
| ENSG00000152578 | 2893 | PWB-iPSC UP | GRIA4 | 0.570435 | | ENSG00000106772 | 158471 | PWB-EC UP | PRUNE2 | 2.331299446 | | ENSG00000113396 | 28965 | PWB-MSC DN | SLC27A6 | -0.894437 |
| ENSG00000178971 | 80169 | PWB-iPSC UP | CTC1 | 0.569441 | | ENSG00000188596 | 144535 | PWB-EC UP | CFAP54 | 2.32985931 | | ENSG00000273136 | 101060684 | PWB-MSC DN | NBPF26 | -0.900074 |
| ENSG00000112159 | 23195 | PWB-iPSC UP | MDN1 | 0.56741 | | ENSG00000134198 | 10100 | PWB-EC UP | TSPAN2 | 2.314224255 | | ENSG00000142046 | 641649 | PWB-MSC DN | TMEM91 | -0.901248 |
| ENSG00000115290 | 2888 | PWB-iPSC UP | GRB14 | 0.566231 | | ENSG00000167037 | 129049 | PWB-EC UP | SGSM1 | 2.305758661 | | ENSG00000173546 | 1464 | PWB-MSC DN | CSPG4 | -0.901561 |
| ENSG00000120963 | 51123 | PWB-iPSC UP | ZNF706 | 0.565258 | | ENSG00000152192 | 5457 | PWB-EC UP | POU4F1 | 2.29256669 | | ENSG00000158296 | 64849 | PWB-MSC DN | SLC13A3 | -0.902188 |
| ENSG00000117335 | 4179 | PWB-iPSC UP | CD46 | 0.562454 | | ENSG00000243896 | 401427 | PWB-EC UP | OR2A7 | 2.279204576 | | ENSG00000134352 | 3572 | PWB-MSC DN | IL6ST | -0.902192 |
| ENSG00000250120 | 56139 | PWB-iPSC UP | PCDHA10 | 0.561155 | | ENSG00000187527 | 344905 | PWB-EC UP | ATP13A5 | 2.277409817 | | ENSG00000156140 | 9508 | PWB-MSC DN | ADAMTS3 | -0.902238 |
| ENSG00000099139 | 5125 | PWB-iPSC UP | PCSK5 | 0.559952 | | ENSG00000140873 | 170692 | PWB-EC UP | ADAMTS18 | 2.2718627 | | ENSG00000269556 | 84548 | PWB-MSC DN | TMEM185A | -0.905716 |
| ENSG00000105875 | 29062 | PWB-iPSC UP | WDR91 | 0.556906 | | ENSG00000039139 | 1767 | PWB-EC UP | DNAH5 | 2.265159271 | | ENSG00000142871 | 3491 | PWB-MSC DN | CCN1 | -0.909576 |
| ENSG00000163820 | 79443 | PWB-iPSC UP | FYCO1 | 0.554556 | | ENSG00000204969 | 56146 | PWB-EC UP | PCDHA2 | 2.263222941 | | ENSG00000148680 | 3363 | PWB-MSC DN | HTR7 | -0.915438 |
| ENSG00000265185 | 26851 | PWB-iPSC UP | SNORD3B-1 | 0.55294 | | ENSG00000173535 | 8794 | PWB-EC UP | TNFRSF10C | 2.263170361 | | ENSG00000115226 | 64838 | PWB-MSC DN | FNDC4 | -0.916499 |
| ENSG00000127603 | 23499 | PWB-iPSC UP | MACF1 | 0.552474 | | ENSG00000215186 | 55889 | PWB-EC UP | GOLGA6B | 2.258295967 | | ENSG00000115935 | 7456 | PWB-MSC DN | WIPF1 | -0.916919 |
| ENSG00000141458 | 4864 | PWB-iPSC UP | NPC1 | 0.551529 | | ENSG00000204956 | 56114 | PWB-EC UP | PCDHGA1 | 2.25715279 | | ENSG00000125355 | 55026 | PWB-MSC DN | TMEM255A | -0.920434 |
| ENSG00000262074 | 780852 | PWB-iPSC UP | SNORD3B-2 | 0.550664 | | ENSG00000227507 | 4050 | PWB-EC UP | LTB | 2.23868732 | | ENSG00000165152 | 84302 | PWB-MSC DN | PGAP4 | -0.922937 |
| ENSG00000132359 | 23108 | PWB-iPSC UP | RAP1GAP2 | 0.550333 | | ENSG00000184408 | 3751 | PWB-EC UP | KCND2 | 2.221550383 | | ENSG00000166450 | 283659 | PWB-MSC DN | PRTG | -0.925154 |
| ENSG00000198000 | 55035 | PWB-iPSC UP | NOL8 | 0.546936 | | ENSG00000180815 | 389840 | PWB-EC UP | MAP3K15 | 2.220693133 | | ENSG00000134440 | 4677 | PWB-MSC DN | NARS1 | -0.92586 |
| ENSG00000213988 | 7643 | PWB-iPSC UP | ZNF90 | 0.545525 | | ENSG00000183770 | 668 | PWB-EC UP | FOXL2 | 2.213182249 | | ENSG00000105278 | 23217 | PWB-MSC DN | ZFR2 | -0.932672 |
| ENSG00000039560 | 26064 | PWB-iPSC UP | RAI14 | 0.543982 | | ENSG00000223839 | 100133036 | PWB-EC UP | FAM95B1 | 2.211777752 | | ENSG00000100321 | 9145 | PWB-MSC DN | SYNGR1 | -0.938961 |
| ENSG00000007341 | 54879 | PWB-iPSC UP | ST7L | 0.5412 | | ENSG00000101017 | 958 | PWB-EC UP | CD40 | 2.200470235 | | ENSG00000131037 | 54869 | PWB-MSC DN | EPS8L1 | -0.939163 |
| ENSG00000073008 | 5817 | PWB-iPSC UP | PVR | 0.540441 | | ENSG00000255408 | 56145 | PWB-EC UP | PCDHA3 | 2.187806932 | | ENSG00000152256 | 5163 | PWB-MSC DN | PDK1 | -0.940536 |
| ENSG00000136141 | 23143 | PWB-iPSC UP | LRCH1 | 0.5394 | | ENSG00000249816 | 157381 | PWB-EC UP | LINC00964 | 2.164480322 | | ENSG00000117586 | 7292 | PWB-MSC DN | TNFSF4 | -0.944344 |
| ENSG00000133858 | 196441 | PWB-iPSC UP | ZFC3H1 | 0.537925 | | ENSG00000130433 | 59285 | PWB-EC UP | CACNG6 | 2.129884104 | | ENSG00000242028 | 25764 | PWB-MSC DN | HYPK | -0.945545 |
| ENSG00000147601 | 7013 | PWB-iPSC UP | TERF1 | 0.537711 | | ENSG00000254221 | 56104 | PWB-EC UP | PCDHGB1 | 2.104224165 | | ENSG00000135312 | 3351 | PWB-MSC DN | HTR1B | -0.945624 |
| ENSG00000122515 | 83637 | PWB-iPSC UP | ZMIZ2 | 0.536008 | | ENSG00000153347 | 153643 | PWB-EC UP | FAM81B | 2.103220957 | | ENSG00000089163 | 23409 | PWB-MSC DN | SIRT4 | -0.946789 |
| ENSG00000133401 | 23037 | PWB-iPSC UP | PDZD2 | 0.535844 | | ENSG00000269155 | 441179 | PWB-EC UP | LOC441179 | 2.101228772 | | ENSG00000150995 | 3708 | PWB-MSC DN | ITPR1 | -0.958824 |
| ENSG00000131067 | 2686 | PWB-iPSC UP | GGT7 | 0.52867 | | ENSG00000274628 | 105379252 | PWB-EC UP | LOC10537925 | 2.090444949 | | ENSG00000164463 | 153222 | PWB-MSC DN | CREBRF | -0.958837 |
| ENSG00000161791 | 91010 | PWB-iPSC UP | FMNL3 | 0.5282 | | ENSG00000105825 | 7980 | PWB-EC UP | TFPI2 | 2.076601006 | | ENSG00000215914 | 8511 | PWB-MSC DN | MMP23A | -0.959998 |
| ENSG00000077044 | 8527 | PWB-iPSC UP | DGKD | 0.523504 | | ENSG00000139597 | 90634 | PWB-EC UP | N4BP2L1 | 2.063001613 | | ENSG00000122824 | 170685 | PWB-MSC DN | NUDT10 | -0.968346 |
| ENSG00000175054 | 545 | PWB-iPSC UP | ATR | 0.523178 | | ENSG00000172014 | 728747 | PWB-EC UP | ANKRD20A4 | 2.060939189 | | ENSG00000165434 | 283209 | PWB-MSC DN | PGM2L1 | -0.968516 |
| ENSG00000147050 | 7403 | PWB-iPSC UP | KDM6A | 0.522961 | | ENSG00000001561 | 22875 | PWB-EC UP | ENPP4 | 2.054550998 | | ENSG00000118160 | 6543 | PWB-MSC DN | SLC8A2 | -0.968932 |
| ENSG00000118276 | 9331 | PWB-iPSC UP | B4GALT6 | 0.52163 | | ENSG00000275553 | 107983955 | PWB-EC UP | ELOA3C | 2.045290937 | | ENSG00000126803 | 3306 | PWB-MSC DN | HSPA2 | -0.969018 |
| ENSG00000173258 | 158399 | PWB-iPSC UP | ZNF483 | 0.518419 | | ENSG00000112232 | 202559 | PWB-EC UP | KHDRBS2 | 2.043685666 | | ENSG00000022267 | 2273 | PWB-MSC DN | FHL1 | -0.970126 |
| ENSG00000102699 | 143 | PWB-iPSC UP | PARP4 | 0.518288 | | ENSG00000107295 | 6456 | PWB-EC UP | SH3GL2 | 2.020340808 | | ENSG00000255794 | 196475 | PWB-MSC DN | RMST | -0.971041 |
| ENSG00000182841 | 91695 | PWB-iPSC UP | RRP7BP | 0.516559 | | ENSG00000271605 | 284021 | PWB-EC UP | MILR1 | 2.011334216 | | ENSG00000164933 | 81034 | PWB-MSC DN | SLC25A32 | -0.971814 |
| ENSG00000134815 | 9704 | PWB-iPSC UP | DHX34 | 0.516025 | | ENSG00000275950 | 102465433 | PWB-EC UP | MIR6724-1 | 1.959303348 | | ENSG00000250241 | 101927359 | PWB-MSC DN | LOC101927359 | -0.974689 |
| ENSG00000185436 | 163702 | PWB-iPSC UP | IFNLR1 | 0.510184 | | ENSG00000005189 | 81691 | PWB-EC UP | REXO5 | 1.947540906 | | ENSG00000123119 | 64168 | PWB-MSC DN | NECAB1 | -0.975209 |
| ENSG00000186777 | 654254 | PWB-iPSC UP | ZNF732 | 0.509356 | | ENSG00000111181 | 6539 | PWB-EC UP | SLC6A12 | 1.944716893 | | ENSG00000225783 | 440823 | PWB-MSC DN | MIAT | -0.97981 |
| ENSG00000215126 | 644019 | PWB-iPSC UP | CBWD6 | 0.508543 | | ENSG00000101160 | 1522 | PWB-EC UP | CTSZ | 1.934779405 | | ENSG00000128655 | 50940 | PWB-MSC DN | PDE11A | -0.980818 |
| ENSG00000167280 | 64772 | PWB-iPSC UP | ENGASE | 0.50811 | | ENSG00000221926 | 10626 | PWB-EC UP | TRIM16 | 1.926587157 | | ENSG00000109339 | 5602 | PWB-MSC DN | MAPK10 | -0.980939 |
| ENSG00000166770 | 100128252 | PWB-iPSC UP | ZNF667-AS1 | 0.508093 | | ENSG00000021355 | 1992 | PWB-EC UP | SERPINB1 | 1.92126458 | | ENSG00000197566 | 57547 | PWB-MSC DN | ZNF624 | -0.983056 |
| ENSG00000232956 | 285958 | PWB-iPSC UP | SNHG15 | 0.506185 | | ENSG00000115828 | 25797 | PWB-EC UP | QPCT | 1.89601825 | | ENSG00000169744 | 9079 | PWB-MSC DN | LDB2 | -0.985956 |
| ENSG00000263934 | 780851 | PWB-iPSC UP | SNORD3A | 0.504783 | | ENSG00000133101 | 8900 | PWB-EC UP | CCNA1 | 1.886799233 | | ENSG00000178201 | 57191 | PWB-MSC DN | VN1R1 | -0.986809 |
| ENSG00000152749 | 160897 | PWB-iPSC UP | GPR180 | 0.504353 | | ENSG00000178033 | 254228 | PWB-EC UP | CALHM5 | 1.870011489 | | ENSG00000144746 | 10550 | PWB-MSC DN | ARL6IP5 | -1.001019 |
| ENSG00000106261 | 7586 | PWB-iPSC UP | ZKSCAN1 | 0.503358 | | ENSG00000118503 | 7128 | PWB-EC UP | TNFAIP3 | 1.867274633 | | ENSG00000243753 | 3139 | PWB-MSC DN | HLA-L | -1.001308 |

| | | | | | | | | | | | | | | |
|---|---|---|---|---|---|---|---|---|---|---|---|---|---|---|
| ENSG00000213676 | 1388 | PWB-iPSC UP | ATF6B | 0.502879 | ENSG00000277379 | 103504739 | PWB-EC UP | MIR6724-3 | 1.838044645 | ENSG00000116761 | 1491 | PWB-MSC DN | CTH | -1.007638 |
| ENSG00000089006 | 27131 | PWB-iPSC UP | SNX5 | 0.502316 | ENSG00000110455 | 84680 | PWB-EC UP | ACCS | 1.826882401 | ENSG00000273079 | 2904 | PWB-MSC DN | GRIN2B | -1.008013 |
| ENSG00000119684 | 27030 | PWB-iPSC UP | MLH3 | 0.502214 | ENSG00000167555 | 84436 | PWB-EC UP | ZNF528 | 1.803642254 | ENSG00000138061 | 1545 | PWB-MSC DN | CYP1B1 | -1.009886 |
| ENSG00000078043 | 9063 | PWB-iPSC UP | PIAS2 | 0.497986 | ENSG00000154265 | 23461 | PWB-EC UP | ABCA5 | 1.801867318 | ENSG00000186951 | 5465 | PWB-MSC DN | PPARA | -1.011069 |
| ENSG00000157741 | 254048 | PWB-iPSC UP | UBN2 | 0.497625 | ENSG00000122420 | 5737 | PWB-EC UP | PTGFR | 1.79821502 | ENSG00000130540 | 25830 | PWB-MSC DN | SULT4A1 | -1.012286 |
| ENSG00000169871 | 81844 | PWB-iPSC UP | TRIM56 | 0.497577 | ENSG00000276203 | 441425 | PWB-EC UP | ANKRD20A3 | 1.788796199 | ENSG00000189409 | 8510 | PWB-MSC DN | MMP23B | -1.013414 |
| ENSG00000152700 | 51128 | PWB-iPSC UP | SAR1B | 0.496333 | ENSG00000164659 | 222223 | PWB-EC UP | KIAA1324L | 1.776045292 | ENSG00000100968 | 4776 | PWB-MSC DN | NFATC4 | -1.013878 |
| ENSG00000108848 | 51747 | PWB-iPSC UP | LUC7L3 | 0.494594 | ENSG00000145365 | 92610 | PWB-EC UP | TIFA | 1.734445772 | ENSG00000136160 | 1910 | PWB-MSC DN | EDNRB | -1.014672 |
| ENSG00000258890 | 90799 | PWB-iPSC UP | CEP95 | 0.492173 | ENSG00000145198 | 90113 | PWB-EC UP | VWA5B2 | 1.71121146 | ENSG00000213420 | 221914 | PWB-MSC DN | GPC2 | -1.016259 |
| ENSG00000254681 | 110006322 | PWB-iPSC UP | PKD1P5-LOC10537 | 0.490017 | ENSG00000154928 | 2047 | PWB-EC UP | EPHB1 | 1.701974755 | ENSG00000157613 | 90993 | PWB-MSC DN | CREB3L1 | -1.022574 |
| ENSG00000090905 | 27327 | PWB-iPSC UP | TNRC6A | 0.479074 | ENSG00000176293 | 7694 | PWB-EC UP | ZNF135 | 1.693366338 | ENSG00000170542 | 5272 | PWB-MSC DN | SERPINB9 | -1.023991 |
| ENSG00000146067 | 54540 | PWB-iPSC UP | FAM193B | 0.477387 | ENSG00000034677 | 25897 | PWB-EC UP | RNF19A | 1.673225006 | ENSG00000174720 | 51574 | PWB-MSC DN | LARP7 | -1.025654 |
| ENSG00000137955 | 5876 | PWB-iPSC UP | RABGGTB | 0.476453 | ENSG00000164647 | 26872 | PWB-EC UP | STEAP1 | 1.671718917 | ENSG00000158560 | 1780 | PWB-MSC DN | DYNC1I1 | -1.030364 |
| ENSG00000075391 | 9462 | PWB-iPSC UP | RASAL2 | 0.472644 | ENSG00000223482 | 728190 | PWB-EC UP | NUTM2A-AS1 | 1.668552784 | ENSG00000114796 | 54800 | PWB-MSC DN | KLHL24 | -1.031899 |
| ENSG00000123560 | 5354 | PWB-iPSC UP | PLP1 | 0.468359 | ENSG00000196562 | 55959 | PWB-EC UP | SULF2 | 1.648525144 | ENSG00000130270 | 148229 | PWB-MSC DN | ATP8B3 | -1.032567 |
| ENSG00000121940 | 23155 | PWB-iPSC UP | CLCC1 | 0.467661 | ENSG00000146232 | 4794 | PWB-EC UP | NFKBIE | 1.645932945 | ENSG00000248730 | 100852408 | PWB-MSC DN | EGFLAM-AS4 | -1.036372 |
| ENSG00000048828 | 23196 | PWB-iPSC UP | FAM120A | 0.463939 | ENSG00000107798 | 3988 | PWB-EC UP | LIPA | 1.636745166 | ENSG00000070193 | 2255 | PWB-MSC DN | FGF10 | -1.037585 |
| ENSG00000135749 | 80003 | PWB-iPSC UP | PCNX2 | 0.462939 | ENSG00000143469 | 255928 | PWB-EC UP | SYT14 | 1.616461965 | ENSG00000266714 | 80022 | PWB-MSC DN | MYO15B | -1.045181 |
| ENSG00000145390 | 54532 | PWB-iPSC UP | USP53 | 0.460628 | ENSG00000144843 | 141 | PWB-EC UP | ADPRH | 1.614693939 | ENSG00000093010 | 1312 | PWB-MSC DN | COMT | -1.047308 |
| ENSG00000095066 | 29911 | PWB-iPSC UP | HOOK2 | 0.458639 | ENSG00000143153 | 481 | PWB-EC UP | ATP1B1 | 1.569466849 | ENSG00000174469 | 26047 | PWB-MSC DN | CNTNAP2 | -1.049625 |
| ENSG00000181450 | 339500 | PWB-iPSC UP | ZNF678 | 0.457885 | ENSG00000168899 | 10791 | PWB-EC UP | VAMP5 | 1.506503512 | ENSG00000161677 | 126119 | PWB-MSC DN | JOSD2 | -1.051071 |
| ENSG00000102710 | 55578 | PWB-iPSC UP | SUPT20H | 0.456848 | ENSG00000167191 | 51704 | PWB-EC UP | GPRC5B | 1.505652106 | ENSG00000182621 | 23236 | PWB-MSC DN | PLCB1 | -1.05208 |
| ENSG00000132424 | 25957 | PWB-iPSC UP | PNISR | 0.453783 | ENSG00000175893 | 340481 | PWB-EC UP | ZDHHC21 | 1.504783907 | ENSG00000261652 | 145788 | PWB-MSC DN | C15orf65 | -1.052682 |
| ENSG00000160336 | 388561 | PWB-iPSC UP | ZNF761 | 0.446356 | ENSG00000176978 | 29952 | PWB-EC UP | DPP7 | 1.497943701 | ENSG00000081181 | 384 | PWB-MSC DN | ARG2 | -1.053203 |
| ENSG00000145439 | 84869 | PWB-iPSC UP | CBR4 | 0.445231 | ENSG00000139178 | 51279 | PWB-EC UP | C1RL | 1.494509464 | ENSG00000142794 | 84224 | PWB-MSC DN | NBPF3 | -1.054105 |
| ENSG00000166402 | 7275 | PWB-iPSC UP | TUB | 0.443354 | ENSG00000112759 | 2030 | PWB-EC UP | SLC29A1 | 1.405780396 | ENSG00000005249 | 5577 | PWB-MSC DN | PRKAR2B | -1.055635 |
| ENSG00000112234 | 26235 | PWB-iPSC UP | FBXL4 | 0.441497 | ENSG00000164251 | 2150 | PWB-EC UP | F2RL1 | 1.396945675 | ENSG00000174600 | 1240 | PWB-MSC DN | CMKLR1 | -1.056841 |
| ENSG00000003509 | 55471 | PWB-iPSC UP | NDUFAF7 | 0.440445 | ENSG00000104689 | 8797 | PWB-EC UP | TNFRSF10A | 1.389496664 | ENSG00000128039 | 79644 | PWB-MSC DN | SRD5A3 | -1.058908 |
| ENSG00000187955 | 7373 | PWB-iPSC UP | COL14A1 | 0.440227 | ENSG00000224914 | 439994 | PWB-EC UP | LINC00863 | 1.386686529 | ENSG00000107611 | 8029 | PWB-MSC DN | CUBN | -1.060665 |
| ENSG00000197948 | 89848 | PWB-iPSC UP | FCHSD1 | 0.435152 | ENSG00000073464 | 1183 | PWB-EC UP | CLCN4 | 1.363294478 | ENSG00000126458 | 6237 | PWB-MSC DN | RRAS | -1.062081 |
| ENSG00000198625 | 4194 | PWB-iPSC UP | MDM4 | 0.42469 | ENSG00000168386 | 11259 | PWB-EC UP | FILIP1L | 1.321105875 | ENSG00000101605 | 8736 | PWB-MSC DN | MYOM1 | -1.063932 |
| ENSG00000111642 | 1108 | PWB-iPSC DN | CHD4 | -0.40352 | ENSG00000050344 | 9603 | PWB-EC UP | NFE2L3 | 1.320325535 | ENSG00000079102 | 862 | PWB-MSC DN | RUNX1T1 | -1.06872 |
| ENSG00000073350 | 3993 | PWB-iPSC DN | LLGL2 | -0.41156 | ENSG00000026103 | 355 | PWB-EC UP | FAS | 1.30074685 | ENSG00000275126 | 8368 | PWB-MSC DN | H4C13 | -1.068898 |
| ENSG00000141551 | 1453 | PWB-iPSC DN | CSNK1D | -0.41457 | ENSG00000188677 | 29780 | PWB-EC UP | PARVB | 1.291475151 | ENSG00000175691 | 58492 | PWB-MSC DN | ZNF77 | -1.06902 |
| ENSG00000116191 | 55103 | PWB-iPSC DN | RALGPS2 | -0.41791 | ENSG00000066926 | 2235 | PWB-EC UP | FECH | 1.204089925 | ENSG00000138468 | 57337 | PWB-MSC DN | SENP7 | -1.071445 |
| ENSG00000198554 | 11169 | PWB-iPSC DN | WDHD1 | -0.42181 | ENSG00000112186 | 10486 | PWB-EC DN | CAP2 | -1.301857642 | ENSG00000070540 | 55062 | PWB-MSC DN | WIPI1 | -1.071838 |
| ENSG00000145860 | 153830 | PWB-iPSC DN | RNF145 | -0.42235 | ENSG00000111885 | 4121 | PWB-EC DN | MAN1A1 | -1.372823774 | ENSG00000146700 | 136853 | PWB-MSC DN | SSC4D | -1.071857 |
| ENSG00000104872 | 55011 | PWB-iPSC DN | PIH1D1 | -0.4233 | ENSG00000105355 | 10226 | PWB-EC DN | PLIN3 | -1.386100668 | ENSG00000183010 | 5831 | PWB-MSC DN | PYCR1 | -1.075161 |
| ENSG00000182287 | 8905 | PWB-iPSC DN | AP1S2 | -0.43673 | ENSG00000277067 | 102724843 | PWB-EC DN | LOC10272484 | -1.391993044 | ENSG00000170915 | 85315 | PWB-MSC DN | PAQR8 | -1.080769 |
| ENSG00000089248 | 10961 | PWB-iPSC DN | ERP29 | -0.44202 | ENSG00000276077 | 102724951 | PWB-EC DN | LOC10272495 | -1.394682963 | ENSG00000183826 | 114781 | PWB-MSC DN | BTBD9 | -1.081456 |
| ENSG00000077942 | 2192 | PWB-iPSC DN | FBLN1 | -0.4437 | ENSG00000274333 | 102724219 | PWB-EC DN | LOC10272421 | -1.404088185 | ENSG00000170820 | 2492 | PWB-MSC DN | FSHR | -1.082363 |
| ENSG00000139990 | 8816 | PWB-iPSC DN | DCAF5 | -0.4452 | ENSG00000275496 | 102724701 | PWB-EC DN | LOC10272470 | -1.438005561 | ENSG00000267040 | 100505549 | PWB-MSC DN | LOC100505549 | -1.085386 |
| ENSG00000164096 | 401152 | PWB-iPSC DN | C4orf3 | -0.45221 | ENSG00000171604 | 51523 | PWB-EC DN | CXXC5 | -1.446467993 | ENSG00000187260 | 349136 | PWB-MSC DN | WDR86 | -1.087064 |
| ENSG00000102241 | 27336 | PWB-iPSC DN | HTATSF1 | -0.45334 | ENSG00000277991 | 102723360 | PWB-EC DN | LOC10272336 | -1.454490176 | ENSG00000125531 | 79025 | PWB-MSC DN | FNDC11 | -1.089107 |
| ENSG00000204977 | 10206 | PWB-iPSC DN | TRIM13 | -0.45528 | ENSG00000128602 | 6608 | PWB-EC DN | SMO | -1.515734302 | ENSG00000165300 | 26050 | PWB-MSC DN | SLITRK5 | -1.091411 |
| ENSG00000077380 | 1781 | PWB-iPSC DN | DYNC1I2 | -0.45531 | ENSG00000271425 | 100132406 | PWB-EC DN | NBPF10 | -1.550901749 | ENSG00000226328 | 100506714 | PWB-MSC DN | NUP50-DT | -1.091885 |
| ENSG00000184697 | 9074 | PWB-iPSC DN | CLDN6 | -0.45877 | ENSG00000201098 | 6084 | PWB-EC DN | RNY1 | -1.571493926 | ENSG00000137094 | 25822 | PWB-MSC DN | DNAJB5 | -1.095464 |
| ENSG00000141646 | 4089 | PWB-iPSC DN | SMAD4 | -0.4588 | ENSG00000127418 | 53834 | PWB-EC DN | FGFRL1 | -1.676983238 | ENSG00000006468 | 2115 | PWB-MSC DN | ETV1 | -1.095576 |
| ENSG00000137693 | 10413 | PWB-iPSC DN | YAP1 | -0.46041 | ENSG00000145908 | 91975 | PWB-EC DN | ZNF300 | -1.679115976 | ENSG00000288015 | 105370224 | PWB-MSC DN | LOC105370224 | -1.096313 |
| ENSG00000127663 | 23030 | PWB-iPSC DN | KDM4B | -0.46084 | ENSG00000130821 | 6535 | PWB-EC DN | SLC6A8 | -1.691715817 | ENSG00000272690 | 107986100 | PWB-MSC DN | LINC02018 | -1.09834 |
| ENSG00000168385 | 4735 | PWB-iPSC DN | SEPTIN2 | -0.46106 | ENSG00000129116 | 23022 | PWB-EC DN | PALLD | -1.748659558 | ENSG00000061918 | 2983 | PWB-MSC DN | GUCY1B1 | -1.104528 |
| ENSG00000113761 | 23567 | PWB-iPSC DN | ZNF346 | -0.46163 | ENSG00000150347 | 84159 | PWB-EC DN | ARID5B | -1.782709707 | ENSG00000105889 | 256227 | PWB-MSC DN | STEAP1B | -1.104684 |
| ENSG00000146085 | 4594 | PWB-iPSC DN | MMUT | -0.46188 | ENSG00000105245 | 9253 | PWB-EC DN | NUMBL | -1.801097254 | ENSG00000106484 | 4232 | PWB-MSC DN | MEST | -1.106534 |
| ENSG00000040933 | 3631 | PWB-iPSC DN | INPP4A | -0.46201 | ENSG00000273703 | 8342 | PWB-EC DN | H2BC14 | -1.804242299 | ENSG00000180543 | 85453 | PWB-MSC DN | TSPYL5 | -1.11712 |
| ENSG00000118898 | 5493 | PWB-iPSC DN | PPL | -0.46319 | ENSG00000184557 | 9021 | PWB-EC DN | SOCS3 | -1.812490721 | ENSG00000115239 | 51130 | PWB-MSC DN | ASB3 | -1.120694 |
| ENSG00000131899 | 3996 | PWB-iPSC DN | LLGL1 | -0.46451 | ENSG00000171621 | 80176 | PWB-EC DN | SPSB1 | -1.813675391 | ENSG00000261455 | 100128822 | PWB-MSC DN | LINC01003 | -1.12129 |
| ENSG00000087152 | 56970 | PWB-iPSC DN | ATXN7L3 | -0.46749 | ENSG00000164176 | 10085 | PWB-EC DN | EDIL3 | -1.828212832 | ENSG00000159761 | 388284 | PWB-MSC DN | C16orf86 | -1.121657 |
| ENSG00000187772 | 389421 | PWB-iPSC DN | LIN28B | -0.46782 | ENSG00000147027 | 83604 | PWB-EC DN | TMEM47 | -1.846787284 | ENSG00000280987 | 9782 | PWB-MSC DN | MATR3 | -1.122884 |
| ENSG00000140829 | 9785 | PWB-iPSC DN | DHX38 | -0.468 | ENSG00000115641 | 2274 | PWB-EC DN | FHL2 | -1.883988491 | ENSG00000066230 | 6550 | PWB-MSC DN | SLC9A3 | -1.123346 |
| ENSG00000135677 | 2799 | PWB-iPSC DN | GNS | -0.46814 | ENSG00000151208 | 9231 | PWB-EC DN | DLG5 | -1.893157111 | ENSG00000179542 | 139065 | PWB-MSC DN | SLITRK4 | -1.125027 |
| ENSG00000103275 | 7329 | PWB-iPSC DN | UBE2I | -0.46891 | ENSG00000167964 | 25837 | PWB-EC DN | RAB26 | -1.90114623 | ENSG00000205710 | 100130311 | PWB-MSC DN | C17orf107 | -1.126772 |
| ENSG00000132842 | 8546 | PWB-iPSC DN | AP3B1 | -0.46996 | ENSG00000109452 | 8821 | PWB-EC DN | INPP4B | -1.910277556 | ENSG00000237036 | 220930 | PWB-MSC DN | ZEB1-AS1 | -1.128518 |
| ENSG00000114126 | 7029 | PWB-iPSC DN | TFDP2 | -0.47173 | ENSG00000157680 | 9162 | PWB-EC DN | DGKI | -1.919130774 | ENSG00000162576 | 54587 | PWB-MSC DN | MXRA8 | -1.129148 |
| ENSG00000171603 | 22883 | PWB-iPSC DN | CLSTN1 | -0.47332 | ENSG00000143341 | 83872 | PWB-EC DN | HMCN1 | -1.947429614 | ENSG00000129675 | 9459 | PWB-MSC DN | ARHGEF6 | -1.136453 |
| ENSG00000124795 | 7913 | PWB-iPSC DN | DEK | -0.47443 | ENSG00000270629 | 25832 | PWB-EC DN | NBPF14 | -1.959651111 | ENSG00000154553 | 27295 | PWB-MSC DN | PDLIM3 | -1.13748 |
| ENSG00000102038 | 6594 | PWB-iPSC DN | SMARCA1 | -0.47489 | ENSG00000022267 | 2273 | PWB-EC DN | FHL1 | -1.981626099 | ENSG00000246130 | 286059 | PWB-MSC DN | LOC286059 | -1.138653 |
| ENSG00000111602 | 8914 | PWB-iPSC DN | TIMELESS | -0.4749 | ENSG00000140263 | 6652 | PWB-EC DN | SORD | -1.990565795 | ENSG00000163565 | 3428 | PWB-MSC DN | IFI16 | -1.139777 |
| ENSG00000131171 | 6451 | PWB-iPSC DN | SH3BGRL | -0.47579 | ENSG00000121716 | 29990 | PWB-EC DN | PILRB | -2.004401881 | ENSG00000175449 | 317671 | PWB-MSC DN | RFESD | -1.142156 |
| ENSG00000143398 | 8394 | PWB-iPSC DN | PIP5K1A | -0.4758 | ENSG00000136153 | 4008 | PWB-EC DN | LMO7 | -2.038002459 | ENSG00000167311 | 116969 | PWB-MSC DN | ART5 | -1.144775 |
| ENSG00000129353 | 57153 | PWB-iPSC DN | SLC44A2 | -0.47582 | ENSG00000187243 | 81557 | PWB-EC DN | MAGED4B | -2.06968936 | ENSG00000261879 | 100130950 | PWB-MSC DN | LOC100130950 | -1.146086 |
| ENSG00000172500 | 9158 | PWB-iPSC DN | FIBP | -0.47653 | ENSG00000154545 | 728239 | PWB-EC DN | MAGED4 | -2.069691026 | ENSG00000151320 | 9472 | PWB-MSC DN | AKAP6 | -1.149377 |
| ENSG00000163904 | 59343 | PWB-iPSC DN | SENP2 | -0.47679 | ENSG00000165655 | 84858 | PWB-EC DN | ZNF503 | -2.077819919 | ENSG00000141506 | 23533 | PWB-MSC DN | PIK3R5 | -1.150897 |
| ENSG00000196792 | 29966 | PWB-iPSC DN | STRN3 | -0.47714 | ENSG00000145087 | 9515 | PWB-EC DN | STXBP5L | -2.110204385 | ENSG00000145632 | 10769 | PWB-MSC DN | PLK2 | -1.151691 |
| ENSG00000111667 | 8078 | PWB-iPSC DN | USP5 | -0.47717 | ENSG00000178498 | 196403 | PWB-EC DN | DTX3 | -2.113562814 | ENSG00000286940 | 102725068 | PWB-MSC DN | MICB-DT | -1.153318 |
| ENSG00000047230 | 56474 | PWB-iPSC DN | CTPS2 | -0.47851 | ENSG00000170390 | 166614 | PWB-EC DN | DCLK2 | -2.114587234 | ENSG00000203952 | 347475 | PWB-MSC DN | CCDC160 | -1.15785 |
| ENSG00000129691 | 9070 | PWB-iPSC DN | ASH2L | -0.47975 | ENSG00000171522 | 5734 | PWB-EC DN | PTGER4 | -2.125453429 | ENSG00000240184 | 5098 | PWB-MSC DN | PCDHGC3 | -1.159962 |
| ENSG00000170027 | 7532 | PWB-iPSC DN | YWHAG | -0.48324 | ENSG00000168785 | 10098 | PWB-EC DN | TSPAN5 | -2.144923145 | ENSG00000198939 | 80108 | PWB-MSC DN | ZFP2 | -1.164214 |
| ENSG00000178252 | 11180 | PWB-iPSC DN | WDR6 | -0.4858 | ENSG00000134762 | 1825 | PWB-EC DN | DSC3 | -2.150573392 | ENSG00000198825 | 22876 | PWB-MSC DN | INPP5F | -1.166083 |
| ENSG00000108219 | 81619 | PWB-iPSC DN | TSPAN14 | -0.48649 | ENSG00000258947 | 10381 | PWB-EC DN | TUBB3 | -2.165036769 | ENSG00000167733 | 374875 | PWB-MSC DN | HSD11B1L | -1.168804 |
| ENSG00000167325 | 6240 | PWB-iPSC DN | RRM1 | -0.4865 | ENSG00000115318 | 84695 | PWB-EC DN | LOXL3 | -2.179868317 | ENSG00000256612 | 1556 | PWB-MSC DN | CYP2B7P | -1.172426 |
| ENSG00000182628 | 348235 | PWB-iPSC DN | SKA2 | -0.48681 | ENSG00000084710 | 22979 | PWB-EC DN | EFR3B | -2.197213402 | ENSG00000189060 | 3005 | PWB-MSC DN | H1-0 | -1.176493 |
| ENSG00000119787 | 64225 | PWB-iPSC DN | ATL2 | -0.48885 | ENSG00000166963 | 4130 | PWB-EC DN | MAP1A | -2.202279597 | ENSG00000225953 | 150538 | PWB-MSC DN | SATB2-AS1 | -1.185937 |
| ENSG00000165458 | 3636 | PWB-iPSC DN | INPPL1 | -0.48912 | ENSG00000091972 | 4345 | PWB-EC DN | CD200 | -2.202318428 | ENSG00000285280 | 105371664 | PWB-MSC DN | LOC105371664 | -1.18892 |
| ENSG00000049449 | 5954 | PWB-iPSC DN | RCN1 | -0.49011 | ENSG00000270722 | 115482718 | PWB-EC DN | RNVU1-31 | -2.258684165 | ENSG00000125170 | 55715 | PWB-MSC DN | DOK4 | -1.191225 |
| ENSG00000181915 | 84890 | PWB-iPSC DN | ADO | -0.49055 | ENSG00000136274 | 23148 | PWB-EC DN | NACAD | -2.289354556 | ENSG00000105559 | 57664 | PWB-MSC DN | PLEKHA4 | -1.19348 |
| ENSG00000130821 | 6535 | PWB-iPSC DN | SLC6A8 | -0.49122 | ENSG00000123384 | 4035 | PWB-EC DN | LRP1 | -2.289575457 | ENSG00000184672 | 138046 | PWB-MSC DN | RALYL | -1.194097 |
| ENSG00000198858 | 91300 | PWB-iPSC DN | R3HDM4 | -0.49125 | ENSG00000103260 | 79006 | PWB-EC DN | METRN | -2.291449282 | ENSG00000159433 | 57519 | PWB-MSC DN | STARD9 | -1.19953 |
| ENSG00000123933 | 10608 | PWB-iPSC DN | MXD4 | -0.49183 | ENSG00000147889 | 1029 | PWB-EC DN | CDKN2A | -2.293962347 | ENSG00000170775 | 2861 | PWB-MSC DN | GPR37 | -1.200865 |
| ENSG00000187325 | 51616 | PWB-iPSC DN | TAF9B | -0.492 | ENSG00000278588 | 8346 | PWB-EC DN | H2BC10 | -2.297183404 | ENSG00000119899 | 26503 | PWB-MSC DN | SLC17A5 | -1.201074 |
| ENSG00000127328 | 117177 | PWB-iPSC DN | RAB3IP | -0.49396 | ENSG00000107984 | 22943 | PWB-EC DN | DKK1 | -2.30500455 | ENSG00000140795 | 91807 | PWB-MSC DN | MYLK3 | -1.202035 |
| ENSG00000167081 | 5090 | PWB-iPSC DN | PBX3 | -0.49464 | ENSG00000165617 | 51339 | PWB-EC DN | DACT1 | -2.318514274 | ENSG00000185522 | 256329 | PWB-MSC DN | LMNTD2 | -1.208002 |
| ENSG00000122550 | 55975 | PWB-iPSC DN | KLHL7 | -0.49525 | ENSG00000132205 | 84034 | PWB-EC DN | EMILIN2 | -2.327811264 | ENSG00000152580 | 285313 | PWB-MSC DN | IGSF10 | -1.214756 |
| ENSG00000125835 | 6628 | PWB-iPSC DN | SNRPB | -0.49547 | ENSG00000205213 | 55366 | PWB-EC DN | LGR4 | -2.335845551 | ENSG00000149596 | 57158 | PWB-MSC DN | JPH2 | -1.217928 |
| ENSG00000102317 | 5935 | PWB-iPSC DN | RBM3 | -0.49639 | ENSG00000105605 | 59284 | PWB-EC DN | CACNG7 | -2.346118988 | ENSG00000213085 | 25790 | PWB-MSC DN | CFAP45 | -1.21899 |
| ENSG00000101115 | 57167 | PWB-iPSC DN | SALL4 | -0.49676 | ENSG00000152953 | 55351 | PWB-EC DN | STK32B | -2.347213277 | ENSG00000137331 | 8870 | PWB-MSC DN | IER3 | -1.219404 |
| ENSG00000078304 | 5527 | PWB-iPSC DN | PPP2R5C | -0.49727 | ENSG00000026508 | 960 | PWB-EC DN | CD44 | -2.348331666 | ENSG00000161912 | 221442 | PWB-MSC DN | ADCY10P1 | -1.219529 |
| ENSG00000103769 | 8766 | PWB-iPSC DN | RAB11A | -0.49789 | ENSG00000081320 | 9262 | PWB-EC DN | STK17B | -2.350410507 | ENSG00000129116 | 23022 | PWB-MSC DN | PALLD | -1.219947 |
| ENSG00000198356 | 439 | PWB-iPSC DN | GET3 | -0.49941 | ENSG00000213420 | 221914 | PWB-EC DN | GPC2 | -2.358955974 | ENSG00000166342 | 81832 | PWB-MSC DN | NETO1 | -1.222525 |
| ENSG00000122378 | 84293 | PWB-iPSC DN | PRXL2A | -0.49969 | ENSG00000160293 | 7410 | PWB-EC DN | VAV2 | -2.361235879 | ENSG00000119630 | 5228 | PWB-MSC DN | PGF | -1.234831 |
| ENSG00000161618 | 126133 | PWB-iPSC DN | ALDH16A1 | -0.50123 | ENSG00000168685 | 3575 | PWB-EC DN | IL7R | -2.362041111 | ENSG00000187624 | 400566 | PWB-MSC DN | C17orf97 | -1.238581 |
| ENSG00000213639 | 5500 | PWB-iPSC DN | PPP1CB | -0.50125 | ENSG00000125675 | 2892 | PWB-EC DN | GRIA3 | -2.367023424 | ENSG00000226752 | 253039 | PWB-MSC DN | CUTALP | -1.252823 |
| ENSG00000143727 | 52 | PWB-iPSC DN | ACP1 | -0.50131 | ENSG00000061918 | 2983 | PWB-EC DN | GUCY1B1 | -2.371705373 | ENSG00000260941 | 644242 | PWB-MSC DN | LINC00622 | -1.25763 |
| ENSG00000001036 | 2519 | PWB-iPSC DN | FUCA2 | -0.50143 | ENSG00000120738 | 1958 | PWB-EC DN | EGR1 | -2.374223218 | ENSG00000160117 | 126549 | PWB-MSC DN | ANKLE1 | -1.263244 |
| ENSG00000137364 | 7172 | PWB-iPSC DN | TPMT | -0.50149 | ENSG00000115290 | 2888 | PWB-EC DN | GRB14 | -2.381342359 | ENSG00000282826 | 100289097 | PWB-MSC DN | FRG1CP | -1.270428 |
| ENSG00000259956 | 29890 | PWB-iPSC DN | RBM15B | -0.50237 | ENSG00000161714 | 113026 | PWB-EC DN | PLCD3 | -2.390192994 | ENSG00000255367 | 101927708 | PWB-MSC DN | LOC101927708 | -1.273159 |
| ENSG00000100426 | 9889 | PWB-iPSC DN | ZBED4 | -0.50353 | ENSG00000169184 | 4330 | PWB-EC DN | MN1 | -2.401660218 | ENSG00000142227 | 2014 | PWB-MSC DN | EMP3 | -1.274174 |
| ENSG00000174238 | 5306 | PWB-iPSC DN | PITPNA | -0.5038 | ENSG00000276903 | 8332 | PWB-EC DN | H2AC16 | -2.402973676 | ENSG00000233056 | 90625 | PWB-MSC DN | ERVH48-1 | -1.279058 |

| | | | | | | | | | | | | | | |
|---|---|---|---|---|---|---|---|---|---|---|---|---|---|---|
| ENSG00000154545 | 728239 | PWB-iPSC DN | MAGED4 | -0.50396 | ENSG00000151150 | 288 | PWB-EC DN | ANK3 | -2.419568022 | ENSG00000115085 | 7535 | PWB-MSC DN | ZAP70 | -1.279484 |
| ENSG00000140262 | 6938 | PWB-iPSC DN | TCF12 | -0.50399 | ENSG00000158163 | 199221 | PWB-EC DN | DZIP1L | -2.42300595 | ENSG00000102287 | 2564 | PWB-MSC DN | GABRE | -1.280577 |
| ENSG00000188895 | 339287 | PWB-iPSC DN | MSL1 | -0.50434 | ENSG00000276180 | 8294 | PWB-EC DN | H4C9 | -2.440907344 | ENSG00000048052 | 9734 | PWB-MSC DN | HDAC9 | -1.280916 |
| ENSG00000113595 | 373 | PWB-iPSC DN | TRIM23 | -0.50505 | ENSG00000166073 | 11245 | PWB-EC DN | GPR176 | -2.446457521 | ENSG00000234377 | 100874222 | PWB-MSC DN | OBI1-AS1 | -1.281249 |
| ENSG00000187243 | 81557 | PWB-iPSC DN | MAGED4B | -0.50599 | ENSG00000091622 | 83394 | PWB-EC DN | PITPNM3 | -2.45228184 | ENSG00000143995 | 4211 | PWB-MSC DN | MEIS1 | -1.283646 |
| ENSG00000090889 | 24137 | PWB-iPSC DN | KIF4A | -0.50685 | ENSG00000157613 | 90993 | PWB-EC DN | CREB3L1 | -2.45411996 | ENSG00000114204 | 5276 | PWB-MSC DN | SERPINI2 | -1.284254 |
| ENSG00000124151 | 8202 | PWB-iPSC DN | NCOA3 | -0.50707 | ENSG00000082014 | 6604 | PWB-EC DN | SMARCD3 | -2.462622129 | ENSG00000107518 | 26033 | PWB-MSC DN | ATRNL1 | -1.285914 |
| ENSG00000075142 | 6717 | PWB-iPSC DN | SRI | -0.5082 | ENSG00000065989 | 5141 | PWB-EC DN | PDE4A | -2.472544281 | ENSG00000106392 | 56913 | PWB-MSC DN | C1GALT1 | -1.287522 |
| ENSG00000169221 | 26000 | PWB-iPSC DN | TBC1D10B | -0.50828 | ENSG00000168140 | 114990 | PWB-EC DN | VASN | -2.473742114 | ENSG00000068831 | 10235 | PWB-MSC DN | RASGRP2 | -1.292384 |
| ENSG00000139651 | 283337 | PWB-iPSC DN | ZNF740 | -0.50843 | ENSG00000105696 | 25789 | PWB-EC DN | TMEM59L | -2.477722616 | ENSG00000164188 | 202151 | PWB-MSC DN | RANBP3L | -1.293268 |
| ENSG00000110925 | 81566 | PWB-iPSC DN | CSRNP2 | -0.50939 | ENSG00000185130 | 8340 | PWB-EC DN | H2BC13 | -2.481854178 | ENSG00000171659 | 2857 | PWB-MSC DN | GPR34 | -1.29634 |
| ENSG00000169925 | 8019 | PWB-iPSC DN | BRD3 | -0.51118 | ENSG00000100417 | 5372 | PWB-EC DN | PMM1 | -2.488185916 | ENSG00000006047 | 51087 | PWB-MSC DN | YBX2 | -1.299945 |
| ENSG00000175756 | 54998 | PWB-iPSC DN | AURKAIP1 | -0.51192 | ENSG00000007866 | 7005 | PWB-EC DN | TEAD3 | -2.520379067 | ENSG00000223770 | 101927356 | PWB-MSC DN | CACNA2D1-AS1 | -1.304847 |
| ENSG00000167470 | 90007 | PWB-iPSC DN | MIDN | -0.51301 | ENSG00000106976 | 1759 | PWB-EC DN | DNM1 | -2.520925537 | ENSG00000268182 | 147670 | PWB-MSC DN | SMIM17 | -1.306202 |
| ENSG00000099942 | 1399 | PWB-iPSC DN | CRKL | -0.51376 | ENSG00000155893 | 92370 | PWB-EC DN | PXYLP1 | -2.567989052 | ENSG00000079385 | 634 | PWB-MSC DN | CEACAM1 | -1.306329 |
| ENSG00000169155 | 23099 | PWB-iPSC DN | ZBTB43 | -0.51382 | ENSG00000123453 | 1757 | PWB-EC DN | SARDH | -2.569589201 | ENSG00000127920 | 2791 | PWB-MSC DN | GNG11 | -1.308254 |
| ENSG00000102935 | 23090 | PWB-iPSC DN | ZNF423 | -0.51452 | ENSG00000176170 | 8877 | PWB-EC DN | SPHK1 | -2.57055979 | ENSG00000241288 | 101927056 | PWB-MSC DN | LINC02614 | -1.31102 |
| ENSG00000274523 | 81554 | PWB-iPSC DN | RCC1L | -0.51507 | ENSG00000144057 | 84620 | PWB-EC DN | ST6GAL2 | -2.571890948 | ENSG00000154783 | 152273 | PWB-MSC DN | FGD5 | -1.311836 |
| ENSG00000185414 | 51263 | PWB-iPSC DN | MRPL30 | -0.51533 | ENSG00000203930 | 286411 | PWB-EC DN | LINC00632 | -2.579495021 | ENSG00000108691 | 6347 | PWB-MSC DN | CCL2 | -1.314872 |
| ENSG00000186575 | 4771 | PWB-iPSC DN | NF2 | -0.51569 | ENSG00000129195 | 54478 | PWB-EC DN | PIMREG | -2.581077934 | ENSG00000147576 | 137872 | PWB-MSC DN | ADHFE1 | -1.316204 |
| ENSG00000143321 | 3068 | PWB-iPSC DN | HDGF | -0.51645 | ENSG00000165912 | 29763 | PWB-EC DN | PACSIN3 | -2.586237944 | ENSG00000196511 | 27010 | PWB-MSC DN | TPK1 | -1.317418 |
| ENSG00000100105 | 23598 | PWB-iPSC DN | PATZ1 | -0.51715 | ENSG00000197301 | 100129940 | PWB-EC DN | HMGA2-AS1 | -2.606731475 | ENSG00000104888 | 57030 | PWB-MSC DN | SLC17A7 | -1.318267 |
| ENSG00000100461 | 55147 | PWB-iPSC DN | RBM23 | -0.51819 | ENSG00000155760 | 8324 | PWB-EC DN | FZD7 | -2.610250772 | ENSG00000166292 | 55273 | PWB-MSC DN | TMEM100 | -1.323713 |
| ENSG00000196233 | 84458 | PWB-iPSC DN | LCOR | -0.51926 | ENSG00000149948 | 8091 | PWB-EC DN | HMGA2 | -2.626169838 | ENSG00000080823 | 5891 | PWB-MSC DN | MOK | -1.324155 |
| ENSG00000128791 | 57045 | PWB-iPSC DN | TWSG1 | -0.51935 | ENSG00000132170 | 5468 | PWB-EC DN | PPARG | -2.626192665 | ENSG00000187554 | 7100 | PWB-MSC DN | TLR5 | -1.324687 |
| ENSG00000135506 | 10956 | PWB-iPSC DN | OS9 | -0.52064 | ENSG00000253716 | 100507316 | PWB-EC DN | MINCR | -2.644614457 | ENSG00000254531 | 90024 | PWB-MSC DN | FLJ20021 | -1.325125 |
| ENSG00000197746 | 5660 | PWB-iPSC DN | PSAP | -0.52071 | ENSG00000235033 | 100505635 | PWB-EC DN | DAAM2-AS1 | -2.647949955 | ENSG00000170113 | 123606 | PWB-MSC DN | NIPA1 | -1.332805 |
| ENSG00000116209 | 9528 | PWB-iPSC DN | TMEM59 | -0.52245 | ENSG00000165323 | 120114 | PWB-EC DN | FAT3 | -2.655706323 | ENSG00000233237 | 79940 | PWB-MSC DN | LINC00472 | -1.337104 |
| ENSG00000000003 | 7105 | PWB-iPSC DN | TSPAN6 | -0.52288 | ENSG00000139211 | 347902 | PWB-EC DN | AMIGO2 | -2.657962959 | ENSG00000130052 | 9754 | PWB-MSC DN | STARD8 | -1.339429 |
| ENSG00000140319 | 6727 | PWB-iPSC DN | SRP14 | -0.52493 | ENSG00000124575 | 3007 | PWB-EC DN | H1-3 | -2.658776371 | ENSG00000007314 | 6329 | PWB-MSC DN | SCN4A | -1.347447 |
| ENSG00000100422 | 64781 | PWB-iPSC DN | CERK | -0.52638 | ENSG00000075407 | 7587 | PWB-EC DN | ZNF37A | -2.666444257 | ENSG00000108556 | 1145 | PWB-MSC DN | CHRNE | -1.351394 |
| ENSG00000144840 | 285282 | PWB-iPSC DN | RABL3 | -0.52691 | ENSG00000183023 | 6546 | PWB-EC DN | SLC8A1 | -2.686857685 | ENSG00000011201 | 3730 | PWB-MSC DN | ANOS1 | -1.354478 |
| ENSG00000144567 | 79137 | PWB-iPSC DN | RETREG2 | -0.52697 | ENSG00000235770 | 646324 | PWB-EC DN | LINC00607 | -2.687604145 | ENSG00000225764 | 101929152 | PWB-MSC DN | P3H2-AS1 | -1.354656 |
| ENSG00000113812 | 93973 | PWB-iPSC DN | ACTR8 | -0.52733 | ENSG00000240184 | 5098 | PWB-EC DN | PCDHGC3 | -2.694383603 | ENSG00000167716 | 124997 | PWB-MSC DN | WDR81 | -1.356267 |
| ENSG00000167193 | 1398 | PWB-iPSC DN | CRK | -0.52791 | ENSG00000183598 | 653604 | PWB-EC DN | H3C13 | -2.736094106 | ENSG00000163357 | 149095 | PWB-MSC DN | DCST1 | -1.359053 |
| ENSG00000159023 | 2035 | PWB-iPSC DN | EPB41 | -0.52804 | ENSG00000276043 | 29128 | PWB-EC DN | UHRF1 | -2.765014863 | ENSG00000212978 | 339803 | PWB-MSC DN | LOC339803 | -1.359145 |
| ENSG00000157191 | 55707 | PWB-iPSC DN | NECAP2 | -0.52805 | ENSG00000173068 | 54796 | PWB-EC DN | BNC2 | -2.770009622 | ENSG00000105825 | 7980 | PWB-MSC DN | TFPI2 | -1.359813 |
| ENSG00000091542 | 54890 | PWB-iPSC DN | ALKBH5 | -0.52903 | ENSG00000167654 | 85300 | PWB-EC DN | ATCAY | -2.774143094 | ENSG00000247416 | 101928847 | PWB-MSC DN | LOC101928847 | -1.359858 |
| ENSG00000111144 | 4048 | PWB-iPSC DN | LTA4H | -0.52945 | ENSG00000249395 | 101805492 | PWB-EC DN | CASC9 | -2.791673203 | ENSG00000271133 | 101927811 | PWB-MSC DN | LOC101927811 | -1.361641 |
| ENSG00000067606 | 5590 | PWB-iPSC DN | PRKCZ | -0.53001 | ENSG00000181649 | 7262 | PWB-EC DN | PHLDA2 | -2.795885005 | ENSG00000184838 | 51334 | PWB-MSC DN | PRR16 | -1.363013 |
| ENSG00000183255 | 754 | PWB-iPSC DN | PTTG1IP | -0.53056 | ENSG00000112599 | 2979 | PWB-EC DN | GUCA1B | -2.801544986 | ENSG00000181804 | 285195 | PWB-MSC DN | SLC9A9 | -1.3653 |
| ENSG00000215301 | 1654 | PWB-iPSC DN | DDX3X | -0.53069 | ENSG00000143768 | 7044 | PWB-EC DN | LEFTY2 | -2.811544363 | ENSG00000105048 | 7138 | PWB-MSC DN | TNNT1 | -1.365959 |
| ENSG00000177733 | 10949 | PWB-iPSC DN | HNRNPA0 | -0.5326 | ENSG00000241749 | 204010 | PWB-EC DN | RPSAP52 | -2.824387655 | ENSG00000182612 | 83882 | PWB-MSC DN | TSPAN10 | -1.375591 |
| ENSG00000117748 | 6118 | PWB-iPSC DN | RPA2 | -0.53318 | ENSG00000116641 | 85440 | PWB-EC DN | DOCK7 | -2.838547666 | ENSG00000171970 | 126295 | PWB-MSC DN | ZNF57 | -1.375908 |
| ENSG00000183624 | 56941 | PWB-iPSC DN | HMCES | -0.53397 | ENSG00000188488 | 5104 | PWB-EC DN | SERPINA5 | -2.841940891 | ENSG00000262655 | 10418 | PWB-MSC DN | SPON1 | -1.385153 |
| ENSG00000124562 | 6631 | PWB-iPSC DN | SNRPC | -0.53412 | ENSG00000171877 | 84978 | PWB-EC DN | FRMD5 | -2.843051472 | ENSG00000204577 | 11025 | PWB-MSC DN | LILRB3 | -1.386237 |
| ENSG00000198960 | 54470 | PWB-iPSC DN | ARMCX6 | -0.53441 | ENSG00000206538 | 389136 | PWB-EC DN | VGLL3 | -2.851315391 | ENSG00000230778 | 100131244 | PWB-MSC DN | ANKRD63 | -1.387094 |
| ENSG00000177383 | 64110 | PWB-iPSC DN | MAGEF1 | -0.53565 | ENSG00000116711 | 5321 | PWB-EC DN | PLA2G4A | -2.85295665 | ENSG00000183607 | 200504 | PWB-MSC DN | GKN2 | -1.392268 |
| ENSG00000102226 | 8237 | PWB-iPSC DN | USP11 | -0.53607 | ENSG00000167771 | 283248 | PWB-EC DN | RCOR2 | -2.856557953 | ENSG00000259073 | 29018 | PWB-MSC DN | FOXN3-AS2 | -1.393236 |
| ENSG00000104081 | 90427 | PWB-iPSC DN | BMF | -0.5385 | ENSG00000143061 | 3321 | PWB-EC DN | IGSF3 | -2.868309683 | ENSG00000003436 | 7035 | PWB-MSC DN | TFPI | -1.404236 |
| ENSG00000039068 | 999 | PWB-iPSC DN | CDH1 | -0.53867 | ENSG00000270885 | 91608 | PWB-EC DN | RASL10B | -2.873049204 | ENSG00000133110 | 10631 | PWB-MSC DN | POSTN | -1.409571 |
| ENSG00000171316 | 55636 | PWB-iPSC DN | CHD7 | -0.53886 | ENSG00000236824 | 618 | PWB-EC DN | BCYRN1 | -2.873572494 | ENSG00000164949 | 2669 | PWB-MSC DN | GEM | -1.410026 |
| ENSG00000097007 | 25 | PWB-iPSC DN | ABL1 | -0.53941 | ENSG00000117069 | 81849 | PWB-EC DN | ST6GALNAC5 | -2.882230609 | ENSG00000166780 | 89927 | PWB-MSC DN | BMERB1 | -1.411966 |
| ENSG00000112514 | 51596 | PWB-iPSC DN | CUTA | -0.53954 | ENSG00000177614 | 79605 | PWB-EC DN | PGBD5 | -2.884372301 | ENSG00000180340 | 2535 | PWB-MSC DN | FZD2 | -1.413307 |
| ENSG00000102858 | 23295 | PWB-iPSC DN | MGRN1 | -0.54013 | ENSG00000100097 | 3956 | PWB-EC DN | LGALS1 | -2.896057576 | ENSG00000108821 | 1277 | PWB-MSC DN | COL1A1 | -1.41927 |
| ENSG00000169679 | 699 | PWB-iPSC DN | BUB1 | -0.5413 | ENSG00000136531 | 6326 | PWB-EC DN | SCN2A | -2.904095624 | ENSG00000183690 | 80258 | PWB-MSC DN | EFHC2 | -1.420555 |
| ENSG00000172273 | 25988 | PWB-iPSC DN | HINFP | -0.54141 | ENSG00000213023 | 84258 | PWB-EC DN | SYT3 | -2.910244156 | ENSG00000140682 | 7041 | PWB-MSC DN | TGFB1I1 | -1.422157 |
| ENSG00000072506 | 3028 | PWB-iPSC DN | HSD17B10 | -0.54171 | ENSG00000164418 | 2898 | PWB-EC DN | GRIK2 | -2.944393849 | ENSG00000187550 | 646643 | PWB-MSC DN | SBK2 | -1.422301 |
| ENSG00000122958 | 9559 | PWB-iPSC DN | VPS26A | -0.54213 | ENSG00000130635 | 1289 | PWB-EC DN | COL5A1 | -2.952699002 | ENSG00000164176 | 10085 | PWB-MSC DN | EDIL3 | -1.425073 |
| ENSG00000101680 | 284217 | PWB-iPSC DN | LAMA1 | -0.54268 | ENSG00000173848 | 10276 | PWB-EC DN | NET1 | -2.956213436 | ENSG00000152931 | 25859 | PWB-MSC DN | PART1 | -1.427539 |
| ENSG00000168496 | 2237 | PWB-iPSC DN | FEN1 | -0.54455 | ENSG00000185634 | 399694 | PWB-EC DN | SHC4 | -2.963203762 | ENSG00000158859 | 9507 | PWB-MSC DN | ADAMTS4 | -1.432901 |
| ENSG00000182512 | 51218 | PWB-iPSC DN | GLRX5 | -0.54469 | ENSG00000140557 | 8128 | PWB-EC DN | ST8SIA2 | -2.965730374 | ENSG00000141854 | 113230 | PWB-MSC DN | MISP3 | -1.434802 |
| ENSG00000035862 | 7077 | PWB-iPSC DN | TIMP2 | -0.54495 | ENSG00000186684 | 339761 | PWB-EC DN | CYP27C1 | -2.976550858 | ENSG00000256576 | 100996246 | PWB-MSC DN | LINC02361 | -1.435996 |
| ENSG00000115234 | 9784 | PWB-iPSC DN | SNX17 | -0.54509 | ENSG00000149256 | 26011 | PWB-EC DN | TENM4 | -2.991808349 | ENSG00000266709 | 84815 | PWB-MSC DN | MGC12916 | -1.436269 |
| ENSG00000205476 | 317762 | PWB-iPSC DN | CCDC85C | -0.54566 | ENSG00000134775 | 80206 | PWB-EC DN | FHOD3 | -2.99321101 | ENSG00000145850 | 91937 | PWB-MSC DN | TIMD4 | -1.44222 |
| ENSG00000151458 | 57182 | PWB-iPSC DN | ANKRD50 | -0.54599 | ENSG00000187688 | 51393 | PWB-EC DN | TRPV2 | -2.998196477 | ENSG00000233806 | 101927289 | PWB-MSC DN | LINC01237 | -1.446283 |
| ENSG00000172057 | 94103 | PWB-iPSC DN | ORMDL3 | -0.54622 | ENSG00000123342 | 4327 | PWB-EC DN | MMP19 | -3.006773997 | ENSG00000158955 | 7484 | PWB-MSC DN | WNT9B | -1.447654 |
| ENSG00000118707 | 60436 | PWB-iPSC DN | TGIF2 | -0.54745 | ENSG00000180730 | 387914 | PWB-EC DN | SHISA2 | -3.011351237 | ENSG00000238098 | 650655 | PWB-MSC DN | ABCA17P | -1.449749 |
| ENSG00000153944 | 124540 | PWB-iPSC DN | MSI2 | -0.54755 | ENSG00000143507 | 11221 | PWB-EC DN | DUSP10 | -3.022912029 | ENSG00000082556 | 4986 | PWB-MSC DN | OPRK1 | -1.451263 |
| ENSG00000174010 | 80311 | PWB-iPSC DN | KLHL15 | -0.54769 | ENSG00000092969 | 7042 | PWB-EC DN | TGFB2 | -3.026517143 | ENSG00000124701 | 10930 | PWB-MSC DN | APOBEC2 | -1.458249 |
| ENSG00000116017 | 1820 | PWB-iPSC DN | ARID3A | -0.54815 | ENSG00000090530 | 55214 | PWB-EC DN | P3H2 | -3.02739397 | ENSG00000163762 | 116441 | PWB-MSC DN | TM4SF18 | -1.460329 |
| ENSG00000014216 | 823 | PWB-iPSC DN | CAPN1 | -0.54824 | ENSG00000154553 | 27295 | PWB-EC DN | PDLIM3 | -3.031302508 | ENSG00000182667 | 50863 | PWB-MSC DN | NTM | -1.462168 |
| ENSG00000144659 | 54977 | PWB-iPSC DN | SLC25A38 | -0.54854 | ENSG00000149380 | 283208 | PWB-EC DN | P4HA3 | -3.048747716 | ENSG00000112394 | 117247 | PWB-MSC DN | SLC16A10 | -1.465298 |
| ENSG00000084693 | 60509 | PWB-iPSC DN | AGBL5 | -0.5493 | ENSG00000151491 | 2059 | PWB-EC DN | EPS8 | -3.062469303 | ENSG00000185634 | 399694 | PWB-MSC DN | SHC4 | -1.465658 |
| ENSG00000079332 | 56681 | PWB-iPSC DN | SAR1A | -0.54946 | ENSG00000163053 | 151473 | PWB-EC DN | SLC16A14 | -3.063074614 | ENSG00000162374 | 1996 | PWB-MSC DN | ELAVL4 | -1.469013 |
| ENSG00000160991 | 80228 | PWB-iPSC DN | ORAI2 | -0.55025 | ENSG00000219438 | 25817 | PWB-EC DN | TAFA5 | -3.074651653 | ENSG00000117707 | 5629 | PWB-MSC DN | PROX1 | -1.481014 |
| ENSG00000116212 | 115353 | PWB-iPSC DN | LRRC42 | -0.55045 | ENSG00000072195 | 10290 | PWB-EC DN | SPEG | -3.076083505 | ENSG00000165181 | 158401 | PWB-MSC DN | SHOC1 | -1.481805 |
| ENSG00000108406 | 79665 | PWB-iPSC DN | DHX40 | -0.55051 | ENSG00000021645 | 9369 | PWB-EC DN | NRXN3 | -3.099508537 | ENSG00000100031 | 2678 | PWB-MSC DN | GGT1 | -1.483582 |
| ENSG00000157227 | 4323 | PWB-iPSC DN | MMP14 | -0.55192 | ENSG00000014914 | 10903 | PWB-EC DN | MTMR11 | -3.104835722 | ENSG00000268738 | 100130086 | PWB-MSC DN | HSFX2 | -1.494601 |
| ENSG00000136436 | 10241 | PWB-iPSC DN | CALCOCO2 | -0.552 | ENSG00000140937 | 1009 | PWB-EC DN | CDH11 | -3.110132394 | ENSG00000196159 | 79633 | PWB-MSC DN | FAT4 | -1.496168 |
| ENSG00000111897 | 57515 | PWB-iPSC DN | SERINC1 | -0.55342 | ENSG00000197457 | 50861 | PWB-EC DN | STMN3 | -3.117761751 | ENSG00000244694 | 442213 | PWB-MSC DN | PTCHD4 | -1.497481 |
| ENSG00000141580 | 56270 | PWB-iPSC DN | WDR45B | -0.55347 | ENSG00000254453 | 100874014 | PWB-EC DN | NAV2-AS2 | -3.128861132 | ENSG00000120500 | 407 | PWB-MSC DN | ARR3 | -1.498025 |
| ENSG00000157954 | 26100 | PWB-iPSC DN | WIPI2 | -0.55495 | ENSG00000185052 | 57419 | PWB-EC DN | SLC24A3 | -3.141898807 | ENSG00000198576 | 23237 | PWB-MSC DN | ARC | -1.49859 |
| ENSG00000120690 | 1997 | PWB-iPSC DN | ELF1 | -0.55521 | ENSG00000104368 | 5327 | PWB-EC DN | PLAT | -3.145619988 | ENSG00000186767 | 139886 | PWB-MSC DN | SPIN4 | -1.498929 |
| ENSG00000117036 | 2117 | PWB-iPSC DN | ETV3 | -0.55558 | ENSG00000105559 | 57664 | PWB-EC DN | PLEKHA4 | -3.181628773 | ENSG00000179776 | 1003 | PWB-MSC DN | CDH5 | -1.499621 |
| ENSG00000174579 | 55167 | PWB-iPSC DN | MSL2 | -0.55672 | ENSG00000106333 | 5118 | PWB-EC DN | PCOLCE | -3.18337543 | ENSG00000153993 | 223117 | PWB-MSC DN | SEMA3D | -1.506086 |
| ENSG00000170779 | 55038 | PWB-iPSC DN | CDCA4 | -0.55684 | ENSG00000128591 | 2318 | PWB-EC DN | FLNC | -3.18988516 | ENSG00000056291 | 10886 | PWB-MSC DN | NPFFR2 | -1.518143 |
| ENSG00000109787 | 51274 | PWB-iPSC DN | KLF3 | -0.55735 | ENSG00000215386 | 388815 | PWB-EC DN | MIR99AHG | -3.202836708 | ENSG00000188747 | 10811 | PWB-MSC DN | NOXA1 | -1.548223 |
| ENSG00000137710 | 5962 | PWB-iPSC DN | RDX | -0.55756 | ENSG00000006016 | 9244 | PWB-EC DN | CRLF1 | -3.213761559 | ENSG00000095370 | 10044 | PWB-MSC DN | SH2D3C | -1.549897 |
| ENSG00000169504 | 25932 | PWB-iPSC DN | CLIC4 | -0.55759 | ENSG00000136010 | 160428 | PWB-EC DN | ALDH1L2 | -3.214976809 | ENSG00000276410 | 3018 | PWB-MSC DN | H2BC3 | -1.550376 |
| ENSG00000156381 | 122416 | PWB-iPSC DN | ANKRD9 | -0.55863 | ENSG00000182272 | 338707 | PWB-EC DN | B4GALNT4 | -3.232263884 | ENSG00000230426 | 104169671 | PWB-MSC DN | LINC01036 | -1.556297 |
| ENSG00000126945 | 3188 | PWB-iPSC DN | HNRNPH2 | -0.56003 | ENSG00000144868 | 66000 | PWB-EC DN | TMEM108 | -3.237260281 | ENSG00000044524 | 2042 | PWB-MSC DN | EPHA3 | -1.561915 |
| ENSG00000044115 | 1495 | PWB-iPSC DN | CTNNA1 | -0.56016 | ENSG00000169439 | 6383 | PWB-EC DN | SDC2 | -3.257500632 | ENSG00000182070 | 23538 | PWB-MSC DN | OR52A1 | -1.562284 |
| ENSG00000006047 | 51087 | PWB-iPSC DN | YBX2 | -0.56061 | ENSG00000004948 | 799 | PWB-EC DN | CALCR | -3.26854673 | ENSG00000187556 | 342977 | PWB-MSC DN | NANOS3 | -1.565556 |
| ENSG00000198324 | 144717 | PWB-iPSC DN | PHETA1 | -0.56168 | ENSG00000128268 | 4248 | PWB-EC DN | MGAT3 | -3.286776952 | ENSG00000145864 | 2561 | PWB-MSC DN | GABRB2 | -1.565585 |
| ENSG00000264364 | 140735 | PWB-iPSC DN | DYNLL2 | -0.56247 | ENSG00000166402 | 7275 | PWB-EC DN | TUB | -3.299204804 | ENSG00000188825 | 100130581 | PWB-MSC DN | LINC00910 | -1.569823 |
| ENSG00000147650 | 29967 | PWB-iPSC DN | LRP12 | -0.56259 | ENSG00000158457 | 340348 | PWB-EC DN | TSPAN33 | -3.308003024 | ENSG00000142552 | 57333 | PWB-MSC DN | RCN3 | -1.571235 |
| ENSG00000148200 | 2649 | PWB-iPSC DN | NR6A1 | -0.56355 | ENSG00000172986 | 727936 | PWB-EC DN | GXYLT2 | -3.308291258 | ENSG00000074660 | 8578 | PWB-MSC DN | SCARF1 | -1.571957 |
| ENSG00000105254 | 1155 | PWB-iPSC DN | TBCB | -0.56487 | ENSG00000164796 | 114788 | PWB-EC DN | CSMD3 | -3.316945547 | ENSG00000013297 | 5010 | PWB-MSC DN | CLDN11 | -1.5761 |
| ENSG00000143514 | 7159 | PWB-iPSC DN | TP53BP2 | -0.56517 | ENSG00000070404 | 10272 | PWB-EC DN | FSTL3 | -3.31800145 | ENSG00000198798 | 4114 | PWB-MSC DN | MAGEB3 | -1.580726 |
| ENSG00000090971 | 57106 | PWB-iPSC DN | NAT14 | -0.56588 | ENSG00000170577 | 10736 | PWB-EC DN | SIX2 | -3.330632626 | ENSG00000236532 | 101927973 | PWB-MSC DN | LINC01695 | -1.586475 |
| ENSG00000105229 | 51588 | PWB-iPSC DN | PIAS4 | -0.56756 | ENSG00000077264 | 5063 | PWB-EC DN | PAK3 | -3.330998338 | ENSG00000092051 | 84502 | PWB-MSC DN | JPH4 | -1.587543 |
| ENSG00000105426 | 5802 | PWB-iPSC DN | PTPRS | -0.56824 | ENSG00000166582 | 201161 | PWB-EC DN | CENPV | -3.339830841 | ENSG00000179598 | 201164 | PWB-MSC DN | PLD6 | -1.591361 |
| ENSG00000067369 | 7158 | PWB-iPSC DN | TP53BP1 | -0.5689 | ENSG00000164484 | 114801 | PWB-EC DN | TMEM200A | -3.342597006 | ENSG00000100351 | 9402 | PWB-MSC DN | GRAP2 | -1.603825 |
| ENSG00000162889 | 9261 | PWB-iPSC DN | MAPKAPK2 | -0.56912 | ENSG00000131435 | 8572 | PWB-EC DN | PDLIM4 | -3.342652179 | ENSG00000164692 | 1278 | PWB-MSC DN | COL1A2 | -1.604363 |
| ENSG00000104774 | 4125 | PWB-iPSC DN | MAN2B1 | -0.56938 | ENSG00000287080 | 8352 | PWB-EC DN | H3C3 | -3.359784146 | ENSG00000142748 | 8547 | PWB-MSC DN | FCN3 | -1.60497 |
| ENSG00000104886 | 55111 | PWB-iPSC DN | PLEKHJ1 | -0.57075 | ENSG00000107779 | 657 | PWB-EC DN | BMPR1A | -3.362778858 | ENSG00000125895 | 55321 | PWB-MSC DN | TMEM74B | -1.605 |

| | | | | | | | | | | | | | | |
|---|---|---|---|---|---|---|---|---|---|---|---|---|---|---|
| ENSG00000124659 | 6903 | PWB-iPSC DN | TBCC | -0.57319 | ENSG00000178187 | 285676 | PWB-EC DN | ZNF454 | -3.368072317 | ENSG00000152430 | 66037 | PWB-MSC DN | BOLL | -1.607713 |
| ENSG00000099804 | 997 | PWB-iPSC DN | CDC34 | -0.57439 | ENSG00000133107 | 7223 | PWB-EC DN | TRPC4 | -3.37087114 | ENSG00000140527 | 56964 | PWB-MSC DN | WDR93 | -1.609437 |
| ENSG00000153044 | 64946 | PWB-iPSC DN | CENPH | -0.57494 | ENSG00000146122 | 23500 | PWB-EC DN | DAAM2 | -3.380671725 | ENSG00000102385 | 1821 | PWB-MSC DN | DRP2 | -1.610441 |
| ENSG00000071564 | 6929 | PWB-iPSC DN | TCF3 | -0.57554 | ENSG00000057019 | 131566 | PWB-EC DN | DCBLD2 | -3.382466249 | ENSG00000229356 | 100861510 | PWB-MSC DN | LRRC3-DT | -1.628643 |
| ENSG00000122779 | 8805 | PWB-iPSC DN | TRIM24 | -0.57573 | ENSG00000156113 | 3778 | PWB-EC DN | KCNMA1 | -3.396278106 | ENSG00000102271 | 56062 | PWB-MSC DN | KLHL4 | -1.631104 |
| ENSG00000076604 | 9618 | PWB-iPSC DN | TRAF4 | -0.57604 | ENSG00000221241 | 692202 | PWB-EC DN | SNORD88A | -3.396918288 | ENSG00000274253 | 283683 | PWB-MSC DN | LOC283683 | -1.632817 |
| ENSG00000181191 | 64219 | PWB-iPSC DN | PJA1 | -0.57704 | ENSG00000172318 | 8708 | PWB-EC DN | B3GALT1 | -3.400957768 | ENSG00000170989 | 1901 | PWB-MSC DN | S1PR1 | -1.635588 |
| ENSG00000048740 | 10659 | PWB-iPSC DN | CELF2 | -0.57817 | ENSG00000071575 | 28951 | PWB-EC DN | TRIB2 | -3.40747958 | ENSG00000171116 | 100506164 | PWB-MSC DN | HSFX1 | -1.638145 |
| ENSG00000140941 | 81631 | PWB-iPSC DN | MAP1LC3B | -0.57854 | ENSG00000213694 | 1903 | PWB-EC DN | S1PR3 | -3.407484764 | ENSG00000168394 | 6890 | PWB-MSC DN | TAP1 | -1.645003 |
| ENSG00000189369 | 23708 | PWB-iPSC DN | GSPT2 | -0.57875 | ENSG00000211448 | 1734 | PWB-EC DN | DIO2 | -3.411424874 | ENSG00000155011 | 27123 | PWB-MSC DN | DKK2 | -1.645619 |
| ENSG00000092036 | 54930 | PWB-iPSC DN | HAUS4 | -0.579 | ENSG00000198205 | 7789 | PWB-EC DN | ZXDA | -3.41425637 | ENSG00000138080 | 11117 | PWB-MSC DN | EMILIN1 | -1.645703 |
| ENSG00000159399 | 3099 | PWB-iPSC DN | HK2 | -0.57927 | ENSG00000148680 | 3363 | PWB-EC DN | HTR7 | -3.424479509 | ENSG00000274333 | 102724219 | PWB-MSC DN | LOC102724219 | -1.646147 |
| ENSG00000152620 | 133686 | PWB-iPSC DN | NADK2 | -0.58052 | ENSG00000227036 | 400619 | PWB-EC DN | LINC00511 | -3.428029461 | ENSG00000254614 | 728975 | PWB-MSC DN | LOC728975 | -1.657207 |
| ENSG00000157404 | 3815 | PWB-iPSC DN | KIT | -0.58059 | ENSG00000167912 | 100505501 | PWB-EC DN | LOC10050550 | -3.45893325 | ENSG00000002745 | 51384 | PWB-MSC DN | WNT16 | -1.657453 |
| ENSG00000162551 | 249 | PWB-iPSC DN | ALPL | -0.58233 | ENSG00000121690 | 91614 | PWB-EC DN | DEPDC7 | -3.460182686 | ENSG00000009694 | 10178 | PWB-MSC DN | TENM1 | -1.657835 |
| ENSG00000162384 | 54987 | PWB-iPSC DN | CZIB | -0.58355 | ENSG00000111704 | 79923 | PWB-EC DN | NANOG | -3.460370064 | ENSG00000198099 | 127 | PWB-MSC DN | ADH4 | -1.661138 |
| ENSG00000116273 | 148479 | PWB-iPSC DN | PHF13 | -0.58391 | ENSG00000165891 | 144455 | PWB-EC DN | E2F7 | -3.468012899 | ENSG00000182916 | 56849 | PWB-MSC DN | TCEAL7 | -1.664872 |
| ENSG00000168268 | 64943 | PWB-iPSC DN | NT5DC2 | -0.58397 | ENSG00000174871 | 254263 | PWB-EC DN | CNIH2 | -3.479022905 | ENSG00000116791 | 1429 | PWB-MSC DN | CRYZ | -1.665986 |
| ENSG00000147044 | 8573 | PWB-iPSC DN | CASK | -0.58397 | ENSG00000162551 | 249 | PWB-EC DN | ALPL | -3.483987044 | ENSG00000105672 | 2116 | PWB-MSC DN | ETV2 | -1.668179 |
| ENSG00000135486 | 3178 | PWB-iPSC DN | HNRNPA1 | -0.58398 | ENSG00000218336 | 55714 | PWB-EC DN | TENM3 | -3.485315744 | ENSG00000273540 | 123624 | PWB-MSC DN | AGBL1 | -1.668332 |
| ENSG00000274286 | 151 | PWB-iPSC DN | ADRA2B | -0.58432 | ENSG00000157322 | 348174 | PWB-EC DN | CLEC18A | -3.48876653 | ENSG00000091592 | 22861 | PWB-MSC DN | NLRP1 | -1.669216 |
| ENSG00000135900 | 65080 | PWB-iPSC DN | MRPL44 | -0.58443 | ENSG00000162576 | 54587 | PWB-EC DN | MXRA8 | -3.49891992 | ENSG00000178623 | 2859 | PWB-MSC DN | GPR35 | -1.670932 |
| ENSG00000104671 | 10671 | PWB-iPSC DN | DCTN6 | -0.58446 | ENSG00000133216 | 2048 | PWB-EC DN | EPHB2 | -3.508672334 | ENSG00000277067 | 102724843 | PWB-MSC DN | LOC102724843 | -1.67434 |
| ENSG00000109685 | 7468 | PWB-iPSC DN | NSD2 | -0.58446 | ENSG00000111087 | 2735 | PWB-EC DN | GLI1 | -3.521695483 | ENSG00000276077 | 102724951 | PWB-MSC DN | LOC102724951 | -1.676133 |
| ENSG00000176619 | 84823 | PWB-iPSC DN | LMNB2 | -0.58496 | ENSG00000180340 | 2535 | PWB-EC DN | FZD2 | -3.524383392 | ENSG00000139508 | 283537 | PWB-MSC DN | SLC46A3 | -1.681563 |
| ENSG00000175130 | 65108 | PWB-iPSC DN | MARCKSL1 | -0.5853 | ENSG00000138134 | 57559 | PWB-EC DN | STAMBPL1 | -3.538425822 | ENSG00000176907 | 56892 | PWB-MSC DN | TCIM | -1.683309 |
| ENSG00000117114 | 23266 | PWB-iPSC DN | ADGRL2 | -0.58562 | ENSG00000179761 | 51268 | PWB-EC DN | PIPOX | -3.539809766 | ENSG00000198796 | 115701 | PWB-MSC DN | ALPK2 | -1.684738 |
| ENSG00000135108 | 23014 | PWB-iPSC DN | FBXO21 | -0.58655 | ENSG00000074416 | 11343 | PWB-EC DN | MGLL | -3.556141284 | ENSG00000028137 | 7133 | PWB-MSC DN | TNFRSF1B | -1.685743 |
| ENSG00000078061 | 369 | PWB-iPSC DN | ARAF | -0.58663 | ENSG00000187634 | 148398 | PWB-EC DN | SAMD11 | -3.559509483 | ENSG00000124588 | 4835 | PWB-MSC DN | NQO2 | -1.690446 |
| ENSG00000126705 | 27245 | PWB-iPSC DN | AHDC1 | -0.58803 | ENSG00000167601 | 558 | PWB-EC DN | AXL | -3.566192608 | ENSG00000204642 | 3134 | PWB-MSC DN | HLA-F | -1.691406 |
| ENSG00000156639 | 60685 | PWB-iPSC DN | ZFAND3 | -0.58813 | ENSG00000268658 | 400680 | PWB-EC DN | LINC00664 | -3.588105214 | ENSG00000130513 | 9518 | PWB-MSC DN | GDF15 | -1.694111 |
| ENSG00000156299 | 7074 | PWB-iPSC DN | TIAM1 | -0.58814 | ENSG00000167157 | 51450 | PWB-EC DN | PRRX2 | -3.595074694 | ENSG00000019991 | 3082 | PWB-MSC DN | HGF | -1.695621 |
| ENSG00000144579 | 58190 | PWB-iPSC DN | CTDSP1 | -0.58846 | ENSG00000164129 | 4889 | PWB-EC DN | NPY5R | -3.598443822 | ENSG00000277991 | 102723360 | PWB-MSC DN | LOC102723360 | -1.697067 |
| ENSG00000184900 | 6612 | PWB-iPSC DN | SUMO3 | -0.59044 | ENSG00000276410 | 3018 | PWB-EC DN | H2BC3 | -3.598588694 | ENSG00000005108 | 221981 | PWB-MSC DN | THSD7A | -1.700819 |
| ENSG00000139880 | 64403 | PWB-iPSC DN | CDH24 | -0.59063 | ENSG00000176490 | 148252 | PWB-EC DN | DIRAS1 | -3.632875416 | ENSG00000135363 | 4005 | PWB-MSC DN | LMO2 | -1.701452 |
| ENSG00000158246 | 115572 | PWB-iPSC DN | TENT5B | -0.59079 | ENSG00000171724 | 57687 | PWB-EC DN | VAT1L | -3.636830387 | ENSG00000151376 | 10873 | PWB-MSC DN | ME3 | -1.704012 |
| ENSG00000072803 | 23291 | PWB-iPSC DN | FBXW11 | -0.5909 | ENSG00000143867 | 130497 | PWB-EC DN | OSR1 | -3.637574674 | ENSG00000163354 | 127579 | PWB-MSC DN | DCST2 | -1.705025 |
| ENSG00000152558 | 114908 | PWB-iPSC DN | TMEM123 | -0.59187 | ENSG00000285294 | 643650 | PWB-EC DN | LINC00842 | -3.680835889 | ENSG00000225756 | 138948 | PWB-MSC DN | DBH-AS1 | -1.705666 |
| ENSG00000114353 | 2771 | PWB-iPSC DN | GNAI2 | -0.59272 | ENSG00000102678 | 2254 | PWB-EC DN | FGF9 | -3.681409175 | ENSG00000154310 | 23043 | PWB-MSC DN | TNIK | -1.708022 |
| ENSG00000171135 | 84522 | PWB-iPSC DN | JAGN1 | -0.5929 | ENSG00000125430 | 9953 | PWB-EC DN | HS3ST3B1 | -3.68455775 | ENSG00000147573 | 84675 | PWB-MSC DN | TRIM55 | -1.7081 |
| ENSG00000032444 | 10908 | PWB-iPSC DN | PNPLA6 | -0.59345 | ENSG00000141576 | 114804 | PWB-EC DN | RNF157 | -3.691235945 | ENSG00000250571 | 2738 | PWB-MSC DN | GLI4 | -1.708329 |
| ENSG00000114270 | 1294 | PWB-iPSC DN | COL7A1 | -0.59353 | ENSG00000168032 | 956 | PWB-EC DN | ENTPD3 | -3.692459951 | ENSG00000134627 | 143689 | PWB-MSC DN | PIWIL4 | -1.710992 |
| ENSG00000103363 | 6923 | PWB-iPSC DN | ELOB | -0.596 | ENSG00000125148 | 4502 | PWB-EC DN | MT2A | -3.698947296 | ENSG00000275496 | 102724701 | PWB-MSC DN | LOC102724701 | -1.713059 |
| ENSG00000173599 | 5091 | PWB-iPSC DN | PC | -0.59739 | ENSG00000114270 | 1294 | PWB-EC DN | COL7A1 | -3.702277528 | ENSG00000157502 | 139221 | PWB-MSC DN | PWWP3B | -1.713716 |
| ENSG00000155893 | 92370 | PWB-iPSC DN | PXYLP1 | -0.59745 | ENSG00000013364 | 9961 | PWB-EC DN | MVP | -3.710232806 | ENSG00000186300 | 148254 | PWB-MSC DN | ZNF555 | -1.718164 |
| ENSG00000030582 | 2896 | PWB-iPSC DN | GRN | -0.59781 | ENSG00000115107 | 55240 | PWB-EC DN | STEAP3 | -3.719554338 | ENSG00000113083 | 4015 | PWB-MSC DN | LOX | -1.719304 |
| ENSG00000177030 | 10522 | PWB-iPSC DN | DEAF1 | -0.59852 | ENSG00000120658 | 55068 | PWB-EC DN | ENOX1 | -3.739756422 | ENSG00000287080 | 8352 | PWB-MSC DN | H3C3 | -1.720199 |
| ENSG00000101384 | 182 | PWB-iPSC DN | JAG1 | -0.60111 | ENSG00000142156 | 1291 | PWB-EC DN | COL6A1 | -3.740329223 | ENSG00000128052 | 3791 | PWB-MSC DN | KDR | -1.724248 |
| ENSG00000196465 | 140465 | PWB-iPSC DN | MYL6B | -0.60145 | ENSG00000189410 | 400745 | PWB-EC DN | SH2D5 | -3.7411746 | ENSG00000165507 | 11067 | PWB-MSC DN | DEPP1 | -1.727909 |
| ENSG00000130584 | 140685 | PWB-iPSC DN | ZBTB46 | -0.60258 | ENSG00000077092 | 5915 | PWB-EC DN | RARB | -3.769528985 | ENSG00000164171 | 3673 | PWB-MSC DN | ITGA2 | -1.731575 |
| ENSG00000130449 | 57688 | PWB-iPSC DN | ZSWIM6 | -0.60307 | ENSG00000266709 | 84815 | PWB-EC DN | MGC12916 | -3.787730642 | ENSG00000049192 | 11174 | PWB-MSC DN | ADAMTS6 | -1.732385 |
| ENSG00000111961 | 23328 | PWB-iPSC DN | SASH1 | -0.60314 | ENSG00000111186 | 81029 | PWB-EC DN | WNT5B | -3.799898734 | ENSG00000231290 | 149773 | PWB-MSC DN | APCDD1L-DT | -1.737864 |
| ENSG00000171843 | 4300 | PWB-iPSC DN | MLLT3 | -0.60344 | ENSG00000231290 | 149773 | PWB-EC DN | APCDD1L-DT | -3.805932975 | ENSG00000258949 | 101927418 | PWB-MSC DN | LOC101927418 | -1.738575 |
| ENSG00000150471 | 23284 | PWB-iPSC DN | ADGRL3 | -0.60347 | ENSG00000148848 | 8038 | PWB-EC DN | ADAM12 | -3.809936267 | ENSG00000152804 | 3087 | PWB-MSC DN | HHEX | -1.739377 |
| ENSG00000005889 | 7543 | PWB-iPSC DN | ZFX | -0.6038 | ENSG00000101134 | 55816 | PWB-EC DN | DOK5 | -3.818660169 | ENSG00000117009 | 8564 | PWB-MSC DN | KMO | -1.741418 |
| ENSG00000106605 | 644 | PWB-iPSC DN | BLVRA | -0.60416 | ENSG00000178776 | 389336 | PWB-EC DN | C5orf46 | -3.823739711 | ENSG00000169297 | 190 | PWB-MSC DN | NR0B1 | -1.743161 |
| ENSG00000134013 | 4017 | PWB-iPSC DN | LOXL2 | -0.60439 | ENSG00000249378 | 401164 | PWB-EC DN | LINC01060 | -3.826705404 | ENSG00000137558 | 51050 | PWB-MSC DN | PI15 | -1.752569 |
| ENSG00000185532 | 5592 | PWB-iPSC DN | PRKG1 | -0.60474 | ENSG00000182752 | 5069 | PWB-EC DN | PAPPA | -3.839940081 | ENSG00000165124 | 79987 | PWB-MSC DN | SVEP1 | -1.75308 |
| ENSG00000167565 | 29946 | PWB-iPSC DN | SERTAD3 | -0.60498 | ENSG00000230778 | 100131244 | PWB-EC DN | ANKRD63 | -3.8468549 | ENSG00000236714 | 101926975 | PWB-MSC DN | LINC01844 | -1.758092 |
| ENSG00000183655 | 64410 | PWB-iPSC DN | KLHL25 | -0.60628 | ENSG00000134602 | 51765 | PWB-EC DN | STK26 | -3.858101951 | ENSG00000113946 | 10686 | PWB-MSC DN | CLDN16 | -1.760125 |
| ENSG00000096696 | 1832 | PWB-iPSC DN | DSP | -0.60659 | ENSG00000196557 | 8912 | PWB-EC DN | CACNA1H | -3.85827022 | ENSG00000167711 | 5345 | PWB-MSC DN | SERPINF2 | -1.763834 |
| ENSG00000164576 | 79685 | PWB-iPSC DN | SAP30L | -0.60739 | ENSG00000233098 | 339260 | PWB-EC DN | LOC339260 | -3.858377297 | ENSG00000151702 | 2313 | PWB-MSC DN | FLI1 | -1.765842 |
| ENSG00000051523 | 1535 | PWB-iPSC DN | CYBA | -0.6075 | ENSG00000148053 | 4915 | PWB-EC DN | NTRK2 | -3.871156931 | ENSG00000152093 | 653275 | PWB-MSC DN | CFC1B | -1.768008 |
| ENSG00000186834 | 10614 | PWB-iPSC DN | HEXIM1 | -0.60818 | ENSG00000181541 | 10586 | PWB-EC DN | MAB21L2 | -3.876227696 | ENSG00000061455 | 93166 | PWB-MSC DN | PRDM6 | -1.769361 |
| ENSG00000197043 | 309 | PWB-iPSC DN | ANXA6 | -0.60835 | ENSG00000162520 | 81493 | PWB-EC DN | SYNC | -3.891725379 | ENSG00000188783 | 5549 | PWB-MSC DN | PRELP | -1.775079 |
| ENSG00000151692 | 9781 | PWB-iPSC DN | RNF144A | -0.61043 | ENSG00000264343 | 388677 | PWB-EC DN | NOTCH2NLA | -3.893068786 | ENSG00000251600 | 101927636 | PWB-MSC DN | LOC101927636 | -1.775517 |
| ENSG00000130479 | 55201 | PWB-iPSC DN | MAP1S | -0.61066 | ENSG00000054356 | 5798 | PWB-EC DN | PTPRN | -3.897843397 | ENSG00000075407 | 7587 | PWB-MSC DN | ZNF37A | -1.777541 |
| ENSG00000135404 | 967 | PWB-iPSC DN | CD63 | -0.61081 | ENSG00000256268 | 105369807 | PWB-EC DN | LINC02454 | -3.899011187 | ENSG00000105605 | 59284 | PWB-MSC DN | CACNG7 | -1.778557 |
| ENSG00000100380 | 6767 | PWB-iPSC DN | ST13 | -0.61118 | ENSG00000034239 | 79645 | PWB-EC DN | EFCAB1 | -3.913333352 | ENSG00000184226 | 5101 | PWB-MSC DN | PCDH9 | -1.786372 |
| ENSG00000106799 | 7046 | PWB-iPSC DN | TGFBR1 | -0.61126 | ENSG00000141052 | 93649 | PWB-EC DN | MYOCD | -3.926028324 | ENSG00000196154 | 6275 | PWB-MSC DN | S100A4 | -1.787952 |
| ENSG00000182307 | 65265 | PWB-iPSC DN | C8orf33 | -0.61154 | ENSG00000116741 | 5997 | PWB-EC DN | RGS2 | -3.934022666 | ENSG00000169169 | 126129 | PWB-MSC DN | CPT1C | -1.79041 |
| ENSG00000140471 | 55180 | PWB-iPSC DN | LINS1 | -0.61223 | ENSG00000196793 | 8187 | PWB-EC DN | ZNF239 | -3.942358043 | ENSG00000204653 | 554235 | PWB-MSC DN | ASPDH | -1.792374 |
| ENSG00000172053 | 5859 | PWB-iPSC DN | QARS1 | -0.61258 | ENSG00000138080 | 11117 | PWB-EC DN | EMILIN1 | -3.954038804 | ENSG00000185760 | 56479 | PWB-MSC DN | KCNQ5 | -1.795053 |
| ENSG00000168566 | 154007 | PWB-iPSC DN | SNRNP48 | -0.61424 | ENSG00000182168 | 8633 | PWB-EC DN | UNC5C | -3.956479429 | ENSG00000100097 | 3956 | PWB-MSC DN | LGALS1 | -1.796132 |
| ENSG00000169635 | 23119 | PWB-iPSC DN | HIC2 | -0.61457 | ENSG00000011465 | 1634 | PWB-EC DN | DCN | -3.963718479 | ENSG00000156253 | 10069 | PWB-MSC DN | RWDD2B | -1.799515 |
| ENSG00000108468 | 10951 | PWB-iPSC DN | CBX1 | -0.61473 | ENSG00000286106 | 101929796 | PWB-EC DN | NOTCH2NLR | -3.971338587 | ENSG00000136698 | 55997 | PWB-MSC DN | CFC1 | -1.802335 |
| ENSG00000167601 | 558 | PWB-iPSC DN | AXL | -0.61493 | ENSG00000185760 | 56479 | PWB-EC DN | KCNQ5 | -3.97792141 | ENSG00000154330 | 5239 | PWB-MSC DN | PGM5 | -1.80477 |
| ENSG00000025039 | 58528 | PWB-iPSC DN | RRAGD | -0.61523 | ENSG00000138696 | 658 | PWB-EC DN | BMPR1B | -3.982130179 | ENSG00000119147 | 84417 | PWB-MSC DN | ECRG4 | -1.813897 |
| ENSG00000145794 | 84466 | PWB-iPSC DN | MEGF10 | -0.61574 | ENSG00000162595 | 9077 | PWB-EC DN | DIRAS3 | -3.988065278 | ENSG00000140287 | 3067 | PWB-MSC DN | HDC | -1.820141 |
| ENSG00000129932 | 83475 | PWB-iPSC DN | DOHH | -0.61618 | ENSG00000164509 | 133396 | PWB-EC DN | IL31RA | -3.994652907 | ENSG00000150687 | 11098 | PWB-MSC DN | PRSS23 | -1.823087 |
| ENSG00000143222 | 51506 | PWB-iPSC DN | UFC1 | -0.61719 | ENSG00000286219 | 100996717 | PWB-EC DN | NOTCH2NLC | -4.002668475 | ENSG00000245694 | 643911 | PWB-MSC DN | CRNDE | -1.826166 |
| ENSG00000162616 | 11080 | PWB-iPSC DN | DNAJB4 | -0.61748 | ENSG00000158164 | 11013 | PWB-EC DN | TMSB15A | -4.002910182 | ENSG00000245060 | 729678 | PWB-MSC DN | LINC00847 | -1.826389 |
| ENSG00000087460 | 2778 | PWB-iPSC DN | GNAS | -0.61755 | ENSG00000147883 | 1030 | PWB-EC DN | CDKN2B | -4.02068253 | ENSG00000104967 | 4858 | PWB-MSC DN | NOVA2 | -1.835412 |
| ENSG00000197321 | 6840 | PWB-iPSC DN | SVIL | -0.61772 | ENSG00000155011 | 27123 | PWB-EC DN | DKK2 | -4.031094434 | ENSG00000183114 | 163933 | PWB-MSC DN | FAM43B | -1.837955 |
| ENSG00000162191 | 51035 | PWB-iPSC DN | UBXN1 | -0.61883 | ENSG00000204011 | 414316 | PWB-EC DN | COL5A1-AS1 | -4.041434638 | ENSG00000026025 | 7431 | PWB-MSC DN | VIM | -1.843957 |
| ENSG00000168556 | 3622 | PWB-iPSC DN | ING2 | -0.61887 | ENSG00000163629 | 5783 | PWB-EC DN | PTPN13 | -4.049813532 | ENSG00000204711 | 138255 | PWB-MSC DN | C9orf135 | -1.849076 |
| ENSG00000103202 | 4833 | PWB-iPSC DN | NME4 | -0.61912 | ENSG00000102452 | 259232 | PWB-EC DN | NALCN | -4.050621874 | ENSG00000237125 | 79804 | PWB-MSC DN | HAND2-AS1 | -1.854679 |
| ENSG00000118298 | 23632 | PWB-iPSC DN | CA14 | -0.61968 | ENSG00000135097 | 4440 | PWB-EC DN | MSI1 | -4.074762008 | ENSG00000125726 | 970 | PWB-MSC DN | CD70 | -1.861144 |
| ENSG00000122026 | 6144 | PWB-iPSC DN | RPL21 | -0.62084 | ENSG00000167653 | 8000 | PWB-EC DN | PSCA | -4.081288426 | ENSG00000152689 | 25780 | PWB-MSC DN | RASGRP3 | -1.865399 |
| ENSG00000130338 | 56995 | PWB-iPSC DN | TULP4 | -0.62148 | ENSG00000082458 | 1741 | PWB-EC DN | DLG3 | -4.094042935 | ENSG00000171587 | 1826 | PWB-MSC DN | DSCAM | -1.86895 |
| ENSG00000163346 | 57326 | PWB-iPSC DN | PBXIP1 | -0.62152 | ENSG00000178773 | 27132 | PWB-EC DN | CPNE7 | -4.099249814 | ENSG00000286106 | 101929796 | PWB-MSC DN | NOTCH2NLR | -1.871808 |
| ENSG00000064961 | 10362 | PWB-iPSC DN | HMG20B | -0.62172 | ENSG00000235602 | 642559 | PWB-EC DN | POU5F1P3 | -4.107346545 | ENSG00000104415 | 8840 | PWB-MSC DN | CCN4 | -1.877526 |
| ENSG00000213190 | 10962 | PWB-iPSC DN | MLLT11 | -0.62193 | ENSG00000148798 | 9118 | PWB-EC DN | INA | -4.111406092 | ENSG00000183715 | 4978 | PWB-MSC DN | OPCML | -1.882922 |
| ENSG00000128268 | 4248 | PWB-iPSC DN | MGAT3 | -0.62201 | ENSG00000072657 | 29953 | PWB-EC DN | TRHDE | -4.111972631 | ENSG00000004799 | 5166 | PWB-MSC DN | PDK4 | -1.885168 |
| ENSG00000116117 | 117583 | PWB-iPSC DN | PARD3B | -0.62204 | ENSG00000131409 | 94030 | PWB-EC DN | LRRC4B | -4.117496114 | ENSG00000171243 | 25928 | PWB-MSC DN | SOSTDC1 | -1.88782 |
| ENSG00000175198 | 5095 | PWB-iPSC DN | PCCA | -0.62207 | ENSG00000168546 | 2675 | PWB-EC DN | GFRA2 | -4.121480082 | ENSG00000225376 | 101928470 | PWB-MSC DN | TMEM246-AS1 | -1.891232 |
| ENSG00000178982 | 27335 | PWB-iPSC DN | EIF3K | -0.62215 | ENSG00000183798 | 90187 | PWB-EC DN | EMILIN3 | -4.131154877 | ENSG00000164220 | 2151 | PWB-MSC DN | F2RL2 | -1.892727 |
| ENSG00000123091 | 26994 | PWB-iPSC DN | RNF11 | -0.62217 | ENSG00000225383 | 207107 | PWB-EC DN | SFTA1P | -4.136077083 | ENSG00000168421 | 399 | PWB-MSC DN | RHOH | -1.917043 |
| ENSG00000099364 | 54620 | PWB-iPSC DN | FBXL19 | -0.62246 | ENSG00000134853 | 5156 | PWB-EC DN | PDGFRA | -4.145552568 | ENSG00000145681 | 1404 | PWB-MSC DN | HAPLN1 | -1.923135 |
| ENSG00000170412 | 55890 | PWB-iPSC DN | GPRC5C | -0.62281 | ENSG00000175832 | 2118 | PWB-EC DN | ETV4 | -4.171138271 | ENSG00000034239 | 79645 | PWB-MSC DN | EFCAB1 | -1.925692 |
| ENSG00000137312 | 10211 | PWB-iPSC DN | FLOT1 | -0.62328 | ENSG00000106484 | 4232 | PWB-EC DN | MEST | -4.17938805 | ENSG00000085514 | 29992 | PWB-MSC DN | PILRA | -1.927627 |
| ENSG00000123472 | 64756 | PWB-iPSC DN | ATPAF1 | -0.6234 | ENSG00000134250 | 4853 | PWB-EC DN | NOTCH2 | -4.183184877 | ENSG00000100095 | 23544 | PWB-MSC DN | SEZ6L | -1.928305 |
| ENSG00000175701 | 205251 | PWB-iPSC DN | MTLN | -0.62446 | ENSG00000286019 | 100996763 | PWB-EC DN | NOTCH2NLB | -4.18910358 | ENSG00000156564 | 57497 | PWB-MSC DN | LRFN2 | -1.952521 |
| ENSG00000102401 | 51566 | PWB-iPSC DN | ARMCX3 | -0.62628 | ENSG00000113739 | 8614 | PWB-EC DN | STC2 | -4.192763724 | ENSG00000283945 | 158035 | PWB-MSC DN | LINC00032 | -1.954834 |
| ENSG00000184602 | 8303 | PWB-iPSC DN | SNN | -0.62727 | ENSG00000159228 | 873 | PWB-EC DN | CBR1 | -4.204385949 | ENSG00000143127 | 8515 | PWB-MSC DN | ITGA10 | -1.958066 |
| ENSG00000132563 | 51308 | PWB-iPSC DN | REEP2 | -0.62733 | ENSG00000153404 | 153478 | PWB-EC DN | PLEKHG4B | -4.222575286 | ENSG00000172969 | 100288801 | PWB-MSC DN | FRG2C | -1.960028 |
| ENSG00000184924 | 391356 | PWB-iPSC DN | PTRHD1 | -0.62852 | ENSG00000104447 | 7227 | PWB-EC DN | TRPS1 | -4.227869943 | ENSG00000157103 | 6529 | PWB-MSC DN | SLC6A1 | -1.961243 |

| | | | | | | | | | | | | | | |
|---|---|---|---|---|---|---|---|---|---|---|---|---|---|---|
| ENSG00000072071 | 22859 | PWB-iPSC DN | ADGRL1 | -0.62907 | ENSG00000223764 | 100130417 | PWB-EC DN | LINC02593 | -4.231129657 | ENSG00000120332 | 63923 | PWB-MSC DN | TNN | -1.966258 |
| ENSG00000178425 | 221294 | PWB-iPSC DN | NT5DC1 | -0.62934 | ENSG00000227053 | 102724094 | PWB-EC DN | MUC12-AS1 | -4.233346113 | ENSG00000170807 | 442721 | PWB-MSC DN | LMOD2 | -1.966896 |
| ENSG00000198034 | 6191 | PWB-iPSC DN | RPS4X | -0.62962 | ENSG00000169715 | 4493 | PWB-EC DN | MT1E | -4.242038746 | ENSG00000079112 | 1015 | PWB-MSC DN | CDH17 | -1.972783 |
| ENSG00000164105 | 8819 | PWB-iPSC DN | SAP30 | -0.63031 | ENSG00000229647 | 105373853 | PWB-EC DN | MYOSLID | -4.250075053 | ENSG00000130612 | 22952 | PWB-MSC DN | CYP2G1P | -1.975859 |
| ENSG00000177096 | 150368 | PWB-iPSC DN | PHETA2 | -0.63057 | ENSG00000134668 | 90853 | PWB-EC DN | SPOCD1 | -4.250878593 | ENSG00000284713 | 107984345 | PWB-MSC DN | SMIM38 | -1.982201 |
| ENSG00000066468 | 2263 | PWB-iPSC DN | FGFR2 | -0.63172 | ENSG00000184985 | 57537 | PWB-EC DN | SORCS2 | -4.259339255 | ENSG00000170180 | 2993 | PWB-MSC DN | GYPA | -1.984426 |
| ENSG00000149679 | 81928 | PWB-iPSC DN | CABLES2 | -0.6337 | ENSG00000249835 | 105379054 | PWB-EC DN | VCAN-AS1 | -4.260081034 | ENSG00000136535 | 10716 | PWB-MSC DN | TBR1 | -1.987529 |
| ENSG00000133639 | 694 | PWB-iPSC DN | BTG1 | -0.63421 | ENSG00000156265 | 56911 | PWB-EC DN | MAP3K7CL | -4.260312088 | ENSG00000127951 | 10875 | PWB-MSC DN | FGL2 | -1.992232 |
| ENSG00000138760 | 950 | PWB-iPSC DN | SCARB2 | -0.63444 | ENSG00000117152 | 5999 | PWB-EC DN | RGS4 | -4.272704916 | ENSG00000172578 | 89857 | PWB-MSC DN | KLHL6 | -1.992991 |
| ENSG00000213753 | 65996 | PWB-iPSC DN | CENPBD1P1 | -0.63514 | ENSG00000144857 | 91653 | PWB-EC DN | BOC | -4.281094101 | ENSG00000196104 | 50859 | PWB-MSC DN | SPOCK3 | -2.002759 |
| ENSG00000117519 | 1266 | PWB-iPSC DN | CNN3 | -0.63555 | ENSG00000276600 | 338382 | PWB-EC DN | RAB7B | -4.293360189 | ENSG00000109099 | 5376 | PWB-MSC DN | PMP22 | -2.006481 |
| ENSG00000176490 | 148252 | PWB-iPSC DN | DIRAS1 | -0.63573 | ENSG00000259721 | 100131315 | PWB-EC DN | LOC10013131 | -4.297740214 | ENSG00000122176 | 2331 | PWB-MSC DN | FMOD | -2.011264 |
| ENSG00000101421 | 128866 | PWB-iPSC DN | CHMP4B | -0.63596 | ENSG00000180155 | 66004 | PWB-EC DN | LYNX1 | -4.306705587 | ENSG00000248858 | 441369 | PWB-MSC DN | FLJ46284 | -2.0128 |
| ENSG00000146833 | 89122 | PWB-iPSC DN | TRIM4 | -0.63657 | ENSG00000137273 | 2295 | PWB-EC DN | FOXF2 | -4.312380018 | ENSG00000183454 | 2903 | PWB-MSC DN | GRIN2A | -2.018378 |
| ENSG00000136943 | 1515 | PWB-iPSC DN | CTSV | -0.63714 | ENSG00000168389 | 84879 | PWB-EC DN | MFSD2A | -4.324598071 | ENSG00000168334 | 165904 | PWB-MSC DN | XIRP1 | -2.025642 |
| ENSG00000141698 | 115024 | PWB-iPSC DN | NT5C3B | -0.63821 | ENSG00000255571 | 254559 | PWB-EC DN | MIR9-3HG | -4.328715535 | ENSG00000166501 | 5579 | PWB-MSC DN | PRKCB | -2.027991 |
| ENSG00000198363 | 444 | PWB-iPSC DN | ASPH | -0.63822 | ENSG00000177234 | 404216 | PWB-EC DN | LINC01561 | -4.331248992 | ENSG00000118997 | 56171 | PWB-MSC DN | DNAH7 | -2.0291 |
| ENSG00000120885 | 1191 | PWB-iPSC DN | CLU | -0.63845 | ENSG00000143333 | 6004 | PWB-EC DN | RGS16 | -4.336160292 | ENSG00000142408 | 59283 | PWB-MSC DN | CACNG8 | -2.032428 |
| ENSG00000088179 | 5775 | PWB-iPSC DN | PTPN4 | -0.63875 | ENSG00000232977 | 100506697 | PWB-EC DN | LINC00327 | -4.346338027 | ENSG00000228824 | 642345 | PWB-MSC DN | MIR4500HG | -2.032796 |
| ENSG00000131914 | 79727 | PWB-iPSC DN | LIN28A | -0.63887 | ENSG00000121440 | 23024 | PWB-EC DN | PDZRN3 | -4.346554146 | ENSG00000162676 | 2672 | PWB-MSC DN | GFI1 | -2.043631 |
| ENSG00000132604 | 7014 | PWB-iPSC DN | TERF2 | -0.63898 | ENSG00000149177 | 5795 | PWB-EC DN | PTPRJ | -4.346701755 | ENSG00000112562 | 64094 | PWB-MSC DN | SMOC2 | -2.050212 |
| ENSG00000168283 | 648 | PWB-iPSC DN | BMI1 | -0.63931 | ENSG00000173320 | 56977 | PWB-EC DN | STOX2 | -4.371397452 | ENSG00000105538 | 54922 | PWB-MSC DN | RASIP1 | -2.055208 |
| ENSG00000188643 | 140576 | PWB-iPSC DN | S100A16 | -0.64127 | ENSG00000204580 | 780 | PWB-EC DN | DDR1 | -4.383985434 | ENSG00000130528 | 3270 | PWB-MSC DN | HRC | -2.058954 |
| ENSG00000158792 | 124044 | PWB-iPSC DN | SPATA2L | -0.64159 | ENSG00000203805 | 196051 | PWB-EC DN | PLPP4 | -4.389926474 | ENSG00000198205 | 7789 | PWB-MSC DN | ZXDA | -2.063473 |
| ENSG00000138175 | 403 | PWB-iPSC DN | ARL3 | -0.64193 | ENSG00000178031 | 92949 | PWB-EC DN | ADAMTSL1 | -4.397070622 | ENSG00000170909 | 126014 | PWB-MSC DN | OSCAR | -2.065605 |
| ENSG00000137726 | 53826 | PWB-iPSC DN | FXYD6 | -0.64229 | ENSG00000187955 | 7373 | PWB-EC DN | COL14A1 | -4.41232413 | ENSG00000185614 | 389119 | PWB-MSC DN | INKA1 | -2.080481 |
| ENSG00000099956 | 6598 | PWB-iPSC DN | SMARCB1 | -0.6426 | ENSG00000151572 | 121601 | PWB-EC DN | ANO4 | -4.423376929 | ENSG00000187323 | 1630 | PWB-MSC DN | DCC | -2.090821 |
| ENSG00000159720 | 9114 | PWB-iPSC DN | ATP6V0D1 | -0.64286 | ENSG00000183762 | 83999 | PWB-EC DN | KREMEN1 | -4.441293594 | ENSG00000228598 | 100874041 | PWB-MSC DN | MACC1-AS1 | -2.094765 |
| ENSG00000132475 | 3021 | PWB-iPSC DN | H3-3B | -0.6434 | ENSG00000067798 | 89795 | PWB-EC DN | NAV3 | -4.457314452 | ENSG00000099869 | 51214 | PWB-MSC DN | IGF2-AS | -2.105074 |
| ENSG00000124299 | 5184 | PWB-iPSC DN | PEPD | -0.64406 | ENSG00000126778 | 6495 | PWB-EC DN | SIX1 | -4.467978832 | ENSG00000258977 | 101928559 | PWB-MSC DN | LINC01467 | -2.12789 |
| ENSG00000099795 | 4713 | PWB-iPSC DN | NDUFB7 | -0.64528 | ENSG00000185274 | 64409 | PWB-EC DN | GALNT17 | -4.474109004 | ENSG00000162645 | 2634 | PWB-MSC DN | GBP2 | -2.132138 |
| ENSG00000184349 | 1946 | PWB-iPSC DN | EFNA5 | -0.64601 | ENSG00000165633 | 196740 | PWB-EC DN | VSTM4 | -4.481612769 | ENSG00000125735 | 8740 | PWB-MSC DN | TNFSF14 | -2.142004 |
| ENSG00000063176 | 56848 | PWB-iPSC DN | SPHK2 | -0.64603 | ENSG00000113578 | 2246 | PWB-EC DN | FGF1 | -4.491336561 | ENSG00000276759 | 100506422 | PWB-MSC DN | LOC100506422 | -2.142699 |
| ENSG00000101849 | 6907 | PWB-iPSC DN | TBL1X | -0.64633 | ENSG00000103257 | 8140 | PWB-EC DN | SLC7A5 | -4.497428634 | ENSG00000131409 | 94030 | PWB-MSC DN | LRRC4B | -2.16227 |
| ENSG00000104765 | 665 | PWB-iPSC DN | BNIP3L | -0.64666 | ENSG00000231574 | 102724550 | PWB-EC DN | LINC02015 | -4.504137485 | ENSG00000162618 | 64123 | PWB-MSC DN | ADGRL4 | -2.165278 |
| ENSG00000116251 | 6146 | PWB-iPSC DN | RPL22 | -0.64852 | ENSG00000082397 | 23136 | PWB-EC DN | EPB41L3 | -4.505974597 | ENSG00000172250 | 94009 | PWB-MSC DN | SERHL | -2.18614 |
| ENSG00000111058 | 79611 | PWB-iPSC DN | ACSS3 | -0.6489 | ENSG00000153976 | 9955 | PWB-EC DN | HS3ST3A1 | -4.512178034 | ENSG00000102359 | 27286 | PWB-MSC DN | SRPX2 | -2.194524 |
| ENSG00000126934 | 5605 | PWB-iPSC DN | MAP2K2 | -0.64916 | ENSG00000063660 | 2817 | PWB-EC DN | GPC1 | -4.528478492 | ENSG00000168542 | 1281 | PWB-MSC DN | COL3A1 | -2.196567 |
| ENSG00000107341 | 54926 | PWB-iPSC DN | UBE2R2 | -0.64916 | ENSG00000235034 | 342918 | PWB-EC DN | C19orf81 | -4.534872939 | ENSG00000244541 | 101927992 | PWB-MSC DN | LINC01213 | -2.200424 |
| ENSG00000170310 | 9482 | PWB-iPSC DN | STX8 | -0.64976 | ENSG00000132821 | 128434 | PWB-EC DN | VSTM2L | -4.541281859 | ENSG00000162944 | 130132 | PWB-MSC DN | RFTN2 | -2.204056 |
| ENSG00000147604 | 6129 | PWB-iPSC DN | RPL7 | -0.6498 | ENSG00000108823 | 6442 | PWB-EC DN | SGCA | -4.542500402 | ENSG00000112936 | 730 | PWB-MSC DN | C7 | -2.207587 |
| ENSG00000119801 | 51646 | PWB-iPSC DN | YPEL5 | -0.6501 | ENSG00000166106 | 170689 | PWB-EC DN | ADAMTS15 | -4.544614251 | ENSG00000164116 | 2982 | PWB-MSC DN | GUCY1A1 | -2.211181 |
| ENSG00000145736 | 2966 | PWB-iPSC DN | GTF2H2 | -0.65039 | ENSG00000162545 | 55450 | PWB-EC DN | CAMK2N1 | -4.545499199 | ENSG00000130226 | 1804 | PWB-MSC DN | DPP6 | -2.220331 |
| ENSG00000127955 | 2770 | PWB-iPSC DN | GNAI1 | -0.65047 | ENSG00000159216 | 861 | PWB-EC DN | RUNX1 | -4.559766653 | ENSG00000176692 | 2303 | PWB-MSC DN | FOXC2 | -2.222001 |
| ENSG00000023171 | 57476 | PWB-iPSC DN | GRAMD1B | -0.65117 | ENSG00000075213 | 10371 | PWB-EC DN | SEMA3A | -4.56420358 | ENSG00000182492 | 633 | PWB-MSC DN | BGN | -2.223006 |
| ENSG00000147912 | 26267 | PWB-iPSC DN | FBXO10 | -0.65174 | ENSG00000113083 | 4015 | PWB-EC DN | LOX | -4.564945376 | ENSG00000134215 | 10451 | PWB-MSC DN | VAV3 | -2.227104 |
| ENSG00000133069 | 9911 | PWB-iPSC DN | TMCC2 | -0.65259 | ENSG00000177335 | 286122 | PWB-EC DN | C8orf31 | -4.572251799 | ENSG00000125804 | 284800 | PWB-MSC DN | FAM182A | -2.233547 |
| ENSG00000088882 | 56265 | PWB-iPSC DN | CPXM1 | -0.65293 | ENSG00000115844 | 1746 | PWB-EC DN | DLX2 | -4.580204664 | ENSG00000250821 | 646145 | PWB-MSC DN | EXOC1L | -2.242353 |
| ENSG00000092445 | 7301 | PWB-iPSC DN | TYRO3 | -0.65482 | ENSG00000174332 | 148979 | PWB-EC DN | GLIS1 | -4.582792079 | ENSG00000205502 | 388125 | PWB-MSC DN | C2CD4B | -2.249581 |
| ENSG00000004059 | 381 | PWB-iPSC DN | ARF5 | -0.65486 | ENSG00000128510 | 51200 | PWB-EC DN | CPA4 | -4.586036316 | ENSG00000162782 | 163589 | PWB-MSC DN | TDRD5 | -2.249727 |
| ENSG00000160993 | 54784 | PWB-iPSC DN | ALKBH4 | -0.65519 | ENSG00000181638 | 286128 | PWB-EC DN | ZFP41 | -4.588873555 | ENSG00000188828 | 441509 | PWB-MSC DN | GLRA4 | -2.258445 |
| ENSG00000149179 | 79096 | PWB-iPSC DN | C11orf49 | -0.65601 | ENSG00000138795 | 51176 | PWB-EC DN | LEF1 | -4.595168758 | ENSG00000213023 | 84258 | PWB-MSC DN | SYT3 | -2.265087 |
| ENSG00000168036 | 1499 | PWB-iPSC DN | CTNNB1 | -0.65657 | ENSG00000018236 | 1272 | PWB-EC DN | CNTN1 | -4.606272821 | ENSG00000140945 | 1012 | PWB-MSC DN | CDH13 | -2.266069 |
| ENSG00000196531 | 4666 | PWB-iPSC DN | NACA | -0.65695 | ENSG00000146648 | 1956 | PWB-EC DN | EGFR | -4.63618812 | ENSG00000115008 | 3552 | PWB-MSC DN | IL1A | -2.266103 |
| ENSG00000229117 | 6171 | PWB-iPSC DN | RPL41 | -0.6573 | ENSG00000164199 | 84059 | PWB-EC DN | ADGRV1 | -4.642483919 | ENSG00000171056 | 83595 | PWB-MSC DN | SOX7 | -2.26707 |
| ENSG00000143878 | 388 | PWB-iPSC DN | RHOB | -0.65874 | ENSG00000091656 | 79776 | PWB-EC DN | ZFHX4 | -4.642655236 | ENSG00000125657 | 8744 | PWB-MSC DN | TNFSF9 | -2.283731 |
| ENSG00000213626 | 81606 | PWB-iPSC DN | LBH | -0.65918 | ENSG00000101311 | 55612 | PWB-EC DN | FERMT1 | -4.654138964 | ENSG00000164241 | 401207 | PWB-MSC DN | C5orf63 | -2.285701 |
| ENSG00000144749 | 26018 | PWB-iPSC DN | LRIG1 | -0.65929 | ENSG00000026559 | 3755 | PWB-EC DN | KCNG1 | -4.664670775 | ENSG00000204179 | 26095 | PWB-MSC DN | PTPN20 | -2.288805 |
| ENSG00000169169 | 126129 | PWB-iPSC DN | CPT1C | -0.66036 | ENSG00000134463 | 79746 | PWB-EC DN | ECHDC3 | -4.681208823 | ENSG00000136826 | 9314 | PWB-MSC DN | KLF4 | -2.290667 |
| ENSG00000109794 | 25854 | PWB-iPSC DN | FAM149A | -0.66067 | ENSG00000101210 | 1917 | PWB-EC DN | EEF1A2 | -4.683843313 | ENSG00000235890 | 54082 | PWB-MSC DN | TSPEAR-AS1 | -2.29219 |
| ENSG00000115457 | 3485 | PWB-iPSC DN | IGFBP2 | -0.66088 | ENSG00000235890 | 54082 | PWB-EC DN | TSPEAR-AS1 | -4.692461806 | ENSG00000118407 | 27145 | PWB-MSC DN | FILIP1 | -2.32606 |
| ENSG00000128203 | 57168 | PWB-iPSC DN | ASPHD2 | -0.66088 | ENSG00000128266 | 2781 | PWB-EC DN | GNAZ | -4.701564139 | ENSG00000166033 | 5654 | PWB-MSC DN | HTRA1 | -2.326174 |
| ENSG00000106829 | 7091 | PWB-iPSC DN | TLE4 | -0.66096 | ENSG00000122862 | 5552 | PWB-EC DN | SRGN | -4.709045367 | ENSG00000164483 | 154075 | PWB-MSC DN | SAMD3 | -2.336076 |
| ENSG00000130313 | 25796 | PWB-iPSC DN | PGLS | -0.66351 | ENSG00000106278 | 5803 | PWB-EC DN | PTPRZ1 | -4.710731106 | ENSG00000172137 | 794 | PWB-MSC DN | CALB2 | -2.348451 |
| ENSG00000154229 | 5578 | PWB-iPSC DN | PRKCA | -0.6653 | ENSG00000251493 | 2297 | PWB-EC DN | FOXD1 | -4.732221387 | ENSG00000127241 | 5648 | PWB-MSC DN | MASP1 | -2.352121 |
| ENSG00000164366 | 133957 | PWB-iPSC DN | CCDC127 | -0.66547 | ENSG00000079931 | 26002 | PWB-EC DN | MOXD1 | -4.733544811 | ENSG00000164107 | 9464 | PWB-MSC DN | HAND2 | -2.369245 |
| ENSG00000075043 | 3785 | PWB-iPSC DN | KCNQ2 | -0.66795 | ENSG00000198598 | 4326 | PWB-EC DN | MMP17 | -4.74676403 | ENSG00000171388 | 8862 | PWB-MSC DN | APLN | -2.370648 |
| ENSG00000014641 | 4190 | PWB-iPSC DN | MDH1 | -0.66851 | ENSG00000186297 | 2558 | PWB-EC DN | GABRA5 | -4.749063799 | ENSG00000134463 | 79746 | PWB-MSC DN | ECHDC3 | -2.372008 |
| ENSG00000160789 | 4000 | PWB-iPSC DN | LMNA | -0.66879 | ENSG00000233608 | 117581 | PWB-EC DN | TWIST2 | -4.758334444 | ENSG00000174429 | 137735 | PWB-MSC DN | ABRA | -2.392343 |
| ENSG00000169855 | 6091 | PWB-iPSC DN | ROBO1 | -0.6699 | ENSG00000163520 | 2199 | PWB-EC DN | FBLN2 | -4.771291716 | ENSG00000250361 | 2994 | PWB-MSC DN | GYPB | -2.395588 |
| ENSG00000205903 | 100131017 | PWB-iPSC DN | ZNF316 | -0.67093 | ENSG00000184838 | 51334 | PWB-EC DN | PRR16 | -4.787616455 | ENSG00000130649 | 1571 | PWB-MSC DN | CYP2E1 | -2.410442 |
| ENSG00000146112 | 170954 | PWB-iPSC DN | PPP1R18 | -0.67133 | ENSG00000162733 | 4921 | PWB-EC DN | DDR2 | -4.78848426 | ENSG00000108622 | 3384 | PWB-MSC DN | ICAM2 | -2.413419 |
| ENSG00000140931 | 123920 | PWB-iPSC DN | CMTM3 | -0.67145 | ENSG00000253414 | 100507420 | PWB-EC DN | LINC01605 | -4.795204575 | ENSG00000177464 | 2828 | PWB-MSC DN | GPR4 | -2.419289 |
| ENSG00000197565 | 1288 | PWB-iPSC DN | COL4A6 | -0.67187 | ENSG00000038427 | 1462 | PWB-EC DN | VCAN | -4.797679639 | ENSG00000270604 | 414778 | PWB-MSC DN | HCG17 | -2.422503 |
| ENSG00000141858 | 90378 | PWB-iPSC DN | SAMD1 | -0.67189 | ENSG00000139329 | 4060 | PWB-EC DN | LUM | -4.805833946 | ENSG00000196604 | 728378 | PWB-MSC DN | POTEF | -2.430214 |
| ENSG00000086848 | 79796 | PWB-iPSC DN | ALG9 | -0.67243 | ENSG00000174343 | 55584 | PWB-EC DN | CHRNA9 | -4.806662228 | ENSG00000230461 | 100505832 | PWB-MSC DN | PROX1-AS1 | -2.467017 |
| ENSG00000165804 | 51222 | PWB-iPSC DN | ZNF219 | -0.67493 | ENSG00000134363 | 10468 | PWB-EC DN | FST | -4.809717201 | ENSG00000161940 | 255877 | PWB-MSC DN | BCL6B | -2.467427 |
| ENSG00000129245 | 9513 | PWB-iPSC DN | FXR2 | -0.67499 | ENSG00000188760 | 130612 | PWB-EC DN | TMEM198 | -4.810769135 | ENSG00000157554 | 2078 | PWB-MSC DN | ERG | -2.467708 |
| ENSG00000154803 | 201163 | PWB-iPSC DN | FLCN | -0.6755 | ENSG00000104435 | 11075 | PWB-EC DN | STMN2 | -4.818232406 | ENSG00000248099 | 3640 | PWB-MSC DN | INSL3 | -2.483129 |
| ENSG00000178184 | 84552 | PWB-iPSC DN | PARD6G | -0.67578 | ENSG00000198597 | 9745 | PWB-EC DN | ZNF536 | -4.821862043 | ENSG00000164122 | 140458 | PWB-MSC DN | ASB5 | -2.492688 |
| ENSG00000130717 | 83549 | PWB-iPSC DN | UCK1 | -0.67602 | ENSG00000165061 | 79698 | PWB-EC DN | ZMAT4 | -4.828962955 | ENSG00000141469 | 6563 | PWB-MSC DN | SLC14A1 | -2.497104 |
| ENSG00000115415 | 6772 | PWB-iPSC DN | STAT1 | -0.67702 | ENSG00000117228 | 2633 | PWB-EC DN | GBP1 | -4.836124243 | ENSG00000152785 | 651 | PWB-MSC DN | BMP3 | -2.504752 |
| ENSG00000175203 | 10540 | PWB-iPSC DN | DCTN2 | -0.67851 | ENSG00000176842 | 10265 | PWB-EC DN | IRX5 | -4.850277867 | ENSG00000183569 | 253190 | PWB-MSC DN | SERHL2 | -2.513327 |
| ENSG00000175550 | 10589 | PWB-iPSC DN | DRAP1 | -0.67889 | ENSG00000187193 | 4501 | PWB-EC DN | MT1X | -4.850939646 | ENSG00000197465 | 2996 | PWB-MSC DN | GYPE | -2.528175 |
| ENSG00000143147 | 23432 | PWB-iPSC DN | GPR161 | -0.67896 | ENSG00000264717 | 100996758 | PWB-EC DN | NPY4R2 | -4.852163142 | ENSG00000077943 | 8516 | PWB-MSC DN | ITGA8 | -2.537931 |
| ENSG00000116791 | 1429 | PWB-iPSC DN | CRYZ | -0.68099 | ENSG00000124813 | 860 | PWB-EC DN | RUNX2 | -4.854696182 | ENSG00000107984 | 22943 | PWB-MSC DN | DKK1 | -2.567676 |
| ENSG00000057935 | 57504 | PWB-iPSC DN | MTA3 | -0.68218 | ENSG00000105419 | 56917 | PWB-EC DN | MEIS3 | -4.863181092 | ENSG00000109205 | 54959 | PWB-MSC DN | ODAM | -2.582944 |
| ENSG00000080709 | 3781 | PWB-iPSC DN | KCNN2 | -0.68223 | ENSG00000134569 | 4038 | PWB-EC DN | LRP4 | -4.891323494 | ENSG00000101384 | 182 | PWB-MSC DN | JAG1 | -2.607308 |
| ENSG00000145687 | 23635 | PWB-iPSC DN | SSBP2 | -0.68266 | ENSG00000154188 | 284 | PWB-EC DN | ANGPT1 | -4.895615444 | ENSG00000152207 | 57105 | PWB-MSC DN | CYSLTR2 | -2.616372 |
| ENSG00000140332 | 7090 | PWB-iPSC DN | TLE3 | -0.68266 | ENSG00000139278 | 11010 | PWB-EC DN | GLIPR1 | -4.904079216 | ENSG00000197046 | 284266 | PWB-MSC DN | SIGLEC15 | -2.624819 |
| ENSG00000163682 | 6133 | PWB-iPSC DN | RPL9 | -0.68284 | ENSG00000138395 | 65061 | PWB-EC DN | CDK15 | -4.90552438 | ENSG00000276289 | 102723475 | PWB-MSC DN | KCNE1B | -2.63852 |
| ENSG00000111266 | 80824 | PWB-iPSC DN | DUSP16 | -0.68329 | ENSG00000206190 | 57194 | PWB-EC DN | ATP10A | -4.908023313 | ENSG00000259134 | 145820 | PWB-MSC DN | LINC00924 | -2.652895 |
| ENSG00000089876 | 55760 | PWB-iPSC DN | DHX32 | -0.68409 | ENSG00000163710 | 26577 | PWB-EC DN | PCOLCE2 | -4.918737296 | ENSG00000164106 | 11341 | PWB-MSC DN | SCRG1 | -2.667409 |
| ENSG00000105373 | 29997 | PWB-iPSC DN | NOP53 | -0.68446 | ENSG00000164932 | 115908 | PWB-EC DN | CTHRC1 | -4.927578819 | ENSG00000125730 | 718 | PWB-MSC DN | C3 | -2.668831 |
| ENSG00000116584 | 9181 | PWB-iPSC DN | ARHGEF2 | -0.68464 | ENSG00000269994 | 440173 | PWB-EC DN | LOC440173 | -4.929333839 | ENSG00000167617 | 148170 | PWB-MSC DN | CDC42EP5 | -2.67242 |
| ENSG00000186918 | 55893 | PWB-iPSC DN | ZNF395 | -0.6862 | ENSG00000019549 | 6591 | PWB-EC DN | SNAI2 | -4.951415068 | ENSG00000205678 | 253017 | PWB-MSC DN | TECRL | -2.691847 |
| ENSG00000117632 | 3925 | PWB-iPSC DN | STMN1 | -0.68621 | ENSG00000240771 | 115557 | PWB-EC DN | ARHGEF25 | -4.959107704 | ENSG00000162039 | 254528 | PWB-MSC DN | MEIOB | -2.695605 |
| ENSG00000114544 | 54946 | PWB-iPSC DN | SLC41A3 | -0.68635 | ENSG00000078098 | 2191 | PWB-EC DN | FAP | -5.00491716 | ENSG00000283196 | 107985911 | PWB-MSC DN | LOC107985911 | -2.699257 |
| ENSG00000182054 | 3418 | PWB-iPSC DN | IDH2 | -0.68706 | ENSG00000100985 | 4318 | PWB-EC DN | MMP9 | -5.007648795 | ENSG00000205856 | 150297 | PWB-MSC DN | C22orf42 | -2.706291 |
| ENSG00000164684 | 619279 | PWB-iPSC DN | ZNF704 | -0.68767 | ENSG00000123989 | 79586 | PWB-EC DN | CHPF | -5.045615961 | ENSG00000162511 | 7805 | PWB-MSC DN | LAPTM5 | -2.738455 |
| ENSG00000181588 | 399664 | PWB-iPSC DN | MEX3D | -0.68788 | ENSG00000138061 | 1545 | PWB-EC DN | CYP1B1 | -5.046605884 | ENSG00000092969 | 7042 | PWB-MSC DN | TGFB2 | -2.738724 |
| ENSG00000100554 | 51382 | PWB-iPSC DN | ATP6V1D | -0.69055 | ENSG00000212724 | 730755 | PWB-EC DN | KRTAP2-3 | -5.070011338 | ENSG00000247774 | 100233209 | PWB-MSC DN | PCED1B-AS1 | -2.749412 |
| ENSG00000096717 | 23411 | PWB-iPSC DN | SIRT1 | -0.69151 | ENSG00000250366 | 100507043 | PWB-EC DN | TUNAR | -5.072397108 | ENSG00000175445 | 4023 | PWB-MSC DN | LPL | -2.754968 |
| ENSG00000147010 | 30011 | PWB-iPSC DN | SH3KBP1 | -0.69297 | ENSG00000103196 | 83716 | PWB-EC DN | CRISPLD2 | -5.072864872 | ENSG00000182870 | 50614 | PWB-MSC DN | GALNT9 | -2.758666 |
| ENSG00000164258 | 4724 | PWB-iPSC DN | NDUFS4 | -0.69355 | ENSG00000164949 | 2669 | PWB-EC DN | GEM | -5.087751169 | ENSG00000106034 | 79974 | PWB-MSC DN | CPED1 | -2.763185 |
| ENSG00000265241 | 9939 | PWB-iPSC DN | RBM8A | -0.69402 | ENSG00000168874 | 84913 | PWB-EC DN | ATOH8 | -5.097726382 | ENSG00000118733 | 118427 | PWB-MSC DN | OLFM3 | -2.764788 |
| ENSG00000171298 | 2548 | PWB-iPSC DN | GAA | -0.69434 | ENSG00000158008 | 2134 | PWB-EC DN | EXTL1 | -5.098385117 | ENSG00000155307 | 64092 | PWB-MSC DN | SAMSN1 | -2.765105 |

| | | | | | | | | | | | | | | | | |
|---|---|---|---|---|---|---|---|---|---|---|---|---|---|---|---|---|
| ENSG00000077264 | 5063 | PWB-iPSC DN | PAK3 | -0.69489 | | ENSG00000117586 | 7292 | PWB-EC DN | TNFSF4 | -5.12294738 | | ENSG00000159251 | 70 | PWB-MSC DN | ACTC1 | -2.782403 |
| ENSG00000180921 | 286077 | PWB-iPSC DN | FAM83H | -0.69601 | | ENSG00000248905 | 342184 | PWB-EC DN | FMN1 | -5.125950639 | | ENSG00000189108 | 26280 | PWB-MSC DN | IL1RAPL2 | -2.79713 |
| ENSG00000138829 | 2201 | PWB-iPSC DN | FBN2 | -0.69688 | | ENSG00000149451 | 80332 | PWB-EC DN | ADAM33 | -5.13142734 | | ENSG00000162367 | 6886 | PWB-MSC DN | TAL1 | -2.799682 |
| ENSG00000116922 | 54955 | PWB-iPSC DN | C1orf109 | -0.69725 | | ENSG00000075043 | 3785 | PWB-EC DN | KCNQ2 | -5.135933095 | | ENSG00000187037 | 353345 | PWB-MSC DN | GPR141 | -2.809019 |
| ENSG00000273136 | 101060684 | PWB-iPSC DN | NBPF26 | -0.69748 | | ENSG00000166250 | 79827 | PWB-EC DN | CLMP | -5.146086787 | | ENSG00000064989 | 10203 | PWB-MSC DN | CALCRL | -2.810045 |
| ENSG00000079462 | 5050 | PWB-iPSC DN | PAFAH1B3 | -0.69813 | | ENSG00000185942 | 286183 | PWB-EC DN | NKAIN3 | -5.152015558 | | ENSG00000130300 | 83483 | PWB-MSC DN | PLVAP | -2.827194 |
| ENSG00000158158 | 26504 | PWB-iPSC DN | CNNM4 | -0.69921 | | ENSG00000137691 | 85016 | PWB-EC DN | CFAP300 | -5.156893336 | | ENSG00000284837 | 339166 | PWB-MSC DN | LOC339166 | -2.837167 |
| ENSG00000170889 | 6203 | PWB-iPSC DN | RPS9 | -0.6997 | | ENSG00000145423 | 6423 | PWB-EC DN | SFRP2 | -5.165423585 | | ENSG00000125810 | 22918 | PWB-MSC DN | CD93 | -2.868511 |
| ENSG00000236104 | 9278 | PWB-iPSC DN | ZBTB22 | -0.69973 | | ENSG00000133083 | 9201 | PWB-EC DN | DCLK1 | -5.172371624 | | ENSG00000120457 | 3762 | PWB-MSC DN | KCNJ5 | -2.878084 |
| ENSG00000155959 | 7411 | PWB-iPSC DN | VBP1 | -0.70043 | | ENSG00000166922 | 6447 | PWB-EC DN | SCG5 | -5.186605781 | | ENSG00000119917 | 3437 | PWB-MSC DN | IFIT3 | -2.896877 |
| ENSG00000092010 | 5720 | PWB-iPSC DN | PSME1 | -0.70111 | | ENSG00000156466 | 392255 | PWB-EC DN | GDF6 | -5.187322396 | | ENSG00000248673 | 104310351 | PWB-MSC DN | LINC01331 | -2.914643 |
| ENSG00000146232 | 4794 | PWB-iPSC DN | NFKBIE | -0.70226 | | ENSG00000114923 | 6508 | PWB-EC DN | SLC4A3 | -5.194407741 | | ENSG00000266200 | 5408 | PWB-MSC DN | PNLIPRP2 | -2.942902 |
| ENSG00000143951 | 51057 | PWB-iPSC DN | WDPCP | -0.70405 | | ENSG00000105894 | 5764 | PWB-EC DN | PTN | -5.243560564 | | ENSG00000184374 | 10584 | PWB-MSC DN | COLEC10 | -2.94427 |
| ENSG00000063241 | 79763 | PWB-iPSC DN | ISOC2 | -0.70443 | | ENSG00000111424 | 7421 | PWB-EC DN | VDR | -5.243810834 | | ENSG00000249343 | 101929082 | PWB-MSC DN | LINC01333 | -2.955002 |
| ENSG00000020256 | 55734 | PWB-iPSC DN | ZFP64 | -0.70539 | | ENSG00000123610 | 7130 | PWB-EC DN | TNFAIP6 | -5.2520072 | | ENSG00000235034 | 342918 | PWB-MSC DN | C19orf81 | -2.966043 |
| ENSG00000128039 | 79644 | PWB-iPSC DN | SRD5A3 | -0.70549 | | ENSG00000168243 | 2786 | PWB-EC DN | GNG4 | -5.258577418 | | ENSG00000126785 | 57381 | PWB-MSC DN | RHOJ | -2.972529 |
| ENSG00000102265 | 7076 | PWB-iPSC DN | TIMP1 | -0.70561 | | ENSG00000102385 | 1821 | PWB-EC DN | DRP2 | -5.260532135 | | ENSG00000169908 | 4071 | PWB-MSC DN | TM4SF1 | -2.98128 |
| ENSG00000115109 | 57669 | PWB-iPSC DN | EPB41L5 | -0.707 | | ENSG00000205212 | 339184 | PWB-EC DN | CCDC144NL | -5.263022517 | | ENSG00000100362 | 5816 | PWB-MSC DN | PVALB | -2.988561 |
| ENSG00000130939 | 10277 | PWB-iPSC DN | UBE4B | -0.70727 | | ENSG00000170961 | 3037 | PWB-EC DN | HAS2 | -5.263813845 | | ENSG00000167800 | 347853 | PWB-MSC DN | TBX10 | -3.046389 |
| ENSG00000108953 | 7531 | PWB-iPSC DN | YWHAE | -0.70761 | | ENSG00000151388 | 81792 | PWB-EC DN | ADAMTS12 | -5.28894737 | | ENSG00000286058 | 440905 | PWB-MSC DN | FAR2P1 | -3.056556 |
| ENSG00000174748 | 6138 | PWB-iPSC DN | RPL15 | -0.71009 | | ENSG00000137868 | 64220 | PWB-EC DN | STRA6 | -5.30065004 | | ENSG00000168702 | 53353 | PWB-MSC DN | LRP1B | -3.058964 |
| ENSG00000215305 | 64601 | PWB-iPSC DN | VPS16 | -0.71048 | | ENSG00000127863 | 55504 | PWB-EC DN | TNFRSF19 | -5.301219122 | | ENSG00000161681 | 50944 | PWB-MSC DN | SHANK1 | -3.07807 |
| ENSG00000114115 | 5947 | PWB-iPSC DN | RBP1 | -0.71122 | | ENSG00000069188 | 54549 | PWB-EC DN | SDK2 | -5.30461261 | | ENSG00000136869 | 7099 | PWB-MSC DN | TLR4 | -3.115876 |
| ENSG00000130545 | 92359 | PWB-iPSC DN | CRB3 | -0.71173 | | ENSG00000087494 | 5744 | PWB-EC DN | PTHLH | -5.327437814 | | ENSG00000069122 | 221395 | PWB-MSC DN | ADGRF5 | -3.123968 |
| ENSG00000111052 | 8825 | PWB-iPSC DN | LIN7A | -0.71255 | | ENSG00000198846 | 9760 | PWB-EC DN | TOX | -5.33488236 | | ENSG00000203878 | 149620 | PWB-MSC DN | CHIAP2 | -3.197815 |
| ENSG00000105204 | 9149 | PWB-iPSC DN | DYRK1B | -0.71392 | | ENSG00000163017 | 72 | PWB-EC DN | ACTG2 | -5.336909678 | | ENSG00000163217 | 27302 | PWB-MSC DN | BMP10 | -3.209681 |
| ENSG00000140807 | 85407 | PWB-iPSC DN | NKD1 | -0.71425 | | ENSG00000074047 | 2736 | PWB-EC DN | GLI2 | -5.355661772 | | ENSG00000128253 | 10739 | PWB-MSC DN | RFPL2 | -3.261398 |
| ENSG00000140391 | 10099 | PWB-iPSC DN | TSPAN3 | -0.71434 | | ENSG00000140807 | 85407 | PWB-EC DN | NKD1 | -5.370463187 | | ENSG00000157680 | 9162 | PWB-MSC DN | DGKI | -3.265418 |
| ENSG00000136110 | 11061 | PWB-iPSC DN | CNMD | -0.71445 | | ENSG00000075223 | 10512 | PWB-EC DN | SEMA3C | -5.374481559 | | ENSG00000280809 | 101929052 | PWB-MSC DN | LINC00836 | -3.304487 |
| ENSG00000128309 | 4357 | PWB-iPSC DN | MPST | -0.71454 | | ENSG00000066248 | 25791 | PWB-EC DN | NGEF | -5.375564364 | | ENSG00000112619 | 5961 | PWB-MSC DN | PRPH2 | -3.314308 |
| ENSG00000175970 | 84747 | PWB-iPSC DN | UNC119B | -0.71639 | | ENSG00000198768 | 164284 | PWB-EC DN | APCDD1L | -5.382572545 | | ENSG00000100721 | 8115 | PWB-MSC DN | TCL1A | -3.316199 |
| ENSG00000186468 | 6228 | PWB-iPSC DN | RPS23 | -0.71655 | | ENSG00000189223 | 654433 | PWB-EC DN | PAX8-AS1 | -5.400198318 | | ENSG00000147113 | 79742 | PWB-MSC DN | DIPK2B | -3.318935 |
| ENSG00000186283 | 64222 | PWB-iPSC DN | TOR3A | -0.71706 | | ENSG00000041353 | 5874 | PWB-EC DN | RAB27B | -5.417980985 | | ENSG00000249378 | 401164 | PWB-MSC DN | LINC01060 | -3.323937 |
| ENSG00000184076 | 29796 | PWB-iPSC DN | UQCR10 | -0.72068 | | ENSG00000197632 | 5055 | PWB-EC DN | SERPINB2 | -5.423036448 | | ENSG00000151962 | 166863 | PWB-MSC DN | RBM46 | -3.33082 |
| ENSG00000100167 | 55964 | PWB-iPSC DN | SEPTIN3 | -0.72127 | | ENSG00000166396 | 8710 | PWB-EC DN | SERPINB7 | -5.423283574 | | ENSG00000278522 | 102724631 | PWB-MSC DN | POTEB3 | -3.385228 |
| ENSG00000164251 | 2150 | PWB-iPSC DN | F2RL1 | -0.72267 | | ENSG00000249306 | 101928176 | PWB-EC DN | LINC01411 | -5.425017012 | | ENSG00000214081 | 100132708 | PWB-MSC DN | CYP4F30P | -3.397456 |
| ENSG00000172379 | 9915 | PWB-iPSC DN | ARNT2 | -0.7227 | | ENSG00000137809 | 22801 | PWB-EC DN | ITGA11 | -5.426746642 | | ENSG00000185272 | 54033 | PWB-MSC DN | RBM11 | -3.453374 |
| ENSG00000140548 | 374655 | PWB-iPSC DN | ZNF710 | -0.72273 | | ENSG00000182667 | 50863 | PWB-EC DN | NTM | -5.434990395 | | ENSG00000185745 | 3434 | PWB-MSC DN | IFIT1 | -3.630775 |
| ENSG00000144909 | 114885 | PWB-iPSC DN | OSBPL11 | -0.72432 | | ENSG00000041982 | 3371 | PWB-EC DN | TNC | -5.436403314 | | ENSG00000119922 | 3433 | PWB-MSC DN | IFIT2 | -3.693711 |
| ENSG00000002330 | 572 | PWB-iPSC DN | BAD | -0.72468 | | ENSG00000245526 | 645323 | PWB-EC DN | LINC00461 | -5.436727815 | | ENSG00000074706 | 26034 | PWB-MSC DN | IPCEF1 | -3.777106 |
| ENSG00000113296 | 7060 | PWB-iPSC DN | THBS4 | -0.72513 | | ENSG00000125285 | 11166 | PWB-EC DN | SOX21 | -5.446480055 | | ENSG00000261780 | 100505817 | PWB-MSC DN | LINC02582 | -3.819091 |
| ENSG00000169714 | 7555 | PWB-iPSC DN | CNBP | -0.72514 | | ENSG00000163485 | 134 | PWB-EC DN | ADORA1 | -5.464917554 | | ENSG00000261517 | 100861552 | PWB-MSC DN | LINC00558 | -3.860948 |
| ENSG00000148158 | 401548 | PWB-iPSC DN | SNX30 | -0.72671 | | ENSG00000160307 | 6285 | PWB-EC DN | S100B | -5.466155561 | | ENSG00000204792 | 102724515 | PWB-MSC DN | LINC01291 | -3.89068 |
| ENSG00000204304 | 5089 | PWB-iPSC DN | PBX2 | -0.72805 | | ENSG00000091986 | 151887 | PWB-EC DN | CCDC80 | -5.466248021 | | ENSG00000086288 | 51314 | PWB-MSC DN | NME8 | -3.94222 |
| ENSG00000083067 | 80036 | PWB-iPSC DN | TRPM3 | -0.72835 | | ENSG00000160886 | 54742 | PWB-EC DN | LY6K | -5.47339933 | | ENSG00000177354 | 118461 | PWB-MSC DN | C10orf71 | -4.189328 |
| ENSG00000111671 | 84727 | PWB-iPSC DN | SPSB2 | -0.72861 | | ENSG00000049540 | 2006 | PWB-EC DN | ELN | -5.475657392 | | ENSG00000196735 | 3117 | PWB-MSC DN | HLA-DQA1 | -4.266776 |
| ENSG00000036672 | 9099 | PWB-iPSC DN | USP2 | -0.72866 | | ENSG00000232530 | 110806283 | PWB-EC DN | LIF-AS1 | -5.47676678 | | ENSG00000231431 | 440910 | PWB-MSC DN | LOC440910 | -4.272994 |
| ENSG00000130203 | 348 | PWB-iPSC DN | APOE | -0.72927 | | ENSG00000116132 | 5396 | PWB-EC DN | PRRX1 | -5.48867232 | | ENSG00000250420 | 729522 | PWB-MSC DN | AACSP1 | -4.29554 |
| ENSG00000149480 | 9219 | PWB-iPSC DN | MTA2 | -0.72936 | | ENSG00000168621 | 2668 | PWB-EC DN | GDNF | -5.4996249 | | ENSG00000141668 | 147381 | PWB-MSC DN | CBLN2 | -4.34688 |
| ENSG00000143622 | 6016 | PWB-iPSC DN | RIT1 | -0.72964 | | ENSG00000136542 | 11227 | PWB-EC DN | GALNT5 | -5.508413949 | | ENSG00000253716 | 100507316 | PWB-MSC DN | MINCR | -4.352669 |
| ENSG00000143862 | 127829 | PWB-iPSC DN | ARL8A | -0.72972 | | ENSG00000128606 | 10234 | PWB-EC DN | LRRC17 | -5.514256026 | | ENSG00000223760 | 285103 | PWB-MSC DN | MED15P9 | -4.575404 |
| ENSG00000186350 | 6256 | PWB-iPSC DN | RXRA | -0.73005 | | ENSG00000144355 | 1745 | PWB-EC DN | DLX1 | -5.534991497 | | ENSG00000180777 | 374860 | PWB-MSC DN | ANKRD30B | -4.611853 |
| ENSG00000135643 | 27345 | PWB-iPSC DN | KCNMB4 | -0.73048 | | ENSG00000130957 | 8789 | PWB-EC DN | FBP2 | -5.541492722 | | ENSG00000164283 | 11082 | PWB-MSC DN | ESM1 | -4.61653 |
| ENSG00000148303 | 6130 | PWB-iPSC DN | RPL7A | -0.73115 | | ENSG00000145934 | 57451 | PWB-EC DN | TENM2 | -5.543184006 | | ENSG00000265766 | 440224 | PWB-MSC DN | CXADRP3 | -4.668567 |
| ENSG00000164292 | 22836 | PWB-iPSC DN | RHOBTB3 | -0.73266 | | ENSG00000131981 | 3958 | PWB-EC DN | LGALS3 | -5.546317583 | | ENSG00000007908 | 6401 | PWB-MSC DN | SELE | -4.70916 |
| ENSG00000006282 | 64847 | PWB-iPSC DN | SPATA20 | -0.73316 | | ENSG00000185904 | 84856 | PWB-EC DN | LINC00839 | -5.558642254 | | ENSG00000105472 | 6320 | PWB-MSC DN | CLEC11A | -4.860706 |
| ENSG00000176641 | 220441 | PWB-iPSC DN | RNF152 | -0.7337 | | ENSG00000164692 | 1278 | PWB-EC DN | COL1A2 | -5.559313984 | | ENSG00000179344 | 3119 | PWB-MSC DN | HLA-DQB1 | -4.897148 |
| ENSG00000185565 | 4045 | PWB-iPSC DN | LSAMP | -0.73371 | | ENSG00000155511 | 2890 | PWB-EC DN | GRIA1 | -5.566833859 | | ENSG00000166351 | 317754 | PWB-MSC DN | POTED | -4.909649 |
| ENSG00000176788 | 10409 | PWB-iPSC DN | BASP1 | -0.73392 | | ENSG00000111728 | 6489 | PWB-EC DN | ST8SIA1 | -5.573602949 | | ENSG00000113262 | 2916 | PWB-MSC DN | GRM6 | -5.157014 |
| ENSG00000165449 | 220963 | PWB-iPSC DN | SLC16A9 | -0.73454 | | ENSG00000150551 | 116372 | PWB-EC DN | LYPD1 | -5.577749318 | | ENSG00000183206 | 388468 | PWB-MSC DN | POTEC | -5.183277 |
| ENSG00000126950 | 59353 | PWB-iPSC DN | TMEM35A | -0.73607 | | ENSG00000212719 | 339263 | PWB-EC DN | LINC02693 | -5.599262214 | | ENSG00000277630 | 100288966 | PWB-MSC DN | LOC100288966 | -5.310581 |
| ENSG00000132932 | 51761 | PWB-iPSC DN | ATP8A2 | -0.7372 | | ENSG00000221963 | 80830 | PWB-EC DN | APOL6 | -5.601564164 | | ENSG00000130957 | 8789 | PWB-MSC DN | FBP2 | -5.622268 |
| ENSG00000166341 | 8642 | PWB-iPSC DN | DCHS1 | -0.73779 | | ENSG00000204531 | 5460 | PWB-EC DN | POU5F1 | -5.601808374 | | ENSG00000100191 | 6527 | PWB-MSC DN | SLC5A4 | -5.87192 |
| ENSG00000073712 | 10979 | PWB-iPSC DN | FERMT2 | -0.73815 | | ENSG00000226806 | 100507600 | PWB-EC DN | LCT-AS1 | -5.630797198 | | ENSG00000205212 | 339184 | PWB-MSC DN | CCDC144NL | -5.886292 |
| ENSG00000116871 | 55700 | PWB-iPSC DN | MAP7D1 | -0.73963 | | ENSG00000145536 | 170690 | PWB-EC DN | ADAMTS16 | -5.670680654 | | ENSG00000224957 | 101927215 | PWB-MSC DN | LINC01266 | -5.962558 |
| ENSG00000111669 | 7167 | PWB-iPSC DN | TPI1 | -0.74008 | | ENSG00000116983 | 51440 | PWB-EC DN | HPCAL4 | -5.693782632 | | ENSG00000178187 | 285676 | PWB-MSC DN | ZNF454 | -6.133223 |
| ENSG00000129195 | 54478 | PWB-iPSC DN | PIMREG | -0.74097 | | ENSG00000233532 | 728192 | PWB-EC DN | LINC00460 | -5.696647515 | | ENSG00000188992 | 149998 | PWB-MSC DN | LIPI | -6.529304 |
| ENSG00000167969 | 1632 | PWB-iPSC DN | ECI1 | -0.74335 | | ENSG00000152977 | 7545 | PWB-EC DN | ZIC1 | -5.732910509 | | ENSG00000233098 | 339260 | PWB-MSC DN | LOC339260 | -6.581565 |
| ENSG00000242498 | 348110 | PWB-iPSC DN | ARPIN | -0.74666 | | ENSG00000125965 | 8200 | PWB-EC DN | GDF5 | -5.75161423 | | ENSG00000215912 | 100287898 | PWB-MSC DN | TTC34 | -6.820221 |
| ENSG00000154917 | 51560 | PWB-iPSC DN | RAB6B | -0.74902 | | ENSG00000120149 | 4488 | PWB-EC DN | MSX2 | -5.760574814 | | ENSG00000233608 | 117581 | PWB-MSC DN | TWIST2 | -7.045603 |
| ENSG00000175287 | 254295 | PWB-iPSC DN | PHYHD1 | -0.74954 | | ENSG00000170561 | 153572 | PWB-EC DN | IRX2 | -5.783679727 | | ENSG00000212719 | 339263 | PWB-MSC DN | LINC02693 | -7.180511 |
| ENSG00000182534 | 439921 | PWB-iPSC DN | MXRA7 | -0.75135 | | ENSG00000176049 | 9832 | PWB-EC DN | JAKMIP2 | -5.816350581 | | ENSG00000147223 | 92129 | PWB-MSC DN | RIPPLY1 | -7.376345 |
| ENSG00000144118 | 5899 | PWB-iPSC DN | RALB | -0.7518 | | ENSG00000176956 | 4062 | PWB-EC DN | LY6H | -5.816949357 | | ENSG00000181638 | 286128 | PWB-MSC DN | ZFP41 | -7.665695 |
| ENSG00000048052 | 9734 | PWB-iPSC DN | HDAC9 | -0.75224 | | ENSG00000172020 | 2596 | PWB-EC DN | GAP43 | -5.820663457 | | ENSG00000242082 | 110806273 | PWB-MSC DN | SLC5A4-AS1 | -7.779052 |
| ENSG00000101782 | 8780 | PWB-iPSC DN | RIOK3 | -0.75224 | | ENSG00000134817 | 187 | PWB-EC DN | APLNR | -5.836742963 | | ENSG00000130193 | 51337 | PWB-MSC DN | THEM6 | -7.920076 |
| ENSG00000133243 | 55643 | PWB-iPSC DN | BTBD2 | -0.75246 | | ENSG00000185567 | 113146 | PWB-EC DN | AHNAK2 | -5.847621003 | | ENSG00000086205 | 2346 | PWB-MSC DN | FOLH1 | -8.135641 |
| ENSG00000173950 | 152002 | PWB-iPSC DN | XXYLT1 | -0.75279 | | ENSG00000233521 | 105372978 | PWB-EC DN | LINC01638 | -5.855796814 | | ENSG00000214049 | 652995 | PWB-MSC DN | UCA1 | -8.716991 |
| ENSG00000058668 | 493 | PWB-iPSC DN | ATP2B4 | -0.7553 | | ENSG00000147041 | 94122 | PWB-EC DN | SYTL5 | -5.86509051 | | ENSG00000198105 | 57209 | PWB-MSC DN | ZNF248 | -9.004243 |
| ENSG00000204308 | 6048 | PWB-iPSC DN | RNF5 | -0.75548 | | ENSG00000125726 | 970 | PWB-EC DN | CD70 | -5.865923798 | | ENSG00000234616 | 8629 | PWB-MSC DN | JRK | -9.207588 |
| ENSG00000169372 | 8738 | PWB-iPSC DN | CRADD | -0.75663 | | ENSG00000170549 | 79192 | PWB-EC DN | IRX1 | -5.87969251 | | ENSG00000180155 | 66004 | PWB-MSC DN | LYNX1 | -9.384043 |
| ENSG00000205978 | 57523 | PWB-iPSC DN | NYNRIN | -0.75728 | | ENSG00000223485 | 101929484 | PWB-EC DN | LINC01615 | -5.903491505 | | ENSG00000160932 | 4061 | PWB-MSC DN | LY6E | -9.757836 |
| ENSG00000156475 | 5521 | PWB-iPSC DN | PPP2R2B | -0.75737 | | ENSG00000139800 | 85416 | PWB-EC DN | ZIC5 | -5.908199639 | | ENSG00000177335 | 286122 | PWB-MSC DN | C8orf31 | -10.52677 |
| ENSG00000132406 | 85013 | PWB-iPSC DN | TMEM128 | -0.75818 | | ENSG00000118785 | 6696 | PWB-EC DN | SPP1 | -5.923552953 | | | | | | |

#### Supplementary File 2 Hippo, Wnt, and ZNF DEGs (FDR < 0.05)

##### Hippo & Wnt Signatures

| id | ENTREZID | Group | ID.SYMBOL | Hippo | Wnt | logFC |
|---|---|---|---|---|---|---|
| ENSG00000075290 | 7479 | PWB-iPSC UP | WNT8B | Yes | Yes | 1.1 |
| ENSG00000150672 | 1740 | PWB-iPSC UP | DLG2 | Yes | Yes | 0.79 |
| ENSG00000081059 | 6932 | PWB-iPSC UP | TCF7 | Yes | Yes | 0.65 |
| ENSG00000141551 | 1453 | PWB-iPSC DN | CSNK1D | Yes | Yes | -0.41 |
| ENSG00000141646 | 4089 | PWB-iPSC DN | SMAD4 | Yes | Yes | -0.46 |
| ENSG00000137693 | 10413 | PWB-iPSC DN | YAP1 | Yes | Yes | -0.46 |
| ENSG00000039068 | 999 | PWB-iPSC DN | CDH1 | Yes | Yes | -0.54 |
| ENSG00000072803 | 23291 | PWB-iPSC DN | FBXW11 | Yes | Yes | -0.59 |
| ENSG00000171843 | 4300 | PWB-iPSC DN | MLLT3 | Yes | Yes | -0.6 |
| ENSG00000168036 | 1499 | PWB-iPSC DN | CTNNB1 | Yes | Yes | -0.66 |
| ENSG00000140807 | 85407 | PWB-iPSC DN | NKD1 | Yes | Yes | -0.71 |
| ENSG00000139174 | 144165 | PWB-iPSC DN | PRICKLE1 | Yes | Yes | -0.94 |
| ENSG00000166710 | 567 | PWB-iPSC DN | B2M | Yes | Yes | -1 |
| ENSG00000181449 | 6657 | PWB-iPSC DN | SOX2 | Yes | Yes | -1.08 |
| ENSG00000115266 | 10297 | PWB-iPSC DN | APC2 | Yes | Yes | -1.22 |
| ENSG00000157240 | 8321 | PWB-iPSC DN | FZD1 | Yes | Yes | -1.22 |
| ENSG00000163251 | 7855 | PWB-iPSC DN | FZD5 | Yes | Yes | -1.26 |
| ENSG00000162552 | 54361 | PWB-iPSC DN | WNT4 | Yes | Yes | -1.44 |
| ENSG00000125845 | 650 | PWB-iPSC DN | BMP2 | Yes | Yes | -1.58 |
| ENSG00000111186 | 81029 | PWB-iPSC DN | WNT5B | Yes | Yes | -1.58 |
| ENSG00000177283 | 8325 | PWB-iPSC DN | FZD8 | Yes | Yes | -1.64 |
| ENSG00000180340 | 2535 | PWB-iPSC DN | FZD2 | Yes | Yes | -2.2 |
| ENSG00000061492 | 7478 | PWB-iPSC DN | WNT8A | Yes | Yes | -3.49 |
| ENSG00000155760 | 8324 | PWB-EC DN | FZD7 | Yes | Yes | -2.61 |
| ENSG00000180340 | 2535 | PWB-EC DN | FZD2 | Yes | Yes | -3.52 |
| ENSG00000111186 | 81029 | PWB-EC DN | WNT5B | Yes | Yes | -3.8 |
| ENSG00000138795 | 51176 | PWB-EC DN | LEF1 | Yes | Yes | -4.6 |
| ENSG00000019549 | 6591 | PWB-EC DN | SNAI2 | Yes | Yes | -4.95 |
| ENSG00000140807 | 85407 | PWB-EC DN | NKD1 | Yes | Yes | -5.37 |
| ENSG00000181449 | 6657 | PWB-EC DN | SOX2 | Yes | Yes | -8.12 |
| ENSG00000154342 | 89780 | PWB-MSC UP | WNT3A | Yes | Yes | 2.02 |
| ENSG00000139174 | 144165 | PWB-MSC UP | PRICKLE1 | Yes | Yes | 0.91 |
| ENSG00000132535 | 1742 | PWB-MSC DN | DLG4 | Yes | Yes | -0.68 |
| ENSG00000180340 | 2535 | PWB-MSC DN | FZD2 | Yes | Yes | -1.41 |
| ENSG00000158955 | 7484 | PWB-MSC DN | WNT9B | Yes | Yes | -1.45 |
| ENSG00000002745 | 51384 | PWB-MSC DN | WNT16 | Yes | Yes | -1.66 |
| ENSG00000125965 | 8200 | PWB-iPSC UP | GDF5 | Yes | | 1.68 |
| ENSG00000152402 | 2977 | PWB-iPSC UP | GUCY1A2 | Yes | | 1.12 |
| ENSG00000166535 | 144568 | PWB-iPSC UP | A2ML1 | Yes | | 0.78 |
| ENSG00000101546 | 79863 | PWB-iPSC UP | RBFA | Yes | | 0.71 |
| ENSG00000073350 | 3993 | PWB-iPSC DN | LLGL2 | Yes | | -0.41 |
| ENSG00000104872 | 55011 | PWB-iPSC DN | PIH1D1 | Yes | | -0.42 |
| ENSG00000131899 | 3996 | PWB-iPSC DN | LLGL1 | Yes | | -0.46 |
| ENSG00000132842 | 8546 | PWB-iPSC DN | AP3B1 | Yes | | -0.47 |
| ENSG00000196792 | 29966 | PWB-iPSC DN | STRN3 | Yes | | -0.48 |
| ENSG00000170027 | 7532 | PWB-iPSC DN | YWHAG | Yes | | -0.48 |
| ENSG00000125835 | 6628 | PWB-iPSC DN | SNRPB | Yes | | -0.5 |
| ENSG00000213639 | 5500 | PWB-iPSC DN | PPP1CB | Yes | | -0.5 |
| ENSG00000186575 | 4771 | PWB-iPSC DN | NF2 | Yes | | -0.52 |
| ENSG00000067606 | 5590 | PWB-iPSC DN | PRKCZ | Yes | | -0.53 |
| ENSG00000205476 | 317762 | PWB-iPSC DN | CCDC85C | Yes | | -0.55 |
| ENSG00000044115 | 1495 | PWB-iPSC DN | CTNNA1 | Yes | | -0.56 |
| ENSG00000264364 | 140735 | PWB-iPSC DN | DYNLL2 | Yes | | -0.56 |
| ENSG00000143514 | 7159 | PWB-iPSC DN | TP53BP2 | Yes | | -0.57 |
| ENSG00000147044 | 8573 | PWB-iPSC DN | CASK | Yes | | -0.58 |
| ENSG00000130479 | 55201 | PWB-iPSC DN | MAP1S | Yes | | -0.61 |
| ENSG00000106799 | 7046 | PWB-iPSC DN | TGFBR1 | Yes | | -0.61 |
| ENSG00000146112 | 170954 | PWB-iPSC DN | PPP1R18 | Yes | | -0.67 |
| ENSG00000178184 | 84552 | PWB-iPSC DN | PARD6G | Yes | | -0.68 |
| ENSG00000108953 | 7531 | PWB-iPSC DN | YWHAE | Yes | | -0.71 |
| ENSG00000111052 | 8825 | PWB-iPSC DN | LIN7A | Yes | | -0.71 |
| ENSG00000166341 | 8642 | PWB-iPSC DN | DCHS1 | Yes | | -0.74 |
| ENSG00000156475 | 5521 | PWB-iPSC DN | PPP2R2B | Yes | | -0.76 |
| ENSG00000148180 | 2934 | PWB-iPSC DN | GSN | Yes | | -0.78 |
| ENSG00000126016 | 154796 | PWB-iPSC DN | AMOT | Yes | | -0.81 |
| ENSG00000108819 | 84687 | PWB-iPSC DN | PPP1R9B | Yes | | -0.91 |
| ENSG00000101144 | 655 | PWB-iPSC DN | BMP7 | Yes | | -0.92 |
| ENSG00000091129 | 4897 | PWB-iPSC DN | NRCAM | Yes | | -1.01 |
| ENSG00000066032 | 1496 | PWB-iPSC DN | CTNNA2 | Yes | | -1.07 |
| ENSG00000163545 | 81788 | PWB-iPSC DN | NUAK2 | Yes | | -1.09 |

##### ZNF Signatures

| id | ENTREZID | Group | ID.SYMBOL | Zinc finger Group name | logFC |
|---|---|---|---|---|---|
| ENSG00000173068 | 54796 | PWB-EC DN | BNC2 | Zinc fingers C2H2-type | -2.77 |
| ENSG00000171604 | 51523 | PWB-EC DN | CXXC5 | Zinc fingers CXXC-type | -1.45 |
| ENSG00000102385 | 1821 | PWB-EC DN | DRP2 | Zinc fingers ZZ-type | -5.26 |
| ENSG00000178498 | 196403 | PWB-EC DN | DTX3 | Ring finger proteins | -2.11 |
| ENSG00000158163 | 199221 | PWB-EC DN | DZIP1L | Zinc fingers C2H2-type | -2.42 |
| ENSG00000120738 | 1958 | PWB-EC DN | EGR1 | Zinc fingers C2H2-type | -2.37 |
| ENSG00000022267 | 2273 | PWB-EC DN | FHL1 | LIM domain containing | -1.98 |
| ENSG00000115641 | 2274 | PWB-EC DN | FHL2 | LIM domain containing | -1.88 |
| ENSG00000111087 | 2735 | PWB-EC DN | GLI1 | Zinc fingers C2H2-type | -3.52 |
| ENSG00000074047 | 2736 | PWB-EC DN | GLI2 | Zinc fingers C2H2-type | -5.36 |
| ENSG00000174332 | 148979 | PWB-EC DN | GLIS1 | Zinc fingers C2H2-type | -4.58 |
| ENSG00000136153 | 4008 | PWB-EC DN | LMO7 | LIM domain containing | -2.04 |
| ENSG00000143867 | 130497 | PWB-EC DN | OSR1 | Zinc fingers C2H2-type | -3.64 |
| ENSG00000154553 | 27295 | PWB-EC DN | PDLIM3 | LIM domain containing | -3.03 |
| ENSG00000131435 | 8572 | PWB-EC DN | PDLIM4 | LIM domain containing | -3.34 |
| ENSG00000121440 | 23024 | PWB-EC DN | PDZRN3 | Zinc fingers TRAF-type | -4.35 |
| ENSG00000132170 | 5468 | PWB-EC DN | PPARG | ome proliferator activated re | -2.63 |
| ENSG00000077092 | 5915 | PWB-EC DN | RARB | Retinoic acid receptors | -3.77 |
| ENSG00000141576 | 114804 | PWB-EC DN | RNF157 | Ring finger proteins | -3.69 |
| ENSG00000019549 | 6591 | PWB-EC DN | SNAI2 | Zinc fingers C2H2-type | -4.95 |
| ENSG00000104447 | 7227 | PWB-EC DN | TRPS1 | Zinc fingers C2H2-type | -4.23 |
| ENSG00000276043 | 29128 | PWB-EC DN | UHRF1 | Ring finger proteins | -2.77 |
| ENSG00000111424 | 7421 | PWB-EC DN | VDR | :lear receptor subfamily 1 gro | -5.24 |
| ENSG00000091656 | 79776 | PWB-EC DN | ZFHX4 | Zinc fingers matrin-type | -4.64 |
| ENSG00000181638 | 286128 | PWB-EC DN | ZFP41 | Zinc fingers C2H2-type | -4.59 |
| ENSG00000152977 | 7545 | PWB-EC DN | ZIC1 | Zinc fingers C2H2-type | -5.73 |
| ENSG00000043355 | 7546 | PWB-EC DN | ZIC2 | Zinc fingers C2H2-type | -7.94 |
| ENSG00000139800 | 85416 | PWB-EC DN | ZIC5 | Zinc fingers C2H2-type | -5.91 |
| ENSG00000165061 | 79698 | PWB-EC DN | ZMAT4 | Zinc fingers matrin-type | -4.83 |
| ENSG00000196793 | 8187 | PWB-EC DN | ZNF239 | Zinc fingers C2H2-type | -3.94 |
| ENSG00000198105 | 57209 | PWB-EC DN | ZNF248 | Zinc fingers C2H2-type | -7.16 |
| ENSG00000145908 | 91975 | PWB-EC DN | ZNF300 | Zinc fingers C2H2-type | -1.68 |
| ENSG00000075407 | 7587 | PWB-EC DN | ZNF37A | Zinc fingers C2H2-type | -2.67 |
| ENSG00000178187 | 285676 | PWB-EC DN | ZNF454 | Zinc fingers C2H2-type | -3.37 |
| ENSG00000165655 | 84858 | PWB-EC DN | ZNF503 | Zinc fingers C2H2-type | -2.08 |
| ENSG00000198597 | 9745 | PWB-EC DN | ZNF536 | Zinc fingers C2H2-type | -4.82 |
| ENSG00000198205 | 7789 | PWB-EC DN | ZXDA | Zinc fingers C2H2-type | -3.41 |
| ENSG00000023445 | 330 | PWB-EC UP | BIRC3 | Ring finger proteins | 2.67 |
| ENSG00000136944 | 4010 | PWB-EC UP | LMX1B | LIM class homeoboxes | 5.48 |
| ENSG00000034677 | 25897 | PWB-EC UP | RNF19A | Ring finger proteins | 1.67 |
| ENSG00000175893 | 340481 | PWB-EC UP | ZDHHC21 | Zinc fingers DHHC-type | 1.5 |
| ENSG00000141497 | 84225 | PWB-EC UP | ZMYND15 | Zinc fingers MYND-type | 3.15 |
| ENSG00000176293 | 7694 | PWB-EC UP | ZNF135 | Zinc fingers C2H2-type | 1.69 |
| ENSG00000160321 | 7757 | PWB-EC UP | ZNF208 | Zinc fingers C2H2-type | 3.14 |
| ENSG00000197134 | 113835 | PWB-EC UP | ZNF257 | Zinc fingers C2H2-type | 4.78 |
| ENSG00000167555 | 84436 | PWB-EC UP | ZNF528 | Zinc fingers C2H2-type | 1.8 |
| ENSG00000198028 | 147741 | PWB-EC UP | ZNF560 | Zinc fingers C2H2-type | 3.75 |
| ENSG00000258405 | 147660 | PWB-EC UP | ZNF578 | Zinc fingers C2H2-type | 2.58 |
| ENSG00000196109 | 163223 | PWB-EC UP | ZNF676 | Zinc fingers C2H2-type | 4.1 |
| ENSG00000269067 | 388523 | PWB-EC UP | ZNF728 | Zinc fingers C2H2-type | 4.3 |
| ENSG00000196350 | 1E+08 | PWB-EC UP | ZNF729 | Zinc fingers C2H2-type | 7.38 |
| ENSG00000197360 | 148198 | PWB-EC UP | ZNF98 | Zinc fingers C2H2-type | 4.81 |
| ENSG00000213973 | 7652 | PWB-EC UP | ZNF99 | Zinc fingers C2H2-type | 5.22 |
| ENSG00000152467 | 284312 | PWB-EC UP | ZSCAN1 | Zinc fingers C2H2-type | 3.28 |
| ENSG00000129691 | 9070 | PWB-iPSC DN | ASH2L | PHD finger proteins | -0.48 |
| ENSG00000161940 | 255877 | PWB-iPSC DN | BCL6B | Zinc fingers C2H2-type | -0.77 |
| ENSG00000168283 | 648 | PWB-iPSC DN | BMI1 | Ring finger proteins | -0.64 |
| ENSG00000111642 | 1108 | PWB-iPSC DN | CHD4 | PHD finger proteins | -0.4 |
| ENSG00000169714 | 7555 | PWB-iPSC DN | CNBP | Zinc fingers CCHC-type | -0.73 |
| ENSG00000124092 | 140690 | PWB-iPSC DN | CTCFL | Zinc fingers C2H2-type | -1.53 |
| ENSG00000177030 | 10522 | PWB-iPSC DN | DEAF1 | Zinc fingers MYND-type | -0.6 |
| ENSG00000099364 | 54620 | PWB-iPSC DN | FBXL19 | Zinc fingers CXXC-type | -0.62 |
| ENSG00000128610 | 389549 | PWB-iPSC DN | FEZF1 | Zinc fingers C2H2-type | -4.64 |
| ENSG00000153266 | 55079 | PWB-iPSC DN | FEZF2 | Zinc fingers C2H2-type | -3.84 |
| ENSG00000106571 | 2737 | PWB-iPSC DN | GLI3 | Zinc fingers C2H2-type | -0.9 |
| ENSG00000169635 | 23119 | PWB-iPSC DN | HIC2 | Zinc fingers C2H2-type | -0.61 |
| ENSG00000172273 | 25988 | PWB-iPSC DN | HINFP | Zinc fingers C2H2-type | -0.54 |
| ENSG00000168556 | 3622 | PWB-iPSC DN | ING2 | PHD finger proteins | -0.62 |
| ENSG00000109787 | 51274 | PWB-iPSC DN | KLF3 | Zinc fingers C2H2-type | -0.56 |
| ENSG00000136826 | 9314 | PWB-iPSC DN | KLF4 | Zinc fingers C2H2-type | -1.38 |

| | | | | | | |
|---|---|---|---|---|---|---|
| ENSG00000125968 | 3397 | PWB-iPSC DN | ID1 | Yes | | -1.36 |
| ENSG00000176887 | 6664 | PWB-iPSC DN | SOX11 | Yes | | -1.39 |
| ENSG00000115738 | 3398 | PWB-iPSC DN | ID2 | Yes | | -1.42 |
| ENSG00000148204 | 286204 | PWB-iPSC DN | CRB2 | Yes | | -1.61 |
| ENSG00000057149 | 6317 | PWB-EC UP | SERPINB3 | Yes | | 3.52 |
| ENSG00000166535 | 144568 | PWB-EC UP | A2ML1 | Yes | | 3.08 |
| ENSG00000023445 | 330 | PWB-EC UP | BIRC3 | Yes | | 2.67 |
| ENSG00000151208 | 9231 | PWB-EC DN | DLG5 | Yes | | -1.89 |
| ENSG00000061918 | 2983 | PWB-EC DN | GUCY1B1 | Yes | | -2.37 |
| ENSG00000007866 | 7005 | PWB-EC DN | TEAD3 | Yes | | -2.52 |
| ENSG00000116641 | 85440 | PWB-EC DN | DOCK7 | Yes | | -2.84 |
| ENSG00000171877 | 84978 | PWB-EC DN | FRMD5 | Yes | | -2.84 |
| ENSG00000092969 | 7042 | PWB-EC DN | TGFB2 | Yes | | -3.03 |
| ENSG00000107779 | 657 | PWB-EC DN | BMPR1A | Yes | | -3.36 |
| ENSG00000134602 | 51765 | PWB-EC DN | STK26 | Yes | | -3.86 |
| ENSG00000138696 | 658 | PWB-EC DN | BMPR1B | Yes | | -3.98 |
| ENSG00000163629 | 5783 | PWB-EC DN | PTPN13 | Yes | | -4.05 |
| ENSG00000082458 | 1741 | PWB-EC DN | DLG3 | Yes | | -4.09 |
| ENSG00000113578 | 2246 | PWB-EC DN | FGF1 | Yes | | -4.49 |
| ENSG00000082397 | 23136 | PWB-EC DN | EPB41L3 | Yes | | -4.51 |
| ENSG00000156466 | 392255 | PWB-EC DN | GDF6 | Yes | | -5.19 |
| ENSG00000074047 | 2736 | PWB-EC DN | GLI2 | Yes | | -5.36 |
| ENSG00000125965 | 8200 | PWB-EC DN | GDF5 | Yes | | -5.75 |
| ENSG00000148204 | 286204 | PWB-EC DN | CRB2 | Yes | | -6.27 |
| ENSG00000104899 | 268 | PWB-MSC UP | AMH | Yes | | 2.32 |
| ENSG00000153162 | 654 | PWB-MSC UP | BMP6 | Yes | | 1.71 |
| ENSG00000189334 | 57402 | PWB-MSC UP | S100A14 | Yes | | 1.64 |
| ENSG00000197905 | 7004 | PWB-MSC UP | TEAD4 | Yes | | 1.31 |
| ENSG00000113645 | 23286 | PWB-MSC UP | WWC1 | Yes | | 0.87 |
| ENSG00000247077 | 192111 | PWB-MSC UP | PGAM5 | Yes | | 0.84 |
| ENSG00000166908 | 79837 | PWB-MSC UP | PIP4K2C | Yes | | 0.64 |
| ENSG00000106066 | 54504 | PWB-MSC UP | CPVL | Yes | | 0.64 |
| ENSG00000166226 | 10576 | PWB-MSC UP | CCT2 | Yes | | 0.64 |
| ENSG00000110958 | 10728 | PWB-MSC UP | PTGES3 | Yes | | 0.47 |
| ENSG00000167553 | 84790 | PWB-MSC UP | TUBA1C | Yes | | 0.45 |
| ENSG00000178184 | 84552 | PWB-MSC DN | PARD6G | Yes | | -0.54 |
| ENSG00000176887 | 6664 | PWB-MSC DN | SOX11 | Yes | | -0.55 |
| ENSG00000146112 | 170954 | PWB-MSC DN | PPP1R18 | Yes | | -0.56 |
| ENSG00000166341 | 8642 | PWB-MSC DN | DCHS1 | Yes | | -0.85 |
| ENSG00000061918 | 2983 | PWB-MSC DN | GUCY1B1 | Yes | | -1.1 |
| ENSG00000196159 | 79633 | PWB-MSC DN | FAT4 | Yes | | -1.5 |
| ENSG00000092969 | 7042 | PWB-MSC DN | TGFB2 | Yes | | -2.74 |
| ENSG00000146047 | 255626 | PWB-iPSC UP | H2BC1 | | Yes | 2.02 |
| ENSG00000156076 | 11197 | PWB-iPSC UP | WIF1 | | Yes | 1.6 |
| ENSG00000105974 | 857 | PWB-iPSC UP | CAV1 | | Yes | 1.54 |
| ENSG00000147869 | 9350 | PWB-iPSC UP | CER1 | | Yes | 1.47 |
| ENSG00000165617 | 51339 | PWB-iPSC UP | DACT1 | | Yes | 1.4 |
| ENSG00000106588 | 5683 | PWB-iPSC UP | PSMA2 | | Yes | 1.08 |
| ENSG00000123104 | 3709 | PWB-iPSC UP | ITPR2 | | Yes | 1.06 |
| ENSG00000132846 | 84327 | PWB-iPSC UP | ZBED3 | | Yes | 1.04 |
| ENSG00000110888 | 65981 | PWB-iPSC UP | CAPRIN2 | | Yes | 0.93 |
| ENSG00000143801 | 5664 | PWB-iPSC UP | PSEN2 | | Yes | 0.89 |
| ENSG00000144642 | 27303 | PWB-iPSC UP | RBMS3 | | Yes | 0.83 |
| ENSG00000164362 | 7015 | PWB-iPSC UP | TERT | | Yes | 0.78 |
| ENSG00000126583 | 5582 | PWB-iPSC UP | PRKCG | | Yes | 0.76 |
| ENSG00000138798 | 1950 | PWB-iPSC UP | EGF | | Yes | 0.66 |
| ENSG00000008710 | 5310 | PWB-iPSC UP | PKD1 | | Yes | 0.64 |
| ENSG00000137841 | 5330 | PWB-iPSC UP | PLCB2 | | Yes | 0.62 |
| ENSG00000150995 | 3708 | PWB-iPSC UP | ITPR1 | | Yes | 0.59 |
| ENSG00000127603 | 23499 | PWB-iPSC UP | MACF1 | | Yes | 0.55 |
| ENSG00000090905 | 27327 | PWB-iPSC UP | TNRC6A | | Yes | 0.48 |
| ENSG00000163904 | 59343 | PWB-iPSC DN | SENP2 | | Yes | -0.48 |
| ENSG00000129691 | 9070 | PWB-iPSC DN | ASH2L | | Yes | -0.48 |
| ENSG00000078304 | 5527 | PWB-iPSC DN | PPP2R5C | | Yes | -0.5 |
| ENSG00000215301 | 1654 | PWB-iPSC DN | DDX3X | | Yes | -0.53 |
| ENSG00000097007 | 25 | PWB-iPSC DN | ABL1 | | Yes | -0.54 |
| ENSG00000122958 | 9559 | PWB-iPSC DN | VPS26A | | Yes | -0.54 |
| ENSG00000105229 | 51588 | PWB-iPSC DN | PIAS4 | | Yes | -0.57 |
| ENSG00000071564 | 6929 | PWB-iPSC DN | TCF3 | | Yes | -0.58 |
| ENSG00000156299 | 7074 | PWB-iPSC DN | TIAM1 | | Yes | -0.59 |
| ENSG00000101384 | 182 | PWB-iPSC DN | JAG1 | | Yes | -0.6 |
| ENSG00000185532 | 5592 | PWB-iPSC DN | PRKG1 | | Yes | -0.6 |
| ENSG00000163346 | 57326 | PWB-iPSC DN | PBXIP1 | | Yes | -0.62 |

| | | | | | |
|---|---|---|---|---|---|
| ENSG00000106689 | 9355 | PWB-iPSC DN | LHX2 | LIM class homeoboxes | -3.37 |
| ENSG00000089116 | 64211 | PWB-iPSC DN | LHX5 | LIM class homeoboxes | -3.51 |
| ENSG00000064042 | 22998 | PWB-iPSC DN | LIMCH1 | LIM domain containing | -1.41 |
| ENSG00000131914 | 79727 | PWB-iPSC DN | LIN28A | Zinc fingers CCHC-type | -0.64 |
| ENSG00000166407 | 4004 | PWB-iPSC DN | LMO1 | LIM domain containing | -3.84 |
| ENSG00000162761 | 4009 | PWB-iPSC DN | LMX1A | LIM class homeoboxes | -3.96 |
| ENSG00000173926 | 115123 | PWB-iPSC DN | MARCHF3 | Ring finger proteins | -0.82 |
| ENSG00000139266 | 92979 | PWB-iPSC DN | MARCHF9 | Ring finger proteins | -0.79 |
| ENSG00000085276 | 2122 | PWB-iPSC DN | MECOM | Zinc fingers C2H2-type | -2.49 |
| ENSG00000181588 | 399664 | PWB-iPSC DN | MEX3D | Ring finger proteins | -0.69 |
| ENSG00000102858 | 23295 | PWB-iPSC DN | MGRN1 | Ring finger proteins | -0.54 |
| ENSG00000179455 | 7681 | PWB-iPSC DN | MKRN3 | Ring finger proteins | -4.21 |
| ENSG00000174579 | 55167 | PWB-iPSC DN | MSL2 | Ring finger proteins | -0.56 |
| ENSG00000149480 | 9219 | PWB-iPSC DN | MTA2 | TA zinc finger domain contain | -0.73 |
| ENSG00000057935 | 57504 | PWB-iPSC DN | MTA3 | TA zinc finger domain contain | -0.68 |
| ENSG00000175745 | 7025 | PWB-iPSC DN | NR2F1 | lear receptor subfamily 2 gro | -5.46 |
| ENSG00000185551 | 7026 | PWB-iPSC DN | NR2F2 | lear receptor subfamily 2 gro | -5.87 |
| ENSG00000148200 | 2649 | PWB-iPSC DN | NR6A1 | lear receptor subfamily 6 grou | -0.56 |
| ENSG00000109685 | 7468 | PWB-iPSC DN | NSD2 | PHD finger proteins | -0.58 |
| ENSG00000152193 | 79596 | PWB-iPSC DN | OBI1 | Ring finger proteins | -1.01 |
| ENSG00000100105 | 23598 | PWB-iPSC DN | PATZ1 | Zinc fingers C2H2-type | -0.52 |
| ENSG00000116273 | 148479 | PWB-iPSC DN | PHF13 | PHD finger proteins | -0.58 |
| ENSG00000105229 | 51588 | PWB-iPSC DN | PIAS4 | Zinc fingers MIZ-type | -0.57 |
| ENSG00000181191 | 64219 | PWB-iPSC DN | PJA1 | Ring finger proteins | -0.58 |
| ENSG00000139174 | 144165 | PWB-iPSC DN | PRICKLE1 | LIM domain containing | -0.94 |
| ENSG00000171016 | 26108 | PWB-iPSC DN | PYGO1 | PHD finger proteins | -1 |
| ENSG00000123091 | 26994 | PWB-iPSC DN | RNF11 | Ring finger proteins | -0.62 |
| ENSG00000113269 | 55819 | PWB-iPSC DN | RNF130 | Ring finger proteins | -0.85 |
| ENSG00000151692 | 9781 | PWB-iPSC DN | RNF144A | Ring finger proteins | -0.61 |
| ENSG00000145860 | 153830 | PWB-iPSC DN | RNF145 | Ring finger proteins | -0.42 |
| ENSG00000176641 | 220441 | PWB-iPSC DN | RNF152 | Ring finger proteins | -0.73 |
| ENSG00000141576 | 114804 | PWB-iPSC DN | RNF157 | Ring finger proteins | -0.99 |
| ENSG00000168159 | 149603 | PWB-iPSC DN | RNF187 | Ring finger proteins | -0.78 |
| ENSG00000204308 | 6048 | PWB-iPSC DN | RNF5 | Ring finger proteins | -0.76 |
| ENSG00000198963 | 6096 | PWB-iPSC DN | RORB | RAR related orphan receptors | -0.89 |
| ENSG00000186350 | 6256 | PWB-iPSC DN | RXRA | Retinoid X receptors | -0.73 |
| ENSG00000103449 | 6299 | PWB-iPSC DN | SALL1 | Zinc fingers C2H2-type | -0.94 |
| ENSG00000256463 | 27164 | PWB-iPSC DN | SALL3 | Zinc fingers C2H2-type | -1.13 |
| ENSG00000101115 | 57167 | PWB-iPSC DN | SALL4 | Zinc fingers C2H2-type | -0.5 |
| ENSG00000124562 | 6631 | PWB-iPSC DN | SNRPC | Zinc fingers matrin-type | -0.53 |
| ENSG00000204335 | 389058 | PWB-iPSC DN | SP5 | Zinc fingers C2H2-type | -2.87 |
| ENSG00000164651 | 221833 | PWB-iPSC DN | SP8 | Zinc fingers C2H2-type | -1.05 |
| ENSG00000151090 | 7068 | PWB-iPSC DN | THRB | Thyroid hormone receptors | -1.09 |
| ENSG00000076604 | 9618 | PWB-iPSC DN | TRAF4 | Zinc fingers TRAF-type | -0.58 |
| ENSG00000204977 | 10206 | PWB-iPSC DN | TRIM13 | Ring finger proteins | -0.46 |
| ENSG00000113595 | 373 | PWB-iPSC DN | TRIM23 | Ring finger proteins | -0.51 |
| ENSG00000122779 | 8805 | PWB-iPSC DN | TRIM24 | Ring finger proteins | -0.58 |
| ENSG00000146833 | 89122 | PWB-iPSC DN | TRIM4 | Ring finger proteins | -0.64 |
| ENSG00000177238 | 493829 | PWB-iPSC DN | TRIM72 | Ring finger proteins | -1.78 |
| ENSG00000100426 | 9889 | PWB-iPSC DN | ZBED4 | Zinc fingers BED-type | -0.5 |
| ENSG00000205189 | 65986 | PWB-iPSC DN | ZBTB10 | Zinc fingers C2H2-type | -0.97 |
| ENSG00000204366 | 221527 | PWB-iPSC DN | ZBTB12 | Zinc fingers C2H2-type | -0.98 |
| ENSG00000236104 | 9278 | PWB-iPSC DN | ZBTB22 | Zinc fingers C2H2-type | -0.7 |
| ENSG00000169155 | 23099 | PWB-iPSC DN | ZBTB43 | Zinc fingers C2H2-type | -0.51 |
| ENSG00000130584 | 140685 | PWB-iPSC DN | ZBTB46 | Zinc fingers C2H2-type | -0.6 |
| ENSG00000160685 | 51043 | PWB-iPSC DN | ZBTB7B | Zinc fingers C2H2-type | -0.85 |
| ENSG00000174460 | 170261 | PWB-iPSC DN | ZCCHC12 | Zinc fingers CCHC-type | -2.03 |
| ENSG00000148516 | 6935 | PWB-iPSC DN | ZEB1 | Zinc fingers C2H2-type | -3.24 |
| ENSG00000169554 | 9839 | PWB-iPSC DN | ZEB2 | Zinc fingers C2H2-type | -6.13 |
| ENSG00000156639 | 60685 | PWB-iPSC DN | ZFAND3 | Zinc fingers AN1-type | -0.59 |
| ENSG00000198939 | 80108 | PWB-iPSC DN | ZFP2 | Zinc fingers C2H2-type | -0.99 |
| ENSG00000152518 | 678 | PWB-iPSC DN | ZFP36L2 | Zinc fingers CCCH-type | -1 |
| ENSG00000020256 | 55734 | PWB-iPSC DN | ZFP64 | Zinc fingers C2H2-type | -0.71 |
| ENSG00000105278 | 23217 | PWB-iPSC DN | ZFR2 | Zinc fingers matrin-type | -1.01 |
| ENSG00000005889 | 7543 | PWB-iPSC DN | ZFX | Zinc fingers C2H2-type | -0.6 |
| ENSG00000152977 | 7545 | PWB-iPSC DN | ZIC1 | Zinc fingers C2H2-type | -3.35 |
| ENSG00000043355 | 7546 | PWB-iPSC DN | ZIC2 | Zinc fingers C2H2-type | -1.06 |
| ENSG00000174963 | 84107 | PWB-iPSC DN | ZIC4 | Zinc fingers C2H2-type | -2.62 |
| ENSG00000139800 | 85416 | PWB-iPSC DN | ZIC5 | Zinc fingers C2H2-type | -0.77 |
| ENSG00000147394 | 7739 | PWB-iPSC DN | ZNF185 | LIM domain containing | -0.78 |
| ENSG00000165804 | 51222 | PWB-iPSC DN | ZNF219 | Zinc fingers C2H2-type | -0.67 |
| ENSG00000198105 | 57209 | PWB-iPSC DN | ZNF248 | Zinc fingers C2H2-type | -3.51 |
| ENSG00000169548 | 129025 | PWB-iPSC DN | ZNF280A | Zinc fingers C2H2-type | -2.71 |

| | | | | | |
|---|---|---|---|---|---|
| ENSG00000066468 | 2263 | PWB-iPSC DN | FGFR2 | Yes | -0.63 |
| ENSG00000132475 | 3021 | PWB-iPSC DN | H3-3B | Yes | -0.64 |
| ENSG00000101849 | 6907 | PWB-iPSC DN | TBL1X | Yes | -0.65 |
| ENSG00000127955 | 2770 | PWB-iPSC DN | GNAI1 | Yes | -0.65 |
| ENSG00000115457 | 3485 | PWB-iPSC DN | IGFBP2 | Yes | -0.66 |
| ENSG00000106829 | 7091 | PWB-iPSC DN | TLE4 | Yes | -0.66 |
| ENSG00000154229 | 5578 | PWB-iPSC DN | PRKCA | Yes | -0.67 |
| ENSG00000140332 | 7090 | PWB-iPSC DN | TLE3 | Yes | -0.68 |
| ENSG00000092010 | 5720 | PWB-iPSC DN | PSME1 | Yes | -0.7 |
| ENSG00000130203 | 348 | PWB-iPSC DN | APOE | Yes | -0.73 |
| ENSG00000073712 | 10979 | PWB-iPSC DN | FERMT2 | Yes | -0.74 |
| ENSG00000108840 | 10014 | PWB-iPSC DN | HDAC5 | Yes | -0.78 |
| ENSG00000160014 | 808 | PWB-iPSC DN | CALM3 | Yes | -0.8 |
| ENSG00000108821 | 1277 | PWB-iPSC DN | COL1A1 | Yes | -0.82 |
| ENSG00000142541 | 23521 | PWB-iPSC DN | RPL13A | Yes | -0.84 |
| ENSG00000078902 | 54472 | PWB-iPSC DN | TOLLIP | Yes | -0.85 |
| ENSG00000181274 | 23401 | PWB-iPSC DN | FRAT2 | Yes | -0.9 |
| ENSG00000106571 | 2737 | PWB-iPSC DN | GLI3 | Yes | -0.9 |
| ENSG00000205213 | 55366 | PWB-iPSC DN | LGR4 | Yes | -0.91 |
| ENSG00000103449 | 6299 | PWB-iPSC DN | SALL1 | Yes | -0.94 |
| ENSG00000169071 | 4920 | PWB-iPSC DN | ROR2 | Yes | -0.96 |
| ENSG00000110492 | 4192 | PWB-iPSC DN | MDK | Yes | -0.96 |
| ENSG00000118503 | 7128 | PWB-iPSC DN | TNFAIP3 | Yes | -0.96 |
| ENSG00000171016 | 26108 | PWB-iPSC DN | PYGO1 | Yes | -1 |
| ENSG00000178585 | 56998 | PWB-iPSC DN | CTNNBIP1 | Yes | -1.01 |
| ENSG00000105559 | 57664 | PWB-iPSC DN | PLEKHA4 | Yes | -1.06 |
| ENSG00000169862 | 1501 | PWB-iPSC DN | CTNND2 | Yes | -1.06 |
| ENSG00000164488 | 168002 | PWB-iPSC DN | DACT2 | Yes | -1.16 |
| ENSG00000125398 | 6662 | PWB-iPSC DN | SOX9 | Yes | -1.24 |
| ENSG00000104332 | 6422 | PWB-iPSC DN | SFRP1 | Yes | -1.25 |
| ENSG00000136826 | 9314 | PWB-iPSC DN | KLF4 | Yes | -1.38 |
| ENSG00000132386 | 5176 | PWB-iPSC DN | SERPINF1 | Yes | -1.4 |
| ENSG00000068078 | 2261 | PWB-iPSC DN | FGFR3 | Yes | -1.45 |
| ENSG00000167779 | 3489 | PWB-iPSC DN | IGFBP6 | Yes | -1.48 |
| ENSG00000147257 | 2719 | PWB-iPSC DN | GPC3 | Yes | -1.51 |
| ENSG00000109339 | 5602 | PWB-iPSC DN | MAPK10 | Yes | -1.51 |
| ENSG00000180694 | 169200 | PWB-iPSC DN | TMEM64 | Yes | -1.59 |
| ENSG00000116729 | 79971 | PWB-iPSC DN | WLS | Yes | -1.66 |
| ENSG00000116574 | 58480 | PWB-iPSC DN | RHOU | Yes | -1.71 |
| ENSG00000172201 | 3400 | PWB-iPSC DN | ID4 | Yes | -1.75 |
| ENSG00000163666 | 8820 | PWB-iPSC DN | HESX1 | Yes | -1.84 |
| ENSG00000167874 | 92162 | PWB-iPSC DN | TMEM88 | Yes | -1.91 |
| ENSG00000110195 | 2348 | PWB-iPSC DN | FOLR1 | Yes | -2.07 |
| ENSG00000152804 | 3087 | PWB-iPSC DN | HHEX | Yes | -2.1 |
| ENSG00000064195 | 1747 | PWB-iPSC DN | DLX3 | Yes | -2.35 |
| ENSG00000169218 | 284654 | PWB-iPSC DN | RSPO1 | Yes | -2.54 |
| ENSG00000162998 | 2487 | PWB-iPSC DN | FRZB | Yes | -2.8 |
| ENSG00000204335 | 389058 | PWB-iPSC DN | SP5 | Yes | -2.87 |
| ENSG00000162761 | 4009 | PWB-iPSC DN | LMX1A | Yes | -3.96 |
| ENSG00000166923 | 26585 | PWB-iPSC DN | GREM1 | Yes | -4.11 |
| ENSG00000138083 | 6496 | PWB-iPSC DN | SIX3 | Yes | -4.56 |
| ENSG00000105880 | 1749 | PWB-iPSC DN | DLX5 | Yes | -4.73 |
| ENSG00000169554 | 9839 | PWB-iPSC DN | ZEB2 | Yes | -6.13 |
| ENSG00000261594 | 100507050 | PWB-EC UP | TPBGL | Yes | 4.14 |
| ENSG00000138083 | 6496 | PWB-EC UP | SIX3 | Yes | 3.31 |
| ENSG00000118503 | 7128 | PWB-EC UP | TNFAIP3 | Yes | 1.87 |
| ENSG00000196562 | 55959 | PWB-EC UP | SULF2 | Yes | 1.65 |
| ENSG00000167191 | 51704 | PWB-EC UP | GPRC5B | Yes | 1.51 |
| ENSG00000273703 | 8342 | PWB-EC DN | H2BC14 | Yes | -1.8 |
| ENSG00000115641 | 2274 | PWB-EC DN | FHL2 | Yes | -1.88 |
| ENSG00000123384 | 4035 | PWB-EC DN | LRP1 | Yes | -2.29 |
| ENSG00000278588 | 8346 | PWB-EC DN | H2BC10 | Yes | -2.3 |
| ENSG00000107984 | 22943 | PWB-EC DN | DKK1 | Yes | -2.31 |
| ENSG00000165617 | 51339 | PWB-EC DN | DACT1 | Yes | -2.32 |
| ENSG00000205213 | 55366 | PWB-EC DN | LGR4 | Yes | -2.34 |
| ENSG00000026508 | 960 | PWB-EC DN | CD44 | Yes | -2.35 |
| ENSG00000120738 | 1958 | PWB-EC DN | EGR1 | Yes | -2.37 |
| ENSG00000276180 | 8294 | PWB-EC DN | H4C9 | Yes | -2.44 |
| ENSG00000185130 | 8340 | PWB-EC DN | H2BC13 | Yes | -2.48 |
| ENSG00000149948 | 8091 | PWB-EC DN | HMGA2 | Yes | -2.63 |
| ENSG00000183598 | 653604 | PWB-EC DN | H3C13 | Yes | -2.74 |
| ENSG00000173848 | 10276 | PWB-EC DN | NET1 | Yes | -2.96 |
| ENSG00000180730 | 387914 | PWB-EC DN | SHISA2 | Yes | -3.01 |

| | | | | | |
|---|---|---|---|---|---|
| ENSG00000275004 | 140883 | PWB-iPSC DN | ZNF280B | Zinc fingers C2H2-type | -0.78 |
| ENSG00000205903 | 1E+08 | PWB-iPSC DN | ZNF316 | Zinc fingers C2H2-type | -0.67 |
| ENSG00000113761 | 23567 | PWB-iPSC DN | ZNF346 | Zinc fingers matrin-type | -0.46 |
| ENSG00000198816 | 140467 | PWB-iPSC DN | ZNF358 | Zinc fingers C2H2-type | -0.86 |
| ENSG00000186918 | 55893 | PWB-iPSC DN | ZNF395 | Zinc fingers C2H2-type | -0.69 |
| ENSG00000102935 | 23090 | PWB-iPSC DN | ZNF423 | Zinc fingers C2H2-type | -0.51 |
| ENSG00000131116 | 126299 | PWB-iPSC DN | ZNF428 | Zinc fingers C2H2-type | -0.92 |
| ENSG00000180035 | 197407 | PWB-iPSC DN | ZNF48 | Zinc fingers C2H2-type | -0.88 |
| ENSG00000198795 | 25925 | PWB-iPSC DN | ZNF521 | Zinc fingers C2H2-type | -1.14 |
| ENSG00000198597 | 9745 | PWB-iPSC DN | ZNF536 | Zinc fingers C2H2-type | -2.01 |
| ENSG00000186300 | 148254 | PWB-iPSC DN | ZNF555 | Zinc fingers C2H2-type | -0.79 |
| ENSG00000171425 | 51545 | PWB-iPSC DN | ZNF581 | Zinc fingers C2H2-type | -0.81 |
| ENSG00000198182 | 84775 | PWB-iPSC DN | ZNF607 | Zinc fingers C2H2-type | -0.85 |
| ENSG00000161914 | 115950 | PWB-iPSC DN | ZNF653 | Zinc fingers C2H2-type | -0.81 |
| ENSG00000164684 | 619279 | PWB-iPSC DN | ZNF704 | Zinc fingers C2H2-type | -0.69 |
| ENSG00000140548 | 374655 | PWB-iPSC DN | ZNF710 | Zinc fingers C2H2-type | -0.72 |
| ENSG00000139651 | 283337 | PWB-iPSC DN | ZNF740 | Zinc fingers C2H2-type | -0.51 |
| ENSG00000130449 | 57688 | PWB-iPSC DN | ZSWIM6 | Zinc fingers SWIM-type | -0.6 |
| ENSG00000173210 | 22885 | PWB-iPSC UP | ABLIM3 | LIM domain containing | 1.08 |
| ENSG00000146215 | 401262 | PWB-iPSC UP | CRIP3 | LIM domain containing | 1.16 |
| ENSG00000163820 | 79443 | PWB-iPSC UP | FYCO1 | Zinc fingers FYVE-type | 0.55 |
| ENSG00000153814 | 221895 | PWB-iPSC UP | JAZF1 | Zinc fingers C2H2-type | 1.07 |
| ENSG00000185513 | 26013 | PWB-iPSC UP | L3MBTL1 | Zinc fingers C2HC-type | 0.95 |
| ENSG00000072163 | 55679 | PWB-iPSC UP | LIMS2 | LIMS zinc finger family | 1.22 |
| ENSG00000048540 | 55885 | PWB-iPSC UP | LMO3 | LIM domain containing | 1.29 |
| ENSG00000145012 | 4026 | PWB-iPSC UP | LPP | Zyxin family | 0.65 |
| ENSG00000145416 | 55016 | PWB-iPSC UP | MARCHF1 | Ring finger proteins | 1.52 |
| ENSG00000015479 | 9782 | PWB-iPSC UP | MATR3 | Zinc fingers matrin-type | 0.71 |
| ENSG00000152601 | 4154 | PWB-iPSC UP | MBNL1 | Zinc fingers CCCH-type | 0.71 |
| ENSG00000198625 | 4194 | PWB-iPSC UP | MDM4 | Ring finger proteins | 0.42 |
| ENSG00000166343 | 118490 | PWB-iPSC UP | MSS51 | Zinc fingers MYND-type | 1.16 |
| ENSG00000186472 | 27445 | PWB-iPSC UP | PCLO | Zinc fingers PCLO-type | 1.07 |
| ENSG00000078043 | 9063 | PWB-iPSC UP | PIAS2 | Zinc fingers MIZ-type | 0.5 |
| ENSG00000178222 | 285498 | PWB-iPSC UP | RNF212 | Ring finger proteins | 1.22 |
| ENSG00000079102 | 862 | PWB-iPSC UP | RUNX1T1 | Zinc fingers MYND-type | 0.74 |
| ENSG00000135899 | 3431 | PWB-iPSC UP | SP110 | PHD finger proteins | 1.32 |
| ENSG00000183718 | 84851 | PWB-iPSC UP | TRIM52 | Ring finger proteins | 0.67 |
| ENSG00000169871 | 81844 | PWB-iPSC UP | TRIM56 | Ring finger proteins | 0.5 |
| ENSG00000100505 | 114088 | PWB-iPSC UP | TRIM9 | Ring finger proteins | 0.79 |
| ENSG00000111424 | 7421 | PWB-iPSC UP | VDR | :lear receptor subfamily 1 gro | 1.82 |
| ENSG00000132846 | 84327 | PWB-iPSC UP | ZBED3 | Zinc fingers BED-type | 1.04 |
| ENSG00000089775 | 7597 | PWB-iPSC UP | ZBTB25 | Zinc fingers C2H2-type | 0.75 |
| ENSG00000185278 | 84614 | PWB-iPSC UP | ZBTB37 | Zinc fingers C2H2-type | 0.82 |
| ENSG00000206077 | 653082 | PWB-iPSC UP | ZDHHC11B | Zinc fingers DHHC-type | 0.92 |
| ENSG00000175893 | 340481 | PWB-iPSC UP | ZDHHC21 | Zinc fingers DHHC-type | 0.61 |
| ENSG00000177108 | 283576 | PWB-iPSC UP | ZDHHC22 | Zinc fingers DHHC-type | 0.9 |
| ENSG00000184307 | 254887 | PWB-iPSC UP | ZDHHC23 | Zinc fingers DHHC-type | 0.64 |
| ENSG00000133858 | 196441 | PWB-iPSC UP | ZFC3H1 | Zinc fingers | 0.54 |
| ENSG00000196670 | 643836 | PWB-iPSC UP | ZFP62 | Zinc fingers C2H2-type | 0.58 |
| ENSG00000106261 | 7586 | PWB-iPSC UP | ZKSCAN1 | Zinc fingers C2H2-type | 0.5 |
| ENSG00000122515 | 83637 | PWB-iPSC UP | ZMIZ2 | Zinc fingers MIZ-type | 0.54 |
| ENSG00000196247 | 51427 | PWB-iPSC UP | ZNF107 | Zinc fingers C2H2-type | 1.11 |
| ENSG00000062370 | 7771 | PWB-iPSC UP | ZNF112 | Zinc fingers C2H2-type | 0.75 |
| ENSG00000197008 | 7697 | PWB-iPSC UP | ZNF138 | Zinc fingers C2H2-type | 0.64 |
| ENSG00000131127 | 7700 | PWB-iPSC UP | ZNF141 | Zinc fingers C2H2-type | 0.71 |
| ENSG00000132010 | 7568 | PWB-iPSC UP | ZNF20 | Zinc fingers C2H2-type | 1.06 |
| ENSG00000160321 | 7757 | PWB-iPSC UP | ZNF208 | Zinc fingers C2H2-type | 2.1 |
| ENSG00000159905 | 7638 | PWB-iPSC UP | ZNF221 | Zinc fingers C2H2-type | 1.55 |
| ENSG00000167380 | 7769 | PWB-iPSC UP | ZNF226 | Zinc fingers C2H2-type | 0.59 |
| ENSG00000196793 | 8187 | PWB-iPSC UP | ZNF239 | Zinc fingers C2H2-type | 1.14 |
| ENSG00000197134 | 113835 | PWB-iPSC UP | ZNF257 | Zinc fingers C2H2-type | 2.47 |
| ENSG00000174652 | 10781 | PWB-iPSC UP | ZNF266 | Zinc fingers C2H2-type | 0.77 |
| ENSG00000182986 | 162967 | PWB-iPSC UP | ZNF320 | Zinc fingers C2H2-type | 0.62 |
| ENSG00000130844 | 55422 | PWB-iPSC UP | ZNF331 | Zinc fingers C2H2-type | 0.61 |
| ENSG00000198185 | 55713 | PWB-iPSC UP | ZNF334 | Zinc fingers C2H2-type | 0.59 |
| ENSG00000196693 | 7582 | PWB-iPSC UP | ZNF33B | Zinc fingers C2H2-type | 0.88 |
| ENSG00000151789 | 79750 | PWB-iPSC UP | ZNF385D | Zinc fingers matrin-type | 0.81 |
| ENSG00000170954 | 55786 | PWB-iPSC UP | ZNF415 | Zinc fingers C2H2-type | 0.87 |
| ENSG00000105136 | 79744 | PWB-iPSC UP | ZNF419 | Zinc fingers C2H2-type | 0.62 |
| ENSG00000197016 | 388566 | PWB-iPSC UP | ZNF470 | Zinc fingers C2H2-type | 0.97 |
| ENSG00000196263 | 57573 | PWB-iPSC UP | ZNF471 | Zinc fingers C2H2-type | 1.07 |
| ENSG00000173258 | 158399 | PWB-iPSC UP | ZNF483 | Zinc fingers C2H2-type | 0.52 |
| ENSG00000229676 | 57615 | PWB-iPSC UP | ZNF492 | Zinc fingers C2H2-type | 6.25 |

| | | | | | |
|---|---|---|---|---|---|
| ENSG00000105559 | 57664 | PWB-EC DN | PLEKHA4 | Yes | -3.18 |
| ENSG00000287080 | 8352 | PWB-EC DN | H3C3 | Yes | -3.36 |
| ENSG00000146122 | 23500 | PWB-EC DN | DAAM2 | Yes | -3.38 |
| ENSG00000111704 | 79923 | PWB-EC DN | NANOG | Yes | -3.46 |
| ENSG00000111087 | 2735 | PWB-EC DN | GLI1 | Yes | -3.52 |
| ENSG00000276410 | 3018 | PWB-EC DN | H2BC3 | Yes | -3.6 |
| ENSG00000102678 | 2254 | PWB-EC DN | FGF9 | Yes | -3.68 |
| ENSG00000155011 | 27123 | PWB-EC DN | DKK2 | Yes | -4.03 |
| ENSG00000183762 | 83999 | PWB-EC DN | KREMEN1 | Yes | -4.44 |
| ENSG00000146648 | 1956 | PWB-EC DN | EGFR | Yes | -4.64 |
| ENSG00000101311 | 55612 | PWB-EC DN | FERMT1 | Yes | -4.65 |
| ENSG00000251493 | 2297 | PWB-EC DN | FOXD1 | Yes | -4.73 |
| ENSG00000188760 | 130612 | PWB-EC DN | TMEM198 | Yes | -4.81 |
| ENSG00000124813 | 860 | PWB-EC DN | RUNX2 | Yes | -4.85 |
| ENSG00000134569 | 4038 | PWB-EC DN | LRP4 | Yes | -4.89 |
| ENSG00000164932 | 115908 | PWB-EC DN | CTHRC1 | Yes | -4.93 |
| ENSG00000145423 | 6423 | PWB-EC DN | SFRP2 | Yes | -5.17 |
| ENSG00000168243 | 2786 | PWB-EC DN | GNG4 | Yes | -5.26 |
| ENSG00000198768 | 164284 | PWB-EC DN | APCDD1L | Yes | -5.38 |
| ENSG00000204531 | 5460 | PWB-EC DN | POU5F1 | Yes | -5.6 |
| ENSG00000151617 | 1909 | PWB-EC DN | EDNRA | Yes | -6.08 |
| ENSG00000115884 | 6382 | PWB-EC DN | SDC1 | Yes | -6.32 |
| ENSG00000125398 | 6662 | PWB-EC DN | SOX9 | Yes | -6.54 |
| ENSG00000198796 | 115701 | PWB-EC DN | ALPK2 | Yes | -6.65 |
| ENSG00000166923 | 26585 | PWB-EC DN | GREM1 | Yes | -7.37 |
| ENSG00000234616 | 8629 | PWB-EC DN | JRK | Yes | -7.62 |
| ENSG00000108821 | 1277 | PWB-EC DN | COL1A1 | Yes | -8.23 |
| ENSG00000212993 | 5462 | PWB-MSC UP | POU5F1B | Yes | 2.35 |
| ENSG00000188906 | 120892 | PWB-MSC UP | LRRK2 | Yes | 1.89 |
| ENSG00000105974 | 857 | PWB-MSC UP | CAV1 | Yes | 1.41 |
| ENSG00000168243 | 2786 | PWB-MSC UP | GNG4 | Yes | 1.28 |
| ENSG00000132846 | 84327 | PWB-MSC UP | ZBED3 | Yes | 1.12 |
| ENSG00000064195 | 1747 | PWB-MSC UP | DLX3 | Yes | 1 |
| ENSG00000146648 | 1956 | PWB-MSC UP | EGFR | Yes | 0.99 |
| ENSG00000157895 | 64897 | PWB-MSC UP | C12orf43 | Yes | 0.95 |
| ENSG00000123349 | 5204 | PWB-MSC UP | PFDN5 | Yes | 0.81 |
| ENSG00000167779 | 3489 | PWB-MSC UP | IGFBP6 | Yes | 0.79 |
| ENSG00000167548 | 8085 | PWB-MSC UP | KMT2D | Yes | 0.75 |
| ENSG00000111640 | 2597 | PWB-MSC UP | GAPDH | Yes | 0.64 |
| ENSG00000102882 | 5595 | PWB-MSC UP | MAPK3 | Yes | 0.61 |
| ENSG00000111142 | 10988 | PWB-MSC UP | METAP2 | Yes | 0.59 |
| ENSG00000143842 | 9580 | PWB-MSC UP | SOX13 | Yes | 0.52 |
| ENSG00000149948 | 8091 | PWB-MSC UP | HMGA2 | Yes | 0.48 |
| ENSG00000060237 | 65125 | PWB-MSC UP | WNK1 | Yes | 0.48 |
| ENSG00000182197 | 2131 | PWB-MSC UP | EXT1 | Yes | 0.45 |
| ENSG00000277157 | 8360 | PWB-MSC DN | H4C4 | Yes | -0.54 |
| ENSG00000286522 | 8358 | PWB-MSC DN | H3C2 | Yes | -0.55 |
| ENSG00000138032 | 5495 | PWB-MSC DN | PPM1B | Yes | -0.56 |
| ENSG00000127955 | 2770 | PWB-MSC DN | GNAI1 | Yes | -0.58 |
| ENSG00000138738 | 11107 | PWB-MSC DN | PRDM5 | Yes | -0.59 |
| ENSG00000274618 | 8361 | PWB-MSC DN | H4C6 | Yes | -0.63 |
| ENSG00000050165 | 27122 | PWB-MSC DN | DKK3 | Yes | -0.63 |
| ENSG00000138814 | 5530 | PWB-MSC DN | PPP3CA | Yes | -0.65 |
| ENSG00000105856 | 26959 | PWB-MSC DN | HBP1 | Yes | -0.71 |
| ENSG00000122707 | 8434 | PWB-MSC DN | RECK | Yes | -0.81 |
| ENSG00000058091 | 5218 | PWB-MSC DN | CDK14 | Yes | -0.83 |
| ENSG00000150995 | 3708 | PWB-MSC DN | ITPR1 | Yes | -0.96 |
| ENSG00000109339 | 5602 | PWB-MSC DN | MAPK10 | Yes | -0.98 |
| ENSG00000100968 | 4776 | PWB-MSC DN | NFATC4 | Yes | -1.01 |
| ENSG00000136160 | 1910 | PWB-MSC DN | EDNRB | Yes | -1.01 |
| ENSG00000070193 | 2255 | PWB-MSC DN | FGF10 | Yes | -1.04 |
| ENSG00000182621 | 23236 | PWB-MSC DN | PLCB1 | Yes | -1.05 |
| ENSG00000275126 | 8368 | PWB-MSC DN | H4C13 | Yes | -1.07 |
| ENSG00000105559 | 57664 | PWB-MSC DN | PLEKHA4 | Yes | -1.19 |
| ENSG00000127920 | 2791 | PWB-MSC DN | GNG11 | Yes | -1.31 |
| ENSG00000108821 | 1277 | PWB-MSC DN | COL1A1 | Yes | -1.42 |
| ENSG00000140682 | 7041 | PWB-MSC DN | TGFB1I1 | Yes | -1.42 |
| ENSG00000276410 | 3018 | PWB-MSC DN | H2BC3 | Yes | -1.55 |
| ENSG00000155011 | 27123 | PWB-MSC DN | DKK2 | Yes | -1.65 |
| ENSG00000105672 | 2116 | PWB-MSC DN | ETV2 | Yes | -1.67 |
| ENSG00000198796 | 115701 | PWB-MSC DN | ALPK2 | Yes | -1.68 |
| ENSG00000154310 | 23043 | PWB-MSC DN | TNIK | Yes | -1.71 |
| ENSG00000287080 | 8352 | PWB-MSC DN | H3C3 | Yes | -1.72 |

| | | | | | |
|---|---|---|---|---|---|
| ENSG00000196268 | 284443 | PWB-iPSC UP | ZNF493 | Zinc fingers C2H2-type | 0.73 |
| ENSG00000203326 | 170958 | PWB-iPSC UP | ZNF525 | Zinc fingers C2H2-type | 1.4 |
| ENSG00000167555 | 84436 | PWB-iPSC UP | ZNF528 | Zinc fingers C2H2-type | 1.12 |
| ENSG00000198633 | 147658 | PWB-iPSC UP | ZNF534 | Zinc fingers C2H2-type | 1.7 |
| ENSG00000167785 | 148156 | PWB-iPSC UP | ZNF558 | Zinc fingers C2H2-type | 1.43 |
| ENSG00000258405 | 147660 | PWB-iPSC UP | ZNF578 | Zinc fingers C2H2-type | 2.43 |
| ENSG00000160229 | 7617 | PWB-iPSC UP | ZNF66 | Zinc fingers C2H2-type | 1.21 |
| ENSG00000197497 | 79788 | PWB-iPSC UP | ZNF665 | Zinc fingers C2H2-type | 0.87 |
| ENSG00000196109 | 163223 | PWB-iPSC UP | ZNF676 | Zinc fingers C2H2-type | 6.05 |
| ENSG00000181450 | 339500 | PWB-iPSC UP | ZNF678 | Zinc fingers C2H2-type | 0.46 |
| ENSG00000173041 | 340252 | PWB-iPSC UP | ZNF680 | Zinc fingers C2H2-type | 0.73 |
| ENSG00000167562 | 55762 | PWB-iPSC UP | ZNF701 | Zinc fingers C2H2-type | 0.9 |
| ENSG00000120963 | 51123 | PWB-iPSC UP | ZNF706 | Zinc fingers C2H2-type | 0.57 |
| ENSG00000227124 | 1E+08 | PWB-iPSC UP | ZNF717 | Zinc fingers C2H2-type | 0.82 |
| ENSG00000268696 | 646864 | PWB-iPSC UP | ZNF723 | Zinc fingers C2H2-type | 8.9 |
| ENSG00000213967 | 730087 | PWB-iPSC UP | ZNF726 | Zinc fingers C2H2-type | 1.49 |
| ENSG00000269067 | 388523 | PWB-iPSC UP | ZNF728 | Zinc fingers C2H2-type | 2.89 |
| ENSG00000196350 | 1E+08 | PWB-iPSC UP | ZNF729 | Zinc fingers C2H2-type | 11.14 |
| ENSG00000186777 | 654254 | PWB-iPSC UP | ZNF732 | Zinc fingers C2H2-type | 0.51 |
| ENSG00000234444 | 728927 | PWB-iPSC UP | ZNF736 | Zinc fingers C2H2-type | 1.58 |
| ENSG00000237440 | 1E+08 | PWB-iPSC UP | ZNF737 | Zinc fingers C2H2-type | 0.81 |
| ENSG00000160336 | 388561 | PWB-iPSC UP | ZNF761 | Zinc fingers C2H2-type | 0.45 |
| ENSG00000196417 | 91661 | PWB-iPSC UP | ZNF765 | Zinc fingers C2H2-type | 0.66 |
| ENSG00000128000 | 163131 | PWB-iPSC UP | ZNF780B | Zinc fingers C2H2-type | 0.99 |
| ENSG00000198482 | 388558 | PWB-iPSC UP | ZNF808 | Zinc fingers C2H2-type | 0.97 |
| ENSG00000198346 | 126017 | PWB-iPSC UP | ZNF813 | Zinc fingers C2H2-type | 0.86 |
| ENSG00000151612 | 152485 | PWB-iPSC UP | ZNF827 | Zinc fingers C2H2-type | 0.62 |
| ENSG00000167766 | 55769 | PWB-iPSC UP | ZNF83 | Zinc fingers C2H2-type | 0.61 |
| ENSG00000197608 | 284371 | PWB-iPSC UP | ZNF841 | Zinc fingers C2H2-type | 0.96 |
| ENSG00000197385 | 344787 | PWB-iPSC UP | ZNF860 | Zinc fingers C2H2-type | 1.21 |
| ENSG00000221923 | 400713 | PWB-iPSC UP | ZNF880 | Zinc fingers C2H2-type | 0.94 |
| ENSG00000214029 | 1.01E+08 | PWB-iPSC UP | ZNF891 | Zinc fingers C2H2-type | 1.36 |
| ENSG00000213988 | 7643 | PWB-iPSC UP | ZNF90 | Zinc fingers C2H2-type | 0.55 |
| ENSG00000197360 | 148198 | PWB-iPSC UP | ZNF98 | Zinc fingers C2H2-type | 8.25 |
| ENSG00000213973 | 7652 | PWB-iPSC UP | ZNF99 | Zinc fingers C2H2-type | 11.12 |
| ENSG00000152467 | 284312 | PWB-iPSC UP | ZSCAN1 | Zinc fingers C2H2-type | 2.12 |
| ENSG00000121413 | 65982 | PWB-iPSC UP | ZSCAN18 | Zinc fingers C2H2-type | 0.6 |
| ENSG00000196350 | 1E+08 | PWB-MSC UP | ZNF729 | Zinc fingers C2H2-type | 10.64 |
| ENSG00000268696 | 646864 | PWB-MSC UP | ZNF723 | Zinc fingers C2H2-type | 9.95 |
| ENSG00000197360 | 148198 | PWB-MSC UP | ZNF98 | Zinc fingers C2H2-type | 8.87 |
| ENSG00000213973 | 7652 | PWB-MSC UP | ZNF99 | Zinc fingers C2H2-type | 8.7 |
| ENSG00000229676 | 57615 | PWB-MSC UP | ZNF492 | Zinc fingers C2H2-type | 8.44 |
| ENSG00000269067 | 388523 | PWB-MSC UP | ZNF728 | Zinc fingers C2H2-type | 3.62 |
| ENSG00000197134 | 113835 | PWB-MSC UP | ZNF257 | Zinc fingers C2H2-type | 3 |
| ENSG00000258405 | 147660 | PWB-MSC UP | ZNF578 | Zinc fingers C2H2-type | 2.82 |
| ENSG00000160321 | 7757 | PWB-MSC UP | ZNF208 | Zinc fingers C2H2-type | 2.41 |
| ENSG00000234444 | 728927 | PWB-MSC UP | ZNF736 | Zinc fingers C2H2-type | 2.28 |
| ENSG00000100505 | 114088 | PWB-MSC UP | TRIM9 | Ring finger proteins | 1.9 |
| ENSG00000278318 | 7772 | PWB-MSC UP | ZNF229 | Zinc fingers C2H2-type | 1.85 |
| ENSG00000198028 | 147741 | PWB-MSC UP | ZNF560 | Zinc fingers C2H2-type | 1.68 |
| ENSG00000146215 | 401262 | PWB-MSC UP | CRIP3 | LIM domain containing | 1.39 |
| ENSG00000127084 | 89846 | PWB-MSC UP | FGD3 | Zinc fingers FYVE-type | 1.37 |
| ENSG00000167555 | 84436 | PWB-MSC UP | ZNF528 | Zinc fingers C2H2-type | 1.27 |
| ENSG00000179059 | 132625 | PWB-MSC UP | ZFP42 | Zinc fingers C2H2-type | 1.22 |
| ENSG00000172818 | 5017 | PWB-MSC UP | OVOL1 | Zinc fingers C2H2-type | 1.14 |
| ENSG00000132846 | 84327 | PWB-MSC UP | ZBED3 | Zinc fingers BED-type | 1.12 |
| ENSG00000130182 | 84891 | PWB-MSC UP | ZSCAN10 | Zinc fingers C2H2-type | 1.12 |
| ENSG00000196132 | 4661 | PWB-MSC UP | MYT1 | Zinc fingers C2H2C-type | 1.1 |
| ENSG00000196653 | 91392 | PWB-MSC UP | ZNF502 | Zinc fingers C2H2-type | 0.98 |
| ENSG00000139174 | 144165 | PWB-MSC UP | PRICKLE1 | LIM domain containing | 0.91 |
| ENSG00000197124 | 91120 | PWB-MSC UP | ZNF682 | Zinc fingers C2H2-type | 0.9 |
| ENSG00000198040 | 7637 | PWB-MSC UP | ZNF84 | Zinc fingers C2H2-type | 0.88 |
| ENSG00000197279 | 7718 | PWB-MSC UP | ZNF165 | Zinc fingers C2H2-type | 0.87 |
| ENSG00000221923 | 400713 | PWB-MSC UP | ZNF880 | Zinc fingers C2H2-type | 0.87 |
| ENSG00000204186 | 57683 | PWB-MSC UP | ZDBF2 | Zinc fingers DBF-type | 0.81 |
| ENSG00000180855 | 10224 | PWB-MSC UP | ZNF443 | Zinc fingers C2H2-type | 0.8 |
| ENSG00000198393 | 7574 | PWB-MSC UP | ZNF26 | Zinc fingers C2H2-type | 0.79 |
| ENSG00000065029 | 7629 | PWB-MSC UP | ZNF76 | Zinc fingers C2H2-type | 0.76 |
| ENSG00000213988 | 7643 | PWB-MSC UP | ZNF90 | Zinc fingers C2H2-type | 0.76 |
| ENSG00000167548 | 8085 | PWB-MSC UP | KMT2D | PHD finger proteins | 0.75 |
| ENSG00000172748 | 169270 | PWB-MSC UP | ZNF596 | Zinc fingers C2H2-type | 0.73 |
| ENSG00000173041 | 340252 | PWB-MSC UP | ZNF680 | Zinc fingers C2H2-type | 0.73 |
| ENSG00000198093 | 65251 | PWB-MSC UP | ZNF649 | Zinc fingers C2H2-type | 0.71 |

| | | | | | |
|---|---|---|---|---|---|
| ENSG00000152804 | 3087 | PWB-MSC DN | HHEX | Yes | -1.74 |
| ENSG00000104415 | 8840 | PWB-MSC DN | CCN4 | Yes | -1.88 |
| ENSG00000171243 | 25928 | PWB-MSC DN | SOSTDC1 | Yes | -1.89 |
| ENSG00000120332 | 63923 | PWB-MSC DN | TNN | Yes | -1.97 |
| ENSG00000166501 | 5579 | PWB-MSC DN | PRKCB | Yes | -2.03 |
| ENSG00000171056 | 83595 | PWB-MSC DN | SOX7 | Yes | -2.27 |
| ENSG00000136826 | 9314 | PWB-MSC DN | KLF4 | Yes | -2.29 |
| ENSG00000107984 | 22943 | PWB-MSC DN | DKK1 | Yes | -2.57 |
| ENSG00000101384 | 182 | PWB-MSC DN | JAG1 | Yes | -2.61 |
| ENSG00000234616 | 8629 | PWB-MSC DN | JRK | Yes | -9.21 |

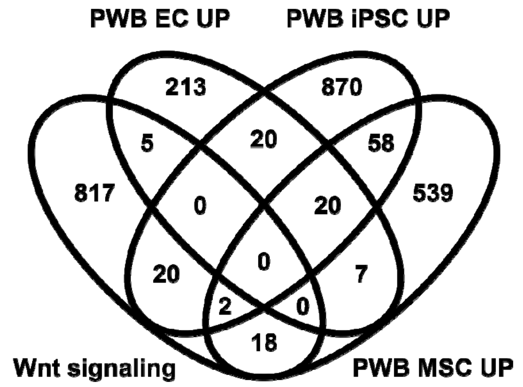


5 DEGs in "Wnt signaling" and "PWB EC UP":
GPRC5B, SIX3, SULF2, TNFAIP3, TPBGL

20 DEGs in "Wnt signaling" and "PWB iPSC UP":
CAPRIN2, CER1, DACT1, DLG2, EGF, H2BC1, ITPR1, ITPR2, MACF1, PKD1, PLCB2
PRKCG, PSEN2, PSMA2, RBMS3, TCF7, TERT, TNRC6A, WIF1, WNT8B

2 DEGs in "Wnt signaling", "PWB iPSC UP" and "PWB MSC UP": CAV1, ZBED3

18 DEGs in "Wnt signaling" and "PWB MSC UP":
C12orf43, DLX3, EGFR, EXT1, GAPDH, GNG4, HMGA2, IGFBP6, KMT2D, LRRK2
MAPK3, METAP2, PFDN5, POU5F1B, PRICKLE1, SOX13, WNK1, WNT3A

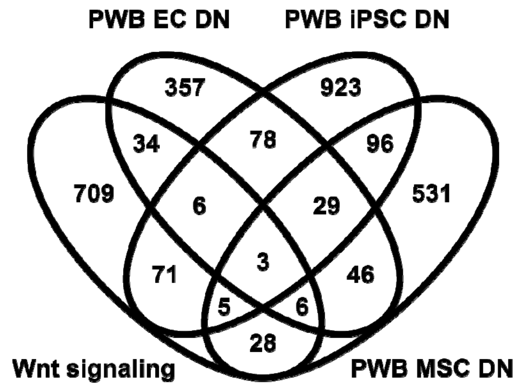


6 common DEGs in "Wnt signaling", "PWB EC DN" and "PWB iPSC DN":
WNT5B, SOX9, SOX2, NKD1, LGR4, GREM1

5 common DEGs in "Wnt signaling", "PWB iPSC DN" and "PWB MSC DN":
GNAI1, HHEX, JAG1, KLF4, MAPK10

6 common DEGs in "Wnt signaling", "PWB EC DN" and "PWB MSC DN":
ALPK2, DKK1, DKK2, H2BC3, H3C3, JRK

| | | | | | |
|---|---|---|---|---|---|
| ENSG00000275023 | 4302 | PWB-MSC UP | MLLT6 | PHD finger proteins | 0.7 |
| ENSG00000090612 | 10795 | PWB-MSC UP | ZNF268 | Zinc fingers C2H2-type | 0.7 |
| ENSG00000089280 | 2521 | PWB-MSC UP | FUS | Zinc fingers RANBP2-type | 0.69 |
| ENSG00000196458 | 1E+08 | PWB-MSC UP | ZNF605 | Zinc fingers C2H2-type | 0.68 |
| ENSG00000162086 | 7627 | PWB-MSC UP | ZNF75A | Zinc fingers C2H2-type | 0.65 |
| ENSG00000156650 | 23522 | PWB-MSC UP | KAT6B | Zinc fingers C2HC-type | 0.64 |
| ENSG00000110851 | 11108 | PWB-MSC UP | PRDM4 | Zinc fingers C2H2-type | 0.63 |
| ENSG00000196247 | 51427 | PWB-MSC UP | ZNF107 | Zinc fingers C2H2-type | 0.63 |
| ENSG00000256223 | 7556 | PWB-MSC UP | ZNF10 | Zinc fingers C2H2-type | 0.62 |
| ENSG00000270647 | 8148 | PWB-MSC UP | TAF15 | Zinc fingers RANBP2-type | 0.61 |
| ENSG00000198464 | 147657 | PWB-MSC UP | ZNF480 | Zinc fingers C2H2-type | 0.61 |
| ENSG00000179195 | 144348 | PWB-MSC UP | ZNF664 | Zinc fingers C2H2-type | 0.61 |
| ENSG00000269343 | 1E+08 | PWB-MSC UP | ZNF587B | Zinc fingers C2H2-type | 0.6 |
| ENSG00000167562 | 55762 | PWB-MSC UP | ZNF701 | Zinc fingers C2H2-type | 0.6 |
| ENSG00000180263 | 55785 | PWB-MSC UP | FGD6 | Zinc fingers FYVE-type | 0.59 |
| ENSG00000237440 | 1E+08 | PWB-MSC UP | ZNF737 | Zinc fingers C2H2-type | 0.59 |
| ENSG00000126746 | 171017 | PWB-MSC UP | ZNF384 | Zinc fingers C2H2-type | 0.56 |
| ENSG00000198482 | 388558 | PWB-MSC UP | ZNF808 | Zinc fingers C2H2-type | 0.56 |
| ENSG00000196172 | 148213 | PWB-MSC UP | ZNF681 | Zinc fingers C2H2-type | 0.55 |
| ENSG00000072609 | 55743 | PWB-MSC UP | CHFR | Ring finger proteins | 0.54 |
| ENSG00000076770 | 55796 | PWB-MSC UP | MBNL3 | Zinc fingers CCCH-type | 0.54 |
| ENSG00000089094 | 84678 | PWB-MSC UP | KDM2B | Zinc fingers CXXC-type | 0.52 |
| ENSG00000070047 | 57661 | PWB-MSC UP | PHRF1 | Ring finger proteins | 0.51 |
| ENSG00000182986 | 162967 | PWB-MSC UP | ZNF320 | Zinc fingers C2H2-type | 0.5 |
| ENSG00000139651 | 283337 | PWB-MSC UP | ZNF740 | Zinc fingers C2H2-type | 0.49 |
| ENSG00000089234 | 8315 | PWB-MSC UP | BRAP | Ring finger proteins | 0.47 |
| ENSG00000147548 | 54904 | PWB-MSC DN | NSD3 | PHD finger proteins | -0.49 |
| ENSG00000146373 | 154214 | PWB-MSC DN | RNF217 | Ring finger proteins | -0.49 |
| ENSG00000177463 | 7182 | PWB-MSC DN | NR2C2 | lear receptor subfamily 2 gro | -0.5 |
| ENSG00000105866 | 6671 | PWB-MSC DN | SP4 | Zinc fingers C2H2-type | -0.53 |
| ENSG00000066422 | 27107 | PWB-MSC DN | ZBTB11 | Zinc fingers C2H2-type | -0.54 |
| ENSG00000172667 | 64393 | PWB-MSC DN | ZMAT3 | Zinc fingers matrin-type | -0.56 |
| ENSG00000156831 | 286053 | PWB-MSC DN | NSMCE2 | Zinc fingers MIZ-type | -0.57 |
| ENSG00000181690 | 5324 | PWB-MSC DN | PLAG1 | Zinc fingers C2H2-type | -0.59 |
| ENSG00000138738 | 11107 | PWB-MSC DN | PRDM5 | Zinc fingers C2H2-type | -0.59 |
| ENSG00000078114 | 10529 | PWB-MSC DN | NEBL | LIM domain containing | -0.6 |
| ENSG00000101752 | 57534 | PWB-MSC DN | MIB1 | Zinc fingers ZZ-type | -0.63 |
| ENSG00000109787 | 51274 | PWB-MSC DN | KLF3 | Zinc fingers C2H2-type | -0.64 |
| ENSG00000188321 | 84527 | PWB-MSC DN | ZNF559 | Zinc fingers C2H2-type | -0.68 |
| ENSG00000147394 | 7739 | PWB-MSC DN | ZNF185 | LIM domain containing | -0.72 |
| ENSG00000176641 | 220441 | PWB-MSC DN | RNF152 | Ring finger proteins | -0.74 |
| ENSG00000177932 | 30832 | PWB-MSC DN | ZNF354C | Zinc fingers C2H2-type | -0.81 |
| ENSG00000161298 | 84911 | PWB-MSC DN | ZNF382 | Zinc fingers C2H2-type | -0.83 |
| ENSG00000121236 | 117854 | PWB-MSC DN | TRIM6 | Ring finger proteins | -0.84 |
| ENSG00000198795 | 25925 | PWB-MSC DN | ZNF521 | Zinc fingers C2H2-type | -0.86 |
| ENSG00000275004 | 140883 | PWB-MSC DN | ZNF280B | Zinc fingers C2H2-type | -0.89 |
| ENSG00000105278 | 23217 | PWB-MSC DN | ZFR2 | Zinc fingers matrin-type | -0.93 |
| ENSG00000022267 | 2273 | PWB-MSC DN | FHL1 | LIM domain containing | -0.97 |
| ENSG00000197566 | 57547 | PWB-MSC DN | ZNF624 | Zinc fingers C2H2-type | -0.98 |
| ENSG00000186951 | 5465 | PWB-MSC DN | PPARA | ome proliferator activated re | -1.01 |
| ENSG00000079102 | 862 | PWB-MSC DN | RUNX1T1 | Zinc fingers MYND-type | -1.07 |
| ENSG00000175691 | 58492 | PWB-MSC DN | ZNF77 | Zinc fingers C2H2-type | -1.07 |
| ENSG00000154553 | 27295 | PWB-MSC DN | PDLIM3 | LIM domain containing | -1.14 |
| ENSG00000198939 | 80108 | PWB-MSC DN | ZFP2 | Zinc fingers C2H2-type | -1.16 |
| ENSG00000154783 | 152273 | PWB-MSC DN | FGD5 | Zinc fingers FYVE-type | -1.31 |
| ENSG00000163357 | 149095 | PWB-MSC DN | DCST1 | Ring finger proteins | -1.36 |
| ENSG00000171970 | 126295 | PWB-MSC DN | ZNF57 | Zinc fingers C2H2-type | -1.38 |
| ENSG00000140682 | 7041 | PWB-MSC DN | TGFB1I1 | LIM domain containing | -1.42 |
| ENSG00000187556 | 342977 | PWB-MSC DN | NANOS3 | Zinc fingers C2HC-type | -1.57 |
| ENSG00000102385 | 1821 | PWB-MSC DN | DRP2 | Zinc fingers ZZ-type | -1.61 |
| ENSG00000135363 | 4005 | PWB-MSC DN | LMO2 | LIM domain containing | -1.7 |
| ENSG00000250571 | 2738 | PWB-MSC DN | GLI4 | Zinc fingers C2H2-type | -1.71 |
| ENSG00000147573 | 84675 | PWB-MSC DN | TRIM55 | Ring finger proteins | -1.71 |
| ENSG00000186300 | 148254 | PWB-MSC DN | ZNF555 | Zinc fingers C2H2-type | -1.72 |
| ENSG00000169297 | 190 | PWB-MSC DN | NR0B1 | lear receptor subfamily 0 gro | -1.74 |
| ENSG00000061455 | 93166 | PWB-MSC DN | PRDM6 | Zinc fingers C2H2-type | -1.77 |
| ENSG00000075407 | 7587 | PWB-MSC DN | ZNF37A | Zinc fingers C2H2-type | -1.78 |
| ENSG00000162676 | 2672 | PWB-MSC DN | GFI1 | Zinc fingers C2H2-type | -2.04 |
| ENSG00000198205 | 7789 | PWB-MSC DN | ZXDA | Zinc fingers C2H2-type | -2.06 |
| ENSG00000136826 | 9314 | PWB-MSC DN | KLF4 | Zinc fingers C2H2-type | -2.29 |
| ENSG00000161940 | 255877 | PWB-MSC DN | BCL6B | Zinc fingers C2H2-type | -2.47 |
| ENSG00000128253 | 10739 | PWB-MSC DN | RFPL2 | Ring finger proteins | -3.26 |
| ENSG00000178187 | 285676 | PWB-MSC DN | ZNF454 | Zinc fingers C2H2-type | -6.13 |

| | | | | | |
|---|---|---|---|---|---|
| ENSG00000181638 | 286128 | PWB-MSC DN | ZFP41 | Zinc fingers C2H2-type | -7.67 |
| ENSG00000198105 | 57209 | PWB-MSC DN | ZNF248 | Zinc fingers C2H2-type | -9 |

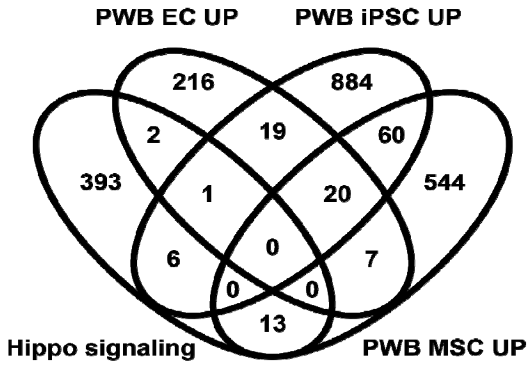


2 common DEGs in "Hippo signaling" and "PWB EC UP": BIRC3, SERPINB3

1 common DEG in "Hippo signaling", "PWB EC UP" and "PWB iPSC UP": A2ML1

6 common DEGs in "Hippo signaling" and "PWB iPSC UP":
GDF5, TCF7, WNT8B, DLG2, GUCY1A2, RBFA

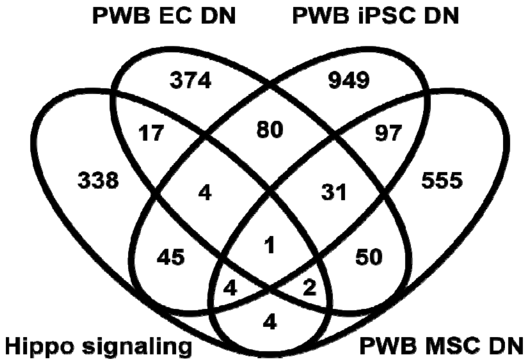


4 common DEGs in "Hippo signaling", "PWB EC DN" and "PWB iPSC DN":
CRB2, NKD1, SOX2, WNT5B

4 common DEGs in "Hippo signaling", "PWB iPSC DN" and "PWB MSC DN":
DCHS1, PARD6G, PPP1R18, SOX11

2 common DEGs in "Hippo signaling", "PWB EC DN" and "PWB MSC DN": GUCY1B1, TGF[illegible]

17 common DEGs in "Hippo signaling" and "PWB EC DN":
BMPR1A, BMPR1B, DLG3, DLG5, DOCK7, EPB41L3, FGF1, FRMD5
FZD7, GDF5, GDF6, GLI2, LEF1, PTPN13, SNAI2, STK26, TEAD3

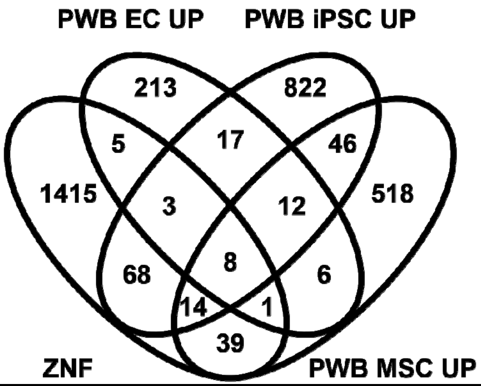


8 common DEGs in "ZNF", "PWB EC UP", "PWB iPSC UP" and "PWB MSC UP":
ZNF208, ZNF257, ZNF528, ZNF578, ZNF728, ZNF729, ZNF98, ZNF99

3 common DEGs in "ZNF", "PWB EC UP" and "PWB iPSC UP": ZNF676, ZSCAN1, ZDHHC21

1 common DEG in "ZNF", "PWB EC UP" and "PWB MSC UP": ZNF560

14 common DEGs in "ZNF", "PWB iPSC UP" and "PWB MSC UP":
CRIP3, TRIM9, ZBED3, ZNF107, ZNF320, ZNF492, ZNF680, ZNF701, ZNF723,
ZNF736, ZNF737, ZNF808, ZNF880, ZNF90

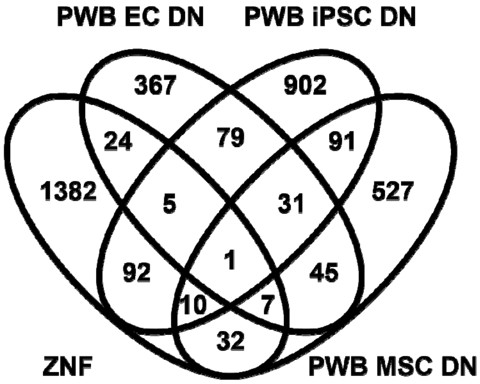


1 common DEGs in "ZNF", "PWB EC DN", "PWB iPSC DN" and "PWB MSC DN": ZNF248

5 common DEGs in "ZNF", "PWB EC DN" and "PWB iPSC DN":
RNF157, ZIC2, ZIC5, ZIC1, ZNF536

10 common DEGs in "ZNF", "PWB iPSC DN" and "PWB MSC DN":
ZNF185, RNF152, KLF3, KLF4, ZNF280B, ZNF521, ZNF555, BCL6B, ZFP2, ZFR2

7 common DEGs in "ZNF", "PWB EC DN" and "PWB MSC DN":
FHL1, PDLIM3, ZNF37A, ZNF454, ZFP41, ZXDA, DRP2

#### Supplementary File 3: Functional enrichment network of DEGs

**1. Functional enrichment network for all DEGs**

| S.No. | Category | ID | Title (or Source) | PWB-EC UP_GeneSet | PWB-EC DN_GeneSet | PWB-iPSC DN_GeneSet | PWB-MSC UP_GeneSet | PWB-iPSC UP_GeneSet | PWB-MSC DN_GeneSet | PWB-EC UP(-logP) | PWB-EC DN(-logP) | PWB-iPSC DN(-logP) | PWB-MSC UP(-logP) | PWB-iPSC UP(-logP) | PWB-MSC DN(-logP) |
|---|---|---|---|---|---|---|---|---|---|---|---|---|---|---|---|
| 31 | GO: Molecular Function | GO:0015278 | calcium-release channel activity | | | | | ITPR1,ITPR2,MCOLN2,RYR1,RYR2,TPCN1,TPCN2 | | 0 | 0 | 0 | 0 | 5.8898574 | 0 |
| 112 | GO: Biological Process | GO:0001944 | vasculature development | ABCC8,ADAMTS18,ANGPT2,CCR3,CD40,CTSZ,CXCL1,CXCL12,CXCL8,EPHB1,GPRC5B,HPSE,IGF1,IL1A,IL1B,LIPA,LPA,MCAM,MIR10B,PGF,PSG4,PSG9,SELE,TDGF1,TDGF1P3,TINAGL1,TNF,TNFAIP3 | ADAM12,ANGPT1,APLNR,BMPR1A,COL1A1,COL1A2,COL3A1,COL5A1,CRB2,CREB3L1,CSPG4,CXXC5,CYP1B1,DCBLD2,DCN,E2F7,EDNRA,EGFR,EGR1,EMILIN1,EMILIN2,ENOX1,EPHB2,FAP,FGF1,FGF9,FOXF2,GDF6,GREM1,HAS2,HMGA2,KREMEN1,LEF1,LGALS1,LGALS3,LIF,LOX,LRP1,MMP19,MMP9,MT1E,MT1X,MT2A,MYOCD,NOTCH2,NRXN3,NTRK2,OSR1,PDGFRA,PDGFRB,PDPN,PITPNM3,PLCD3,PPARG,PRRX1,PRRX2,PTGER4,PTHLH,PTN,PTPRJ,RUNX1,SDC2,SEMA3A,SEMA3C,SERPINB7,SFRP2,SIX1,SMO,SOCS3,SPHK1,STRA6,TAFA5,TGFB2,TGFBI,THBS2,TNFRSF19,TNFSF4,VAV2,VSTM4,WNT5B | ABCA1,ABL1,ACTA2,ADGRG1,ADRA2B,ALDH1A2,AMOT,APOE,ATP2B4,BASP1,BMP7,BTG1,C3,CCL5,CHD7,CITED2,CLIC4,CNMD,COL1A1,COL3A1,CRB2,CRKL,CTNNB1,CXCL10,CXCR4,CYP26A1,CYP26C1,DLX3,EFNB2,ENPP2,EPHB2,EPHB3,ERAP1,FBN2,FBXW11,FERMT2,FGFR2,FLCN,FOLR1,FRZB,FZD5,FZD8,GAA,GLI3,GPC3,GREM1,GRN,HDAC5,HDAC9,HES1,HHEX,HK2,HS6ST2,HSPB1,ID1,IGFBP2,ITGAX,ITGB1BP1,ITGB8,JAG1,JCAD,JUNB,KIT,KLF4,LAMA1,LGALS1,LLGL2,LOXL1,LOXL2,MARCKSL1,MB,MDK,MMP14,NFKB2,NR2F1,NR2F2,NRCAM,NTN1,PAK4,PAX6,PIK3R3,PLCD3,PNPLA6,PRICKLE1,PRKCA,PRKCZ,PTH1R,PTN,RAB11A,RALB,RHOB,ROBO1,RSPO1,RXRA,SASH1,SERPINF1,SERPINF2,SFRP1,SH3KBP1,SIRT1,SNX17,SOX11,SPHK2,STAT1,SUFU,TGFBI,TGFBR1,THBS4,TNFAIP3,TNFRSF19,TNMD,TNNI3,TWSG1,VASH1,VCAM1,WNT4,WNT5B,YAP1 | | | ADAMTS3,ADAMTS6,ADGRF5,AKT3,APLN,BMP10,BMPER,C1GALT1,C3,CALCRL,CCL2,CCN1,CDH13,CDH5,CEACAM1,CFC1,CFC1B,CITED2,COL1A1,COL1A2,COL3A1,CREB3L1,CSPG4,CTH,CUL7,CYP1B1,CYSLTR2,EMILIN1,ERG,ESM1,ETV2,FAM43B,FBN2,FGF10,FLI1,FOXC2,GADD45A,GPR4,HAND2,HAPLN1,HDAC9,HGF,HHEX,HIF3A,HLA-F,IL1A,ITGAV,ITGB1,ITGB1BP1,JAG1,KDR,KLF4,LDB2,LGALS1,LMO2,LOX,LOXL2,LTBP1,MEIS1,MIB1,NFATC4,PGF,PLK2,PPFIA3,PRKCB,PROX1,RASIP1,RECK,RHOJ,RRAS,S1PR1,SDC2,SELE,SERPINF2,SLC4A7,SMOC2,SOX11,SOX7,SRPX2,SVEP1,TAL1,TGFB2,THSD7A,TMEM100,TNFRSF1B,TNFSF4,TNN,TRPC1,TSPAN12,VAV3,ZAP70 | 4.38478 | 10 | 10 | 0 | 0 | 10 |
| 113 | GO: Biological Process | GO:0048514 | blood vessel morphogenesis | ABCC8,ADAMTS18,ANGPT2,CCR3,CD40,CTSZ,CXCL1,CXCL12,CXCL8,EPHB1,HPSE,IL1A,IL1B,LPA,MCAM,MIR10B,PGF,PSG4,PSG9,TDGF1,TDGF1P3,TINAGL1,TNF,TNFAIP3 | ADAM12,ANGPT1,APLNR,BMPR1A,COL3A1,CRB2,CREB3L1,CSPG4,CXXC5,CYP1B1,DCBLD2,DCN,E2F7,EDNRA,EGFR,EMILIN1,EMILIN2,EPHB2,FAP,FGF1,FGF9,GDF6,GREM1,HAS2,HMGA2,KREMEN1,LEF1,LGALS1,LGALS3,LIF,LOX,LRP1,MMP19,MMP9,MT1E,MT1X,MT2A,MYOCD,NOTCH2,NRXN3,NTRK2,PDGFRA,PDGFRB,PLCD3,PPARG,PRRX1,PRRX2,PTGER4,PTN,PTPRJ,RUNX1,SDC2,SEMA3A,SFRP2,SIX1,SMO,SPHK1,STRA6,TAFA5,TGFB2,TGFBI,THBS2,TNFRSF19,VAV2,VSTM4,WNT5B | ABCA1,ABL1,ADGRG1,ADRA2B,AMOT,APOE,ATP2B4,BASP1,BTG1,C3,CCL5,CHD7,CITED2,CLIC4,CNMD,COL3A1,CRB2,CTNNB1,CXCL10,CXCR4,EFNB2,ENPP2,EPHB2,EPHB3,ERAP1,FBN2,FBXW11,FERMT2,FGFR2,FLCN,FOLR1,FZD5,FZD8,GREM1,GRN,HDAC5,HDAC9,HES1,HHEX,HK2,HS6ST2,HSPB1,ID1,IGFBP2,ITGAX,ITGB1BP1,ITGB8,JAG1,JCAD,JUNB,KLF4,LAMA1,LGALS1,LOXL2,MARCKSL1,MB,MDK,MMP14,NFKB2,NR2F1,NR2F2,NRCAM,NTN1,PAK4,PIK3R3,PLCD3,PNPLA6,PRKCA,PRKCZ,PTN,RAB11A,RALB,RHOB,ROBO1,RSPO1,RXRA,SASH1,SERPINF1,SERPINF2,SFRP1,SH3KBP1,SIRT1,SOX11,STAT1,TGFBI,TGFBR1,THBS4,TNFAI | | | ADGRF5,AKT3,APLN,BMPER,C1GALT1,C3,CALCRL,CCL2,CCN1,CDH13,CDH5,CEACAM1,CITED2,COL3A1,CREB3L1,CSPG4,CUL7,CYP1B1,CYSLTR2,EMILIN1,ERG,ESM1,ETV2,FBN2,FGF10,FLI1,FOXC2,GADD45A,GPR4,HAND2,HAPLN1,HDAC9,HGF,HHEX,HIF3A,HLA-F,IL1A,ITGAV,ITGB1,ITGB1BP1,JAG1,KDR,KLF4,LDB2,LGALS1,LMO2,LOX,LOXL2,MEIS1,MIB1,NFATC4,PGF,PLK2,PRKCB,PROX1,RASIP1,RECK,RHOJ,RRAS,S1PR1,SDC2,SERPINF2,SMOC2,SOX11,SOX7,SRPX2,TAL1,TGFB2,THSD7A,TMEM100,TNN,TRPC1,TSPAN12,VAV3,ZAP70 | 4.02078 | 10 | 10 | 0 | 0 | 10 |
| 115 | GO: Biological Process | GO:0001568 | blood vessel development | ABCC8,ADAMTS18,ANGPT2,CCR3,CD40,CTSZ,CXCL1,CXCL12,CXCL8,EPHB1,HPSE,IL1A,IL1B,LIPA,LPA,MCAM,MIR10B,PGF,PSG4,PSG9,TDGF1,TDGF1P3,TINAGL1,TNF,TNFAIP3 | ADAM12,ANGPT1,APLNR,BMPR1A,COL1A1,COL1A2,COL3A1,COL5A1,CRB2,CREB3L1,CSPG4,CXXC5,CYP1B1,DCBLD2,DCN,E2F7,EDNRA,EGFR,EGR1,EMILIN1,EMILIN2,EPHB2,FAP,FGF1,FGF9,GDF6,GREM1,HAS2,HMGA2,KREMEN1,LEF1,LGALS1,LGALS3,LIF,LOX,LRP1,MMP19,MMP9,MT1E,MT1X,MT2A,MYOCD,NOTCH2,NRXN3,NTRK2,OSR1,PDGFRA,PDGFRB,PLCD3,PPARG,PRRX1,PRRX2,PTGER4,PTHLH,PTN,PTPRJ,RUNX1,SDC2,SEMA3A,SEMA3C,SERPINB7,SFRP2,SIX1,SMO,SOCS3,SPHK1,STRA6,TAFA5,TGFB2,TGFBI,THBS2,TNFRSF19,TNFSF4,VAV2,VSTM4,WNT5B | ABCA1,ABL1,ACTA2,ADGRG1,ADRA2B,ALDH1A2,AMOT,APOE,ATP2B4,BASP1,BMP7,BTG1,C3,CCL5,CHD7,CITED2,CLIC4,CNMD,COL1A1,COL3A1,CRB2,CRKL,CTNNB1,CXCL10,CXCR4,CYP26A1,DLX3,EFNB2,ENPP2,EPHB2,EPHB3,ERAP1,FBN2,FBXW11,FERMT2,FGFR2,FLCN,FOLR1,FZD5,FZD8,GAA,GLI3,GPC3,GREM1,GRN,HDAC5,HDAC9,HES1,HHEX,HK2,HS6ST2,HSPB1,ID1,IGFBP2,ITGAX,ITGB1BP1,ITGB8,JAG1,JCAD,JUNB,KLF4,LAMA1,LGALS1,LLGL2,LOXL1,LOXL2,MARCKSL1,MB,MDK,MMP14,NFKB2,NR2F1,NR2F2,NRCAM,NTN1,PAK4,PAX6,PIK3R3,PLCD3,PNPLA6,PRICKLE1,PRKCA,PRKCZ,PTH1R,PTN,RAB11A,RALB,RHOB,ROBO1,RSPO1,RXRA,SASH1,SERPINF1,SERPINF2,SFRP1,SH3KBP1,SIRT1,SNX17,SOX11,SPHK2,STAT1,SUFU,TGFBI,TGFBR1,THBS4,TNFAIP3,TNFRSF19,TNMD,TNNI3,TWSG1,VASH1,WNT4,WNT5B,YAP1 | | | ADAMTS6,ADGRF5,AKT3,APLN,BMP10,BMPER,C1GALT1,C3,CALCRL,CCL2,CCN1,CDH13,CDH5,CEACAM1,CFC1,CFC1B,CITED2,COL1A1,COL1A2,COL3A1,CREB3L1,CSPG4,CUL7,CYP1B1,CYSLTR2,EMILIN1,ERG,ESM1,ETV2,FBN2,FGF10,FLI1,FOXC2,GADD45A,GPR4,HAND2,HAPLN1,HDAC9,HGF,HHEX,HIF3A,HLA-F,IL1A,ITGAV,ITGB1,ITGB1BP1,JAG1,KDR,KLF4,LDB2,LGALS1,LMO2,LOX,LOXL2,LTBP1,MEIS1,MIB1,NFATC4,PGF,PLK2,PRKCB,PROX1,RASIP1,RECK,RHOJ,RRAS,S1PR1,SDC2,SERPINF2,SMOC2,SOX11,SOX7,SRPX2,TAL1,TGFB2,THSD7A,TMEM100,TNFSF4,TNN,TRPC1,TSPAN12,VAV3,ZAP70 | 3.67406 | 10 | 10 | 0 | 0 | 10 |
| 128 | GO: Biological Process | GO:0007156 | homophilic cell adhesion via plasma membrane adhesion molecules | CDH22,PCDHA10,PCDHA2,PCDHA3,PCDHGA1,PCDHGA2,PCDHGA5,PCDHGB1,PCDHGB2,PSG4,PSG9 | AMIGO2,CDH11,CDH6,CRB2,DSC3,FAT3,HMCN1,L1CAM,PALLD,PCDHGC3,SDK2,TENM3,TENM4 | CDH1,CDH10,CDH11,CDH24,CDH4,CDH6,CDH9,CLSTN1,CRB2,DCHS1,FAT1,FAT3,NRCAM,PALLD,PCDH17,PCDH18,PCDH19,PCDH7,PRTG,ROBO1,SDK2,TENM4 | CDHR1,CLSTN3,DCHS2,PCDHA3,PCDHA7,PCDHB12,PCDHB16,PCDHB18P,PCDHB3,PCDHB5,PCDHB7,PCDHB8,PCDHGA1,PCDHGA2,PCDHGA6,PCDHGB1,PCDHGB5 | CDHR1,CNTN2,DCHS2,L1CAM,PCDHA10,PCDHA3,PCDHA5,PCDHA7,PCDHA9,PCDHGA1,PCDHGA4,PCDHGA5,PCDHGA6,PCDHGA8,PCDHGA9,PCDHGB1,PCDHGB2,PCDHGB3,PCDHGB5,PKD1,PVR | CDH13,CDH17,CDH5,CEACAM1,DCHS1,DSCAM,FAT4,ITGB1,PALLD,PCDH10,PCDH7,PCDH9,PCDHGC3,PRTG,SDK2,TENM4 | 5.44017 | 3.071988 | 2.9510906 | 4.607038 | 5.6048177 | 3.4799982 |
| 178 | GO: Biological Process | GO:0016055 | Wnt signaling pathway | | ALPK2,APCDD1L,CD44,COL1A1,CTHRC1,DAAM2,DACT1,DKK1,DKK2,EDNRA,EGFR,EGR1,FERMT1,FGF9,FOXD1,FZD2,FZD7,GLI1,GREM1,HMGA2,JRK,KREMEN1,LEF1,LGR4,LRP1,LRP4,NANOG,NET1,NKD1,PLEKHA4,POU5F1,SDC1,SFRP2,SHISA2,SNAI2,SOX2,SOX9,TMEM198,WNT5B | ABL1,APC2,APOE,BMP2,CDH1,COL1A1,CSNK1D,CTNNB1,CTNNBIP1,CTNND2,DACT2,DDX3X,DLX3,DLX5,FBXW11,FERMT2,FGFR2,FGFR3,FOLR1,FRAT2,FRZB,FZD1,FZD2,FZD5,FZD8,GLI3,GPC3,GREM1,HESX1,HHEX,ID4,IGFBP2,IGFBP6,KLF4,LGR4,LMX1A,MDK,MLLT3,NKD1,PBXIP1,PIAS4,PLEKHA4,PRICKLE1,PYGO1,ROR2,RSPO1,SALL1,SENP2,SFRP1,SIX3,SOX2,SOX9,SP5,TBL1X,TIAM1,TLE3,TLE4,TMEM64,TMEM88,TNFAIP3,TOLLIP,WLS,WNT4,WNT5B,WNT8A,YAP1,ZEB2 | | | | 0 | 10 | 10 | 0 | 0 | 0 |
| 182 | GO: Biological Process | GO:0090090 | negative regulation of canonical Wnt signaling pathway | | CTHRC1,DACT1,DKK1,DKK2,EGR1,FERMT1,GLI1,GREM1,KREMEN1,LRP4,NANOG,NKD1,SFRP2,SNAI2,SOX2,SOX9,WNT5B | APC2,APOE,BMP2,CDH1,CTNNB1,CTNNBIP1,FRZB,FZD1,FZD5,GLI3,GPC3,GREM1,IGFBP2,IGFBP6,MDK,MLLT3,NKD1,PRICKLE1,ROR2,SFRP1,SOX2,SOX9,TLE3,TLE4,TMEM64,TMEM88,WNT5B,WNT8A | | | | 0 | 10 | 10 | 0 | 0 | 0 |

| | | | | | | | | | | | | | | | |
|---|---|---|---|---|---|---|---|---|---|---|---|---|---|---|---|
| 245 | GO: Biological Process | GO:0030111 | regulation of Wnt signaling pathway | | ALPK2,APCDD1L,COL1A1,CTHRC1,DAAM2,DACT1,DKK1,DKK2,EGFR,EGR1,FERMT1,FGF9,FZD7,GLI1,GREM1,HMGA2,JRK,KREMEN1,LEF1,LGR4,LRP1,LRP4,NANOG,NKD1,PLEKHA4,POU5F1,SFRP2,SHISA2,SNAI2,SOX2,SOX9,TMEM198,WNT5B | ABL1,APC2,APOE,BMP2,CDH1,COL1A1,CSNK1D,CTNNB1,CTNNBIP1,CTNND2,DACT2,DDX3X,DLX5,FGFR2,FGFR3,FOLR1,FRZB,FZD1,FZD5,GLI3,GPC3,GREM1,HHEX,ID4,IGFBP2,IGFBP6,LGR4,LMX1A,MDK,MLLT3,NKD1,PBXIP1,PLEKHA4,PRICKLE1,PYGO1,ROR2,RSPO1,SALL1,SENP2,SFRP1,SIX3,SOX2,SOX9,TBL1X,TIAM1,TLE3,TLE4,TMEM64,TMEM88,TNFAIP3,TOLLIP,WLS,WNT5B,WNT8A,YAP1,ZEB2 | | | | 0 | 10 | 10 | 0 | 0 | 0 |
| 256 | GO: Biological Process | GO:0060828 | regulation of canonical Wnt signaling pathway | | COL1A1,CTHRC1,DAAM2,DACT1,DKK1,DKK2,EGFR,EGR1,FERMT1,FGF9,FZD7,GLI1,GREM1,JRK,KREMEN1,LGR4,LRP4,NANOG,NKD1,PLEKHA4,POU5F1,SFRP2,SNAI2,SOX2,SOX9,TMEM198,WNT5B | APC2,APOE,BMP2,CDH1,COL1A1,CSNK1D,CTNNB1,CTNNBIP1,CTNND2,DDX3X,DLX5,FGFR2,FGFR3,FOLR1,FRZB,FZD1,FZD5,GLI3,GPC3,GREM1,HHEX,IGFBP2,IGFBP6,LGR4,LMX1A,MDK,MLLT3,NKD1,PLEKHA4,PRICKLE1,PYGO1,ROR2,RSPO1,SFRP1,SOX2,SOX9,TBL1X,TLE3,TLE4,TMEM64,TMEM88,WLS,WNT5B,WNT8A,YAP1,ZEB2 | | | | 0 | 10 | 10 | 0 | 0 | 0 |
| 257 | GO: Biological Process | GO:0060070 | canonical Wnt signaling pathway | | COL1A1,CTHRC1,DAAM2,DACT1,DKK1,DKK2,EDNRA,EGFR,EGR1,FERMT1,FGF9,FOXD1,FZD2,FZD7,GLI1,GREM1,JRK,KREMEN1,LEF1,LGR4,LRP4,NANOG,NET1,NKD1,PLEKHA4,POU5F1,SDC1,SFRP2,SNAI2,SOX2,SOX9,TMEM198,WNT5B | APC2,APOE,BMP2,CDH1,COL1A1,CSNK1D,CTNNB1,CTNNBIP1,CTNND2,DDX3X,DLX5,FGFR2,FGFR3,FOLR1,FRAT2,FRZB,FZD1,FZD2,FZD5,FZD8,GLI3,GPC3,GREM1,HESX1,HHEX,IGFBP2,IGFBP6,KLF4,LGR4,LMX1A,MDK,MLLT3,NKD1,PLEKHA4,PRICKLE1,PYGO1,ROR2,RSPO1,SFRP1,SOX2,SOX9,TBL1X,TLE3,TLE4,TMEM64,TMEM88,WLS,WNT4,WNT5B,WNT8A,YAP1,ZEB2 | | | | 0 | 10 | 10 | 0 | 0 | 0 |
| 288 | GO: Biological Process | GO:0098773 | skin epidermis development | | DKK1,EGFR,FERMT1,FST,GAL,GLI2,IGFBP5,LGR4,LRP4,MSX2,RUNX1,SMO,SOX21,SOX9,TFAP2C,TGFB2,TNFRSF19,TRPS1 | AP3B1,CDH1,CLDN4,CTNNB1,CTSV,DLX3,FGFR2,FLG,FST,GNAS,HRNR,IGFBP5,KLF4,KRT17,LGR4,LHX2,LLGL1,LLGL2,MSX2,PIAS4,SFN,SMAD4,SOX21,SOX9,STMN1,TNFRSF19 | | | | 0 | 10 | 5.7290462 | 0 | 0 | 0 |
| 289 | GO: Biological Process | GO:0030178 | negative regulation of Wnt signaling pathway | | ALPK2,APCDD1L,CTHRC1,DACT1,DKK1,DKK2,EGR1,FERMT1,FGF9,GLI1,GREM1,HMGA2,KREMEN1,LRP1,LRP4,NANOG,NKD1,SFRP2,SHISA2,SNAI2,SOX2,SOX9,WNT5B | APC2,APOE,BMP2,CDH1,CTNNB1,CTNNBIP1,FRZB,FZD1,FZD5,GLI3,GPC3,GREM1,IGFBP2,IGFBP6,MDK,MLLT3,NKD1,PRICKLE1,ROR2,SFRP1,SIX3,SOX2,SOX9,TLE3,TLE4,TMEM64,TMEM88,TOLLIP,WNT5B,WNT8A | | | | 0 | 10 | 5.7138345 | 0 | 0 | 0 |
| 315 | GO: Biological Process | GO:0030177 | positive regulation of Wnt signaling pathway | | COL1A1,DAAM2,DACT1,DKK1,DKK2,EGFR,FGF9,JRK,LEF1,LGR4,NKD1,PLEKHA4,POU5F1,SFRP2,TMEM198,WNT5B | ABL1,BMP2,COL1A1,CSNK1D,CTNNB1,DDX3X,DLX5,FGFR2,FGFR3,GPC3,HHEX,LGR4,LMX1A,MLLT3,NKD1,PBXIP1,PLEKHA4,ROR2,RSPO1,SALL1,SFRP1,TBL1X,TNFAIP3,WLS,WNT5B,WNT8A,YAP1,ZEB2 | | | | 0 | 5.224359 | 10 | 0 | 0 | 0 |
| 321 | GO: Biological Process | GO:0003158 | endothelium development | F2RL1,HPSE,ICAM1,IL1B,LIPA,PSG4,PSG9,TNF,ZDHHC21 | ADAMTS12,ATOH8,BMPR1A,EDNRA,EGFR,FGF1,FZD2,PDPN,POU3F2,S1PR3,SMO,TNFRSF19 | ATOH8,BTG1,CLDN3,CLIC4,CNMD,CTNNB1,CXCL10,CXCR4,F2RL1,FBN2,FZD1,FZD2,HHEX,ID1,ITGAX,JAG1,MARCKSL1,NR2F2,POU3F2,PTPRS,RDX,RHOB,SH3KBP1,SMAD4,TGFBR1,TJP3,TNFRSF19,TNMD,ZEB1 | | | ADAMTS12,CDH5,CEACAM1,CUL7,EDNRB,ETV2,FBN2,FOXC2,FZD2,HHEX,JAG1,KDR,MEIS1,MIB1,PDE4D,PLCB1,PROX1,S1PR1,TAL1,TMEM100 | 3.41102 | 2.09341 | 4.7426526 | 0 | 0 | 4.6929017 |
| 346 | GO: Biological Process | GO:1901342 | regulation of vasculature development | ABCC8,ANGPT2,CCR3,CD40,CXCL8,HPSE,IL1A,IL1B,LPA,PGF,PSG4,PSG9,TNF,TNFAIP3 | ADAM12,APLNR,CREB3L1,CYP1B1,DCN,EMILIN1,EMILIN2,FGF1,GREM1,HMGA2,LGALS3,LIF,LRP1,MMP9,PPARG,PTGER4,PTN,RUNX1,SFRP2,SPHK1,TAFA5,TGFB2,THBS2 | ABL1,AMOT,ATP2B4,BASP1,BTG1,C3,CCL5,CNMD,CTNNB1,CXCL10,CXCR4,ENPP2,ERAP1,FLCN,GREM1,GRN,HHEX,HK2,HSPB1,ID1,ITGAX,ITGB8,JCAD,KLF4,MDK,NTN1,PAK4,PRKCA,PTN,RHOB,SASH1,SERPINF1,SFRP1,SIRT1,SOX11,STAT1,THBS4,TNFAIP3,TNMD,TWSG1,VASH1,WNT4 | | | AKT3,BMPER,C3,CCL2,CCN1,CDH5,CEACAM1,CREB3L1,CYP1B1,CYSLTR2,EMILIN1,FOXC2,GADD45A,GPR4,HGF,HHEX,HLA-F,IL1A,ITGB1,KDR,KLF4,MEIS1,PGF,PLK2,PRKCB,RECK,RHOJ,RRAS,S1PR1,SMOC2,SOX11,TGFB2,TNN,TSPAN12 | 3.49692 | 2.821797 | 2.5900495 | 0 | 0 | 5.1742765 |
| 404 | GO: Biological Process | GO:0045446 | endothelial cell differentiation | F2RL1,HPSE,ICAM1,IL1B,LIPA,PSG4,PSG9,TNF,ZDHHC21 | | ATOH8,BTG1,CLDN3,CLIC4,CNMD,CTNNB1,CXCR4,F2RL1,FBN2,FZD1,FZD2,HHEX,ID1,JAG1,NR2F2,POU3F2,PTPRS,RDX,SMAD4,TGFBR1,TJP3,TNFRSF19,TNMD,ZEB1 | | | CDH5,CEACAM1,CUL7,EDNRB,ETV2,FBN2,FZD2,HHEX,JAG1,KDR,MEIS1,MIB1,PDE4D,PLCB1,PROX1,S1PR1,TAL1,TMEM100 | 3.90876 | 0 | 3.7949915 | 0 | 0 | 4.5649014 |
| 430 | GO: Biological Process | GO:0043542 | endothelial cell migration | ANGPT2,CD40,CXCL12,IGF1,LPA,MIR10B,PSG4,PSG9,TDGF1,TDGF1P3,TNF | | ABL1,AMOT,APOE,ATOH8,ATP2B4,CXCR4,EFNB2,GREM1,GRN,HDAC5,HDAC9,HSPB1,ID1,ITGB1BP1,JCAD,KLF4,LOXL2,NR2F2,PATZ1,PIK3R3,PRKCA,PRKCZ,PTN,RHOB,ROBO1,SASH1,SERPINF1,SIRT1,STAT1,TGFBR1,TMSB4X,VASH1,ZEB2 | | | AKT3,BMP10,BMPER,CDH13,CDH5,CEACAM1,CYP1B1,FOXC2,GADD45A,HDAC9,ITGAV,ITGB1,ITGB1BP1,KDR,KLF4,LMO2,LOXL2,PLK2,PROX1,RECK,RHOJ,RRAS,SEMA3D,SMOC2,SRPX2,THSD7A,TMSB4X,ZAP70 | 3.03133 | 0 | 2.4816535 | 0 | 0 | 5.131281 |
| 481 | GO: Biological Process | GO:0035567 | non-canonical Wnt signaling pathway | | CTHRC1,DAAM2,DACT1,DKK1,FZD2,FZD7,NKD1,PLEKHA4,SFRP2,WNT5B | ABL1,CSNK1D,FRZB,FZD1,FZD2,FZD5,FZD8,GPC3,MLLT3,NKD1,PLEKHA4,PRICKLE1,ROR2,RSPO1,SFRP1,TIAM1,WNT4,WNT5B | | | | 0 | 4.188631 | 5.683391 | 0 | 0 | 0 |
| 490 | GO: Biological Process | GO:1904018 | positive regulation of vasculature development | ANGPT2,CCR3,CD40,CXCL8,HPSE,IL1A,IL1B,PGF | ADAM12,APLNR,CYP1B1,EMILIN1,EMILIN2,FGF1,GREM1,HMGA2,LGALS3,MMP9,RUNX1,SFRP2,SPHK1 | | | | AKT3,BMPER,C3,CCN1,CDH5,CYP1B1,CYSLTR2,EMILIN1,FOXC2,HGF,IL1A,ITGB1,KDR,KLF4,MEIS1,PGF,PRKCB,RRAS,SMOC2,TNN | 2.63278 | 2.339608 | 0 | 0 | 0 | 4.3941047 |
| 498 | GO: Biological Process | GO:0001570 | vasculogenesis | CXCL1,CXCL12,PSG4,PSG9,TDGF1,TDGF1P3 | ANGPT1,APLNR,FGF1,FGF9,HAS2,MYOCD,NTRK2,PDGFRB,PTPRJ,SEMA3A,SMO | | | | CEACAM1,CITED2,CUL7,ETV2,FBN2,FLI1,HHEX,ITGAV,KDR,MIB1,RASIP1,RRAS,SOX7,TAL1,TMEM100,TRPC1 | 2.22944 | 2.665552 | 0 | 0 | 0 | 4.3008135 |
| 521 | GO: Biological Process | GO:2000050 | regulation of non-canonical Wnt signaling pathway | | DAAM2,DACT1,DKK1,NKD1,PLEKHA4,SFRP2,WNT5B | ABL1,CSNK1D,GPC3,MLLT3,NKD1,PLEKHA4,RSPO1,SFRP1,TIAM1,WNT5B | | | | 0 | 4.315363 | 4.2229832 | 0 | 0 | 0 |
| 528 | GO: Biological Process | GO:2000052 | positive regulation of non-canonical Wnt signaling | | DKK1,NKD1,PLEKHA4,WNT5B | ABL1,CSNK1D,GPC3,MLLT3,NKD1,PLEKHA4,RSPO1,SFRP1,WNT5B | | | | 0 | 2.696513 | 5.7227742 | 0 | 0 | 0 |
| 545 | GO: Biological Process | GO:0090263 | positive regulation of canonical Wnt signaling pathway | | COL1A1,DAAM2,DACT1,DKK2,EGFR,FGF9,JRK,LGR4,PLEKHA4,POU5F1,SFRP2,TMEM198 | COL1A1,CSNK1D,DDX3X,DLX5,FGFR2,FGFR3,GPC3,HHEX,LGR4,LMX1A,PLEKHA4,ROR2,RSPO1,SFRP1,TBL1X,WLS,WNT8A,YAP1,ZEB2 | | | | 0 | 4.153268 | 4.0121924 | 0 | 0 | 0 |
| 577 | GO: Biological Process | GO:0043534 | blood vessel endothelial cell migration | ANGPT2,CD40,CXCL12,IGF1,LPA,MIR10B,TDGF1,TDGF1P3,TNF | | | | | AKT3,CDH5,CYP1B1,FOXC2,GADD45A,HDAC9,ITGB1,ITGB1BP1,KDR,KLF4,LMO2,PLK2,RECK,RHOJ,SEMA3D,SRPX2,THSD7A,TMSB4X,ZAP70 | 3.39693 | 0 | 0 | 0 | 0 | 4.1588975 |
| 583 | GO: Biological Process | GO:0060071 | Wnt signaling pathway, planar cell polarity | | CTHRC1,DACT1,DKK1,FZD2,FZD7,NKD1,PLEKHA4,SFRP2 | ABL1,FZD1,FZD2,GPC3,MLLT3,NKD1,PLEKHA4,PRICKLE1,ROR2,SFRP1,TIAM1 | | | | 0 | 4.054132 | 3.4012688 | 0 | 0 | 0 |
| 655 | GO: Biological Process | GO:0045601 | regulation of endothelial cell differentiation | IL1B,PSG4,PSG9,TNF,ZDHHC21 | | | | | CDH5,CEACAM1,CUL7,ETV2,HHEX,JAG1,PLCB1,TAL1,TMEM100 | 3.09014 | 0 | 0 | 0 | 0 | 3.3740574 |
| 660 | GO: Biological Process | GO:2000095 | regulation of Wnt signaling pathway, planar cell polarity | | DACT1,DKK1,NKD1,PLEKHA4,SFRP2 | ABL1,GPC3,MLLT3,NKD1,PLEKHA4,SFRP1 | | | | 0 | 3.645657 | 2.7940525 | 0 | 0 | 0 |
| 685 | GO: Biological Process | GO:0010594 | regulation of endothelial cell migration | ANGPT2,CD40,IGF1,LPA,MIR10B,PSG4,PSG9,TDGF1,TDGF1P3,TNF | | | | | AKT3,BMP10,BMPER,CEACAM1,FOXC2,GADD45A,HDAC9,ITGB1BP1,KDR,KLF4,PLK2,PROX1,RECK,RHOJ,RRAS,SMOC2,SRPX2,TMSB4X,ZAP70 | 3.27783 | 0 | 0 | 0 | 0 | 2.8898286 |

| | | | | | | | | | | | | | | | |
|---|---|---|---|---|---|---|---|---|---|---|---|---|---|---|---|
| 719 | GO: Biological Process | GO:2000096 | positive regulation of Wnt signaling pathway, planar | | DKK1,NKD1,PLEKHA4 | ABL1,GPC3,MLLT3,NKD1,PLEKHA4 | | | | 0 | 2.484989 | 3.3426264 | 0 | 0 | 0 |
| 795 | GO: Biological Process | GO:0010595 | positive regulation of endothelial cell migration | CD40,IGF1,LPA,MIR10B,TDGF1,TDGF1P3 | | | | | AKT3,FOXC2,HDAC9,ITGB1BP1,KDR,PLK2,PROX1,RHOJ,RRAS,SMOC2,SRPX2,TMSB4X,ZAP70 | 2.29348 | 0 | 0 | 0 | 0 | 2.8746247 |
| 822 | GO: Biological Process | GO:0001974 | blood vessel remodeling | | AXL,EDNRA,ELN,LIF,LRP1,SEMA3C,TGFB2 | | | | AXL,CEACAM1,ERG,FGF10,FOXC2,JAG1,S1PR1,TGFB2,TMBIM1 | 0 | 2.2299 | 0 | 0 | 0 | 2.7561804 |
| 908 | GO: Biological Process | GO:0061028 | establishment of endothelial barrier | F2RL1,HPSE,ICAM1,IL1B,TNF,ZDHHC21 | | | | | | 4.44688 | 0 | 0 | 0 | 0 | 0 |
| 952 | GO: Biological Process | GO:1904994 | regulation of leukocyte adhesion to vascular | CXCL12,ICAM1,SELE,TNF,ZDHHC21 | | | | | | 4.15794 | 0 | 0 | 0 | 0 | 0 |
| 989 | GO: Biological Process | GO:0061042 | vascular wound healing | HPSE,MCAM,TNF,TNFAIP3 | | | | | | 4.0107 | 0 | 0 | 0 | 0 | 0 |
| 1143 | GO: Biological Process | GO:0061756 | leukocyte adhesion to vascular endothelial cell | CXCL12,ICAM1,SELE,TNF,ZDHHC21 | | | | | | 3.44494 | 0 | 0 | 0 | 0 | 0 |
| 1147 | GO: Biological Process | GO:0001885 | endothelial cell development | F2RL1,HPSE,ICAM1,IL1B,TNF,ZDHHC21 | | | | | | 3.42843 | 0 | 0 | 0 | 0 | 0 |
| 1199 | GO: Biological Process | GO:0010575 | positive regulation of vascular | HPSE,IL1A,IL1B,SULF2 | | | | | | 3.28801 | 0 | 0 | 0 | 0 | 0 |
| 1300 | GO: Biological Process | GO:0010573 | vascular endothelial growth factor production | HPSE,IL1A,IL1B,SULF2,TNF | | | | | | 3.06361 | 0 | 0 | 0 | 0 | 0 |
| 1307 | GO: Biological Process | GO:0048010 | vascular endothelial growth factor receptor signaling | | | | | | AXL,CCL2,EMILIN1,FGF10,FOXC2,HHEX,KDR,PGF,PRKCB | 0 | 0 | 0 | 0 | 0 | 3.0445166 |
| 1320 | GO: Biological Process | GO:1904997 | regulation of leukocyte adhesion | TNF,ZDHHC21 | | | | | | 3.02541 | 0 | 0 | 0 | 0 | 0 |
| 1384 | GO: Biological Process | GO:1903140 | regulation of establishment of | IL1B,TNF,ZDHHC21 | | | | | | 2.90099 | 0 | 0 | 0 | 0 | 0 |
| 1386 | GO: Biological Process | GO:1901550 | regulation of endothelial cell | IL1B,TNF,ZDHHC21 | | | | | | 2.90099 | 0 | 0 | 0 | 0 | 0 |
| 1393 | GO: Biological Process | GO:0001935 | endothelial cell proliferation | | | ALDH1A2,APOE,ATOH8,BMP2,CNMD,CYBA,DLK1,FGFR3,HMGB2,ITGB1BP1,ITGB8,JCAD,LOXL2,MDK,MMP14,NR2F1,NR2F2,PLA2G2A,PRKCA,SIRT1,STAT1,TGFBR1,THBS4,TNMD,VASH1,ZEB2 | | | | 0 | 0 | 2.8960027 | 0 | 0 | 0 |
| 1419 | GO: Biological Process | GO:0061757 | leukocyte adhesion to arterial | TNF,ZDHHC21 | | | | | | 2.85214 | 0 | 0 | 0 | 0 | 0 |
| 1574 | GO: Biological Process | GO:0061154 | endothelial tube morphogenesis | | | CTNNB1,CXCL10,CXCR4,ITGAX,MARCKSL1,RHOB,SH3KBP1 | | | | 0 | 0 | 2.6492897 | 0 | 0 | 0 |
| 1575 | GO: Biological Process | GO:0003159 | morphogenesis of an endothelium | | | CTNNB1,CXCL10,CXCR4,ITGAX,MARCKSL1,RHOB,SH3KBP1 | | | | 0 | 0 | 2.6492897 | 0 | 0 | 0 |
| 1748 | GO: Biological Process | GO:0008591 | regulation of Wnt signaling pathway, | | DKK1,WNT5B | | | | | 0 | 2.477216 | 0 | 0 | 0 | 0 |
| 1835 | GO: Biological Process | GO:1904953 | Wnt signaling pathway involved in midbrain | | | CSNK1D,CTNNB1,FZD1,LMX1A | | | | 0 | 0 | 2.351673 | 0 | 0 | 0 |
| 1838 | GO: Biological Process | GO:0061316 | canonical Wnt signaling pathway involved in heart | | | CTNNB1,PYGO1,TMEM88,WNT8A | | | | 0 | 0 | 2.351673 | 0 | 0 | 0 |
| 1941 | GO: Biological Process | GO:0007223 | Wnt signaling pathway, calcium | | DKK1,FZD2,WNT5B | | | | | 0 | 2.21635 | 0 | 0 | 0 | 0 |
| 1986 | GO: Biological | GO:0090244 | Wnt signaling | | DKK1,SFRP2 | | | | | 0 | 2.093145 | 0 | 0 | 0 | 0 |
| 1992 | GO: Biological Process | GO:0090081 | regulation of heart induction by | | DKK1,POU5F1 | | | | | 0 | 2.093145 | 0 | 0 | 0 | 0 |
| 1999 | GO: Biological | GO:0038031 | non-canonical Wnt | | FZD7,NKD1 | | | | | 0 | 2.093145 | 0 | 0 | 0 | 0 |
| 2207 | Mouse Phenotype | MP:0001614 | abnormal blood vessel morphology | | ADAMTS12,ANGPT1,APLNR,CACNA1H,CD44,COL1A1,COL3A1,COL5A1,CRB2,DCBLD2,DDR1,DLX1,DLX2,EDNRA,EGFR,ELN,EMILIN1,EPHA3,FBP2,FGF9,FGFRL1,FHOD3,FLNC,GDNF,GFPT2,HAS2,LEF1,LEFTY2,LIF,LOX,MAB21L2,MMP9,MYOCD,NOTCH2,OLFML2B,OSR1,PAX3,PDGFRA,PDGFRB,PDPN,PLAT,PPARG,PRRX1,PTGER4,PTHLH,PTPRJ,RARB,RGS2,RUNX1,RUNX2,S1PR3,SDC2,SEMA3C,SHISAL1,SIX1,SLC4A3,SLC8A1,SMO,SOCS3,SOX2,SPP1,STRA6,TGFB2,THBS2,TNFSF4,TUB,VAV2,VSTM4,ZIC2 | | | | ABRA,ADAMTS12,ADAMTS3,ADAMTS6,ADGRL4,AGBL1,APLN,BMP10,BMPER,C1GALT1,CALCRL,CCL2,CCN1,CDH13,CDH5,CFAP45,CFC1,CFC1B,CITED2,COL1A1,COL3A1,COMT,CTH,CUBN,CUL7,CYP2E1,DUSP3,EMILIN1,EPHA3,ERG,ETV2,FBN2,FBP2,FGD5,FGF10,FLI1,FOLH1,FOXC2,GPR4,GUCY1A1,HAND2,HGF,HHEX,HIF3A,IL1A,IL6ST,ITGAV,ITGB1,JAG1,KDR,KLF3,LAMC1,LMO2,LOX,LOXL2,LPL,LTBP1,LTBP4,MATR3,MEIS1,MIB1,PGF,PLEKHA1,PLVAP,PPARA,PRDM6,PRPH2,RASIP1,RECK,RHOJ,RRAS,S1PR1,SDC2,SEMA3D,SLC4A7,TAL1,TGFB2,TLR4,TMBIM1,TMEM100,TMSB4X,TNFSF4,TSPAN12,TTN,VAV3,VIM | 0 | 5.04957 | 0 | 0 | 0 | 10 |
| 2219 | Mouse Phenotype | MP:0000259 | abnormal vascular development | | ADAMTS12,ANGPT1,CD44,CRB2,DCBLD2,EDNRA,ELN,FGFRL1,FHOD3,FLNC,HAS2,LEF1,LEFTY2,MMP9,MYOCD,OSR1,PAX3,PDGFRA,PDGFRB,PLAT,PRRX1,PTGER4,PTPRJ,RUNX1,RUNX2,SDC2,SLC8A1,SMO,SPP1,TGFB2,THBS2 | | | | ABRA,ADAMTS12,ADAMTS3,ADGRL4,BMP10,C1GALT1,CCL2,CCN1,CDH5,CUBN,CUL7,DUSP3,ERG,ETV2,FLI1,FOLH1,FOXC2,GPR4,HAND2,HHEX,IL1A,ITGAV,JAG1,KDR,LMO2,LTBP1,MATR3,MEIS1,MIB1,PGF,PRDM6,RASIP1,RECK,RHOJ,RRAS,S1PR1,SDC2,TAL1,TGFB2,TLR4,TMEM100,TMSB4X,TTN,VAV3,VIM | 0 | 3.576524 | 0 | 0 | 0 | 10 |
| 2534 | Mouse Phenotype | MP:0003711 | pathological neovascularization | ACE2,ANGPT2,HPSE,ICAM1,LMX1B,LPA,PGF | | | | | | 3.89587 | 0 | 0 | 0 | 0 | 0 |
| 2671 | Mouse Phenotype | MP:0001634 | internal hemorrhage | | | | | | ADAMTS3,ADGRL4,C1GALT1,COL1A1,EPHA3,ERG,FGL2,FLI1,FOXC2,FSHR,GPR4,HGF,IL6ST,ITGAV,JAG1,LAMC1,LOX,PILRA,RECK,RHOJ,RUNX1T1,SEC23A,SVEP1,TFPI,TLR4,TMSB4X,TSPAN12 | 0 | 0 | 0 | 0 | 0 | 3.360282 |
| 2726 | Mouse Phenotype | MP:0006083 | abnormal blood vessel elastic tissue morphology | | COL1A1,COL3A1,ELN,EMILIN1,LOX | | | | | 0 | 3.194123 | 0 | 0 | 0 | 0 |
| 2761 | Mouse Phenotype | MP:0000249 | abnormal blood vessel physiology | ACE2,ANGPT2,CTSS,FAS,HPSE,ICAM1,IL1B,LMX1B,LPA,PGF,PSG4,PSG9 | | | | | | 3.07088 | 0 | 0 | 0 | 0 | 0 |
| 3048 | Pathway | M7253 | KEGG_FOCAL_ADHESION | | COL1A1,COL1A2,COL3A1,COL5A1,COL6A1,COL6A3,EGFR,FLNC,ITGA11,PAK3,PDGFRA,PDGFRB,SHC4,SPP1,THBS2,TNC,VAV2 | | | | AKT3,COL1A1,COL1A2,COL3A1,HGF,ITGA10,ITGA2,ITGA8,ITGAV,ITGB1,KDR,LAMC1,MAPK10,MYLK3,PGF,PIK3R5,PRKCB,SHC4,TNN,VAV3 | 0 | 4.614769 | 0 | 0 | 0 | 4.9998083 |

| | | | | | | | | | | | | | | | |
|---|---|---|---|---|---|---|---|---|---|---|---|---|---|---|---|
| 3093 | Pathway | M2890 | KEGG_CALCIUM_SIGNALING_PATHWAY | | | | | ADCY2,ADCY8,ADCY9,ATP2A3,CALML6,GNA14,GRM1,ITPR1,ITPR2,NTSR1,PDGFRB,PHKA1,PLCB2,PLCE1,PRKCG,RYR1,RYR2,TACR1 | | 0 | 0 | 0 | 0 | 4.8800228 | 0 |
| 3132 | Pathway | M39830 | WP_HIPPO_SIGNALING_REGULATION_PATHWAYS | | | CDH1,CDH10,CDH11,CDH24,CDH4,CDH6,CDH9,CTNNB1,FGFR2,FGFR3,GNAI2,GNAS,KIT,NF2,PRKCA,PRKCE,PRKCZ | | | | 0 | 0 | 4.0786336 | 0 | 0 | 0 |
| 3139 | Pathway | M2642 | KEGG_TGF_BETA_SIGNALING_PATHWAY | | BMPR1A,BMPR1B,CDKN2B,DCN,FST,GDF5,GDF6,LEFTY2,TGFB2,THBS2 | | | | | 0 | 4.027845 | 0 | 0 | 0 | 0 |
| 3153 | Pathway | M11355 | KEGG_TIGHT_JUNCTION | | | CASK,CLDN3,CLDN4,CLDN6,CRB3,CTNNA1,CTNNA2,CTNNB1,EPB41,GNAI1,GNAI2,IGSF5,LLGL1,LLGL2,PARD6G,PPP2R2B,PRKCA,PRKCE,PRKCZ,TJP3 | | | | 0 | 0 | 3.8353035 | 0 | 0 | 0 |
| 3188 | Pathway | M19428 | KEGG_WNT_SIGNALING_PATHWAY | | | APC2,CTNNB1,CTNNBIP1,FBXW11,FRAT2,FZD1,FZD2,FZD5,FZD8,MAPK10,NKD1,PPP2R5C,PRICKLE1,PRKCA,SENP2,SFRP1,SMAD4,TBL1X,WNT4,WNT5B,WNT8A | | | | 0 | 0 | 3.4674713 | 0 | 0 | 0 |
| 4526 | Gene Family | 20 | Clustered protocadherins | PCDHA10,PCDHA2,PCDHA3,PCDHGA1,PCDHGA2,PCDHGA5,PCDHGB1,PCDHGB2 | | | PCDHA3,PCDHA7,PCDHB12,PCDHB16,PCDHB17P,PCDHB18P,PCDHB3,PCDHB5,PCDHB7,PCDHB8,PCDHGA1,PCDHGA2,PCDHGA6,PCDHGB1,PCDHGB5 | PCDHA10,PCDHA3,PCDHA5,PCDHA7,PCDHA9,PCDHGA1,PCDHGA4,PCDHGA5,PCDHGA6,PCDHGA8,PCDHGA9,PCDHGB1,PCDHGB2,PCDHGB3,PCDHGB5,PCDHGB8P | | 10 | 0 | 0 | 10 | 10 | 0 |
| 5257 | Coexpression | M5891 | HALLMARK_HYPOXIA | | COL5A1,DCN,EGFR,GPC1,LOX,MT1E,MT2A,SDC2,STC2,TGFBI | BNIP3L,BTG1,CAVIN1,CITED2,CXCR4,FBP1,GAA,GPC1,GPC3,HK2,IER3,NCAN,PLIN2,PNRC1,PRKCA,PYGM,RRAGD,S100A4,SAP30,TGFBI,TNFAIP3,TPI1 | | | BGN,CCN1,CITED2,ENO3,ERO1A,IER3,KDELR3,KLHL24,LOX,PDK1,PGF,S100A4,SDC2,TMEM45A,TPST2 | 0 | 2.060036 | 3.9120565 | 0 | 0 | 3.3550968 |
| 5627 | Coexpression | M5895 | HALLMARK_WNT_BETA_CATENIN_SIG | | DKK1,LEF1,NKD1,WNT5B | CTNNB1,FZD1,FZD8,GNAI1,HDAC5,JAG1,NKD1,WNT5B | | | | 0 | 1.971718 | 3.2750273 | 0 | 0 | 0 |

**2. Functional enrichment network for Wnt and Hippo_DEGs**

| S.No. | Category | ID | Title (or Source) | PWB EC DEG - Hippo | PWB EC DEG - Wnt | PWB MSC DEG - Wnt | PWB MSC DEG - Hippo | PWB iPSC DEG - Hippo | PWB iPSC DEG - Wnt | PWB EC DEG - Hippo(-logP) | PWB EC DEG - Wnt(-logP) | PWB MSC DEG- Wnt(-logP) | PWB MSC DEG - Hippo(-logP) | PWB iPSC DEG - Hippo(-logP) | PWB iPSC DEG - Wnt(-logP) |
|---|---|---|---|---|---|---|---|---|---|---|---|---|---|---|---|
| 15 | GO: Molecular Function | GO:0005125 | cytokine activity | GDF5,GDF6,TGFB2,WNT5B | | | BMP6,TGFB2,WNT16,WNT3A,WNT9B | BMP2,BMP7,GDF5,WNT4,WNT5B,WNT8A,WNT8B | BMP2,CER1,GREM1,RPL13A,WNT4,WNT5B,WNT8A,WNT8B | 3.476183672 | 0 | 0 | 4.987997888 | 4.967078902 | 4.221863701 |
| 223 | GO: Biological Process | GO:0035239 | tube morphogenesis | BMPR1A,CRB2,DLG5,FGF1,FZD2,GDF6,GLI2,LEF1,NKD1,SOX2,TGFB2,WNT5B | CD44,CTHRC1,DACT1,EDNRA,EGFR,FGF9,FOXD1,FZD2,GLI1,GREM1,HMGA2,KREMEN1,LEF1,LGR4,LRP1,NKD1,SFRP2,SOX2,SOX9,TNFAIP3,WNT5B | CAV1,DLX3,EGFR,ETV2,EXT1,FGF10,FZD2,HHEX,HMGA2,JAG1,KLF4,KMT2D,NFATC4,PRICKLE1,PRKCB,RECK,SOSTDC1,SOX7,TNN,WNK1,WNT3A,WNT9B | DCHS1,FAT4,FZD2,PARD6G,PRICKLE1,SOX11,TGFB2,WNT3A,WNT9B | AMOT,BMP2,BMP7,CDH1,CRB2,CTNNB1,DCHS1,FBXW11,FZD1,FZD2,FZD5,FZD8,ID1,ID2,LLGL1,NKD1,NRCAM,PARD6G,PRICKLE1,PRKCZ,SMAD4,SOX11,SOX2,TCF7,TGFBR1,WNT4,WNT5B,WNT8B,YAP1 | ABL1,APOE,BMP2,CAV1,CDH1,CTNNB1,CTNNBIP1,DACT1,DLX3,DLX5,EGF,FBXW11,FERMT2,FGFR2,FGFR3,FOLR1,FZD1,FZD2,FZD5,FZD8,GLI3,GPC3,GREM1,HDAC5,HESX1,HHEX,IGFBP2,JAG1,KLF4,LGR4,MDK,NKD1,PKD1,PRICKLE1,PRKCA,RSPO1,SALL1,SERPINF1,SFRP1,SMAD4,SOX2,SOX9,TCF7,TERT,TNFAIP3,WNT4,WNT5B,WNT8B,YAP1,ZEB2 | 10 | 10 | 10 | 4.617811115 | 10 | 10 |
| 291 | GO: Biological Process | GO:0001944 | vasculature development | BMPR1A,CRB2,FGF1,GDF6,LEF1,TGFB2,WNT5B | COL1A1,EDNRA,EGFR,EGR1,FGF9,GPRC5B,GREM1,HMGA2,KREMEN1,LEF1,LRP1,SFRP2,TNFAIP3,WNT5B | CAV1,COL1A1,DLX3,EGFR,ETV2,FGF10,HHEX,HMGA2,JAG1,KLF4,KMT2D,NFATC4,PRICKLE1,PRKCB,RECK,SOX7,TNN,WNK1 | | AMOT,BMP7,CRB2,CTNNB1,FBXW11,FZD5,FZD8,ID1,LLGL2,NRCAM,PRICKLE1,PRKCZ,SOX11,TGFBR1,WNT4,WNT5B,YAP1 | ABL1,APOE,CAV1,COL1A1,CTNNB1,DLX3,EGF,FBXW11,FERMT2,FGFR2,FOLR1,FRZB,FZD5,FZD8,GLI3,GPC3,GREM1,HDAC5,HHEX,IGFBP2,JAG1,KLF4,MDK,PKD1,PRICKLE1,PRKCA,RSPO1,SERPINF1,SFRP1,TERT,TNFAIP3,WNT4,WNT5B,YAP1 | 3.042208699 | 5.54865372 | 10 | 0 | 10 | 10 |
| 406 | GO: Biological Process | GO:0098609 | cell-cell adhesion | CRB2,DLG5,GLI2,LEF1,SOX2,TGFB2 | | CAV1,EGFR,EXT1,JAG1,KLF4,PPP3CA,PRICKLE1,SOX13,WNK1,WNT3A | BMP6,DCHS1,FAT4,PRICKLE1,TGFB2,WNT3A | AP3B1,B2M,BMP2,BMP7,CDH1,CRB2,CTNNA1,CTNNA2,CTNNB1,DCHS1,NF2,NRCAM,PRICKLE1,PRKCZ,SOX2,WNT4 | ABL1,B2M,BMP2,CAV1,CDH1,CTNNB1,CTNND2,GLI3,IGFBP2,JAG1,KLF4,MDK,PKD1,PRICKLE1,PRKCA,PRKG1,SOX2,SOX9,TNFAIP3,WNT4 | 2.571286801 | 0 | 2.783959483 | 2.85355699 | 10 | 10 |
| 673 | GO: Biological Process | GO:0003158 | endothelium development | BMPR1A,FGF1,FZD2 | | EDNRB,EGFR,ETV2,FZD2,HHEX,JAG1,PLCB1 | | CTNNB1,FZD1,FZD2,ID1,SMAD4,TGFBR1 | CTNNB1,FZD1,FZD2,HHEX,JAG1,SMAD4 | 2.517124799 | 0 | 5.321524819 | 0 | 4.288710586 | 2.955538106 |
| 1109 | GO: Biological Process | GO:0001570 | vasculogenesis | | | CAV1,ETV2,HHEX,SOX7 | | AMOT,CTNNB1,YAP1 | CAV1,CTNNB1,FGFR2,HHEX,YAP1 | 0 | 0 | 2.815536238 | 0 | 1.89105044 | 2.771658665 |
| 1322 | GO: Biological Process | GO:0001569 | branching involved in blood vessel morphogenesis | | EDNRA,LEF1,SFRP2 | | | | ABL1,CTNNB1,MDK | 0 | 3.32681211 | 0 | 0 | 0 | 2.469292777 |
| 1799 | GO: Biological Process | GO:1905555 | positive regulation of blood vessel branching | | | | | | ABL1,MDK | 0 | 0 | 0 | 0 | 0 | 4.09943726 |
| 2271 | GO: Biological Process | GO:0001974 | blood vessel remodeling | | | EXT1,FGF10,JAG1 | | | | 0 | 0 | 2.599490949 | 0 | 0 | 0 |
| 2393 | GO: Biological Process | GO:0097746 | blood vessel diameter maintenance | | | CAV1,EDNRB,EGFR,EXT1 | | | | 0 | 0 | 2.458944811 | 0 | 0 | 0 |
| 4248 | Mouse Phenotype | MP:0001614 | abnormal blood vessel morphology | | | | | | APOE,CAV1,CDH1,COL1A1,DDX3X,DLX3,DLX5,FGFR2,FOLR1,FZD5,GLI3,GPC3,HHEX,JAG1,MDK,PKD1,ROR2,SERPINF1,SOX2,YAP1 | 0 | 0 | 0 | 0 | 0 | 2.800120572 |
| 4846 | Pathway | M7253 | KEGG_FOCAL_ADHESION | | | CAV1,COL1A1,EGFR,MAPK10,MAPK3,PRKCB,TNN | | | CAV1,COL1A1,CTNNB1,EGF,MAPK10,PRKCA,PRKCG | 0 | 0 | 4.8884092 | 0 | 0 | 3.150758645 |
| 8189 | Coexpression | M5930 | HALLMARK_EPITHELIAL_MESENCHYMAL_TRANSITION | | CD44,COL1A1,CTHRC1,DKK1,GREM1,LRP1,SDC1,SNAI2,TNFAIP3 | | | | COL1A1,FERMT2,FZD8,GREM1,IGFBP2,SFRP1,TNFAIP3 | 0 | 10 | 0 | 0 | 0 | 4.726502954 |
| 8193 | Coexpression | M5903 | HALLMARK_NOTCH_SIGNALING | | | | | FBXW11,FZD1,FZD5 | FBXW11,FZD1,FZD5,JAG1,PRKCA,PSEN2 | 0 | 0 | 0 | 0 | 4.244183454 | 10 |
| 8291 | Coexpression | M5902 | HALLMARK_APOPTOSIS | BIRC3,LEF1,TGFB2 | | | | | BMP2,CAV1,CTNNB1,IGFBP6,PLCB2,PSEN2 | 3.2217636 | 0 | 0 | 0 | 0 | 4.271426097 |
| 8341 | Coexpression | M5953 | HALLMARK_KRAS_SIGNALING_UP | | FGF9,GPRC5B,H2BC3,SOX9,TNFAIP3 | | | | BMP2,KLF4,SOX9,TNFAIP3 | 0 | 4.23718037 | 0 | 0 | 0 | 2.032737877 |
| 8354 | Coexpression | M5923 | HALLMARK_PI3K_AKT_MTOR_SIGNALING | | | EGFR,MAPK10,PLCB1,PRKCB | | | ITPR2,MAPK10,TIAM1 | 0 | 0 | 3.942244206 | 0 | 0 | 2.035383212 |
| 8402 | Coexpression | M5891 | HALLMARK_HYPOXIA | | | CAV1,EGFR,EXT1,GAPDH | | | CAV1,GPC3,PRKCA,TNFAIP3 | 0 | 0 | 2.883644 | 0 | 0 | 2.032737877 |
| 8444 | Coexpression | M5890 | HALLMARK_TNFA_SIGNALING_VIA_NFKB | | CD44,EGR1,TNFAIP3 | | | | BMP2,JAG1,KLF4,TNFAIP3 | 0 | 2.08631637 | 0 | 0 | 0 | 2.032737877 |
| 8467 | Coexpression | M5907 | HALLMARK_ESTROGEN_RESPONSE_LATE | | | | | | CAV1,CDH1,FGFR3,KLF4,MDK,TIAM1 | 0 | 0 | 0 | 0 | 0 | 3.75389724 |
| 8544 | Coexpression | M5937 | HALLMARK_GLYCOLYSIS | | CD44,EGFR,SDC1,SOX9 | | | | | 0 | 3.10949492 | 0 | 0 | 0 | 0 |
| 8708 | Coexpression | M5919 | HALLMARK_HEDGEHOG_SIGNALING | | | | | AMOT,NRCAM | | 0 | 0 | 0 | 0 | 2.494727902 | 0 |
| 8726 | Coexpression | M5892 | HALLMARK_CHOLESTEROL_HOMEOSTASIS | | | | | | CTNNB1,GNAI1,JAG1 | 0 | 0 | 0 | 0 | 0 | 2.456963808 |
| 8900 | Coexpression | M5913 | HALLMARK_INTERFERON_GAMMA_RESPONSE | | | | | | B2M,PSMA2,PSME1,TNFAIP3 | 0 | 0 | 0 | 0 | 0 | 2.032737877 |
| 769 | GO: Biological Process | GO:0045446 | endothelial cell differentiation | | | EDNRB,ETV2,FZD2,HHEX,JAG1,PLCB1 | | CTNNB1,FZD1,FZD2,ID1,SMAD4,TGFBR1 | CTNNB1,FZD1,FZD2,HHEX,JAG1,SMAD4 | 0 | 0 | 4.622744219 | 0 | 4.663604401 | 3.304390978 |

**3. Functional enrichment network for ZNF_DEGs**

| S.No. | Category | ID | Title (or Source) | PWB EC DEG - ZNF_GeneSet | PWB MSC DEG - ZNF_GeneSet | PWB iPSC DEG - ZNF_GeneSet | PWB EC DEG(-logP) | PWB MSC DEG(-logP) | PWB iPSC DEG(-logP) |
|---|---|---|---|---|---|---|---|---|---|
| 5 | GO: Molecular Function | GO:0003700 | DNA-binding transcription factor activity | EGR1,GLI1,GLI2,GLIS1,OSR1,PPARG,RARB,SNAI2,TRPS1,VDR,ZFHX4,ZFP41,ZIC1,ZIC2,ZIC5,ZNF208,ZNF239,ZNF248,ZNF257,ZNF300,ZNF37A,ZNF536,ZNF560,ZNF676,ZNF729,ZNF98,ZNF99,ZSCAN1 | BCL6B,GFI1,GLI4,KLF3,KLF4,KMT2D,MYT1,NR2C2,OVOL1,PLAG1,PPARA,PRDM4,PRDM5,SP4,ZBED3,ZBTB11,ZFP2,ZFP41,ZFP42,ZNF10,ZNF107,ZNF165,ZNF208,ZNF229,ZNF248,ZNF257,ZNF268,ZNF280B,ZNF37A,ZNF382,ZNF384,ZNF443,ZNF492,ZNF502,ZNF521,ZNF555,ZNF559,ZNF560,ZNF57,ZNF587B,ZNF596,ZNF605,ZNF624,ZNF649,ZNF664,ZNF680,ZNF681,ZNF682,ZNF723,ZNF729,ZNF736,ZNF737,ZNF740,ZNF75A,ZNF76,ZNF77,ZNF808,ZNF90,ZNF98,ZNF99,ZSCAN10 | BCL6B,CTCFL,DEAF1,FEZF1,FEZF2,GLI3,HIC2,HINFP,HOMEZ,KLF3,KLF4,LHX2,LHX5,LMX1A,MECOM,NR2F1,NR2F2,NR6A1,PATZ1,RORB,RXRA,SALL1,SALL3,SALL4,SP110,SP5,SP8,THRB,VDR,ZBED3,ZBED4,ZBTB10,ZBTB12,ZBTB22,ZBTB25,ZBTB37,ZBTB43,ZBTB46,ZBTB7B,ZEB1,ZEB2,ZFP2,ZFP62,ZFX,ZIC1,ZIC2,ZIC4,ZIC5,ZKSCAN1,ZNF107,ZNF112,ZNF138,ZNF141,ZNF20,ZNF208,ZNF219,ZNF221,ZNF226,ZNF239,ZNF248,ZNF257,ZNF266,ZNF280A,ZNF280B,ZNF316,ZNF331,ZNF33B,ZNF358,ZNF395,ZNF423,ZNF48,ZNF492,ZNF493,ZNF521,ZNF525,ZNF536,ZNF555,ZNF558,ZNF581,ZNF66,ZNF676,ZNF678,ZNF680,ZNF704,ZNF710,ZNF717,ZNF723,ZNF726,ZNF729,ZNF732,ZNF736,ZNF737,ZNF740,ZNF780B,ZNF808,ZNF83,ZNF841,ZNF891,ZNF90,ZNF98,ZNF99,ZSCAN1,ZSCAN18 | 10 | 10 | 10 |
| 13 | GO: Molecular Function | GO:0061630 | ubiquitin protein ligase activity | BIRC3,DTX3,PDZRN3,RNF157,RNF19A,UHRF1 | BRAP,CHFR,DCST1,MIB1,RFPL2,RNF152,RNF217,TRIM55,TRIM6,TRIM9 | MARCHF1,MGRN1,MKRN3,MSL2,PJA1,RNF11,RNF130,RNF144A,RNF152,RNF157,RNF5,TRIM13,TRIM23,TRIM24,TRIM4,TRIM52,TRIM56,TRIM72,TRIM74,TRIM9 | 3.39341222 | 4.388874389 | 10 |
| 19 | GO: Molecular Function | GO:0003682 | chromatin binding | | | BMI1,CHD4,CNBP,CTCFL,FEZF2,GLI3,HINFP,ING2,KLF3,KLF4,L3MBTL1,LHX2,MTA2,MTA3,NSD2,OBI1,PATZ1,PHF13,RXRA,SALL1,SALL4,THRB,TRIM24,ZEB1,ZFX,ZIC2 | 0 | 0 | 10 |
| 22 | GO: Molecular Function | GO:0042805 | actinin binding | LMO7,PDLIM3,PDLIM4,PPARG | | | 5.29151667 | 0 | 0 |
| 54 | GO: Cellular Component | GO:0000118 | histone deacetylase complex | | | CHD4,ING2,MECOM,MTA2,MTA3,SALL1 | 0 | 0 | 3.81650834 |
| 78 | Pathway | M734 | REACTOME_RNA_POLYMERASE_II_TRANSCRIPTION | GLI2,PPARG,RARB,VDR,ZNF135,ZNF208,ZNF248,ZNF257,ZNF300,ZNF37A,ZNF454,ZNF528,ZNF560,ZNF676,ZNF729,ZNF99 | KLF4,KMT2D,LMO2,NR0B1,NR2C2,PPARA,TAF15,ZFP2,ZNF10,ZNF208,ZNF248,ZNF257,ZNF26,ZNF268,ZNF320,ZNF354C,ZNF37A,ZNF382,ZNF443,ZNF454,ZNF480,ZNF492,ZNF521,ZNF528,ZNF555,ZNF559,ZNF560,ZNF596,ZNF605,ZNF624,ZNF649,ZNF664,ZNF680,ZNF681,ZNF682,ZNF701,ZNF729,ZNF736,ZNF737,ZNF740,ZNF75A,ZNF77,ZNF99 | ASH2L,BMI1,CHD4,GLI3,ING2,KLF4,L3MBTL1,LMO1,MDM4,MTA2,NR2F1,NR6A1,RORB,RXRA,THRB,VDR,ZFP2,ZKSCAN1,ZNF112,ZNF138,ZNF141,ZNF20,ZNF208,ZNF221,ZNF226,ZNF248,ZNF257,ZNF266,ZNF320,ZNF331,ZNF334,ZNF33B,ZNF415,ZNF419,ZNF470,ZNF471,ZNF483,ZNF492,ZNF493,ZNF521,ZNF528,ZNF555,ZNF558,ZNF607,ZNF665,ZNF676,ZNF678,ZNF680,ZNF701,ZNF704,ZNF706,ZNF710,ZNF717,ZNF726,ZNF729,ZNF732,ZNF736,ZNF737,ZNF740,ZNF761,ZNF860,ZNF99 | 10 | 10 | 10 |
| 149 | Gene Family | 71 | Nuclear hormone receptors | PPARG,RARB,VDR | | NR2F1,NR2F2,NR6A1,RORB,RXRA,THRB,VDR | 3.42416582 | 0 | 10 |
| 37 | GO: Biological Process | GO:0016567 | protein ubiquitination | | | BMI1,MARCHF1,MARCHF3,MARCHF9,MDM4,MGRN1,MKRN3,MSL2,OBI1,PJA1,PRICKLE1,RNF11,RNF130,RNF144A,RNF152,RNF157,RNF187,RNF5,TRAF4,TRIM13,TRIM23,TRIM24,TRIM4,TRIM52,TRIM56,TRIM72,TRIM74,TRIM9 | 0 | 0 | 10 |
| 43 | GO: Biological Process | GO:0048384 | retinoic acid receptor signaling pathway | PPARG,RARB,ZIC2,ZNF536 | | | 5.0869325 | 0 | 0 |

### Supplementary File 4: GO Functional Enrichments of DEGs

#### 1. Term_GO_IPSC_PWB 4221 vs control 52521

| | | Ont | N | Up | Down | P.Up | P.Down |
|---|---|---|---|---|---|---|---|
| GO:0048731 | system development | BP | 3853 | 155 | 451 | 1 | 8.21E-33 |
| GO:0048513 | animal organ development | BP | 2749 | 99 | 351 | 1 | 1.74E-31 |
| GO:0048856 | anatomical structure development | BP | 4699 | 183 | 512 | 1 | 4.72E-30 |
| GO:0007275 | multicellular organism development | BP | 4306 | 170 | 480 | 1 | 5.17E-30 |
| GO:0048869 | cellular developmental process | BP | 3394 | 122 | 401 | 1 | 3.49E-29 |
| GO:0032502 | developmental process | BP | 5011 | 194 | 532 | 1 | 1.41E-28 |
| GO:0030154 | cell differentiation | BP | 3226 | 118 | 384 | 1 | 2.33E-28 |
| GO:0007399 | nervous system development | BP | 2055 | 84 | 276 | 0.999928 | 1.70E-27 |
| GO:0009653 | anatomical structure morphogenesis | BP | 2218 | 85 | 285 | 0.999998 | 2.74E-25 |
| GO:0032501 | multicellular organismal process | BP | 5608 | 238 | 564 | 1 | 1.85E-24 |
| GO:0022008 | neurogenesis | BP | 1403 | 52 | 203 | 0.999931 | 8.65E-24 |
| GO:0048519 | negative regulation of biological process | BP | 4511 | 151 | 474 | 1 | 2.25E-23 |
| GO:0051239 | regulation of multicellular organismal process | BP | 2531 | 93 | 304 | 1 | 3.00E-22 |
| GO:0048699 | generation of neurons | BP | 1324 | 49 | 191 | 0.999895 | 3.42E-22 |
| GO:0050793 | regulation of developmental process | BP | 2139 | 74 | 268 | 1 | 5.28E-22 |
| GO:0007417 | central nervous system development | BP | 842 | 23 | 139 | 0.999995 | 1.97E-21 |
| GO:0048523 | negative regulation of cellular process | BP | 3989 | 135 | 422 | 1 | 1.03E-20 |
| GO:0030182 | neuron differentiation | BP | 1195 | 45 | 174 | 0.999665 | 1.26E-20 |
| GO:2000026 | regulation of multicellular organismal development | BP | 1709 | 63 | 223 | 0.99999 | 2.81E-20 |
| GO:0009888 | tissue development | BP | 1524 | 58 | 205 | 0.999923 | 3.93E-20 |
| GO:0050789 | regulation of biological process | BP | 9077 | 377 | 798 | 1 | 4.84E-20 |
| GO:0051094 | positive regulation of developmental process | BP | 1102 | 42 | 163 | 0.99928 | 5.17E-20 |
| GO:0065007 | biological regulation | BP | 9627 | 406 | 835 | 1 | 6.19E-20 |
| GO:0060322 | head development | BP | 665 | 15 | 116 | 0.999998 | 6.74E-20 |
| GO:0009605 | response to external stimulus | BP | 1750 | 54 | 224 | 1 | 2.50E-19 |
| GO:0007420 | brain development | BP | 627 | 15 | 110 | 0.999991 | 4.23E-19 |
| GO:0007155 | cell adhesion | BP | 1126 | 61 | 160 | 0.746397 | 6.15E-18 |
| GO:0022610 | biological adhesion | BP | 1129 | 61 | 160 | 0.753609 | 7.94E-18 |
| GO:0050896 | response to stimulus | BP | 6858 | 273 | 627 | 1 | 5.66E-17 |
| GO:0048518 | positive regulation of biological process | BP | 4969 | 166 | 484 | 1 | 7.58E-17 |
| GO:0008283 | cell proliferation | BP | 1535 | 43 | 196 | 1 | 9.54E-17 |
| GO:0040011 | locomotion | BP | 1455 | 55 | 188 | 0.999909 | 1.37E-16 |
| GO:0042221 | response to chemical | BP | 3470 | 128 | 362 | 1 | 3.07E-16 |
| GO:0048522 | positive regulation of cellular process | BP | 4441 | 148 | 440 | 1 | 3.10E-16 |
| GO:0045595 | regulation of cell differentiation | BP | 1513 | 53 | 192 | 0.999994 | 4.00E-16 |
| GO:0045597 | positive regulation of cell differentiation | BP | 791 | 32 | 121 | 0.991102 | 4.30E-16 |
| GO:0042127 | regulation of cell proliferation | BP | 1272 | 40 | 169 | 0.999998 | 4.62E-16 |
| GO:0030900 | forebrain development | BP | 319 | 6 | 67 | 0.999861 | 5.76E-16 |
| GO:0048468 | cell development | BP | 1807 | 71 | 218 | 0.999948 | 8.07E-16 |
| GO:0051707 | response to other organism | BP | 615 | 15 | 101 | 0.999985 | 1.30E-15 |
| GO:0006928 | movement of cell or subcellular component | BP | 1673 | 68 | 205 | 0.999715 | 1.42E-15 |
| GO:0005488 | binding | MF | 12299 | 521 | 991 | 1 | 1.42E-15 |
| GO:0043207 | response to external biotic stimulus | BP | 616 | 15 | 101 | 0.999986 | 1.45E-15 |
| GO:0050794 | regulation of cellular process | BP | 8526 | 349 | 740 | 1 | 1.48E-15 |
| GO:0051240 | positive regulation of multicellular organismal process | BP | 1373 | 52 | 177 | 0.999851 | 1.58E-15 |
| GO:0009887 | animal organ morphogenesis | BP | 857 | 36 | 126 | 0.987616 | 2.27E-15 |
| GO:0016477 | cell migration | BP | 1143 | 41 | 153 | 0.999858 | 7.40E-15 |
| GO:0035295 | tube development | BP | 846 | 34 | 123 | 0.993581 | 1.17E-14 |
| GO:0009607 | response to biotic stimulus | BP | 646 | 18 | 102 | 0.999925 | 1.32E-14 |
| GO:0060429 | epithelium development | BP | 930 | 28 | 131 | 0.999986 | 1.75E-14 |

| | | | | | | | |
|---|---|---|---|---|---|---|---|
| GO:0007154 | cell communication | BP | 4941 | 200 | 469 | 1 | 4.58E-14 |
| GO:0023052 | signaling | BP | 4919 | 202 | 467 | 1 | 5.28E-14 |
| GO:0000902 | cell morphogenesis | BP | 928 | 35 | 129 | 0.998559 | 7.69E-14 |
| GO:0051241 | negative regulation of multicellular organismal process | BP | 960 | 25 | 132 | 1 | 8.79E-14 |
| GO:0008150 | biological_process | BP | 13798 | 611 | 1075 | 1 | 9.56E-14 |
| GO:0051960 | regulation of nervous system development | BP | 812 | 27 | 117 | 0.999703 | 1.04E-13 |
| GO:0032989 | cellular component morphogenesis | BP | 1014 | 41 | 137 | 0.9964 | 1.11E-13 |
| GO:0098609 | cell-cell adhesion | BP | 658 | 45 | 101 | 0.148114 | 1.12E-13 |
| GO:0007165 | signal transduction | BP | 4517 | 179 | 434 | 1 | 1.24E-13 |
| GO:0005576 | extracellular region | CC | 3062 | 140 | 317 | 0.99967 | 1.57E-13 |
| GO:0044421 | extracellular region part | CC | 2589 | 108 | 277 | 0.99998 | 2.16E-13 |
| GO:0048870 | cell motility | BP | 1261 | 51 | 159 | 0.998667 | 3.62E-13 |
| GO:0051674 | localization of cell | BP | 1261 | 51 | 159 | 0.998667 | 3.62E-13 |
| GO:0010033 | response to organic substance | BP | 2618 | 81 | 278 | 1 | 4.76E-13 |
| GO:0050767 | regulation of neurogenesis | BP | 723 | 26 | 106 | 0.997932 | 5.42E-13 |
| GO:0023051 | regulation of signaling | BP | 2963 | 110 | 306 | 1 | 6.95E-13 |
| GO:0048585 | negative regulation of response to stimulus | BP | 1369 | 40 | 168 | 1 | 7.39E-13 |
| GO:0051093 | negative regulation of developmental process | BP | 776 | 26 | 111 | 0.99955 | 7.64E-13 |
| GO:0048583 | regulation of response to stimulus | BP | 3371 | 121 | 339 | 1 | 8.30E-13 |
| GO:0009966 | regulation of signal transduction | BP | 2606 | 91 | 275 | 1 | 1.40E-12 |
| GO:0010646 | regulation of cell communication | BP | 2923 | 108 | 301 | 1 | 1.68E-12 |
| GO:0060284 | regulation of cell development | BP | 811 | 28 | 113 | 0.999412 | 2.67E-12 |
| GO:0009892 | negative regulation of metabolic process | BP | 2573 | 71 | 271 | 1 | 2.70E-12 |
| GO:0010605 | negative regulation of macromolecule metabolic process | BP | 2373 | 63 | 254 | 1 | 2.94E-12 |
| GO:0021537 | telencephalon development | BP | 215 | 5 | 47 | 0.995814 | 3.05E-12 |
| GO:0048666 | neuron development | BP | 988 | 39 | 130 | 0.997453 | 3.45E-12 |
| GO:0035239 | tube morphogenesis | BP | 686 | 28 | 100 | 0.984743 | 3.65E-12 |
| GO:0022603 | regulation of anatomical structure morphogenesis | BP | 897 | 34 | 121 | 0.998141 | 3.93E-12 |
| GO:0045664 | regulation of neuron differentiation | BP | 593 | 23 | 90 | 0.988031 | 5.00E-12 |
| GO:0008285 | negative regulation of cell proliferation | BP | 556 | 24 | 86 | 0.954286 | 5.44E-12 |
| GO:0051704 | multi-organism process | BP | 1923 | 57 | 214 | 1 | 6.51E-12 |
| GO:0007423 | sensory organ development | BP | 447 | 12 | 74 | 0.999492 | 6.52E-12 |
| GO:0051172 | negative regulation of nitrogen compound metabolic process | BP | 2017 | 49 | 222 | 1 | 7.06E-12 |
| GO:0061564 | axon development | BP | 467 | 18 | 76 | 0.979343 | 7.97E-12 |
| GO:0031324 | negative regulation of cellular metabolic process | BP | 2160 | 58 | 234 | 1 | 8.26E-12 |
| GO:0045596 | negative regulation of cell differentiation | BP | 581 | 19 | 88 | 0.99855 | 9.75E-12 |
| GO:0002376 | immune system process | BP | 2155 | 64 | 233 | 1 | 1.13E-11 |
| GO:0030855 | epithelial cell differentiation | BP | 471 | 14 | 76 | 0.998781 | 1.22E-11 |
| GO:0005886 | plasma membrane | CC | 3917 | 210 | 377 | 0.928353 | 1.23E-11 |
| GO:0007166 | cell surface receptor signaling pathway | BP | 2347 | 84 | 249 | 1 | 1.33E-11 |
| GO:0048646 | anatomical structure formation involved in morphogenesis | BP | 883 | 31 | 118 | 0.999513 | 1.38E-11 |
| GO:0010648 | negative regulation of cell communication | BP | 1167 | 35 | 145 | 0.999999 | 1.39E-11 |
| GO:0045893 | positive regulation of transcription, DNA-templated | BP | 1255 | 27 | 153 | 1 | 1.53E-11 |
| GO:0023057 | negative regulation of signaling | BP | 1170 | 36 | 145 | 0.999998 | 1.68E-11 |
| GO:0005575 | cellular_component | CC | 14436 | 659 | 1104 | 1 | 1.84E-11 |
| GO:0005615 | extracellular space | CC | 2428 | 95 | 255 | 0.999998 | 2.05E-11 |
| GO:0072359 | circulatory system development | BP | 899 | 40 | 119 | 0.973868 | 2.10E-11 |
| GO:0071944 | cell periphery | CC | 4021 | 217 | 384 | 0.915621 | 2.11E-11 |
| GO:0003002 | regionalization | BP | 252 | 8 | 50 | 0.98175 | 2.74E-11 |
| GO:0051716 | cellular response to stimulus | BP | 5751 | 222 | 516 | 1 | 2.86E-11 |

| 2. Term_GO_MSC_PWB 4221 vs control 52521 | | Ont | N | Up | Down | P.Up | P.Down |
|---|---|---|---|---|---|---|---|
| GO:0048646 | anatomical structure formation involved in morphogenesis | BP | 944 | 33 | 91 | 0.647589 | 1.36E-13 |
| GO:0006396 | RNA processing | BP | 1017 | 86 | 12 | 3.02E-13 | 1 |
| GO:0072359 | circulatory system development | BP | 949 | 33 | 90 | 0.659523 | 4.81E-13 |
| GO:0048514 | blood vessel morphogenesis | BP | 513 | 15 | 57 | 0.852821 | 3.07E-11 |
| GO:0001568 | blood vessel development | BP | 587 | 20 | 62 | 0.671632 | 3.46E-11 |
| GO:0072358 | cardiovascular system development | BP | 623 | 21 | 64 | 0.692957 | 5.38E-11 |
| GO:0001944 | vasculature development | BP | 614 | 21 | 63 | 0.667607 | 8.08E-11 |
| GO:0001525 | angiogenesis | BP | 431 | 14 | 50 | 0.721867 | 1.14E-10 |
| GO:0031012 | extracellular matrix | CC | 412 | 15 | 48 | 0.554856 | 2.35E-10 |
| GO:0062023 | collagen-containing extracellular matrix | CC | 323 | 10 | 41 | 0.755125 | 3.78E-10 |
| GO:0071944 | cell periphery | CC | 4106 | 149 | 242 | 0.596061 | 5.63E-09 |
| GO:0009653 | anatomical structure morphogenesis | BP | 2299 | 75 | 153 | 0.886926 | 5.65E-09 |
| GO:0035239 | tube morphogenesis | BP | 731 | 23 | 66 | 0.810301 | 5.84E-09 |
| GO:0032501 | multicellular organismal process | BP | 5746 | 197 | 317 | 0.900892 | 8.22E-09 |
| GO:0005886 | plasma membrane | CC | 3999 | 144 | 236 | 0.635785 | 8.78E-09 |
| GO:0051239 | regulation of multicellular organismal process | BP | 2616 | 79 | 168 | 0.979543 | 1.08E-08 |
| GO:0007166 | cell surface receptor signaling pathway | BP | 2406 | 75 | 157 | 0.951953 | 1.31E-08 |
| GO:0048869 | cellular developmental process | BP | 3484 | 115 | 210 | 0.917944 | 1.66E-08 |
| GO:0016070 | RNA metabolic process | BP | 4212 | 216 | 148 | 2.09E-08 | 0.997616 |
| GO:0050793 | regulation of developmental process | BP | 2202 | 71 | 145 | 0.90107 | 3.14E-08 |
| GO:0005730 | nucleolus | CC | 1000 | 72 | 17 | 3.36E-08 | 0.999999 |
| GO:0006139 | nucleobase-containing compound metabolic process | BP | 5304 | 259 | 173 | 3.39E-08 | 0.999995 |
| GO:1901360 | organic cyclic compound metabolic process | BP | 5634 | 272 | 191 | 3.41E-08 | 0.99997 |
| GO:0023052 | signaling | BP | 5023 | 157 | 280 | 0.994688 | 5.00E-08 |
| GO:0046483 | heterocycle metabolic process | BP | 5427 | 263 | 182 | 5.05E-08 | 0.999977 |
| GO:0007165 | signal transduction | BP | 4613 | 142 | 261 | 0.995616 | 5.18E-08 |
| GO:0022613 | ribonucleoprotein complex biogenesis | BP | 482 | 43 | 2 | 8.19E-08 | 1 |
| GO:0006725 | cellular aromatic compound metabolic process | BP | 5457 | 263 | 185 | 8.63E-08 | 0.999956 |
| GO:0030198 | extracellular matrix organization | BP | 318 | 11 | 36 | 0.62642 | 8.98E-08 |
| GO:0030154 | cell differentiation | BP | 3314 | 109 | 198 | 0.918031 | 1.03E-07 |
| GO:0090304 | nucleic acid metabolic process | BP | 4692 | 232 | 157 | 1.11E-07 | 0.999902 |
| GO:2000026 | regulation of multicellular organismal development | BP | 1766 | 53 | 120 | 0.955407 | 1.20E-07 |
| GO:0000244 | spliceosomal tri-snRNP complex assembly | BP | 25 | 9 | 0 | 1.41E-07 | 1 |
| GO:0035295 | tube development | BP | 897 | 30 | 72 | 0.73421 | 1.44E-07 |
| GO:0005887 | integral component of plasma membrane | CC | 1218 | 63 | 90 | 0.003835 | 1.64E-07 |
| GO:0007154 | cell communication | BP | 5043 | 158 | 278 | 0.994254 | 1.73E-07 |
| GO:0034641 | cellular nitrogen compound metabolic process | BP | 5912 | 279 | 192 | 1.90E-07 | 0.999999 |
| GO:0038023 | signaling receptor activity | MF | 734 | 17 | 62 | 0.98706 | 1.94E-07 |
| GO:0060089 | molecular transducer activity | MF | 785 | 18 | 65 | 0.990419 | 2.00E-07 |
| GO:0048583 | regulation of response to stimulus | BP | 3462 | 114 | 203 | 0.921397 | 2.76E-07 |
| GO:0045595 | regulation of cell differentiation | BP | 1556 | 47 | 107 | 0.939013 | 3.69E-07 |
| GO:0048856 | anatomical structure development | BP | 4822 | 171 | 266 | 0.734088 | 3.77E-07 |
| GO:0031226 | intrinsic component of plasma membrane | CC | 1278 | 66 | 92 | 0.003218 | 3.80E-07 |
| GO:0002040 | sprouting angiogenesis | BP | 107 | 3 | 18 | 0.759146 | 5.59E-07 |
| GO:0032502 | developmental process | BP | 5137 | 181 | 279 | 0.774992 | 6.61E-07 |
| GO:0043062 | extracellular structure organization | BP | 360 | 17 | 37 | 0.176965 | 6.81E-07 |
| GO:0005576 | extracellular region | CC | 3130 | 105 | 185 | 0.869854 | 6.92E-07 |
| GO:0030199 | collagen fibril organization | BP | 49 | 1 | 12 | 0.841234 | 6.93E-07 |
| GO:1904018 | positive regulation of vasculature development | BP | 157 | 4 | 22 | 0.834367 | 8.52E-07 |
| GO:0007275 | multicellular organism development | BP | 4413 | 157 | 245 | 0.706987 | 9.03E-07 |
| GO:0000353 | formation of quadruple SL/U4/U5/U6 snRNP | BP | 11 | 6 | 0 | 9.59E-07 | 1 |
| GO:0045291 | mRNA trans splicing, SL addition | BP | 11 | 6 | 0 | 9.59E-07 | 1 |
| GO:0000365 | mRNA trans splicing, via spliceosome | BP | 11 | 6 | 0 | 9.59E-07 | 1 |
| GO:0010467 | gene expression | BP | 4896 | 235 | 175 | 1.08E-06 | 0.997856 |
| GO:0030239 | myofibril assembly | BP | 60 | 1 | 13 | 0.895041 | 1.09E-06 |
| GO:0005102 | signaling receptor binding | MF | 1164 | 38 | 84 | 0.804069 | 1.17E-06 |
| GO:0004888 | transmembrane signaling receptor activity | MF | 575 | 15 | 50 | 0.939584 | 1.33E-06 |
| GO:0000387 | spliceosomal snRNP assembly | BP | 50 | 11 | 0 | 1.56E-06 | 1 |
| GO:0051716 | cellular response to stimulus | BP | 5864 | 182 | 309 | 0.998457 | 1.90E-06 |
| GO:1901342 | regulation of vasculature development | BP | 273 | 4 | 30 | 0.991332 | 1.97E-06 |
| GO:0045765 | regulation of angiogenesis | BP | 245 | 4 | 28 | 0.981261 | 2.00E-06 |

| | | | | | | | |
|---|---|---|---|---|---|---|---|
| GO:0007155 | cell adhesion | BP | 1164 | 55 | 83 | 0.033653 | 2.18E-06 |
| GO:0044421 | extracellular region part | CC | 2630 | 93 | 158 | 0.68261 | 2.33E-06 |
| GO:0003008 | system process | BP | 1390 | 42 | 95 | 0.927623 | 2.38E-06 |
| GO:0022610 | biological adhesion | BP | 1169 | 56 | 83 | 0.025702 | 2.58E-06 |
| GO:0007507 | heart development | BP | 523 | 16 | 46 | 0.810018 | 2.62E-06 |
| GO:0005201 | extracellular matrix structural constituent | MF | 143 | 6 | 20 | 0.431257 | 2.74E-06 |
| GO:0032963 | collagen metabolic process | BP | 87 | 2 | 15 | 0.835166 | 3.55E-06 |
| GO:0007611 | learning or memory | BP | 214 | 3 | 25 | 0.986519 | 4.73E-06 |
| GO:0098797 | plasma membrane protein complex | CC | 470 | 20 | 42 | 0.283836 | 4.81E-06 |
| GO:0055002 | striated muscle cell development | BP | 137 | 2 | 19 | 0.963903 | 5.50E-06 |
| GO:0055001 | muscle cell development | BP | 150 | 3 | 20 | 0.917818 | 5.78E-06 |
| GO:0098802 | plasma membrane receptor complex | CC | 150 | 10 | 20 | 0.050731 | 5.78E-06 |
| GO:0051241 | negative regulation of multicellular organismal process | BP | 1012 | 35 | 73 | 0.675397 | 6.27E-06 |
| GO:0045766 | positive regulation of angiogenesis | BP | 140 | 4 | 19 | 0.762304 | 7.58E-06 |
| GO:0043235 | receptor complex | CC | 322 | 15 | 32 | 0.209442 | 7.76E-06 |
| GO:0010469 | regulation of signaling receptor activity | BP | 128 | 4 | 18 | 0.698169 | 7.92E-06 |
| GO:0044459 | plasma membrane part | CC | 2228 | 92 | 135 | 0.127386 | 1.01E-05 |
| GO:0031224 | intrinsic component of membrane | CC | 3943 | 139 | 217 | 0.736815 | 1.03E-05 |
| GO:0048584 | positive regulation of response to stimulus | BP | 1894 | 59 | 118 | 0.929073 | 1.17E-05 |
| GO:0048468 | cell development | BP | 1855 | 55 | 116 | 0.966913 | 1.19E-05 |
| GO:0009966 | regulation of signal transduction | BP | 2666 | 81 | 156 | 0.977708 | 1.23E-05 |
| GO:0097525 | spliceosomal snRNP complex | CC | 85 | 13 | 0 | 1.30E-05 | 1 |
| GO:1903670 | regulation of sprouting angiogenesis | BP | 64 | 1 | 12 | 0.909711 | 1.37E-05 |
| GO:0022603 | regulation of anatomical structure morphogenesis | BP | 944 | 22 | 68 | 0.993642 | 1.39E-05 |
| GO:0010927 | cellular component assembly involved in morphogenesis | BP | 97 | 1 | 15 | 0.973962 | 1.40E-05 |
| GO:1903672 | positive regulation of sprouting angiogenesis | BP | 36 | 1 | 9 | 0.741165 | 1.45E-05 |
| GO:0042221 | response to chemical | BP | 3540 | 114 | 197 | 0.95525 | 1.61E-05 |
| GO:0010646 | regulation of cell communication | BP | 2992 | 89 | 171 | 0.990848 | 1.64E-05 |
| GO:0090050 | positive regulation of cell migration involved in sprouting angiogen | BP | 21 | 1 | 7 | 0.545277 | 1.69E-05 |
| GO:0120114 | Sm-like protein family complex | CC | 101 | 14 | 0 | 1.94E-05 | 1 |
| GO:0045596 | negative regulation of cell differentiation | BP | 599 | 25 | 48 | 0.286559 | 1.95E-05 |
| GO:0070887 | cellular response to chemical stimulus | BP | 2649 | 94 | 154 | 0.669875 | 2.02E-05 |
| GO:0005615 | extracellular space | CC | 2465 | 89 | 145 | 0.59725 | 2.08E-05 |
| GO:0007612 | learning | BP | 125 | 2 | 17 | 0.947405 | 2.20E-05 |
| GO:0090049 | regulation of cell migration involved in sprouting angiogenesis | BP | 38 | 1 | 9 | 0.759908 | 2.33E-05 |
| GO:0055003 | cardiac myofibril assembly | BP | 22 | 0 | 7 | 1 | 2.38E-05 |
| GO:0023051 | regulation of signaling | BP | 3035 | 91 | 172 | 0.989226 | 2.49E-05 |
| GO:0048333 | mesodermal cell differentiation | BP | 30 | 3 | 8 | 0.09684 | 2.62E-05 |
| GO:0048731 | system development | BP | 3955 | 140 | 215 | 0.718622 | 2.65E-05 |

| **3. GO_EC_PWB 4221 vs Control52521** | | Ont | N | Up | Down | P.Up | P.Down |
|---|---|---|---|---|---|---|---|
| GO:0048731 | system development | BP | 4036 | 62 | 259 | 0.336967 | 8.15E-38 |
| GO:0007275 | multicellular organism development | BP | 4507 | 66 | 275 | 0.501371 | 7.85E-37 |
| GO:0048856 | anatomical structure development | BP | 4932 | 70 | 284 | 0.621633 | 1.13E-33 |
| GO:0032502 | developmental process | BP | 5262 | 75 | 290 | 0.609279 | 4.67E-31 |
| GO:0048513 | animal organ development | BP | 2907 | 43 | 197 | 0.480148 | 7.77E-30 |
| GO:0032501 | multicellular organismal process | BP | 5880 | 97 | 305 | 0.07549 | 2.57E-28 |
| GO:0009653 | anatomical structure morphogenesis | BP | 2363 | 34 | 168 | 0.557037 | 3.79E-27 |
| GO:0009888 | tissue development | BP | 1656 | 26 | 135 | 0.372255 | 6.07E-27 |
| GO:0030154 | cell differentiation | BP | 3400 | 49 | 210 | 0.556328 | 1.47E-26 |
| GO:0048869 | cellular developmental process | BP | 3570 | 52 | 216 | 0.522877 | 2.99E-26 |
| GO:2000026 | regulation of multicellular organismal development | BP | 1810 | 30 | 134 | 0.252245 | 9.31E-23 |
| GO:0050793 | regulation of developmental process | BP | 2267 | 36 | 154 | 0.313386 | 1.42E-22 |
| GO:0007399 | nervous system development | BP | 2114 | 27 | 147 | 0.793243 | 1.96E-22 |
| GO:0007165 | signal transduction | BP | 4732 | 71 | 248 | 0.405655 | 1.15E-21 |
| GO:0023052 | signaling | BP | 5143 | 81 | 261 | 0.218133 | 4.29E-21 |
| GO:0045595 | regulation of cell differentiation | BP | 1598 | 21 | 119 | 0.720701 | 2.79E-20 |
| GO:0001503 | ossification | BP | 345 | 3 | 50 | 0.881395 | 4.00E-20 |
| GO:0007154 | cell communication | BP | 5166 | 79 | 259 | 0.322513 | 4.32E-20 |
| GO:0031012 | extracellular matrix | CC | 443 | 11 | 56 | 0.060249 | 1.49E-19 |
| GO:0048468 | cell development | BP | 1891 | 30 | 130 | 0.336508 | 3.13E-19 |
| GO:0009887 | animal organ morphogenesis | BP | 932 | 13 | 84 | 0.603121 | 3.25E-19 |
| GO:0007166 | cell surface receptor signaling pathway | BP | 2480 | 46 | 155 | 0.048358 | 3.87E-19 |
| GO:0048699 | generation of neurons | BP | 1357 | 18 | 105 | 0.69392 | 4.99E-19 |
| GO:0022008 | neurogenesis | BP | 1445 | 18 | 109 | 0.787703 | 6.14E-19 |
| GO:0051239 | regulation of multicellular organismal process | BP | 2700 | 47 | 163 | 0.106851 | 9.94E-19 |
| GO:0062023 | collagen-containing extracellular matrix | CC | 350 | 9 | 48 | 0.07105 | 2.37E-18 |
| GO:0005886 | plasma membrane | CC | 4102 | 89 | 215 | 2.23E-05 | 5.12E-18 |
| GO:0030182 | neuron differentiation | BP | 1221 | 14 | 96 | 0.856259 | 7.60E-18 |
| GO:0071944 | cell periphery | CC | 4211 | 90 | 218 | 3.55E-05 | 1.12E-17 |
| GO:0051094 | positive regulation of developmental process | BP | 1190 | 24 | 94 | 0.065983 | 1.32E-17 |
| GO:0045596 | negative regulation of cell differentiation | BP | 625 | 13 | 64 | 0.126021 | 1.82E-17 |
| GO:0051093 | negative regulation of developmental process | BP | 837 | 14 | 75 | 0.335205 | 4.67E-17 |
| GO:0030198 | extracellular matrix organization | BP | 337 | 5 | 45 | 0.545462 | 8.06E-17 |
| GO:0035295 | tube development | BP | 923 | 18 | 79 | 0.127634 | 8.59E-17 |
| GO:0001501 | skeletal system development | BP | 446 | 4 | 52 | 0.892324 | 1.06E-16 |
| GO:0007155 | cell adhesion | BP | 1211 | 37 | 92 | 1.49E-05 | 3.66E-16 |
| GO:0022610 | biological adhesion | BP | 1216 | 38 | 92 | 6.73E-06 | 4.73E-16 |
| GO:0061448 | connective tissue development | BP | 241 | 2 | 37 | 0.868628 | 5.27E-16 |
| GO:0051716 | cellular response to stimulus | BP | 6009 | 79 | 274 | 0.881698 | 1.36E-15 |
| GO:0016477 | cell migration | BP | 1238 | 30 | 92 | 0.004212 | 1.44E-15 |
| GO:0040007 | growth | BP | 875 | 8 | 74 | 0.943763 | 1.74E-15 |
| GO:0043062 | extracellular structure organization | BP | 382 | 8 | 46 | 0.194769 | 1.99E-15 |
| GO:0040011 | locomotion | BP | 1561 | 33 | 106 | 0.019185 | 3.24E-15 |
| GO:0002062 | chondrocyte differentiation | BP | 112 | 1 | 25 | 0.807444 | 4.46E-15 |
| GO:0048666 | neuron development | BP | 1018 | 11 | 80 | 0.881296 | 6.61E-15 |
| GO:0050789 | regulation of biological process | BP | 9457 | 126 | 378 | 0.933931 | 1.71E-14 |
| GO:0001655 | urogenital system development | BP | 301 | 5 | 39 | 0.44639 | 2.70E-14 |
| GO:0044459 | plasma membrane part | CC | 2286 | 59 | 134 | 6.86E-06 | 3.85E-14 |
| GO:0072359 | circulatory system development | BP | 971 | 19 | 76 | 0.11719 | 4.24E-14 |
| GO:0051216 | cartilage development | BP | 182 | 2 | 30 | 0.745792 | 4.71E-14 |
| GO:0035239 | tube morphogenesis | BP | 755 | 16 | 65 | 0.086021 | 4.72E-14 |
| GO:0042127 | regulation of cell proliferation | BP | 1356 | 29 | 94 | 0.023969 | 4.82E-14 |
| GO:0005576 | extracellular region | CC | 3295 | 78 | 173 | 3.93E-06 | 5.29E-14 |
| GO:0051241 | negative regulation of multicellular organismal process | BP | 1059 | 20 | 80 | 0.140866 | 5.66E-14 |
| GO:0010646 | regulation of cell communication | BP | 3063 | 43 | 164 | 0.629015 | 6.12E-14 |
| GO:0048870 | cell motility | BP | 1365 | 32 | 94 | 0.005131 | 7.17E-14 |

| | | | | | | | |
|---|---|---|---|---|---|---|---|
| GO:0051674 | localization of cell | BP | 1365 | 32 | 94 | 0.005131 | 7.17E-14 |
| GO:0007167 | enzyme linked receptor protein signaling pathway | BP | 923 | 14 | 73 | 0.476275 | 8.54E-14 |
| GO:0051240 | positive regulation of multicellular organismal process | BP | 1482 | 33 | 99 | 0.0095 | 9.01E-14 |
| GO:0023051 | regulation of signaling | BP | 3105 | 46 | 165 | 0.473192 | 9.63E-14 |
| GO:0072001 | renal system development | BP | 269 | 5 | 36 | 0.354422 | 1.01E-13 |
| GO:0001822 | kidney development | BP | 255 | 5 | 35 | 0.314244 | 1.05E-13 |
| GO:0048523 | negative regulation of cellular process | BP | 4184 | 54 | 204 | 0.86255 | 1.70E-13 |
| GO:0001649 | osteoblast differentiation | BP | 192 | 2 | 30 | 0.771901 | 2.02E-13 |
| GO:0006928 | movement of cell or subcellular component | BP | 1785 | 34 | 111 | 0.062914 | 2.52E-13 |
| GO:0050896 | response to stimulus | BP | 7174 | 99 | 304 | 0.772981 | 2.61E-13 |
| GO:0050794 | regulation of cellular process | BP | 8859 | 117 | 356 | 0.938642 | 2.83E-13 |
| GO:0009966 | regulation of signal transduction | BP | 2736 | 38 | 149 | 0.650311 | 3.76E-13 |
| GO:0008283 | cell proliferation | BP | 1636 | 32 | 104 | 0.052605 | 4.39E-13 |
| GO:0072210 | metanephric nephron development | BP | 34 | 1 | 14 | 0.39287 | 4.46E-13 |
| GO:0048598 | embryonic morphogenesis | BP | 517 | 5 | 50 | 0.87527 | 6.75E-13 |
| GO:0048519 | negative regulation of biological process | BP | 4730 | 59 | 221 | 0.929836 | 7.81E-13 |
| GO:0045597 | positive regulation of cell differentiation | BP | 847 | 11 | 67 | 0.693512 | 9.97E-13 |
| GO:0048518 | positive regulation of biological process | BP | 5201 | 76 | 236 | 0.507569 | 1.73E-12 |
| GO:0048522 | positive regulation of cellular process | BP | 4632 | 64 | 216 | 0.708764 | 2.18E-12 |
| GO:0048589 | developmental growth | BP | 590 | 5 | 53 | 0.93398 | 2.42E-12 |
| GO:0060429 | epithelium development | BP | 991 | 13 | 73 | 0.690009 | 2.81E-12 |
| GO:0005102 | signaling receptor binding | MF | 1210 | 23 | 83 | 0.114833 | 3.27E-12 |
| GO:0031175 | neuron projection development | BP | 911 | 11 | 69 | 0.778718 | 3.35E-12 |
| GO:0005201 | extracellular matrix structural constituent | MF | 148 | 4 | 25 | 0.169859 | 3.57E-12 |
| GO:0065007 | biological regulation | BP | 10013 | 135 | 386 | 0.918988 | 3.78E-12 |
| GO:0060284 | regulation of cell development | BP | 837 | 13 | 65 | 0.445635 | 5.10E-12 |
| GO:0048729 | tissue morphogenesis | BP | 606 | 6 | 53 | 0.879039 | 6.78E-12 |
| GO:0030199 | collagen fibril organization | BP | 49 | 0 | 15 | 1 | 1.02E-11 |
| GO:0051960 | regulation of nervous system development | BP | 831 | 14 | 64 | 0.32563 | 1.08E-11 |
| GO:0048608 | reproductive structure development | BP | 380 | 7 | 40 | 0.317547 | 1.11E-11 |
| GO:0000902 | cell morphogenesis | BP | 960 | 12 | 70 | 0.746466 | 1.33E-11 |
| GO:0001656 | metanephros development | BP | 77 | 1 | 18 | 0.67744 | 1.37E-11 |
| GO:0009790 | embryo development | BP | 919 | 11 | 68 | 0.788088 | 1.40E-11 |
| GO:0061458 | reproductive system development | BP | 383 | 7 | 40 | 0.324553 | 1.41E-11 |
| GO:0048646 | anatomical structure formation involved in morphogene | BP | 963 | 17 | 70 | 0.239562 | 1.53E-11 |
| GO:0032989 | cellular component morphogenesis | BP | 1052 | 12 | 74 | 0.844817 | 1.78E-11 |
| GO:0070848 | response to growth factor | BP | 644 | 10 | 54 | 0.462995 | 2.14E-11 |
| GO:0061061 | muscle structure development | BP | 569 | 7 | 50 | 0.725915 | 2.34E-11 |
| GO:0120039 | plasma membrane bounded cell projection morphogene | BP | 617 | 7 | 52 | 0.798203 | 4.29E-11 |
| GO:0007267 | cell-cell signaling | BP | 1391 | 20 | 88 | 0.556793 | 5.08E-11 |
| GO:0048858 | cell projection morphogenesis | BP | 621 | 7 | 52 | 0.803517 | 5.43E-11 |
| GO:0060485 | mesenchyme development | BP | 253 | 7 | 31 | 0.077407 | 5.46E-11 |
| GO:0030155 | regulation of cell adhesion | BP | 585 | 12 | 50 | 0.147584 | 6.31E-11 |
| GO:0044421 | extracellular region part | CC | 2764 | 66 | 143 | 2.22E-05 | 6.35E-11 |
